# Stepwise GRN co-option in the evolution of a dipteran respiratory organ

**DOI:** 10.64898/2026.02.28.708779

**Authors:** Yaoming Yang, Xiaobo Zhang, Xiaodong Xu, Jiayao Ren, Lina Shi, Zhuoheng Jiang, Tianlong Cai, Ying Zhen

## Abstract

The origin of morphological novelty lies at the heart of evolutionary biology, yet how ecological shifts drive its emergence through rewiring of gene regulatory networks (GRNs) remains poorly understood. Here, we dissect the roles of GRN co-option in the development and evolution of the prothoracic respiratory organ (PROD), a key innovation facilitating aquatic respiration in dipteran pupae. Ancestral-state reconstruction across over 200 species reveals that PROD originated ∼241 Ma in the common ancestor of Diptera, with a major increase in structural complexity in schizophoran flies ∼66-70 Ma. These transitions coincide with the Carnian Pluvial Episode and the K-Pg mass extinction, periods of profound climatic and ecological upheaval that likely imposed selection pressure for efficient pupal gas exchange in flooded or submerged habitats. Using comparative transcriptomics, scRNA-seq, and genetic manipulations in *Drosophila melanogaster* and *Aedes albopictus*, we demonstrate that PROD is a prothoracic wing serial homolog, specified by a *cut-*centered spiracle GRN and arising through heterochronic activation of an appendage GRN. In schizophoran flies, increase in complexity of PROD involved additional co-option of a *breathless-*centered tracheal GRN. Our findings reveal a stepwise model of hierarchical GRN co-option and provide potential links between geo-climatic events to the origin and escalation of morphological novelty. This *eco-evo-devo* framework provides a mechanistic and generalizable paradigm for how ecological challenges are translated into morphological innovations over macroevolutionary timescales.

## Introduction

The origin of morphological novelty, particularly new anatomical structures without clear antecedents, represents a fundamental challenge in evolutionary biology. Pioneering work in evolutionary developmental biology (*evo-devo*) has revealed that many novel traits arise through the modification and co-option of existing developmental programs, often manifesting as serial homologs of ancestral appendages. For example, insect wings^1–8^, arthropod limbs^9–11^, and their derivatives such as beetle horns and treehopper helmets^12–14^, are built by redeploying deeply conserved gene regulatory networks (GRNs) governing appendage development. Similarly, wing pigmentation patterns illustrate how modular GRNs can be rewired to generate functional innovation^15,16^. Despite these advances, a critical gap remains: we lack a mechanistic understanding of how ecological shifts across macroevolutionary timescales drive the hierarchical co-option of GRNs to produce structurally and functionally novel organs.

We focus on the pupal prothoracic dorsal respiratory organ (PROD), a structure present in most dipteran lineages, including *Drosophila melanogaster*. PROD develops from the larval prothoracic (T1) dorsal imaginal disc and is characterized by a simple tubular morphology with snorkel-like extension, an adaptation associated with respiration in aquatic or semi-aquatic pupal habitat^17–19^. In *D. melanogaster*, the imaginal disc that gives rise to the PROD may be patterned during embryogenesis by the conserved morphogens *wingless* (*wg*), *decapentaplegic* (*Dpp*), and *Distal-less* (*Dll*) (Fig. 1a)^20–22^. Previous studies have reported that manipulation of *Hox* gene *Sex combs reduced* (*Scr*) induced wing-like outgrowths in T1^2,23,24^, while *Ultrabithorax* (*Ubx*) loss-of-function lead to ectopic PROD formation in the meso(T2)- and metathorax(T3)^1^. These observations suggest a shared developmental origin of PROD and thoracic appendages. Together with its morphological diversity and ecological significance in Diptera, PROD provides an opportunity to investigate how GRN modifications generate morphological novelty that facilitate niche adaptation.

**Fig. 1.**
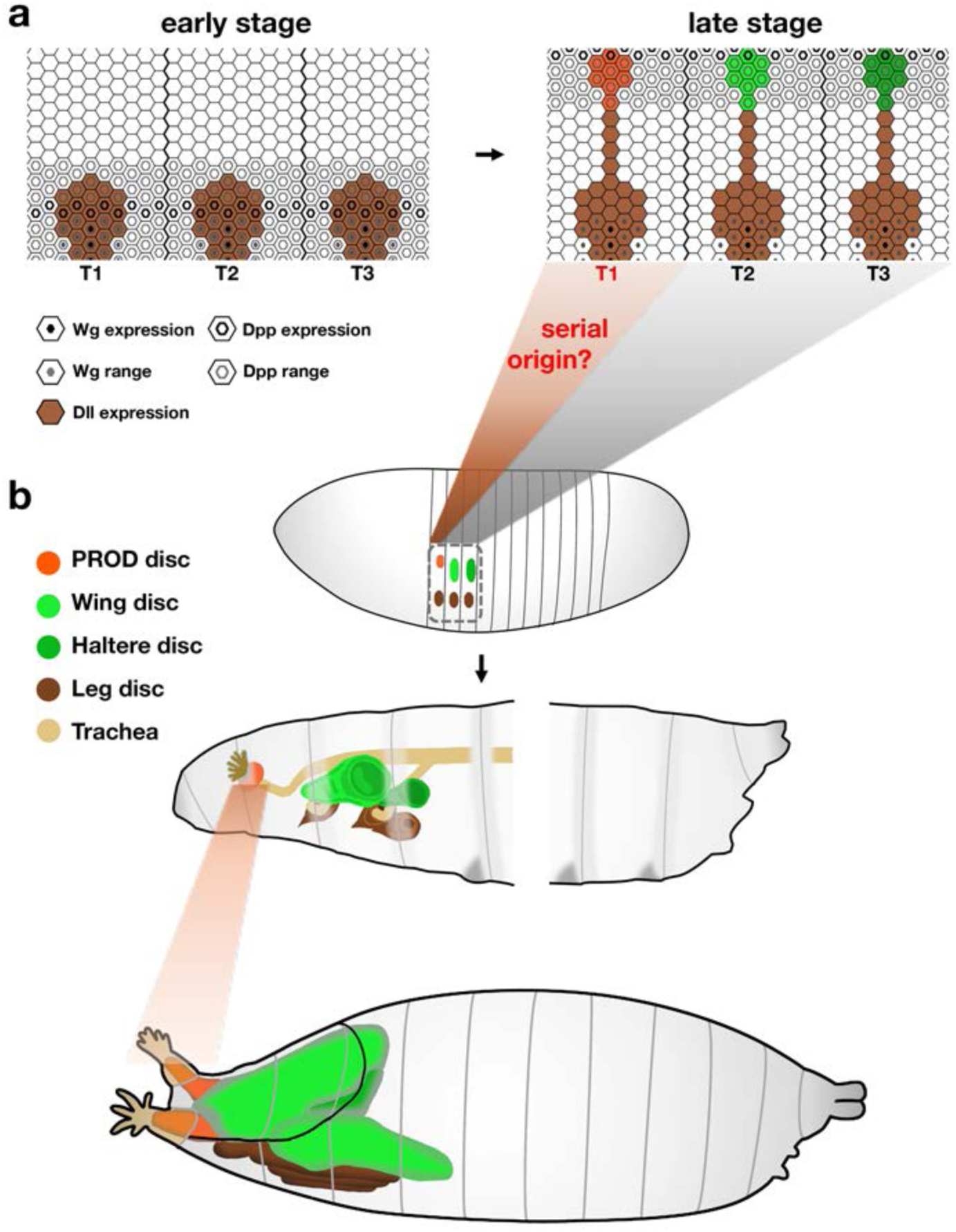
Fate map of thoracic discs in *D. melanogaster*. Images adapted from^20–22,61^. **(a)** The earliest embryonic thoracic positional cues in *D. melanogaster*—*wingless* (*wg*), *decapentaplegic* (*Dpp*), and *Distal-less* (*Dll*)—mediate cell fate determination of the wing, haltere, and, potentially, the pupal prothoracic dorsal respiratory organ (PROD). (b) Fate map of thoracic discs in *D. melanogaster*. Wing and haltere discs give rise to the wings, halteres, and thoracic notum; leg discs give rise to the legs; and the PROD disc gives rise to the anterior spiracle and humeral callus in the adult, and to the PROD at the pupal stage.

In this study, we integrate phylogenomics, comparative transcriptomics, single-cell RNA sequencing (scRNA-seq), and functional genetics across divergent dipteran lineages to investigate the developmental and evolutionary basis of PROD. We first reconstructed the phylogenetic origin of PROD, leveraging newly generated genomes of *Deuterophlebia wuyiensis* and *Philorus* sp., species in Dipteran basal lineages. We then identified conserved GRNs underlying PROD development across *De. wuyiensis*, *Aedes albopictus*, and *D. melanogaster* through comparative transcriptome analyses. Using scRNA-seq and cell lineage tracing, we further mapped cell-specific gene expression profiles and the developmental origin of the PROD disc of *D. melanogaster*. Finally, we performed gene manipulation experiments in both *A. albopictus* and *D. melanogaster* to dissect the functional roles of key regulators. Together, this integrative approach allows us to address three center questions: (1) When and in what ecological context did PRODs originated and modified during dipteran evolution? (2) Does PRODs share a developmental origin and GRN architecture with the wing, supporting its identity as a serial homolog? (3) How have stepwise modifications, including heterochrony and co-option of tracheal programs, shaped the increasing complexity of PROD in derived lineages? By bridging macroevolutionary patterns with developmental mechanisms, our work links ecological transitions to hierarchical co-option of GRNs in the origin and diversification of morphological novelty.

## Results

### A single origin of PROD followed by ecological diversification

To resolve the evolutionary origin of PROD, we first established a robust phylogeny of Diptera by integrating newly generated genomic data with a comprehensive set of publicly available sequences. Our analysis includes 235 species spanning 123 of approximately 157 extant dipteran families, covering 90 of the 95 species-rich families (>100 species), thereby resolving key uncertainties in dipteran phylogeny that have persisted across previous studies (Fig. 2a-c; Supplementary Fig. S1, Tables S1-S3). As part of this effort, we report high-quality, chromosome-level genome assemblies for *Deuterophlebia wuyiensis* (Deuterophlebiidae) and *Philorus* sp. (Blephariceridae), two rare basal lineages critical for rooting morphological and developmental innovations in the order. These genomic resources significantly improve representation of early-diverging Diptera and provide a solid foundation for accurate ancestral state reconstruction (ASR) of PROD across the radiation of true flies.

**Fig. 2.**
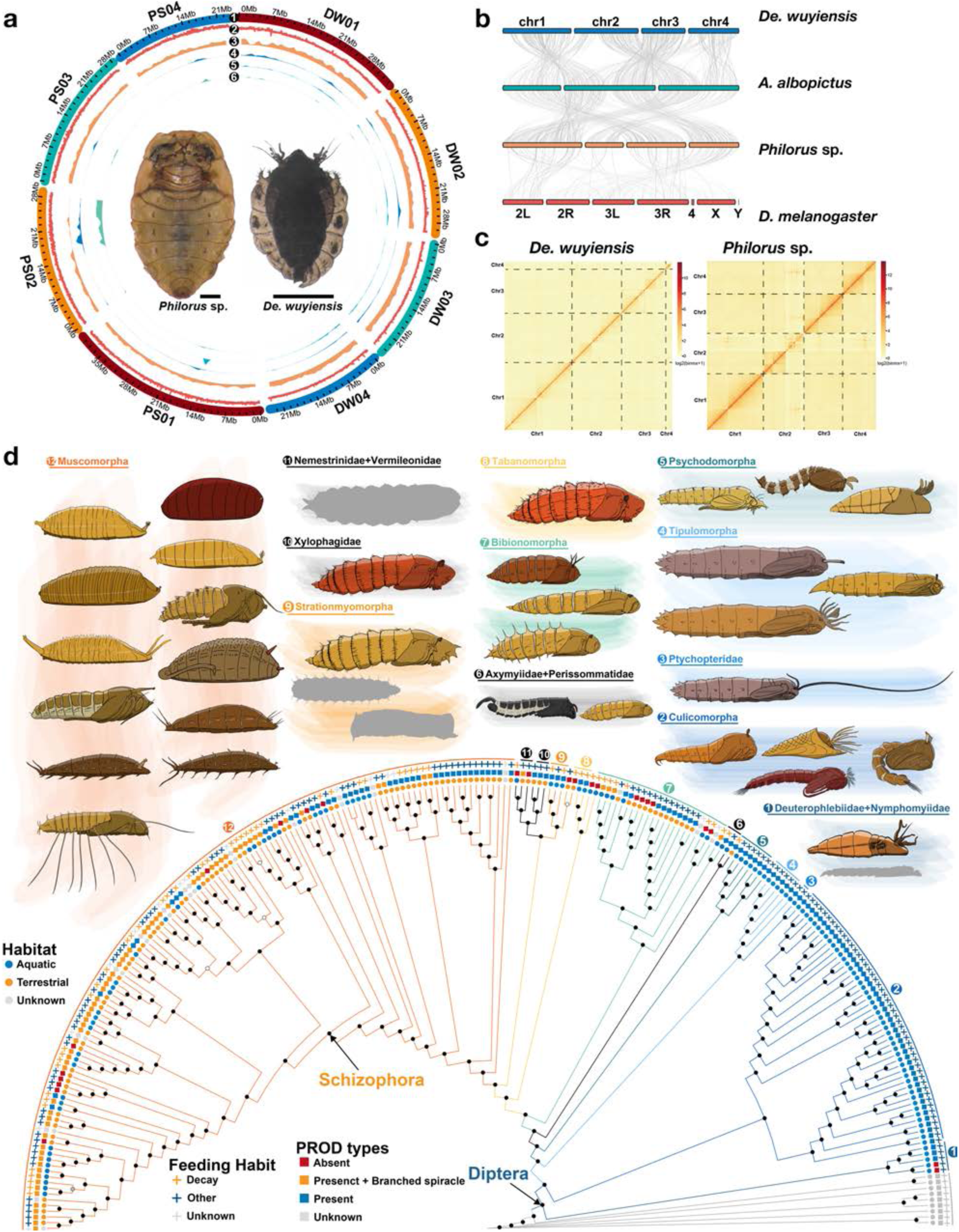
Phylogenomic framework for Diptera and diversity of PRODs. **(a)** Chromosome-level genome assemblies of *Deuterophlebia wuyiensis* and *Philorus* sp representing two key basal lineages. (Scale bars = 1.0 mm). From outer to inner track: chromosome length, GC content, gene density, SINE density, LINE density, and LTR density. **(b)** Macrosynteny among the genomes of *De. wuyiensis*, *A. albopictus*, *Philorus* sp., and *D. melanogaster*. **(c)** Genome-wide contact maps for *De. wuyiensis* and *Philorus* sp.; color indicates the log-transformed interaction strength between genomic bins. **(d)** Phylogenomic tree of Diptera based on nuclear gene sets, annotated with PROD morphology and ecology. Circles at internal nodes indicate branch support (SH-aLRT/UFBoot2), color-coded as black (≥98), grey (≥80), and white (<80). The outer band of plus-sign markers, the middle band of squares, and the inner band of circles denote feeding habitats, PROD types, and larval/pupal habitats, respectively. Group numbers at the tips correspond to the PROD illustrations in the upper panel.

To ensure robust phylogenetic inference, we constructed a high-quality gene matrix of 1,404 single-copy orthologs with at least 80% taxon occupancy across the 235 species, a level of completeness comparable to leading insect phylogenomic studies^25^ (Supplementary Fig. S2). We applied both maximum likelihood (ML) partitioning models and the coalescent-based ASTRAL framework to account for gene tree heterogeneity^26,27^. The resulting topology strongly supports the monophyly of all major dipteran infraorders, with high bootstrap support (> 95) across deep nodes. Notably, Deuterophlebiidae and Nymphomyiidae form a well-supported clade representing the earliest-diverging lineage within Diptera, resolving a long-standing ambiguity arising from their convergent morphology and conflicting placements in prior phylogenetic studies^28–32^. Our analysis further confirms the monophyly of Culicomorpha, Tipulomorpha, and Ptychopteridae (bootstrap > 95) (Fig. 2d), with Culicomorpha as the sister group to all other true flies, followed by (Tipulomorpha + Ptychopteridae). Within Brachycera, where previous phylogenies have shown weak resolution, we recover strongly supported relationships (bootstrap > 95). Together, this well-resolved phylogeny establishes a reliable foundation for accurate ASR of PROD evolution.

To reconstruct the evolutionary history of PROD, we compiled phenotypic data from extensive taxonomic literature and monographs, covering 109 of the 123 sampled families included in our phylogeny^33–55^ (Fig. 3; Supplementary Table S4). Based on morphological and histological evidence, we categorized PROD into three types: absent, present as a simple snorkel-like structure (e.g., in mosquito pupae), and modified with more complex tracheal-spiracular branching (e.g., in *D. melanogaster* pupae). Simple PROD are predominantly found in species in basal lineages with aquatic or semi-aquatic pupal stages^17,18^, while the branched form is typically associated with semi-terrestrial, saprophagous habitats.

**Fig. 3.**
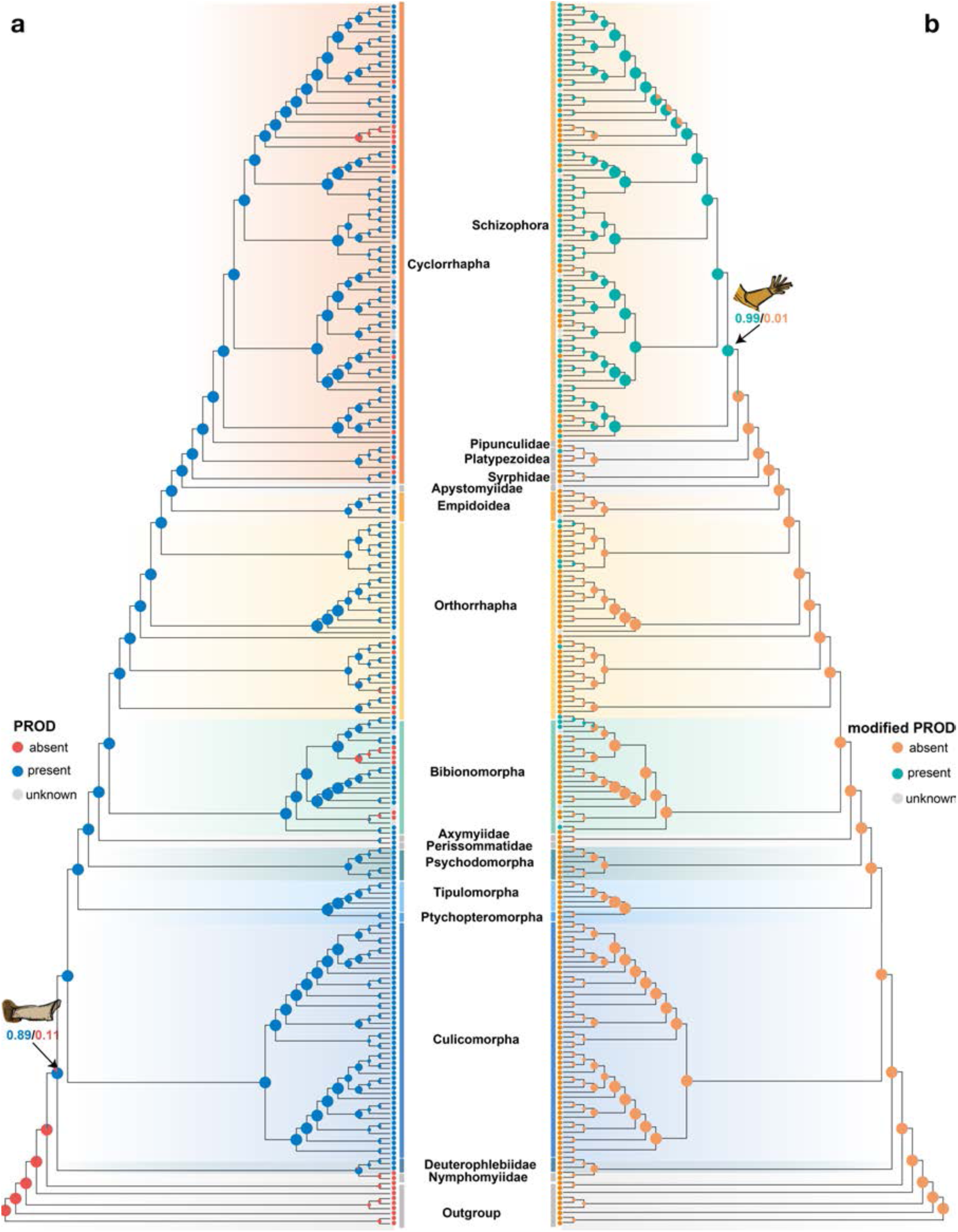
Ancestral state reconstruction of PROD across Diptera and outgroups using ER model. **(a)** Ancestral state reconstruction of PROD presence/absence across the dipteran phylogeny, showing a high inferred probability of a simple PROD in the common ancestor of Diptera. **(b)** Reconstructed probability of a modified PROD with branched spiracles, indicating multiple inferred gains across Diptera, with a particularly pronounced increase at the base of Schizophora.

ASR under our time-calibrated phylogeny strongly supports a single origin of PROD at the most recent common ancestor of Diptera (posterior probability = 0.86; Fig. 3a, 4a), at approximately 241 million years ago (95% HPD: 230-260 Ma). The complex, branched form of PROD evolved multiple times independently, most notably within the Schizophora clade ∼71 Ma (posterior probability = 0.99; 95% HPD: 64-76 Ma; Fig. 3b,4a; Supplementary Figs. S3-S5). The origin of PROD overlapped with the Carnian Pluvial Episode, a period of intense global rainfall and widespread flooding. Basal nematoceran lineages exhibit diverse aquatic-adapted PROD morphologies, consistent with selection for efficient respiration in submerged environments^17,18^. The emergence of complex, branched form of PROD in Schizophora coincided with the K-Pg mass extinction (∼66 Ma) and a major radiation of this detritivorous fly lineage into oxygen-poor decaying substrates, where branched spiracles may reduce the risk of suffocation.

**Fig. 4.**
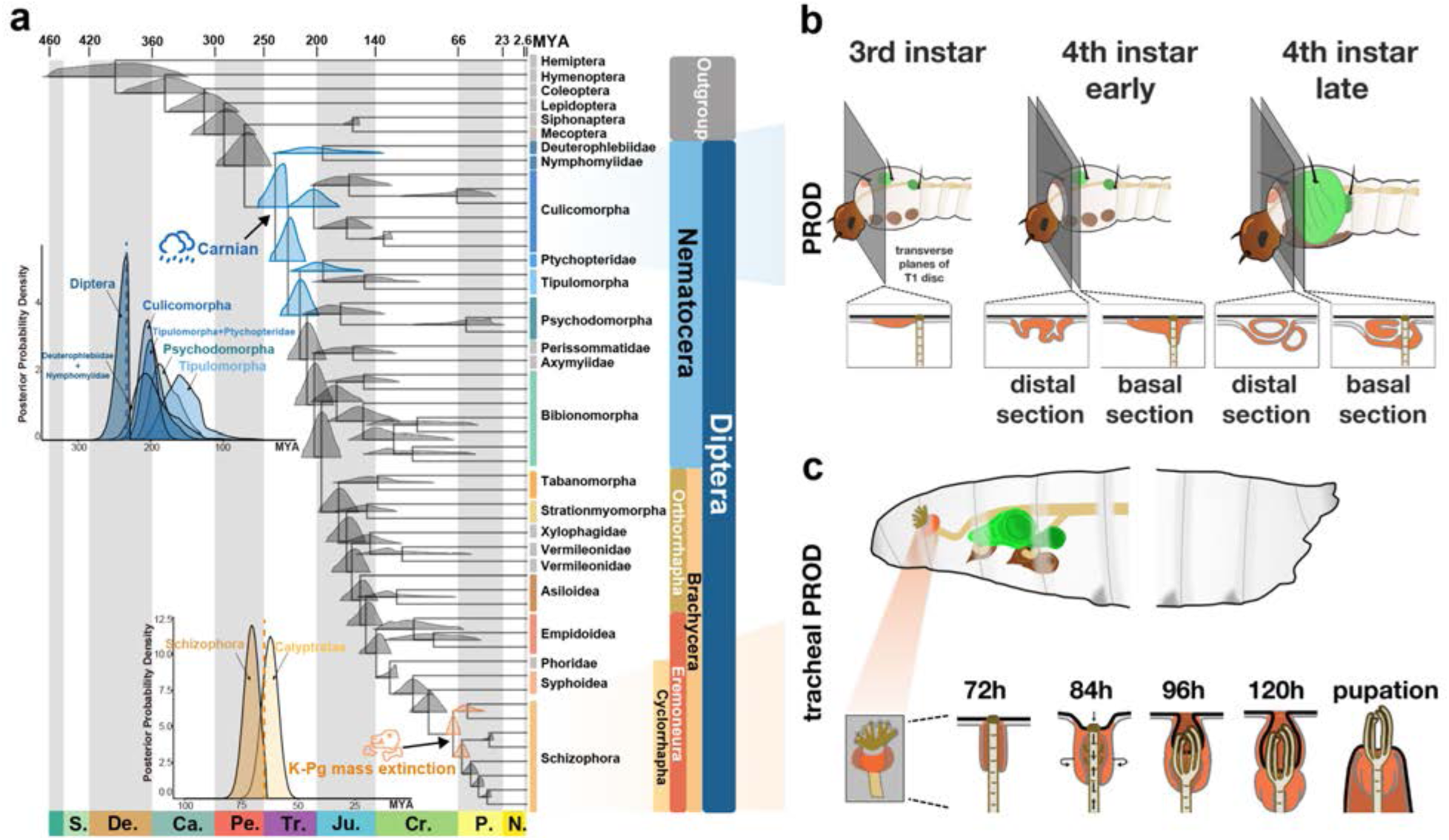
Divergence times and conserved morphogenesis of dipteran PRODs. **(a)** Fossil-calibrated divergence times for Diptera inferred using skew-normal relaxed clock models, based on 47 representative dipteran species and outgroups. Posterior age estimates for key clades are shown as density curves. The timing of two major geological events—the Carnian Pluvial Episode and the Cretaceous-Paleogene (K-Pg) mass extinction—is indicated by blue and orange color. **(b)** Schematic of PROD morphogenesis in *A. albopictus*. A single-layered epithelial PROD disc (orange, transverse section of 3rd instar) folds and elongates along the proximal-distal axis to form a snorkel-like respiratory organ in 4th instar larvae (orange, transverse sections). A subset of disc cells contributes to the surrounding “felt-chamber” structure (not highlighted), which encases the spiracular branch without generating additional tracheal branches. **(c)** Schematic of PROD morphogenesis in *D. melanogaster*. A single-layered epithelial PROD disc folds, interacts with the spiracular tracheal branch (grey), and subsequently everts to form a cone-like respiratory organ with associated tracheal-spiracular branches in the pupa.

Together, these analyses support a model of PROD evolution characterized by a single origin in the dipteran ancestor, followed by repeated structural modification potentially in response to shifting ecological pressures. While temporal congruence between trait evolution and environmental upheaval does not establish causation, such correlations suggest with that ecological opportunity may drive morphological innovation^17,18,56–59^. Rather than representing convergent gains of a similar organ, the distribution of PROD across Diptera is best explained by deep homology and stepwise modification, providing a foundation for investigating the developmental and genetic mechanisms underlying its diversification.

### Conserved morphogenesis processes in PROD development

To evaluate the hypothesis of a single origin of dipteran PROD and to identify conserved developmental mechanisms, we characterized the morphogenesis of the PROD disc in *D. melanogaster* and *A. albopictus* (Fig. 4b, c; Supplementary Fig. S6). In both species, PROD develops from the prothoracic (T1) dorsal imaginal disc, which undergoes epidermal invagination to form a two-layered sac that ultimately gives rise to the adult prothoracic spiracle, a process well documented in *D. melanogaster*^21,60,61^ and some nematoceran species^17–19^. At early stages (72 hours after egg laying [AEL] in *D. melanogaster*), the T1 disc is morphologically similar to other thoracic imaginal discs (Fig. 4b; Supplementary Fig. S6a). In most nematocerans, a subset of T1 dorsal imaginal cells migrates along the T1 spiracular trachea toward the dorsal tracheal trunk early in the final instar, whereas the remaining single-layered cells protrude distally, akin to wing morphogenesis. In *A. albopictus*, this protrusion further tubulates along the proximal-distal (P/D) axis to form a trachea-independent snorkel (Fig. 4c).

In *D. melanogaster* larvae, by 84 h AEL, comparable PROD imaginal cells migrate along the T1 spiracular trachea, envelop the anterior dorsal tracheal trunk, and initiate the formation of pupal tracheal-spiracular branches between 84 and 96 h AEL (Supplementary Fig. S6b). This epithelial-tracheal interaction is a prerequisite for eversion of the pupal distal spiracular structure along the P/D axis, during the transition to the 108-120 h AEL wandering larvae stage and pupation. In contrast, the wing disc remains drop-shaped and only enlarges in size throughout the third instar larvae, with distinct features such as the distal wing blade and veins appearing only after pupation. Notably, the part of wing disc that give rise to proximal region of the adult wing interacts with a non-spiracular trachea branch to mediate the formation of the adult air-sac^62^. Similar tracheal engagement is observed in abdominal imaginal cells which interact with the abdominal tracheal system to form adult abdominal spiracle and surrounding body wall^63–65^. These suggest that local epithelial-tracheal coordination in *D. melanogaster* PROD is a recurring developmental module deployed across multiple body regions including the wing disc and abdominal imaginal cells.

The conserved morphogenesis of dipteran PROD and shared developmental outcomes (pupal protrusion and adult prothoracic spiracle) support its origin from a common developmental program in the dipteran ancestor. However, the recruitment of the spiracular tracheal branch at the pupal stage appears to be a derived trait restricted to Brachycera such as *D. melanogaster* and absent in mosquitoes and most nematoceran lineages^17–19^. This distinction highlights a key evolutionary transition: while the basic PROD architecture is ancestral, its integration with the tracheal system has been co-opted and modified in more derived clades. Together, these conserved and divergent features during morphogenesis also identify informative developmental time windows (e.g., 84 h AEL in *D. melanogaster*) for tissue-specific RNA-seq of PROD, enabling the dissection of GRNs underlying both the conserved and lineage-specific components in PROD development.

### Transcriptomic profiling reveals shared and divergent GRNs in PROD evolution

To dissect the GRNs underlying PROD development, we performed tissue-specific RNA-seq of PROD imaginal discs at 2-4 selected developmental stages across a phylogenetically broad set of dipteran species: *De. wuyiensis*, *A. albopictus*, and *D. melanogaster*, representing the basal-most Diptera, Culicomorpha and Schizophora, respectively (Fig. 5a). As a developmental reference, we included wing imaginal discs, motivated by their shared origin from ventral embryonic primordia, and are patterned by conserved morphogens *wg*, *Dpp*, and *Dll*^20–22^ (Fig. 1). We further integrated transcriptomes from outgroup species with available prothoracic dorsal imaginal tissue (or corresponding tissues) and wing imaginal tissue data, including *Trypoxylus dichotomus*, *Onthophagus taurus*, *Homalodisca vitripennis*, and *Eupolyphaga carinata*^13,14,66^ (Fig. 5c and Supplementary Table S6). For cross-species comparison of differentially expressed genes (DEGs), we applied the “metagene” approach^67^, mapping orthologous groups identified by Orthofinder2^68^ onto *D. melanogaster* reference genes, resulting in 4785 one-to-one orthologs across species. Comparative analyses using these dataset enables us to identify both ancestral conserved GRNs and lineage-specific GRNs in PROD.

**Fig. 5.**
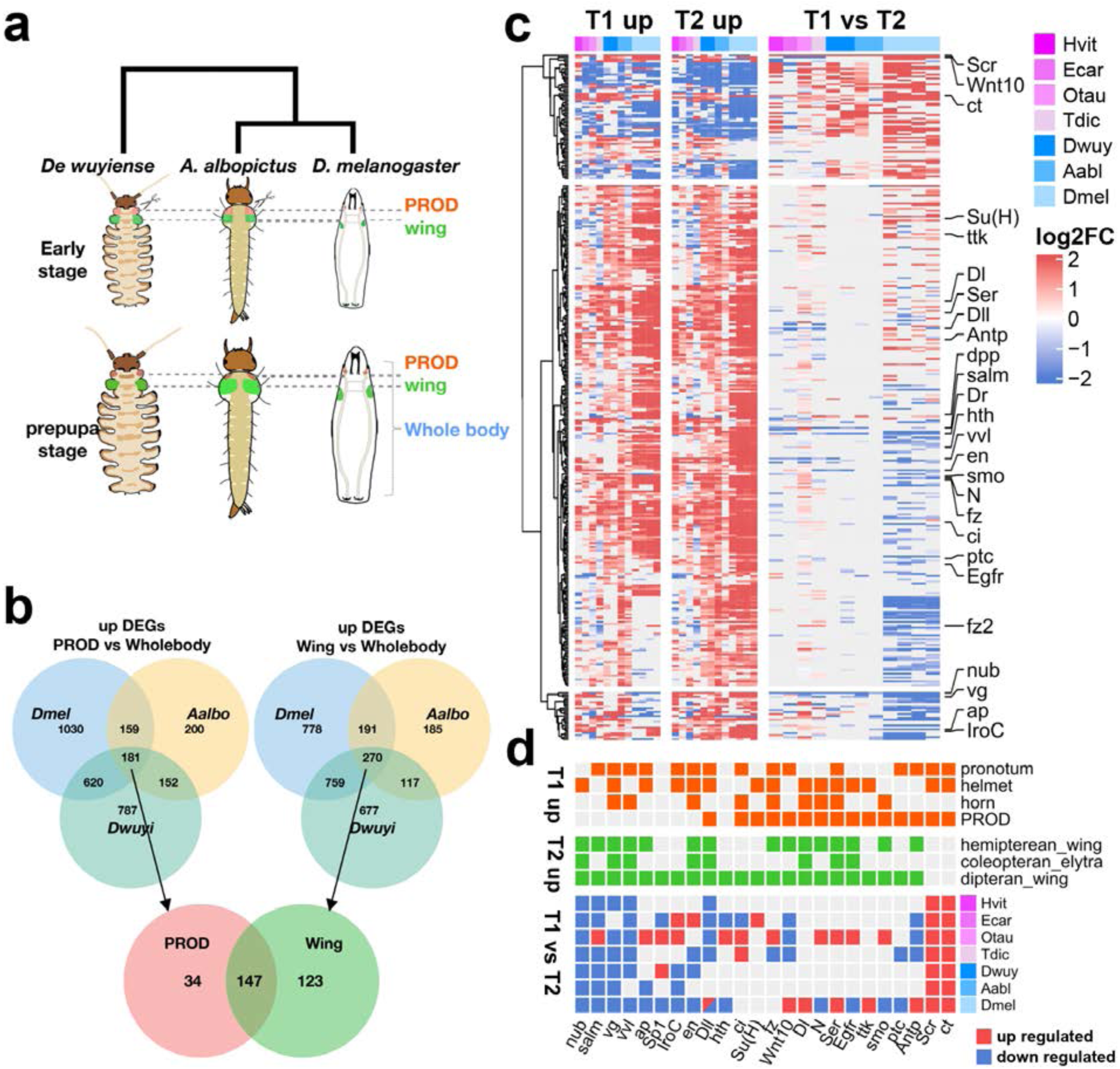
Phylogeny-informed transcriptomics reveal conserved GRNs underlying dipteran PROD and wing development. **(a)** Experimental design for tissue-specific RNA-seq of PROD and wing discs in three dipteran species: *De. wuyiensis* (basal lineage), *A. albopictus* (Culicomorpha), and *D. melanogaster* (Schizophora). For each species, at least two developmental stages were sampled per tissue. **(b)** Venn diagrams summarizing differentially expressed genes (DEGs) upregulated in PROD (left) and wing (right) relative to whole-body samples in the three dipteran species; and a combined diagram (bottom) showing DEGs shared between, or unique to, PROD and wing. **(c)** Differential gene expression analysis of thoracic tissues (PROD or WING) compared to whole-body controls (WHB) is shown in the first two columns, while PROD versus WING comparison is displayed in the third column on the right. The analysis includes seven species: three dipteran species—*De. wuyiensis* (Dwuy), *A. albopictus* (Aalb), and *D. melanogaster* (Dmel)—along with four outgroup species—*Trypoxylus dichotomus* (Tdic), *Onthophagus taurus* (Otau), *Homalodisca vitripennis* (Hvit), and *Eupolyphaga carinata* (Ecar)^13,14,66^. **(d)** Schematic summary of expression patterns of core patterning genes implicated in PROD morphogenesis. In*D. melanogaster*, *Dll* is upregulated in PROD relative to wing at 84 h AEL and downregulated at 120 h AEL, indicating a heterochronic shift in P/D axis deployment.

To identify conserved GRNs for PROD and wing development, we performed DEG analysis comparing PROD or wing discs to whole-body control at selected developmental stages in each species (Fig. 5b; Supplementary Figs. S7). Genes upregulated in either tissue at any stage were aggregated (Figs. 5b-c), revealing 692, 1,740, and 1,990 PROD-upregulated genes and 763, 1,823, and 1,998 wing-upregulated genes in *De. wuyiensis*, *A. albopictus*, and *D. melanogaster*, respectively (Fig. 5c). Intersecting these across three species, we identified 304 genes that were upregulated across species in either PROD or wing imaginal discs (Fig. 5c). Nearly half of these genes are shared between wing and PROD, including canonical patterning genes in the appendage morphogenesis: *Notch* (*N*), *Delta* (*Dl*), *Serrate* (*Ser*), *cubitus interruptus* (*ci*), *patched* (*ptc*), *engrailed* (*en*), *homothorax* (*hth*), *Distal-less* (*Dll*), and the *Hox* gene *Antennapedia* (*Antp*) (see GO and KEGG enrichments in Supplementary Fig. S8).

To uncover lineage-specific GRNs in PROD evolution, we compared expression of these 304 DEGs between PROD and wing across dipterans and outgroups (Fig. 5c and Fig. 5d; Supplementary Fig. S9 and S10). Notably, the homeobox gene *cut* (*ct*) and the Hox gene *Sex combs reduced* (*Scr*) are consistently upregulated in PROD relative to wing in all species (Fig. 5c and Fig. 5d; Supplementary Fig. S9). In *D. melanogaster*, *ct* is a master regulator of abdominal spiracle development, where it acts within a conserved GRN (referred to as spiracle GRN), together with *escargot* (*esg*) and other factors, to specify spiracular fate and mediate tracheal-epithelial morphogenesis^63–65^. Similarly, *ct* functions as a key Hox-dependent factor in posterior spiracle development, and the posterior-spiracle GRN has been implicated in the evolution of novel adult posterior appendages^69,70^. We therefore propose that *ct* may have an ancestral role in specifying the developmental fate of the adult spiracle and the pupal respiratory organ in T1.

Several key developmental genes display dipteran-specific expression patterns. *Dll*, a central regulator of proximal-distal (P/D) patterning, shows a transient expression pulse during *D. melanogaster* PROD development (Supplementary Fig. S9), suggesting a heterochronic deployment of the conserved arthropod appendage GRN^8,9^, which includes regulators defining the P/D (*hth*, *Dll*)^71–75^, dorsal-ventral (D/V; *N*, *Ser*, *Dl*)^20,76–80^, and anterior-posterior (A/P; *ci*, *en*, *ptc*) axes ^81–84^. In *D. melanogaster*, *Antp* has been shown to activate *Dll* by repressing its corepressor *hth*^85^, demonstrating a regulatory link between *Antp* and appendage patterning.

Notably, *Antp* is mostly downregulated in T1 relative to wing in outgroups, whereas dipterans show unchanged or elevated *Antp* expression in T1/PROD (Fig. 5d; Supplementary Fig. S10), consistent with a Diptera-specific role of *Antp* in regulating the appendage GRN underlying PROD development. Furthermore, *D. melanogaster* exhibits pronounced alterations in *N/Dl* signaling, which is center to both appendage patterning and tracheal cell-fate determination and branching morphogenesis—where *btl*-high expressing cells activate *Dl* to repress neighboring tracheal cell fates via *N*^62,63,86^ (referred to as trachea GRN). Regulatory changes in this pathway likely contributes to the more complex branched PROD in brachyceran flies.

These results suggest that PROD development shares a conserved appendage GRN with insect wing, but diverges through stepwise co-option of two additional GRNs, including a *ct*-mediated spiracle GRN and a putative tracheal GRN involving *btl* and *N/Dl* (Fig. 5d). To validate this hypothesis and dissect the roles of these candidate GRNs in PROD development, we leveraged scRNA-seq and genetic perturbations in *D. melanogaster* to dissect the spatiotemporal dynamics of PROD specification and morphogenesis.

### scRNA-seq and *in vivo* expression support GRN co-option in *D. melanogaster* PROD

To test whether PROD shares a deep developmental homology with the wing, we performed scRNA-seq of the *D. melanogaster* PROD disc at 120 h AEL, complemented by spatial-temporal expression analysis (Fig. 6; Supplementary Fig. S11-S12). Similar to the wing disc, the PROD tissue comprises four major cell types: epithelial cells (presumptive imaginal cells), tracheal cells, muscle precursors, and hemocytes. Within the epithelial compartment, we detected high expression of *ct* and *Antp*, genes associated with respiratory fate and thoracic identity, confirming its prothoracic, spiracle-related identity (Fig. 6b; Supplementary Table S6).

**Fig. 6.**
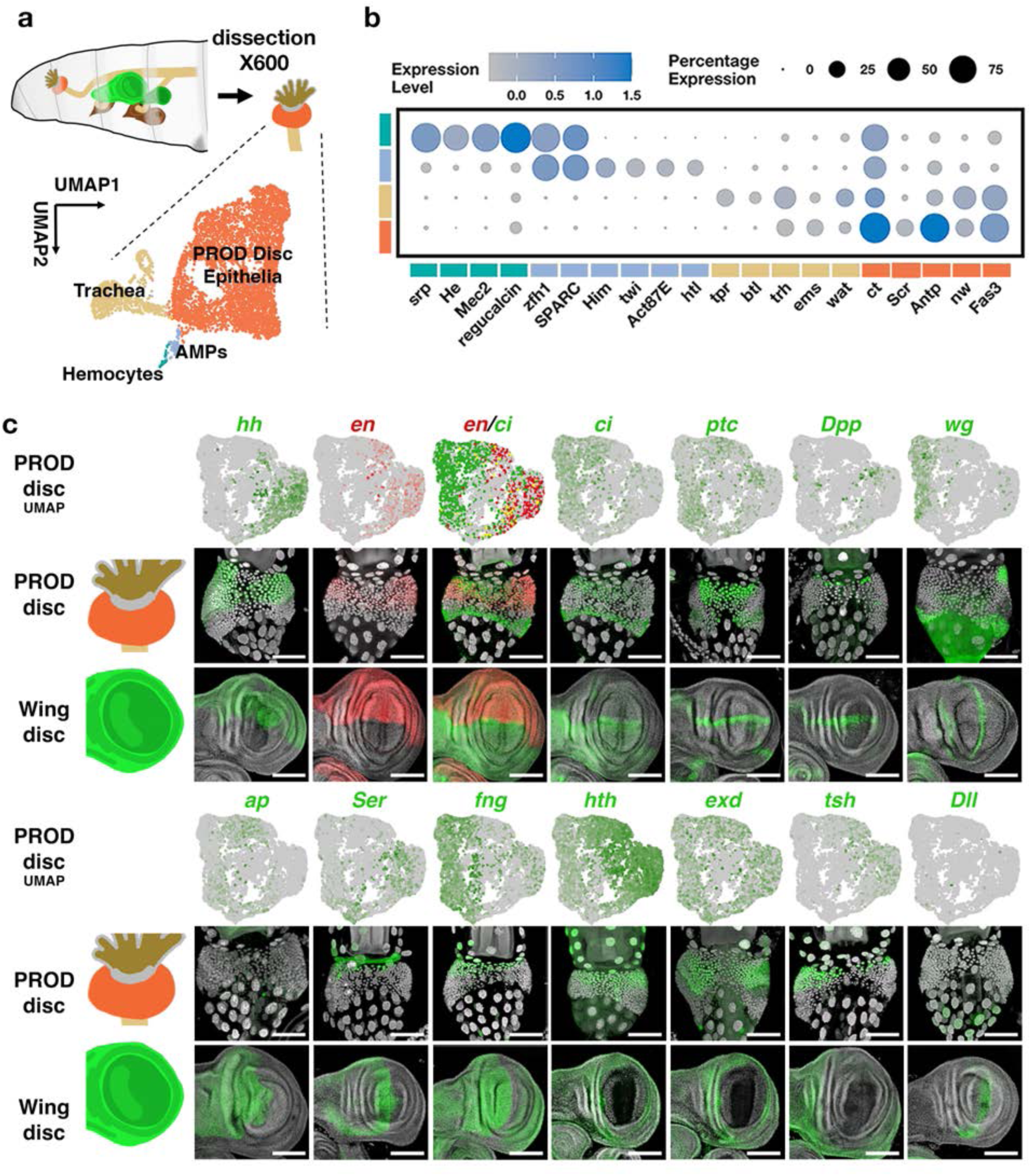
scRNA-seq and spatial expression patterns of appendage GRN in the *D. melanogaster* PROD. (a-b) scRNA-seq analysis of *D. melanogaster* PROD discs. **(a)** UMAP projection of PROD disc scRNA-seq revealing four major cell types: epithelial cells, tracheal cells, hemocytes, and adult muscle precursors. **(b)** Marker gene expression used to assign cell-type identities. **(c)** Expression of developmental patterning genes in epithelial cells of the PROD disc compared to the wing disc, revealed by scRNA-seq, antibody staining, and in vivo reporter assays. These patterns identify epithelial cells as the main imaginal compartment and nominate candidate appendage and spiracle GRN components for functional testing. Scale bars: 50 μm (PROD disc), 100 μm (wing disc).

We next compared the gene regulatory logic of PROD and wing development by focusing on the conserved appendage GRN. In the wing, compartmentalization follows a canonical hierarchy: A/P patterning (*en*, *hedgehog* [*hh*]; *ci*, *ptc*, and *Dpp*) precedes D/V patterning (*ap*, *fringe* [*fng*]); *Ser*, and *wg*), which in turn forming P/D axis through activation of *Dll*, mediated via *wg* and repressed by *hth* and *exd*^71–84^. While the PROD disc retains conserved expression patterns of A/P genes, D/V gene expression is markedly divergent from that in the wing (Fig. 6c). The P/D axis genes *hth* and *exd* are broadly expressed across the disc, while *Dll* expression is nearly undetectable at this stage (Fig. 6c).

To resolve the dynamics of *Dll* expression, we examined time-series transcriptome data and *Dllmd23-GAL4* enhancer trap reporter, and revealed that *Dll* expression peaks at 84 h AEL in the PROD disc, significantly earlier compared to the wing disc (Fig. 7a). This is consistent with the downregulation of *hth* (a corepressor of *Dll*) at 84 h AEL in the PROD disc relative to the wing disc (Fig. 5c; Supplementary Fig. S9), and aligns with the onset of PROD morphogenesis (Fig. 4c and Supplementary Fig. S9). This heterochronic shift in the P/D axis suggest changes in appendage GRN timing contribute to PROD outgrowth at pupal stage.

**Fig. 7.**
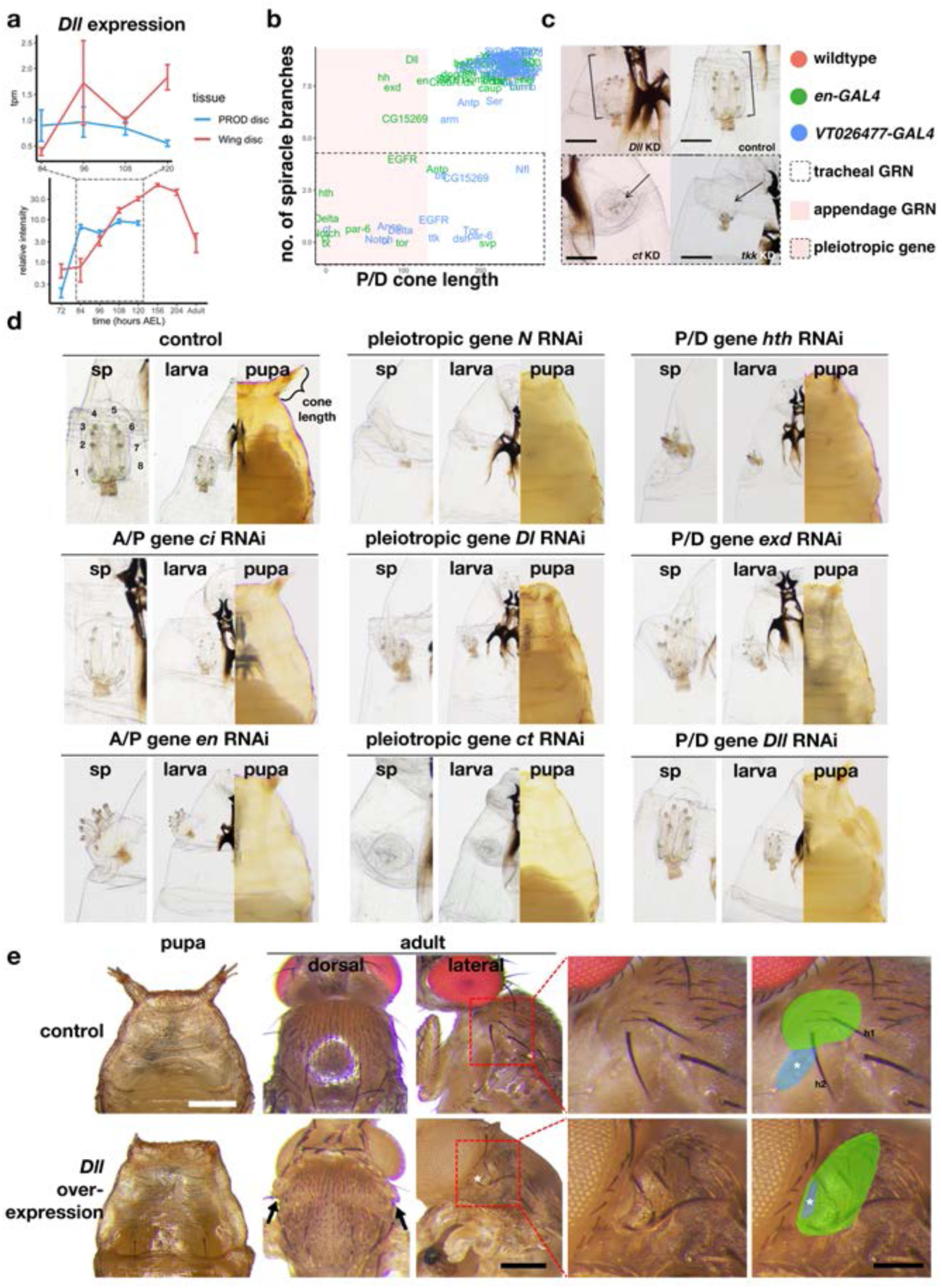
Three partially separable GRN modules underlying PROD morphogenesis in *D. melanogaster*. **(a)** Heterochronic expression patterns of *Dll* between the PROD disc and wing disc reveal differences in organ development timing, using RNA-seq and *Dllmd23-GAL4* reporter assays. **(b)** Summary of RNAi phenotypes for 70 candidate genes using *en-GAL4* (green) and *VT026477-GAL4* (blue) drivers. Each gene name position represents the mean cone length (*x*-axis) and number of tracheal-spiracular branches (*y*-axis) of its knockdown. Wild-type controls in red. Phenotypes cluster into four classes: no effect, altered cone length only, altered branch number only, and pleiotropic effects on both cone length and branch number. **(c)** Representative examples each phenotypic class. Brackets indicate the length of invaginated epithelial tube (future PROD cone after eversion, A/P axis); arrows mark positions of normal branch formation. Scale bars: 100 μm. **(d)** Representative RNAi phenotypes of key appendage GRN components (*en-GAL4* driver) with disrupted PROD morphogenesis, confirming that PROD depends on an appendage GRN. **(e)** *en-GAL4*-driven *Dll* overexpression eliminates the PROD cone, induces a protruding humeral callus in the adult anterior spiracle region, and causes posterior wing defects. Scale bar: 100 μm.

We further examined the spiracle and trachea GRNs, mediated by *ct* and *btl*, respectively. High expression of *ct* in the PROD disc, observed both *in vivo* and in scRNA-seq data (Fig. 6b; Supplementary Figs. S11 and S13), is similar to that observed in abdominal^64,65^ and terminal^69,70^ primordia that give rise to larval and adult posterior spiracles, which supports a shared developmental program for respiratory fate specification. In contrast, *btl* expression is predominantly expressed in tracheal cells, with spatial pattern recapitulated by *btl-GAL4*, which marks the innermost layer of cells surrounding the larval T1 spiracular branch (Supplementary Figs. S12 and S13). This expression pattern resembles tracheal-epithelial interactions in wing disc^62^ and abdominal imaginal cells^63–65^, and support that a *btl*-centered tracheal GRN^86^ contributing to tracheal recruitment in *D. melanogaster* PROD.

Together, these data suggest that PROD development potentially integrates three modular programs: a heterochronic-shifted appendage GRN, a *cut*-dependent spiracle GRN, and a *btl*-centered tracheal GRN to build a novel respiratory structure in pupae.

### Three GRNs underlying modular morphogenesis in *D. melanogaster* PROD

To test whether the GRNs and candidate genes identified through RNA-seq and scRNA-seq are functionally required for PROD development, we conducted RNAi screen of 70 candidate genes in *D. melanogaster*, using PROD size and spiracular branch number as quantitative phenotypic readouts (Fig. 7b, c; Supplementary Fig. S14). To ensure spatial targeting, we evaluated a panel of GAL4 drivers, including those marking appendage patterning genes and enhancer trap lines (enhancers.starklab.org; Supplementary Fig. S11 and Table S7). Two drivers were selected for dual validation: *en-GAL4*, a broadly expressed driver in the PROD disc, and *VT026477-GAL4*, a more restricted driver with weak but PROD-specific expression (Supplementary Fig. S11).

We performed parallel RNAi knockdowns using both *GAL4* drivers. As expected, *en-GAL4* knockdowns produced more severe phenotypes, often associated with embryonic lethality, consistent with its broader and stronger expression (Fig. 7b; Supplementary Fig. S14 and Table S9). In contrast, *VT026477-GAL4* allowed viable assessment of PROD morphology with minimal pleiotropy (Supplementary Fig. S14 and Table S9). Knockdown of the Hox genes *Antp* and *Scr* both disrupted PROD development, as expected, with *Antp* showing a complete loss of PROD, including loss of tracheal-spiracular branches (Fig. S14).

We demonstrated that key components of the appendage GRN—including A/P genes (*en*, *ci*), D/V genes (*N*, *Dl*), and especially P/D genes (*hth*, *exd*, *Dll*)—are essential for PROD morphogenesis in *D. melanogaster* (Fig. 7b, d). To test whether the heterochronic shift of the P/D axis is functionally important, we prolonged *Dll* expression in the PROD disc via *en-GAL4*. Paradoxically, sustained *Dll* expression eliminated the pupal respiratory protrusion and induced ectopic outgrowth in the adult humeral callus, the cuticular structure surrounding anterior spiracle (Fig. 7e). This phenotype indicates that precise temporal control of *Dll* is required for proper P/D patterning. The fact that *en-GAL4*-driven *Dll* overexpression also caused posterior malformations in the wing underscores a serially repeated appendage field in T1 and T2 marked by *en*. Together with the shared expression of A/P, D/V, and P/D patterning systems in PROD and wing, support the hypothesis that the PROD is a serial homolog of wing—structures derived from repeated morphogenic cues of the conserved appendage GRN along the body axis, such as wings and halteres^20–22^.

RNAi of spiracle GRN gene *ct* and trachea GRN genes (*btl*, *Dl*, *N*) also produced predicted phenotypes (Figs. 7b-d). Knockdown of *ct* generated complete loss of PROD and its tracheal-spiracular branches, consistent with its broad expression within PROD epithelial cells (Supplementary Fig. S11). This supports a dominant role of *ct* in PROD development analogous to its role in abdominal and terminal spiracle morphogenesis^64,65,69,70^. Furthermore, we identified *par-6*, *tx*, and *Ser* as additional regulators required for proper formation of the spiracular opening on each branch, suggesting they operate downstream of the *ct*-mediated spiracular GRN (Supplementary Fig. S15 [panel 52, 59, and 67]).

For the trachea GRN, knockdown of *N* and *Dl* caused pleiotropic effects, i.e., defective tracheal-spiracular branches consistent with their known roles in early tracheal development^86^ (Fig. 7d), as well as shortened PROD cone length. To refine this pathway, we focused on *btl*, which functions upstream of *N* and *Dl* in embryonic trachea GRN to mediate branching morphogenesis. *btl* RNAi recapitulated the branching defects without affecting cone length, indicating that tracheal branching and distal cone morphogenesis are modular traits (Supplementary Fig. S14 [panel 9]). Additional genes (*Ance*, *CG15269*, *ttk*) were also identified as contributing to spiracular branch morphogenesis (Supplementary Fig. S14).

Collectively, these experiments indicate that *D. melanogaster* PROD morphogenesis can be dissected into at least three partially separable GRN modules: a *ct*-centered spiracle GRN that specifies overall PROD identity and opening formation, an appendage GRN that patterns the distal cone, and a tracheal GRN that controls the size and number of tracheal-spiracular branches. In *D. melanogaster*, components of these modules also interact in part through the *N/Dl/Ser* signaling pathway.

### Functional dissection of *A. albopictus* PROD reveals ancestral GRNs and lineage-specific co-option

To determine whether the GRNs underlying PROD development in *D. melanogaster* are evolutionarily conserved across Diptera, we also performed RNAi in the mosquito *A. albopictus*, a distantly related lineage in Culicomorpha. We assessed the functional roles of over 20 key genes whose orthologs affect PROD development in *D. melanogaster* (Figs. 8-10; Supplementary information, Figs. S16-S23 and Tables S10-S12). RNAi efficiency in injected larvae was validated by qPCR (Supplementary Fig. S15 and S16), and larval lethality, developmental defects, and morphometrics were measured. We found that, with the exception of *Aa-ems*, *Aa-btl*, and *Aa-trh*, knockdown of all tested genes affected PROD development in *A. albopictus* (Supplementary Fig. S17 and Table S12).

**Fig. 8.**
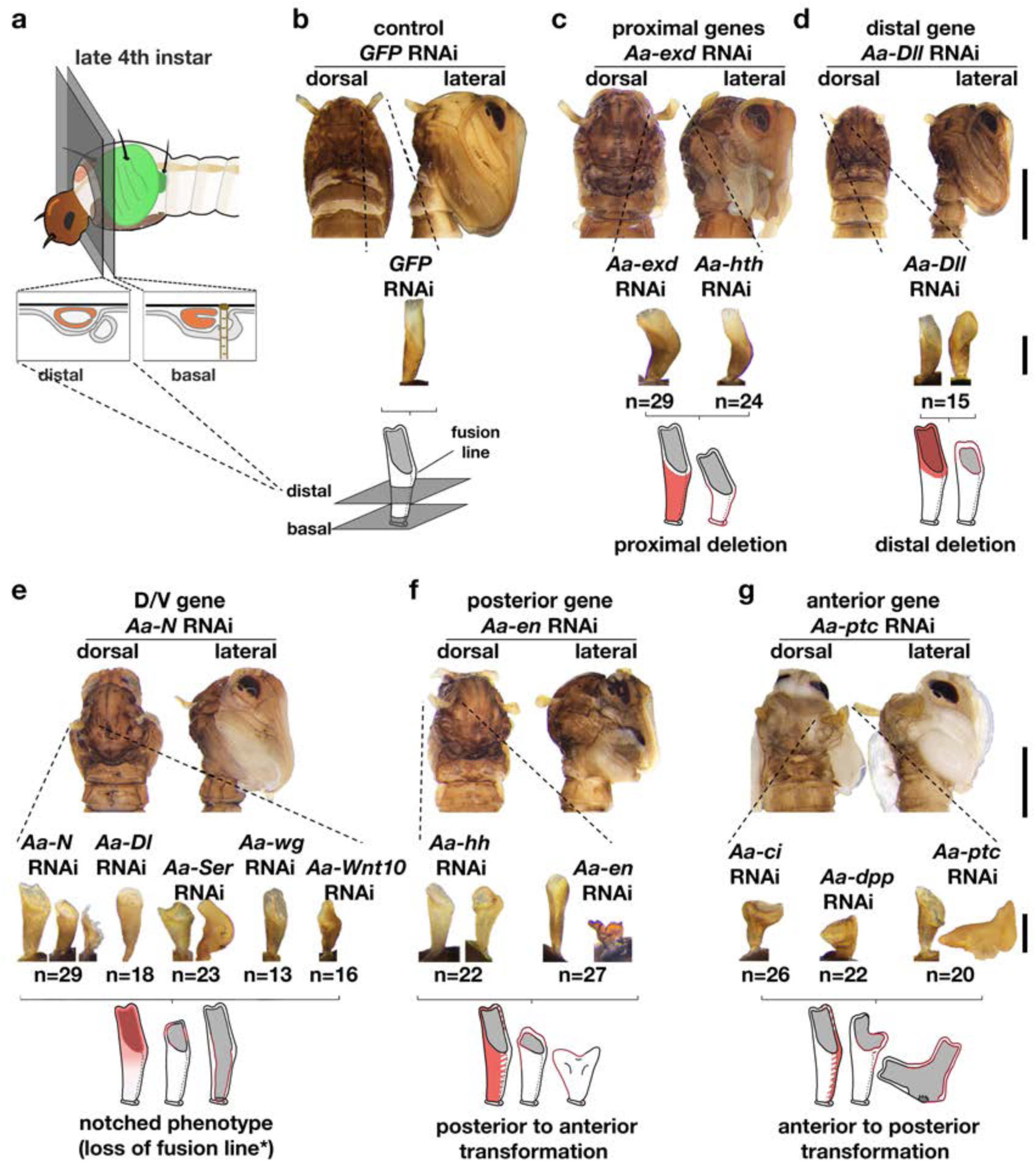
Systematic RNAi of core appendage GRN genes in *A. albopictus* reveals conserved mechanisms underlying PROD morphogenesis. **(a)** Schematic of the larval PROD disc in *A. albopictus*, showing its proximal-distal organization and the corresponding regions in the snorkel-like PROD at the pupal stage (see **b**). **(b)** The *dsGFP* injection control, showing normal PROD morphology and illustrating two transvers plans. **(c-d)** RNAi knockdowns of proximal-distal patterning gene: *Aa-hth* and *Aa-exd* knockdowns produce L-shaped organs with enlarged openings due to proximal region deletion, while *Aa-Dll* knockdowns yield organs with smaller, rounded openings reflecting distal region deletion. Cartoon depicting the inferred deleted regions. **(e)** Dorsal-ventral patterning gene knockdowns (*Aa-N*, *Aa-Dl*) produce rounded openings and exhibit tubeless defects, while the wing notching mirrors classic *N* signaling pathway defects in *D. melanogaster*. **(f)** Knockdowns of anterior-posterior genes (*Aa-en*, *Aa-ptc*) generate PROD with polygonal openings and curled tubes. Scale bar: 500 μm (upper panels), 200 μm (lower panels).

**Fig. 9.**
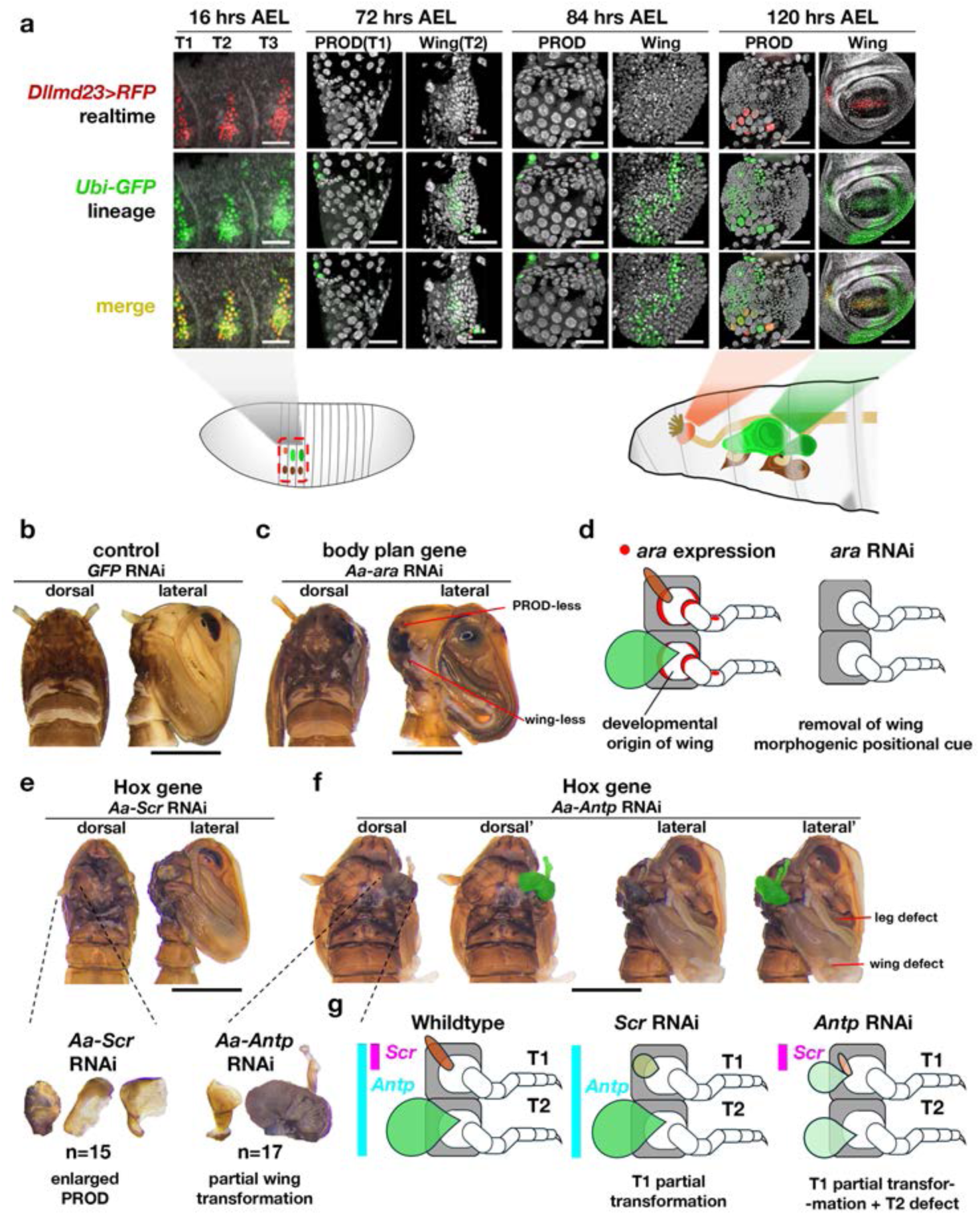
Lineage tracing and Hox/homeobox RNAi support a PROD-wing serial homology. **(a)** G-TRACE lineage tracing in *D. melanogaster* driven by *Dll^md23^-GAL4. GFP*-positive clones show that the PROD disc and wing disc both derive from embryonic ventral *Dll*-expressing thoracic appendage primordia patterned by positional cues from *wg*/*Dpp*/*Dll*. This confirms a shared developmental origin for the PROD and wing in *D. melanogaster*. **(b)** RNAi of GFP in *A. albopictus* pupae shows normal phenotype and serves as the control. **(c)** RNAi targeting *Aa-ara*, a serial patterning gene, eliminates both PROD and wing, indicating a shared serial identity. **(d)** Schematic of conserved body plan gene *ara* expression across Insecta, delineating developmental origin of wing/wing serial-homologs (adapted from ref^8^). (e–f) RNAi of the *Hox* genes (*Aa-Scr*, *Aa-Antp*) in *A. albopictus* induces partial T1-to-T2 transformations, resulting in larger PROD lacking openings and cuticle texture, resembling ectopic wings. Red lines highlight defective appendages (e.g., wings, legs). **(g)** Schematic of *Hox* gene expression domains in dipteran thorax, illustrating the regulatory context of PROD specification. Together, these data support that the PROD and wing develop from serially homologous thoracic fields defined by *ara* and Hox expression. Scale bar: 500 µm.

**Fig. 10.**
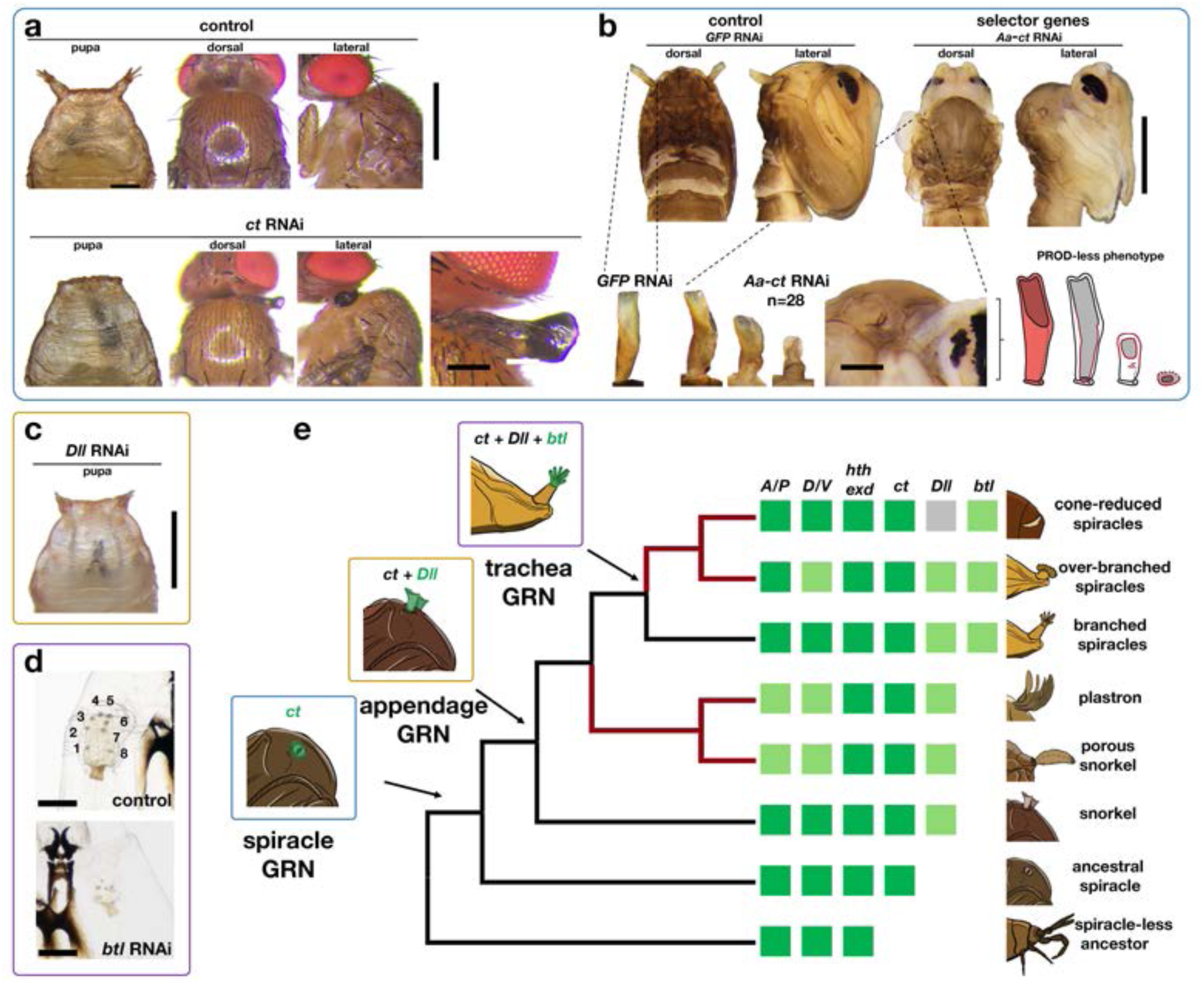
Model for stepwise co-option of GRN in PROD evolution. **(a)** *ct* knockdown in *D. melanogaster* eliminates the PROD and its tracheal-spiracular branches, and induces overgrowth in the adult humeral region, consistent with a homeotic-like transformation. These phenotypes support a *ct*-centered spiracle GRN in specifying PROD/adult spiracle fate. Scale bars: 500 μm or 200 μm (magnified view). **(b)** *Aa-ct* knockdown in *A. albopictus* disrupts PROD development, with severe cases showing complete loss, confirming conservation of the *ct*-centered spiracle GRN across Diptera. Scale bars: 500 μm (pupae), 200 μm (magnified views). **(c)** Perturbation of *Dll* in *D. melanogaster* (see also Fig. 7d-e and Fig. 8c-d for *A. albopictus*) disrupt PROD morphogenesis, demonstrating the importance of a heterochronic appendage GRN module. Scale bar: 500 μm. **(d)** *btl* knockdown in *D. melanogaster* specifically abolishes spiracular branch formation without affecting cone, implicating a *btl*-dependent tracheal GRN in elaborating branching. In contrast, *Aa-btl* knockdown in *A. albopictus* does not produce detectable PROD defects, suggesting that tracheal contribution is not required for snorkel-type PRODs. Scale bar: 500 μm. **(e)** Working model of stepwise GRN co-option in dipteran PROD evolution. We propose that: (1) an ancestral *ct*-centered spiracle GRN established spiracular field in T1; (2) this field was patterned by co-option of appendage GRN, with heterochronic expression of *Dll*; and (3) in Schizophora, a *btl–N/Dl* tracheal GRN was recruited to generate branched PRODs. Red branches are hypothesized regulatory changes that produce natural morphological diversity, recapitulated by mutant phenotypes observed in *A. albopictus* and *D. melanogaster*; colored boxes denote GRN components implicated at each stage. This modular, hierarchical assembly provides a mechanistic framework for how a novel respiratory organ evolved through the sequential integration of conserved developmental programs, and offers testable hypotheses for dissecting the genetic basis of PROD diversity across Diptera.

The conserved roles of these patterning genes in appendage development were confirmed by comparison with known *D. melanogaster* mutant phenotypes. Knockdown of P/D patterning gene *Aa-Dll* resulted in truncated legs and antennae and reduced wings, closely matching *D. melanogaster Dll* mutants^71,72^ (Supplementary Fig. S18a, d). Silencing the *Dll* corepressors *Aa-hth* and *Aa-exd* produced similar defects, consistent with phenotypes in both arthropods^8^ and *D. melanogaster hth*^73,74^ and *exd*^75^ mutants (Supplementary Fig. S18b, c). Knockdown of D/V patterning genes (*Aa-N*, *Aa-Dl*, *Aa-Ser*) caused leg and antennal defects and notched wings (Supplementary Fig. S19a-d), phenocopying *D. melanogaster* mutants^76–79^. Notably, disruption of *N/Dl* signaling more strongly affected the segmentation of the multi-annulated mosquito antenna (Supplementary Fig. S19b). *Aa-wg* and *Aa-Wnt10* knockdowns produced wing and leg phenotypes similar to *D. melanogaster wg* alleles^20,80^ (Supplementary Fig. S19e, f). A/P patterning gene knockdowns (*Aa-hh*, *Aa-ptc*; *Aa-hh*, *Aa-en*; *Aa-ci*) induced defects in legs, antennae, and wings, including blistered wings—a hallmark of polarity loss in *D. melanogaster*^81–84^ (Supplementary Fig. S20). In addition, knockdowns of *Aa-araucan* (*Aa-ara*) led to significantly malformed compound eyes in *A. albopictus* (Fig. 9c), a phenotype similar to that seen in *D. melanogaster* mutants^87^. *Aa-ct* knockdowns resulted in notched wings and minor patterning defects in the compound eye (Fig. 10b, Supplementary Fig. S21a, e), similar to the phenotypic traits observed in *D. melanogaster ct* mutants^88–90^. Together these comparisons confirm the robustness of RNAi in *A. albopictus* and indicate strong conservation of key gene function across Diptera, forming a foundation for interpreting PROD phenotypes.

We next focused on PROD phenotypes in *A. albopictus* RNAi knockdowns, integrating our results with morphological observations of mosquito PROD morphogenesis (Fig. 4c) and previous studies^18,19,21^. Disruption of patterning genes in appendage GRN produced predictable, axis-specific defects in *A. albopictus* PROD, consistent with their roles in typical appendage morphogenesis and confirming that PROD development deploys a conserved GRN. Knockdown of P/D genes, *Aa-hth*/*Aa-exd* produced uniformly L-shaped organs with enlarged openings on shortened tubes, reflecting proximal deletion (Fig. 8c). *Aa-Dll* disruption led to rounder, and often deformed openings, reflecting distal deletion (Fig. 8d). D/V gene knockdowns produced shortened PRODs with rounded openings, resembling the *Aa-Dll* phenotype, along with a distinct tubulation defect (Fig. 8e). This tubulation failure was most severe in *Aa-N/Aa-Dl* knockdowns, which is consistent with eliminated positional cue along the fusion line (Fig. 8a), analogous to the notched-wing phenotype in both *D. melanogaster N/Dl* mutants^91^ and in *A. albopictus N/Dl* RNAi knockdown wings (Supplementary Fig. S19b, c). A/P knockdowns caused deformed polygonal openings and severe tubulation defects (Fig. 8f, g), reminiscent of A/P mirror-image transformations observed in *D. melanogaster* wing^82^.

For genes in spiracle GRN, knockdown of *Aa-ct* knockdown resulted in complete absence of the PROD (Fig. 10b), whereas *Aa-tx* and *Aa-par-6* knockdowns caused only mild defects in the spiracular opening (Supplementary Fig. S21c, d). However, knockdown of the key genes of tracheal GRN in *A. albopictus Aa-btl* and *Aa-trh* resulted in no detectable PROD defects within the sensitivity and survival limits of our assay, indicating that mosquito PROD morphogenesis does not depend on tracheal GRN components (Supplementary Fig. S22). This contrasts sharply with their essential roles in *D. melanogaster*, where *btl* and *trh* are required for tracheal-spiracular integration.

Taken together, the functional data from *D. melanogaster* and *A. albopictus* indicate that PROD morphogenesis is built upon an evolutionarily conserved set of GRN modules. A *ct*-centered spiracle GRN is required for establishing PROD identity, and an appendage GRN involving canonical A/P, D/V, and P/D patterning genes controls the overall shape and distal morphology of PROD. In contrast, the role of the tracheal GRN diverges between two species. Whereas *btl* and *trh* are essential for tracheal-spiracular integration in *D. melanogaster* PROD, knockdown of them produces no detectable PROD defects *in A. albopictus*. This is consistent with our hypothesis that ancestral dipteran PROD was patterned by conserved appendage GRN and *ct*-centered spiracle GRN, and that recruitment of the tracheal GRN represents a later innovation linked to the evolution of complex, branched PROD in the Schizophora.

### Cell lineage tracing, RNAi and homeotic transformations reveal PROD-wing serial homology

To test whether PROD and wing discs initially developed from the same serially repeated positional cues patterned during embryogenesis, we conducted genetic mosaic lineage tracing using G-TRACE^92^ in *D. melanogaster*. Driven by *Dllmd23-GAL4* enhancer trap, this system labels embryonic *Dll*-expressing progenitor cells and their descendants. We found that PROD epithelial cells originate from embryonic thoracic appendage primordia induced by *Wg/Dpp*, identical to wing disc progenitors (Fig. 9a). During embryogenesis, ventral *Dll*-expressing cells give rise to leg discs or migrate dorsally to form wing/haltere precursors^20,22^ (Fig. 1a). Because *Dll* expression halts after embryogenesis and resumes near the end of larval development^93^, *GFP*-positive cells in early larvae stage mark exclusively embryonic *Dll* progenitors. The presence of *GFP*-positive cells in both wing and PROD discs confirms their shared embryonic origins from serially repeated thoracic appendage primordia.

Further evidence for PROD-wing serial homology comes from RNAi experiments in *A. albopictus*. Knockdown of *Aa-ara*, a homeobox gene required for dorsal appendage development, abolished both PROD and wing development with a pronounced hunched-back posture, while the legs and antennae remained unaffected (Fig. 9c, d). This specific co-loss demonstrates that PROD and wing develop from serial-homologous regions in T1 and T2, which are defined by two parallel strips of *ara* expression at basal leg segments where wing and its serial homologs derive from (Fig. 9d)^8,94^. Moreover, knockdown of *Aa-Antp* resulted in an ectopic wing-like outgrowth in T1, while residual *Aa-Scr* expression preserved partial PROD fate (Fig. 9f, g). Knockdown of *Aa-Scr* induced an enlarged, opening-less PROD with smooth cuticular texture, a phenotype suggestive of partial homeotic transformation toward a wing-like identity (Fig. 9e, g).

Collectively, these results demonstrate that PRODs and wings share a common embryonic origin and a core positional identity governed by conserved patterning genes. Their development is orchestrated by serially deployed regulatory programs in T1 and T2, and PROD can be partially converted to wing-like structure through perturbation of Hox genes *Antp* and *Scr*. These findings support the conclusion that the dipteran PROD is a serial homolog of the wing, derived from similar thoracic appendage primordia, rather than an independently evolved respiratory outgrowths.

### Stepwise GRN co-option in PROD evolution

Our findings support a model in which dipteran PRODs are serial homologs of the wing, sharing same developmental origin from dorsal appendage primordium, and diversified through the stepwise co-option and modification of preexisting GRNs. Although appendage GRN provides the foundational patterning logic governing A/P, D/V, and P/D axes, two key questions remain: which regulators specify PROD fate, and how additional features—such as the elaborated spiracular branches in Schizophora—were incorporated during evolution.

To address the first question, we focused on transcription factors that are consistently upregulated in PRODs across Diptera. Among the candidates (*ct*, *tx*, *par-6*), only *ct* knockdown eliminated PROD formation in both *D. melanogaster* and *A. albopictus* (Fig. 10a, b; Supplementary Fig. S21). In *D. melanogaster*, *ct* loss also produced a pupal/adult protrusion phenotype resembling that caused by prolonged *Dll* expression, and in some cases induced ectopic wing-like tissue (Fig. 10a). Together with the conserved T1-specific upregulation of *ct* across Diptera and outgroup insects—including Hemiptera and Coleoptera (Fig. 5c)—and its well-established role in maintaining abdominal and terminal spiracular fate^64,65,69,70^, these observations indicate that *ct* playing a central role in PROD and adult spiracle identity. Our data confirmed that the *ct*-centered spiracular GRN, conserved in insect respiratory development, has been co-opted to specify the fate of PROD.

Within this framework, the PROD emerges a spiracular outgrowth specified by high *ct* activity and patterned by a heterochronic pulse of *Dll* (Fig. 7a, e). The early transient *Dll* expression in the PROD, coinciding with reduced *hth* expression, suggests that distal identity is imposed on a *ct*-positive spiracular field. Co-expression of *Antp* and *Dll* in the distal PROD domain (Fig. 6c; Supplementary Fig. S6a) raises the possibility that *Dll* activation is mediated through *Antp*-dependent repression of *hth*, analogous to the regulatory logic underlying antenna-leg differentiation^85^, although this remains to be tested.

To understand how complex, branched PRODs evolved in Schizophora, we examined the contribution of the tracheal GRN. In *D. melanogaster*, epithelial-tracheal interactions mediated by *N/Dl* signaling coordinate PROD spiracular branching and opening formation. Given that *btl* acts upstream of *N/Dl* in embryonic tracheogenesis^86^, we tested its role in PROD development. *btl* knockdown phenocopied *N* knockdown, indicating that *btl* promotes spiracular branch formation via *N/Dl* signaling (Fig. 7d, 10d, Supplementary S14 [panel 9]). Moreover, *Ser* specifically modulates branch number, while *Dl* influences opening number, revealed by reciprocal effects in overexpression assays: *Dl* overexpression increased spiracular openings but reduced branches, whereas *Ser* overexpression promoted branching with decreased openings (Supplementary Fig. S23). *In vivo* and scRNAseq analyses confirmed that *btl* and *Dl* are co-expressed in tracheal cells associated with spiracular branches (Supplementary Fig. S11 and S24). These results support a *btl-N/Dl/Ser* module that drives spiracular branching morphogenesis in *D. melanogaster* PROD. This represents a key developmental divergence from the wing disc, which interacts with a non-spiracular tracheal branch^62^, indicating that the epithelial-tracheal interactions are deployed in different local GRN contexts to generate divergent structures.

In *A. albopictus*, however, knockdown of *Aa-btl* or *Aa-trh* had no detectable defects in PROD morphology within the survival and sensitivity limits of our systemic RNAi assays (Supplementary Fig. S22). While RNAi efficiency varies across tissues in *A. albopictus*, the absence of consistent PROD defects, contrasted with the robust branching defects observed in *D. melanogaster*, strongly suggests that the tracheal GRN does not play an essential role in PROD morphogenesis in mosquitoes. This functional divergence between species supports the hypothesis that integration of the *btl*-dependent tracheal GRN into PROD development occurred specifically in the schizophoran lineage, coinciding with the evolution of complex, branched respiratory structures.

We therefore propose a stepwise model for PROD evolution: (1) an ancestral *ct*-specified spiracular field was established in prothorax; (2) in Diptera, this field was patterned by co-option of appendage GRN, with a heterochronic pulse of *Dll* enabling distal outgrowth in pupae; and (3) in Schizophora, a *btl-N/Dl/Ser* tracheal GRN was recruited to generate more complex, branched PRODs (Fig. 10e). This modular, hierarchical assembly explains how a serial homolog of the wing acquired a novel respiratory identity and, in select lineages, evolved increased complexity. The PROD thus exemplifies how new organs arise not through de novo innovation, but through the sequential integration of ancient developmental programs, a principle likely shared across the evolution of morphological novelty.

## Discussion

Our study integrates phylogenomics, ASR, and cross-species functional genetics to resolve the origin and developmental basis of the dipteran pupal respiratory organ (PROD). By bridging deep-time evolutionary patterns with experimentally dissected GRNs, we provide a mechanistic view of how a novel organ arises through the co-option and reassembly of ancient developmental programs. We identify the transcription factor *ct* as a key determinant of PROD fate, extending its established role in spiracle specification^64,65,69,70^ to a dorsal, serially homologous appendage field. We show that PROD deploys an appendage GRN—shared with the wing—including canonical A/P, D/V, and P/D patterning systems, and that in schizophoran flies, a *btl*-centered tracheal GRN was independently recruited to drive branching morphogenesis. Together, these findings reveal that the PROD is a composite structure built through the stepwise integration of modular GRNs, a paradigm for the evolution of morphological novelty.

Our densely sampled Diptera phylogeny, based on genomic and transcriptomic data from 235 dipteran species, resolves key relationships within Brachycera and supports the monophyly of major infraorders with high confidence, offering a more robust and taxonomically comprehensive backbone than previous studies. On this tree, ASR supports a single origin of PROD at the base of Diptera, followed by with multiple independent gains of tracheal-spiracular branching, most notably in the Schizophora. The temporal overlap of these evolutionary events with major geological perturbations — the Carnian Pluvial Episode and the K-Pg mass extinction — is consistent with the hypothesis that environmental upheavals created selective pressures favoring novel respiratory strategies^17–19^, which aligns with broader patterns of animal radiation following mass extinction or climate shifts^58,59^.

By integrating developmental genetics in *D. melanogaster* and *A. albopictus* with comparative transcriptomics, we establish the PROD as a serial homolog of the wing. In both species, the PROD derives from the T1 dorsal imaginal disc, and undergoes stereotyped invagination and protrusion to forms the adult prothoracic spiracle. Single-cell and bulk RNA-seq show that PROD and wing discs share a core appendage GRN, including canonical A/P (*ci*, *en*, *ptc*, *Dpp*), D/V (*N*, *Dl*, *Ser*, *wg*, *ap*, *fng*), and P/D (*hth*, *exd*, *Dll*) patterning genes. Genetic lineage tracing in *D. melanogaster* and RNAi of patterning genes in *A. albopictus* confirm that PROD and wing epithelial cells originate from serially repeated morphogenic cues, providing direct developmental evidence that the PROD is not an independent outgrowth, but a modified dorsal appendage.

We note that in both species, knockdown of *Scr* or *Antp* alone result in a partial PROD-to-wing transformation, unlike the classic T3-to-T2 transformation in *Ubx⁻/⁻* mutants^1^. A full humeral-to-wing transformation has been reported by manipulating regulators of Hox expression^24^, suggesting that *Scr* and *Antp* likely act together to stabilize PROD fate. In *A. albopictus*, *Antp* RNAi induces a partial PROD-to-wing transformation, while in *D. melanogaster Antp*^-/-^ and *Antp*^-/-^; *ct*^-/-^ mutants in *D. melanogaster* show overgrowth in humeral callus^95^, a phenotype that has previously been interpreted as wing transformation^2,23^. Alternative interpretations are: for example, PRODs might be viewed as a novel trait without a clear serial homolog (analogous to larval prolegs in Lepidoptera^96^), or primarily as serial homolog of abdominal spiracles with secondary co-option of appendage GRN components. However, the shared embryonic origin from thoracic appendage primordia, the requirement for canonical appendage GRN, and the partial PROD-to-wing transformations induced by *Antp* knockdown together argue more strongly for a wing serial homology.

Within this shared framework, PROD identity is distinguished by the integration of a *ct*-centered spiracular GRN and a heterochronic deployment of appendage GRNs. *ct* is consistently upregulated in PRODs relative to wings. Loss of *ct* abolishes PROD formation in both *D. melanogaster* and *A. albopictus*, and in some cases, led to ectopic wing-like outgrowth in *D. melanogaster*. These results, together with the conserved role of *ct* in abdominal spiracle specification, support a model in which high *ct* activity establishes a spiracular fate field in T1, upon which an appendage GRN is superimposed. The transient pulse of *Dll* expression in the PROD, earlier than in the wing and coinciding with reduced *hth* expression, further suggests that this heterochronic shift in the P/D axis was instrumental in driving the distal respiratory protrusion on a spiracular foundation. Together, these findings demonstrate that the PROD evolved through the modular co-option of an appendage GRN onto a *ct*-defined spiracular field, a mechanism that reconciles its wing serial homolog and novel respiratory morphology.

Our cross-species functional analyses reveal how lineage-specific GRN recruitment contributes to the diversification of PROD. In both *D. melanogaster* (branched tracheal spiracle PROD) and *A. albopictus* (snorkel-like PROD), perturbation of appendage GRN components (*Dll*, *hth*, *exd*, *N*, *Dl*, *Ser*, *wg*/*Wnt*, *hh*, *en*, *ptc*, *ci*) and spiracle GRN components (*ct*, *tx*, *par-6*) produce stereotypical defects in PRODs, underscoring deep conservation of PROD core patterning logic. In contrast, the role of tracheal GRN components show lineage specificity. In *D. melanogaster*, *btl* acts upstream of *N/Dl* during tracheal-spiracle branching development, with *btl, N* and *Ser* knockdowns causing severe PROD defects. In *A. albopictus*, however, knockdown of *Aa-btl* or *Aa-trh* has no detectable effect on PROD morphogenesis, indicating that tracheal co-option is not required. These functional differences support a stepwise co-option model in which an ancestral PROD—built on a *ct*-specified spiracular field and patterned by an appendage GRN—was later elaborated in Schizophora through recruitment of tracheal GRNs.

These data also highlight the modularity of the underlying GRNs. In *D. melanogaster*, manipulating *N* or its two ligands decouple distal cone morphogenesis from tracheal branching, indicating that distinct aspects of PROD morphology—branch number, opening number, and cone length—are controlled by partially separable GRNs. Such modularity likely facilitated the independent gain of branched PRODs in particular lineages without compromising the core function of the organ. The repeated deployment of transcription factors such as *Dll* and *ct* across thoracic and abdominal imaginal tissues further illustrates spatiotemporal shifts in GRN activity can generate novel morphologies. More broadly, this is consistent with proposals that T1 wing serial homologs can serve as substrates for morphological innovation^1–3^, as seen in treehopper helmet and beetle horns^12,13^. Our work adds PROD to this class of co-opted structures and points to a specific regulatory combination—high *ct* activity coupled with a heterochronic *Dll* activation—that may represent one possible mechanism for developing protrusive organs from serially homologous fields.

Several important questions remain to be explored as the field moves toward a more complete understanding of PROD evolution. While our phylogenomic sampling is broad, transcriptional and functional data are currently concentrated in a few representative species, primarily *D. melanogaster* and *A. albopictus*. Expanding to basal nematocerans and non-model brachycerans will be essential for refining the timing and sequence of GRN recruitment across Diptera. The regulatory hierarchies within and between the spiracle, appendage, and tracheal modules also require deeper dissection: targeted perturbations, epistasis tests, and chromatin profiling will be needed to resolve gene regulatory logic and cross-module interactions. Moreover, while our data strongly support PROD-wing serial homology, recent work (e.g., larval prolegs in Lepidoptera^96^) underscores the need for more rigorous, comparative developmental analyses, including direct lineage tracing in additional taxa, are necessary to test the extent of developmental system conservation. Finally, integrating fossil evidence, direct fitness measurement and ecological data on larval and pupal habitats will be critical to move beyond correlation and assess whether environmental shifts directly shaped PROD diversification. Notably, the mutant phenotypes observed in *A. albopictus* and *D. melanogaster* recapitulate natural PROD morphologies found in unsampled dipteran lineages, suggesting that our functional analyses have uncovered not only developmental mechanisms but also plausible evolutionary paths. This offers testable hypotheses for how regulatory changes could generate the natural diversity of PROD form across the order (Fig. 10e).

Despite these open questions, our results establish a foundational *eco-evo-devo* model for the origin of a novel organ. The dipteran PROD arises not through de novo innovation, but through the stepwise co-option and reconfiguration of ancient GRNs. This modular assembly enabled morphological diversification without compromising core function, a principle likely shared across the evolution of morphological novelty. By tracing the developmental rewiring of a serial homolog across deep time, our work illustrates how new organs emerge from the layered integration of conserved programs, offering a general framework for understanding the origins of evolutionary innovation.

## Acknowledgement

We thank Dr. Tian Xu, Dr. Xianjue Ma, Dr. Xiaojun Xie, Dr. Siqian Feng, Dr. Hongjun Shi, and Dr. Yi Sun for their advice and support on genetic analyses; Dr. Jiangfan Xiu and Lei Liu for their help in establishing laboratory mosquito colony; Mingyuan Zhen and Tianzhu Guo for their assistance in lab animal rearing; and Xuhongyi Zheng for his suggestions on sample collection and analyses. We thank the Westlake Supercomputer Center for HPC and computation assistance.

## Author Contributions

Conceptualization: YY, XZ, XX, YZ

Supervision: YZ

Field investigation: YY, XZ, XX, ZJ

Phylogenetic analyses: XX, CT

Genetic and transcriptomic analyses: YY, XZ, XX, SL, JR

Writing: YY, XX, YZ

## Data availability

All sequencing data and genome assemblies for this study are available under BioProject: PRJNA1226360. The dataset includes PacBio HiFi reads, Illumina genome sequencing, Hi-C, RNA-seq, and scRNA-seq data. Genomic and phylogenetic datasets, comparative transcriptomic analyses, and functional phenotypic validation data are available on Figshare under the project titled “Stepwise co-option of gene networks drives morphological innovation in flies” and can be accessed via the following links: https://figshare.com/s/a70f1a5970f330e5c6d9; https://figshare.com/s/fb8d0451d9a108bf1250.

## Materials and Methods

### Fly and mosquito husbandry

All *D. melanogaster* stocks, experimental crosses, and developmental stage-specific analyses were maintained on standard cornmeal agar medium at 25°C with 60% relative humidity and a 12:12 hour light-dark cycle. The *D. melanogaster* stocks were obtained from stock centers and other labs (Table S6). *GAL4* driver lines were selected based on expression patterns relevant to the PROD disc, identified through prior studies and online databases (FlyBase.org, enhancers.starklab.org). For lineage-tracing experiments, the G-TRACE system (Gal4 technique for real-time and clonal expression) was implemented using a *UAS-RFP*, *Ubi-frt-stop-frt-GFP*, *UAS-FLP* genotype (BDSC28280). For analyses that requires staging, flies were staged by collecting eggs laid within a 2-hour window on apple juice agar plates supplemented with yeast paste to ensure synchronization of developmental stages.

*A. albopictus* colonies were obtained from Guizhou Medical University and kept under biosafety level 1 guidelines in an isolated insectary equipped with a double-door entry system to prevent accidental release. Mosquitoes were reared at 27°C with 80% relative humidity and a 12:12 hour light-dark cycle. Larvae were housed in plastic trays filled with dechlorinated water, fed daily with finely ground fish food, and maintained at a density of ∼200 larvae per liter to ensure uniform development. Pupae were collected daily and transferred to fine-mesh cages (<0.5 mm mesh) for emergence, where adults were provided with 10% (w/v) sterile sucrose solution. Two feeding methods were used: defibrinated sheep blood (#R54016, Thermo Fisher Scientific, Waltham, MA, U.S.A.); live mouse blood under approved Institutional Animal Care and Use Committee (IACUC) protocols (No. AP#22-089-ZY), adhering to national ethical guidelines of China. Gravid females were transferred to oviposition cages with wet filter paper and dechlorinated water. Eggs were collected on wet filter paper over a 24-hour period, air-dried at room temperature for 48 hours, and hatched synchronously by submerging them in deoxygenated water prepared by boiling and cooling.

### *ct.E-GAL4* Plasmids and Transgenesis

The *ct.E* enhancer fragment was amplified from Canton-S genomic DNA using forward primer (5’-GAATTCTGAGAAACTTGGGTGTCC-3’) and reverse primer (5’-GAATTCCATCGCACAATGCA-3’). The PCR products were purified and inserted into the *pBPGAL4.2::VP16Uw* vector (gift from Dr. Yi Sun), upstream of the *GAL4* promotor, using the ClonExpress Ultra One Step Cloning Kit V2 (#C116-01, Vazyme, Nanjing, Jiangsu, China) following the manufacturer’s instructions. The clone products were transformed into competent *E. coli* DB3.1 cells (#D1025S, Beyotime Biotechnology, Shanghai, China) and positive clones were confirmed by sequencing. The attP-PhiC31-mediated transgene of verified recombinant *ct.E-GAL4* plasmid were performed at Fungene Biotech (http://www.fungene.tech) using the BDCS25709 background fly with the *X* chromosome cross switched to *w,nos-phiC31*. Transgenic flies were identified using *w* eye color rescue. The activity of *ct.E-GAL4* was verified by crossing with *UAS-GFP*.

### Sample Collection and Identification

Specimens of *De. wuyiensis* were collected from the Wuyi Mountains in Fujian Province, China (27°44′55.52″ N, 117°40′40.77″ E and 27°52′13.33″ N, 117°51′48.69″ E), while *Philorus* sp. specimens were obtained from Yaowang Mountain in Quzhou, Zhejiang Province, China (28.75714611° N, 118.97632728° E). Both locations feature fast-flowing, rocky riverbeds, which are typical habitats for these species during their larval and pupal stages. Larvae and pupae were collected from submerged stones. Immediately after collection, the specimens were placed into plastic petri dishes containing water from their respective habitats. The specimens were preliminarily sorted and identified on site under a SZ810 stereomicroscope (CNOPTEC, Chongqing, China) by examining species-specific morphological characteristics. Distinctive traits, such as ornamental body patterns and unique respiratory organ structures, were used to differentiate between *De. wuyiensis* and *Philorus* specimens (refer to Zheng et al. (2022) for detailed protocols ^97^). Following sorting, the fresh samples were temporarily preserved in sealed containers with dry ice during transportation and later stored at -80°C in the laboratory.

For species identification, DNA was extracted from a subset of specimens, and a fragment of the *cox1* gene was amplified using standard PCR ^98,99^. The amplified DNA fragments were sequenced and compared to GenBank and BOLD databases for species confirmation. Specimens were preserved in 95% ethanol, RNA-later (#AM7021, Thermo Fisher Scientific, Waltham, MA, U.S.A.), or on dry ice for subsequent sequencing and morphological examination.

### Second-Generation Sequencing Workflow

DNA was extracted using CTAB method from pooled samples of each species. DNA integrity was confirmed through agarose gel electrophoresis, and quantified using a Qubit 3.0 fluorometer (Thermo Fisher Scientific, Waltham, MA, U.S.A.). Genomic DNA was fragmented using a Covaris sonicator (Covaris, Woburn, MA, U.S.A.) and sequencing libraries were prepared using the Nextera DNA Flex Library Prep Kit (Illumina, San Diego, CA, U.S.A.). The final libraries have average insert sizes of ∼150 bp. Sequencing was performed on the NovaSeq 6000 platform (Illumina, San Diego, CA, U.S.A.). A total of 27.95 GB and 14.98 GB of clean data were obtained for *De. wuyiensis* and *Philorus* sp., respectively, corresponding to 189,413,324 pairs of reads for *De. wuyiensis* and 102,610,577 pairs of reads for *Philorus* sp.

### Third-Generation SMRT Sequencing Workflow

High-quality genomic DNA was extracted from pooled samples collected in bulk. The DNA was fragmented using the Megaruptor system (Diagenode, Belgium) to achieve fragment sizes of approximately 15-20 kb, followed by end repair and adapter ligation to construct a PCR-free SMRTbell library. The SMRTbell libraries were quantified and subjected to HiFi sequencing on the PacBio Revio platform (Pacific Biosciences, Menlo Park, CA, U.S.A.). Raw sequencing data were stored in the subreads.bam format and processed using PacBio’s SMRTlink v12.0 software (Pacific Biosciences, Menlo Park, CA, U.S.A.). Post-sequencing, a total of 25.61 Gb and 49.83 Gb of high-quality clean data were generated for *De. wuyiensis* and *Philorus* sp., respectively. The average read lengths were 15.27 kb and 17.24 kb, with maximum read lengths up to 44.64 kb and 47.04 kb for the two species, respectively.

### Hi-C Library Preparation and Analysis

For Hi-C library construction, fresh tissue samples were cross-linked, digested with DPN II, and ligated to preserve chromatin interactions. The DNA was sheared, biotinylated, and purified to capture interaction pairs. Library quality was assessed using Qubit 3.0 Fluorometer (Life Technologies, Carlsbad, CA, U.S.A.) and Agilent 2100 Bioanalyzer (Agilent Technologies, Santa Clara, CA, U.S.A.). Effective library concentration was quantified by qPCR. Libraries were sequenced on the NovaSeq X Plus (Illumina, San Diego, CA, U.S.A.), and raw data were processed with HICUP (v0.8.0) ^100^ to extract Valid Interaction Pairs (VIPs) for anchoring and scaffolding contigs. For *De. wuyiensis*, Hi-C sequencing generated 33,908,871 read pairs, with 8,354,110 VIPs identified. For *Philorus* sp., 123,320,854 read pairs were produced, yielding 35,902,009 VIPs.

### Transcriptomes Sequencing for Genome Annotation

Transcriptomes of *De. wuyiensis* and *Philorus* sp. were sequenced to facilitate transcript-based annotation. For *De. wuyiensis*, samples included the whole body of larvae and pupae, as well as the PROD and wing discs from both the early stage of the final instar and the final instar larvae. For *Philorus* sp., the whole body of final instar larvae was used. Total RNA was extracted using the TRIzol reagent (#15596018CN, Thermo Fisher Scientific, USA). RNAseq libraries were constructed using TruSeq RNA and DNA Library Preparation Kits v2 (Illumina, Inc., San Diego, CA, U.S.A.), following the manufacturer’s protocols. Libraries were sequenced on the Illumina Novaseq 6000 platform (Illumina, San Diego, CA, U.S.A.). A total of 55.8 GB and 2.52 GB of clean RNA-seq data were obtained for *De. wuyiensis* and *Philorus* sp., respectively.

### Genome Assembly

The genome assembly was initiated using Hifiasm (v0.19.8) ^101,102^ for long-read sequencing data. To remove potential redundancies inherent in diploid or polyploid genomes, Purge_dups (v1.2.6) ^103^ and Purge_Haplotigs (v1.1.2) ^104^ were employed sequentially. Subsequent error correction was performed with Pilon (v1.24) ^105^, leveraging high-accuracy second-generation DNA sequencing data to rectify base-level errors and improve the overall quality of the assembly. Quast (v5.2.0) ^106^ was used to evaluate the quality of genome assembly by integrating both second- and third-generation sequencing data.

To achieve chromosome-level scaffolding, two complementary approaches were utilized: Juicer combined with 3D-DNA (v180922) ^107^, and AllHIC (v0.9.8) ^108^. These pipelines integrated Hi-C interaction data to order and orient contigs into chromosome-scale scaffolds. The obtained assemblies were visualized and manually curated using Juicebox (v2.20.00) ^109^, allowing for the identification and correction of mis-assemblies based on the consistency and quality of Hi-C contact maps. This meticulous manual adjustment resulted in a highly accurate and contiguous genome, suitable for downstream analyses.

### Genome Structural Annotation

Genome structural annotation was performed using two complementary strategies. The first approach is de novo, transcript- and homology-based annotation. The genome was first soft-masked using EarlGrey (v5.0.0) ^110–112^. BRAKER3 (v3.0.8)^113^ was employed for de novo gene prediction,, integrating RNA-Seq data to enhance annotation accuracy. Transcript-based annotation was conducted using both the reference-free PASA pipeline (v2.5.3)^114,115^ and a reference-guided approach using HISAT2 (v2.2.1) ^116^, StringTie (v2.2.1) ^117^, and TransDecoder (v5.7.1) ^118^. Homology-based annotation was conducted using GeMoMa (v1.7.1) ^119,120^ and miniprot (v0.13) ^121^, leveraging protein alignments from closely related species including *A. aegypti*, *Chironomus riparius*, *D. melanogaster*, and *Polypedilum vanderplanki*. The annotations from these sources were then consolidated using MAKER (v3.01.03) ^122^ or EvidenceModeler (v2.1.0) ^123^. The quality and completeness of the annotations were subsequently evaluated using BUSCO (v5.0.0) ^124^.

The second strategy employed EGAPx (v0.3.2) ^125^, the publicly accessible version of the updated NCBI Eukaryotic Genome Annotation Pipeline. The assembled genome FASTA file, the organism’s taxonomic identifier (taxid: 7147, Diptera), and corresponding RNA-seq data were provided as inputs. EGAPx selected appropriate protein sets and Hidden Markov Model (HMM) profiles based on the taxid. Protein sequences were aligned to the assembly using miniport ^121^, while RNA-seq reads were aligned using STAR ^126^. These alignments were subsequently processed by Gnomon ^127^ for gene prediction. In the initial phase, Gnomon chained short alignments into putative gene models, which were then augmented with ab initio predictions derived from HMM models.

Based on BUSCO assessment, the final genome annotations of *De. wuyiensis* and *Philorus* sp. were determined using EGAPx and manual strategies, respectively.

### Genome Functional Annotation

Following structural annotation, functional annotation of the predicted genes was conducted. EggNOG-Mapper (v5.0)^128,129^ was utilized to assign functional categories based on orthology relationships, leveraging the EggNOG database to predict gene functions, metabolic pathways, and potential protein domains. This tool classifies genes into Clusters of Orthologous Groups (COGs), enabling the identification of conserved functional roles within the Insecta taxonomic scope (Taxonomy ID: Insecta, 50557). Additionally, PANTHER (Protein ANalysis THrough Evolutionary Relationships, v14)^130^ was employed to further annotate gene functions, focusing on Gene Ontology (GO) terms and protein families. PANTHER provided insights into the molecular functions, biological processes, and cellular components associated with each gene, enhancing the functional landscape of the annotated genome. By integrating results from both EggNOG-Mapper and PANTHER, a comprehensive functional annotation was achieved, supporting downstream analyses such as pathway enrichment, gene network construction, and comparative genomics.

### Transcriptomes Assembly for Phylogenetic Analysis

All RNA-seq data used for phylogenic analysis in this study was first assessed using FastQC (v0.12.0)^131^ to evaluate quality. Low-quality bases and adapter sequences were removed using Trimmomatic (v 0.39)^132^, with parameters: LEADING:5, TRAILING:5, SLIDINGWINDOW:4:20, MINLEN:36. Cleaned reads were then assembled *de novo* using Trinity (v2.14)^133^. The longest isoform for each gene was extracted using Trinity’s get_longest_isoform_seq_per_trinity_gene.pl script, providing representative sequences for phylogenetic matrix construction.

### Phylogenetic analysis

Species included were selected by three main criteria: 1) broad taxonomic representation at the family level, ensuring the inclusion of families within Diptera with publicly available genomic or transcriptomic data; 2) data quality, prioritizing species with high BUSCO completeness (>90%) and high-quality assemblies; and 3) irreplaceable samples, where species with lower BUSCO completeness were selected due to their unique phylogenetic position and the limited availability of data. This approach resulted in the selection of 235 dipteran species, which collectively provide comprehensive coverage of dipteran diversity (Table S1).

To identify single-copy orthologs (USCOs), genomic and transcriptomic data were aligned to the diptera_odb10 dataset from BUSCO (v5.0.0) ^124^ using the workflow detailed in the “script2_BUSCO_extraction.sh” of PLWS (v1.0.6) ^134^. This process identified 1,404 universal single-copy genes (USCOs) shared across at least 80% of the selected species.

USCOs were translated into protein sequences using TransDecoder (v5.5.0) ^118^ and aligned with MAFFT (v7.520) ^135^ under default parameters. Conserved regions were retained, and low-quality or highly variable regions were trimmed using TrimAl (v1.4.1)^136^ with the ‘automated1’ algorithm. The trimmed protein sequences were concatenated into a single dataset using FASconCAT-G (v1.05.1)^137^, resulting in a final matrix for phylogenetic analysis.

### Fossil calibration and molecular dating

We selected 47 representative dipteran species and performed MCMCtree analyses using the phylogeny constructed with IQ-TREE, estimating divergence times with 12 fossils as calibration points. To assess the robustness of our estimates, we tested three calibration schemes: (1) a uniform distribution with soft bounds (2.5% minimum and maximum), (2) a skew-normal distribution assigning higher probability near the minimum bound (assuming it closely approximates clade divergence), and (3) a Cauchy distribution with heavy tails, accommodating more extreme age estimates (Supplementary Figs. S3-S5).

Molecular dating analyses were performed using MCMCtree(v4.9) ^138^ with the LG+G amino acid substitution model. We estimated evolutionary rates via the mean substitution rate parameter (rgene_gamma) according to the chosen tree topology and calibration scheme, and set the prior for rate variance (σ²) as Gamma(1,10) to allow violations of strict clock assumptions.

Each analysis ran for 20,000 iterations, following a burn-in of 100,000 iterations and a sampling frequency of 1,000. Convergence was confirmed across five independent chains when effective sample sizes exceeded 200 and posterior means were consistent. The final consensus tree was derived by averaging converged chains, and all age estimates are reported with 95% highest posterior density (HPD) intervals.

### Ancestral-state reconstruction

Ancestral-state reconstruction was conducted using the phylogeny from IQ-TREE (amino acid data) combined with PROD morphological data. To infer the ancestral states, we employed maximum likelihood (ML) methods using the fitMk function in phytools (v2.4-4)^139,140^, comparing three discrete character models: equal-rates (“ER”), symmetric rate (“SYM”), and all-rates-different (“ARD”). The optimal model, determined by Akaike Information Criterion (AIC), was used for ancestral-state reconstruction. Stochastic mapping was performed using the make.simmap function, conducting 2,000 simulations under a Markov Chain Monte Carlo (MCMC) framework with a burn-in of 10,000 and sampling every 50 iterations. This approach estimated posterior probabilities of ancestral states, ensuring robust and consistent results.

### Tissue-Specific RNA Sequencing and analysis

For tissue-specific transcriptomic analysis, PROD and wing tissues were dissected from *De. wuyiensis* and *A. albopictus*. The developmental stage was empirically staged based on the observable morphology of the imaginal discs. Larvae with large, slightly to fully pigmented discs were classified as late-stage prepupating individuals (∼5 days since hatching); while those with smaller, milky-white discs were considered early-stage individuals (∼4 days since hatching).

Dissections were performed under a stereomicroscope in cold phosphate-buffered saline (PBS) on ice. For both species, 5-10 pairs of tissues were pooled per replicate to ensure sufficient RNA yield. Dissected tissues were flash-frozen in liquid nitrogen and stored at −80°C until RNA extraction.

For *D. melanogaster*, PROD and wing discs were dissected from individual specimens at defined developmental stages. Dissections were performed under a SZ810 stereomicroscope (CNOPTEC, Chongqing, China) in cold PBS on ice. Two to three pairs of imaginal discs were collected per sample at early stages (84 and 96 hours after egg laying, AEL) and late stages (108 and 120 hours AEL). The dissected tissues were immediately transferred into MACS Tissue Storage Solution (Miltenyi Biotec, Cologne, NRW, Germany) using a bent-tip needle to minimize the introduction of excess fluid, and were stored at −80°C until RNA extraction.

For bulk RNA-seq, total RNA was extracted from dissected tissues or whole-body samples of *De. wuyiensis* and *A. albopictus* using Eastep Super Total RNA Extraction Kit (#LS1040, Promega Corporation, Madison, WI, U.S.A.). RNAseq libraries were prepared using TruSeq RNA and DNA Library Preparation Kits v2 (Illumina, Inc., San Diego, CA, U.S.A.), following the manufacturer’s protocols. The libraries were sequenced on a NovaSeq 6000 platform (Illumina, Inc., San Diego, CA, U.S.A.), generating 150-bp paired-end reads. For SMART-seq2, the *D. melanogaster* samples were processed using SMART-Seq Single Cell Kit (Takara, Tokyo, Japan) according to the manufacturer’s recommendations. The cDNA products were heat-denatured and circularized using a splint oligo sequence to form single-strand circular DNAs as the final library, which was sequenced on a DNBSEQ sequencing platform (BGI Genomics, Shenzhen, Guangdong, China), generating 100-bp paired-end reads.

### Transcriptome Differential Expression Analysis

Quality control of raw RNA-seq data was performed using FastQC (v0.12.0) ^131^ to evaluate basic quality metrics, followed by trimming with Trimmomatic (v 0.39) ^132^ to remove adapter sequences and low-quality bases. The following parameters were used for trimming: LEADING:5, TRAILING:5, SLIDINGWINDOW:4:20, MINLEN:36. For alignment, HISAT2 (v2.2.1) ^116^ was used to align the trimmed reads to the respective reference genomes: GCF_000001215.4 for *D. melanogaster*, GCA_018104305.1 for *A. albopictus*, GCA_023509865.1 for *T. dichotomus*, GCA_036711975.1 for *O. taurus*, GCA_021130785.2 for *H. vitripennis*, and our newly-assembled genome of *De. wuyiensis*. Following alignment, StringTie (v2.2.1) ^117^ was used to assemble transcripts and estimate their expression levels. To generate gene expression counts for differential expression analysis, the script prepDE.py from StringTie was used. For *E. carinata* quantification, the reference transcriptome^14^ were used for index; and kallisto^141^ and sleuth^142^ pipeline was used to obtain gene count table. Differential expression analysis was carried out using DESeq2 (v1.46.0) ^143^, and plots were generated using ggplot2 (v3.5.1) ^144^ in R. Gene Ontology (GO) enrichment analysis was performed using the ClusterProfiler package (v4.14.4) ^145,146^. Pathway enrichment analysis was conducted using the Kyoto Encyclopedia of Genes and Genomes (KEGG) database to identify enriched signaling pathways.

For cross-species comparison of differentially expressed genes, we used Orthofinder2 (v2.5.5) ^68^ to find orthologous genes across the seven species. Using the “metagene” method in Geirsdottir et al. 2019^67^, we used *D. melanogaster* genes as the reference to breakdown the shared Orthogroups of the other 6 species into metagenes to establish the one-to-one relationships of genes between species. For recently evolved paralogs in *D. melanogaster*, we kept the more conserved copy with shorter branch length in Orthofinder2 gene tree. We proceeded cross-species comparison using metagene as unit by adding the transcript counts within a metagene for the rest of the 6 species.

### 10X Genomics scRNA-seq analysis

For 10X Genomics scRNA-seq, *D. melanogaster* PROD discs were dissected from larvae at 120 hrs AEL. Dissections were performed under SZ810 stereomicroscopes (CNOPTEC, Chongqing, China) in cold DNase, RNase, and Protease-free, sterile PBS (#ST478-500ml, Beyotime Biotechnology, Shanghai, China). To preserve cell viability, the dissected tissues were immediately placed into ice-cold MACS Tissue Storage Solution (#130-100-008, Miltenyi Biotec, Cologne, NRW, Germany) and kept on ice throughout the procedure. Approximately 600 PROD discs from over 300 larvae, corresponding to around 600,000 cells, were pooled to guarantee an adequate cell number.

The tissue was then digested in 1 mg/mL collagenase (#C0130-100MG, MilliporeSigma, St. Louis, MO, U.S.A.) and 1 mg/mL papain (#10108014001; MilliporeSigma, St. Louis, MO, U.S.A.). Once dissociation was complete, the cell suspension was filtered through a 40 µm cell strainer (Corning, New York, U.S.A.) to remove clumps and debris, ensuring a single-cell population. The resulting suspension was counted using a hemocytometer and adjusted to a concentration of approximately 1,000-3,000 cells/µL. Cells were processed using the 10X Genomic microfluidics system at Lianchuan (https://www.lc-bio.com). scRNA-seq libraries were generated using the Chromium Single Cell 3’ Reagent Kit v3 following the manufacturer’s protocol (10X Genomics, Pleasanton, CA, U.S.A.). The quality of the final libraries was evaluated using an Agilent Bioanalyzer 2100 (Agilent Technologies, Santa Clara, CA, U.S.A.). Libraries were then sequenced on an Illumina NovaSeq 6000 platform with paired-end reads.

### scRNAseq Data Processing and Analysis

Raw sequencing data in FASTQ format were aligned to *D. melanogaster* reference genome (GCF_000001215.4) and processed using Cell Ranger (v6.0.0) (10X Genomics, Pleasanton, CA, U.S.A.). A gene expression matrix was generated and further analyzed using Seurat 4 ^147^ in R. The analysis pipeline in Seurat 4 began with quality control to filter out low-quality cells. Cells with a low number of detected genes or a high percentage of mitochondrial gene expression (>10%) were excluded from the dataset. The data were then normalized using the LogNormalize method, which account for differences in sequencing depth across cells. Next, highly variable genes were identified using FindVariableFeatures function, which was then followed by dimensionality reduction using Principal Component Analysis (PCA). The top principal components were selected for clustering, and the cells were grouped into distinct clusters using the Louvain algorithm in Seurat. These clusters were visualized using UMAP and t-SNE. Differential expression analysis was performed using Seurat’s FindMarkers function, which identified marker genes specific to each cluster.

### Imaginal disc dissection and immunostaining

*D. melanogaster* imaginal discs were dissected from larvae under a SZ810 stereomicroscope (CNOPTEC, Chongqing, China) in cold phosphate-buffered saline (PBS; #C0221A, Beyotime Biotechnology, Shanghai, China). The excised PROD and wing discs were transferred with a bent-tip needle into ice-cold 4% paraformaldehyde fixative solution (#P0099, Beyotime Biotechnology, Shanghai, China) and incubated for 20 minutes at room temperature to ensure optimal fixation. For *D. melanogaster* embryo dissection, the eggs were mounted on glass slide and chorion was crack open by a blunt-ended forceps. The embryos with vitelline were then mounted on the tape and then applied with 4% paraformaldehyde fixative solution. After incubation for 20 minutes at room temperature, vitelline was removed with a sharp needle and blunt-ended forceps.

After fixation, the tissues were washed three times with PBS and subsequently permeabilized with PBST (1 L PBS with 3 mL Triton X-100; #T8787, MilliporeSigma, St. Louis, MO, U.S.A.) for 30 minutes at room temperature. For immunostaining, the tissues were treated with a blocking solution (100 mL PBST with 5 g bovine serum albumin, #ST2249, Beyotime Biotechnology, Shanghai, China; 5 mL normal goat serum, #C0265, Beyotime Biotechnology, Shanghai, China) for 1 hour at room temperature to reduce non-specific binding. Primary antibodies were diluted in the blocking solution and applied to the samples overnight at 4°C. After the primary antibody incubation, the tissues were washed three times with PBST and incubated with secondary antibodies for 1 hour at room temperature in the dark. Following secondary antibody incubation, the tissues were washed again three times with PBST. The antibodies used in this study are listed in Supplementary Table S7. The processed samples were mounted on slides with ProLong Gold Antifade Mountant with DNA Stain DAPI (#P36931, Thermo Fisher Scientific, Waltham, MA, U.S.A.) for nuclear staining. Z-stack confocal images were acquired using a FV3000 confocal laser scanning microscope (Olympus Corporation, Tokyo, Japan) and subsequently analyzed using Fiji^148^.

### *D. melanogaster* PROD phenotyping

Wandering *D. melanogaster* larvae 120 hrs AEL were dissected in PBS under a stereomicroscope to remove internal tissues while preserving the epidermis and tracheae. The dissected samples were immersed in 5% lactic acid (#V000448, MilliporeSigma, St. Louis, MO, U.S.A.) at room temperature for 10-15 minutes to clear tissues and improve visibility of tracheae. Specimens were then transferred to a glass slide containing glycerol. Using a fine-tipped probe, tissues were gently flattened with glycerol to make the tracheal branches visible and to avoid excessive distortion. The number of PROD branches of the samples were examined and imaged under an Olympus CKX53 microscope (Olympus Corporation, Tokyo, Japan). Pupae were collected and fixed in 95 % ethanol and their PROD size were measured and imaged under an Olympus SZX16 stereomicroscopes equipped with software CellSens Dimension 3.1.1 (Olympus Corporation, Tokyo, Japan).

### Design and Synthesis of dsRNA

The relevant *A. albopictus* ortholog sequences were used to design dsRNA primers that minimize off-target effects. Potential siRNA sites were identified with the DSIR tool (biodev.extra.cea.fr/DSIR). The primers were designed in low off-target risk regions (∼200-500 nt)(Table S9).

Each dsRNA target region included T7 promoter sequences on both forward and reverse primers. The transcription reaction was carried out using a T7 High Yield RNA Transcription Kit (#TR101, Vazyme Biotech Co., Ltd., Nanjing, Jiangsu, China), following the manufacturer’s protocol. The dsRNA products were purified with VAHTS RNA Clean Beads (#N412, Vazyme Biotech Co., Ltd., Nanjing, Jiangsu, China), resuspended in nuclease-free water at 5 µg/µL, and stored at -80°C until use.

### dsRNA Injection and Electroporation

Larval dsRNA injections were performed under a stereomicroscope (Nikon SMZ 745T, Nikon, Tokyo, Japan) using an FemtoJet 4i microinjector system with a TransferMan 4r Micromanipulator (Eppendorf AG, Hamburg, Germany). Each larva received 0.1 µL of dsRNA solution (500 ng per larva) injected into the sclerotized head capsule, where the cuticle’s resilience prevents leakage, ensuring high efficiency. Electroporation was performed immediately after injection using an ECM 830 Square Wave Electroporation System (Harvard Bioscience, Holliston, MA, USA), with electrodes applied to the larval thorax. Based on previous reports ^149–154^ and our preliminary tests, we optimized an electroporation protocol with five pulses at 100 ms intervals and 900 ms rest intervals. Voltage levels of 7V, 9V, 12V, 15V, and 20V were tested, and 7V was found to cause no physical damage to the larvae. Thus, all subsequent RNAi experiments were conducted at 7V for electroporation. To assess gene knockdown, surviving larvae were collected 24 hours after injection and electroporation for qPCR. For each group, 4 replicates were prepared, with 10 larvae per replicate. Immediately after collection, larvae were flash-frozen in liquid nitrogen and stored at -80°C to preserve RNA integrity.

Individuals (pupa, prepupa that failed to pupate) were preserved in 95% ethanol for endpoint observation. The prepupa were dissected to remove the larval cuticle. The pupal phenotypes of PROD, wing, leg, antennae, and compound eye are assessed and compared to the *D. melanogaster* studies. The *A. albopictus* PRODs were measured and imaged under an Olympus SZX16 stereomicroscopes equipped with software CellSens Dimension 3.1.1 (Olympus Corporation, Tokyo, Japan) and analyzed in R.

### qPCR and Data Analysis

Total RNA was extracted and DNase-treated using Eastep Super Total RNA Extraction Kit following the manufacturer’s protocol (#LS1040, Promega Corporation, Madison, WI, U.S.A.). 1 µg of total RNA from each sample was reverse transcribed into cDNA using Eastep RT Master Mix (#LS2050, Promega Corporation, Madison, WI, U.S.A.), following the manufacturer’s instructions.

Gene-specific primers were listed in Table S10. qPCR experiments were performed using ChamQ Universal SYBR qPCR Master Mix (#Q711-02, Vazyme Biotech Co., Ltd., Nanjing, Jiangsu, China) in a qTOWERiris qPCR system (Analytik Jena, Jena, Thuringia, Germany). Each run began with an initial denaturation at 95°C for 5 minutes, followed by 40 cycles of 95°C for 15 seconds and 60°C for 30 seconds. The normalized expression levels were calculated using the 2^(-ΔΔCt) method relative to the reference gene *ribosomal protein l32* (*Aa-rpl32*) as suggested by previous *A. albopictus* study ^155^. Statistical analyses were conducted using GraphPad Prism (GraphPad Software, San Diego, CA, U.S.A.).

