## supplemental text, figures and tables for "Stepwise GRN co-option in the evolution of a dipteran respiratory organ"

### Supplementary Materials

#### Supplementary Notes on genome assemblies of *De. wuyiensis* and *Philorus* sp.

We combined third generation sequencing, HiC sequencing and second generation sequencing data to assemble the genomes of *Deuterophlebia wuyiensis* and *Philorus* sp. (Supplementary information, Fig. S1 and Table S3). For *De. wuyiensis*, a total of 25.61 Gb of third-generation sequencing data (average coverage depth: 233×) and 27.95 GB of second-generation sequencing data (average coverage depth: 434×) were generated. For *Philorus* sp., 49.83 GB of third-generation sequencing data (average coverage depth: 492×) and 14.98 GB of second-generation sequencing data (average coverage depth: 223×) were obtained.

We obtained near-chromosome genome assemblies of both species. The assembled genome of *De. wuyiensis* is 125.47 Mb, comprises of 37 scaffolds, with a N50 of 30.03 Mb. The assembled genome of *Philorus* sp. is 128.99 Mb with 76 contigs, a N50 of 29.07 Mb. Both genomes have similar GC content as *D. melanogaster*. BUSCO analysis confirms the high completeness of both assemblies, with 96.9% and 96.6% complete BUSCOs for *De. wuyiensis* and *Philorus* sp., respectively (Supplementary information, Table S3).

*De. wuyiensis* exhibits a total repetitive element content of 26.72% (33.52 Mb), including retroelements constituting 8.14% of the genome. Among retroelements, LTR elements are the most abundant (6.52%), while LINEs account for 1.61%. In contrast, *Philorus* sp. shows a lower repetitive content (15.56%, 20.07 Mb), dominated by LTR retroelements (8.12%). These differences suggest distinct evolutionary dynamics in repetitive element accumulation between the two species (Fig 2a; Supplementary information, Table S3).

The number of genes annotated in genomes of *De. wuyiensis* and *Philorus* sp. are 10,586 and 12,774, respectively. Gene structure analysis reveals notable differences in gene length and intron-exon organization between the two species. *De. wuyiensis* features an average gene length of 5,232 bp, with exons and introns averaging 257 bp and 308 bp, respectively. while *Philorus* sp. shows slightly smaller genes (4,134 bp) and longer exons (293 bp).

In summary, we provide high-quality genomic resources for *De. wuyiensis* and *Philorus* sp., representing the first genome assemblies for their respective families (Deuterophlebiidae and Blephariceridae), with Deuterophlebiidae being a basal lineage of Diptera based on morphological and molecular evidence, and Blephariceridae previously proposed as closely related to Deuterophlebiidae in morphological studies. These genomic resources offer a solid foundation for future comparative genomic and evolutionary developmental studies.

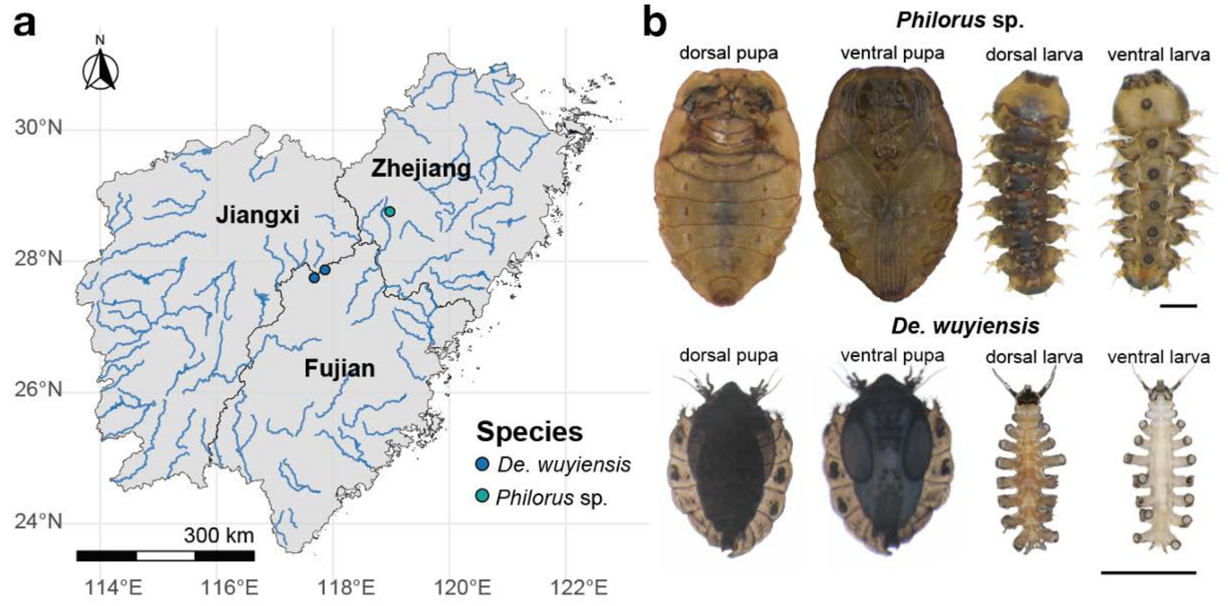


**Figure S1. Two high quality genomes of *De. wuyiensis* and *Philorus* sp.** (**a**) Distribution of the sampled species in Zhejiang and Fujian provinces, China. Blue lines are water systems. (**b**) Images of the sampled species. Scale bars = 1.0 mm. The color in the heatmap represents the log value of the interaction strength of the corresponding genome bin pair.

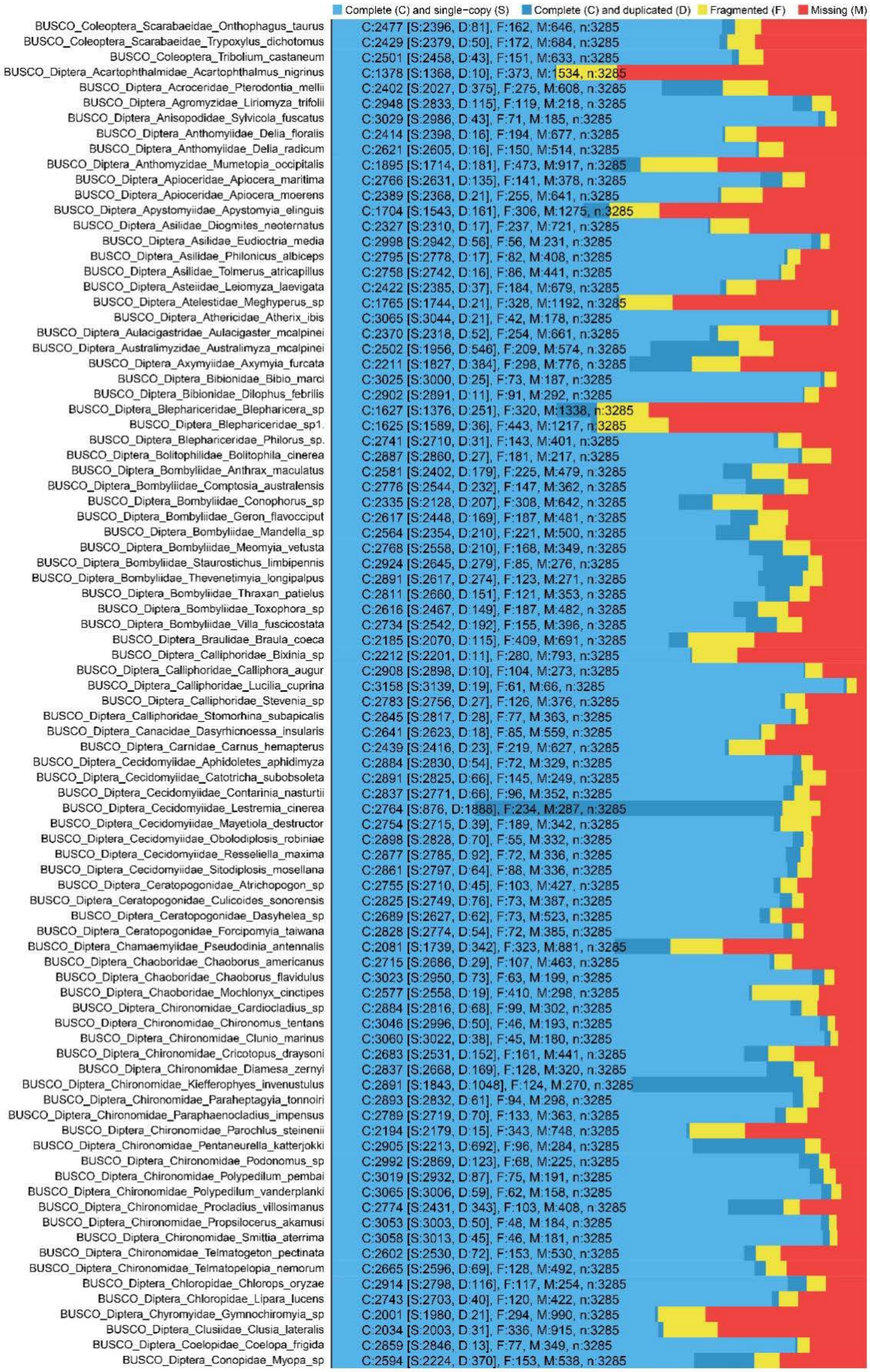
Complete (C) and single-copy (S)
Complete (C) and duplicated (D)
Fragmented (F)
Missing (M)
BUSCO_Coleoptera_Scarabaeidae_Onthophagus_taurus C:2477 [S:2396, D:81], F:162, M:646, n:3285
BUSCO_Coleoptera_Scarabaeidae_Trypoxylus_dichotomus C:2429 [S:2379, D:50], F:172, M:684, n:3285
BUSCO_Coleoptera_Tribolium_castaneum C:2501 [S:2458, D:43], F:151, M:633, n:3285
BUSCO_Diptera_Acartophthalmidae_Acartophthalmus_nigrinus C:1378 [S:1368, D:10], F:373, M:1534, n:3285
BUSCO_Diptera_Acroceridae_Pterodontia_mellii C:2402 [S:2027, D:375], F:275, M:608, n:3285
BUSCO_Diptera_Agromyzidae_Liriomyza_trifolii C:2948 [S:2833, D:115], F:119, M:218, n:3285
BUSCO_Diptera_Anisopodidae_Sylvicola_fuscatus C:3029 [S:2986, D:43], F:71, M:185, n:3285
BUSCO_Diptera_Anthomyiidae_Delia_floralis C:2414 [S:2398, D:16], F:194, M:677, n:3285
BUSCO_Diptera_Anthomyiidae_Delia_radicum C:2621 [S:2605, D:16], F:150, M:514, n:3285
BUSCO_Diptera_Anthomyzidae_Mumetopia_occipitalis C:1895 [S:1714, D:181], F:473, M:917, n:3285
BUSCO_Diptera_Apioceridae_Apiocera_maritima C:2766 [S:2631, D:135], F:141, M:378, n:3285
BUSCO_Diptera_Apioceridae_Apiocera_moerens C:2389 [S:2368, D:21], F:255, M:641, n:3285
BUSCO_Diptera_Apystomyiidae_Apystomyia_elinguis C:1704 [S:1543, D:161], F:306, M:1275, n:3285
BUSCO_Diptera_Asilidae_Diogmites_neoternatus C:2327 [S:2310, D:17], F:237, M:721, n:3285
BUSCO_Diptera_Asilidae_Eudioctria_media C:2998 [S:2942, D:56], F:56, M:231, n:3285
BUSCO_Diptera_Asilidae_Philonicus_albiceps C:2795 [S:2778, D:17], F:82, M:408, n:3285
BUSCO_Diptera_Asilidae_Tolmerus_atricapillus C:2758 [S:2742, D:16], F:86, M:441, n:3285
BUSCO_Diptera_Asteiidae_Leiomyza_laevigata C:2422 [S:2385, D:37], F:184, M:679, n:3285
BUSCO_Diptera_Atelestidae_Meghyperus_sp C:1765 [S:1744, D:21], F:328, M:1192, n:3285
BUSCO_Diptera_Athericidae_Atherix_ibis C:3065 [S:3044, D:21], F:42, M:178, n:3285
BUSCO_Diptera_Aulacigastridae_Aulacigaster_mcalpinei C:2370 [S:2318, D:52], F:254, M:661, n:3285
BUSCO_Diptera_Australimyzidae_Australimyza_mcalpinei C:2502 [S:1956, D:546], F:209, M:574, n:3285
BUSCO_Diptera_Axymyiidae_Axymyia_furcata C:2211 [S:1827, D:384], F:298, M:776, n:3285
BUSCO_Diptera_Bibionidae_Bibio_marci C:3025 [S:3000, D:25], F:73, M:187, n:3285
BUSCO_Diptera_Bibionidae_Dilophus_febrilis C:2902 [S:2891, D:11], F:91, M:292, n:3285
BUSCO_Diptera_Blephariceridae_Blepharicera_sp C:1627 [S:1376, D:251], F:320, M:1338, n:3285
BUSCO_Diptera_Blephariceridae_sp1. C:1625 [S:1589, D:36], F:443, M:1217, n:3285
BUSCO_Diptera_Blephariceridae_Philorus_sp. C:2741 [S:2710, D:31], F:143, M:401, n:3285
BUSCO_Diptera_Bolitophilidae_Bolitophila_cinerea C:2887 [S:2860, D:27], F:181, M:217, n:3285
BUSCO_Diptera_Bombyliidae_Anthrax_maculatus C:2581 [S:2402, D:179], F:225, M:479, n:3285
BUSCO_Diptera_Bombyliidae_Comptosia_australensis C:2776 [S:2544, D:232], F:147, M:362, n:3285
BUSCO_Diptera_Bombyliidae_Conophorus_sp C:2335 [S:2128, D:207], F:308, M:642, n:3285
BUSCO_Diptera_Bombyliidae_Geron_flavocciput C:2617 [S:2448, D:169], F:187, M:481, n:3285
BUSCO_Diptera_Bombyliidae_Mandella_sp C:2564 [S:2354, D:210], F:221, M:500, n:3285
BUSCO_Diptera_Bombyliidae_Meomyia_vetusta C:2768 [S:2558, D:210], F:168, M:349, n:3285
BUSCO_Diptera_Bombyliidae_Staurostichus_limbipennis C:2924 [S:2645, D:279], F:85, M:276, n:3285
BUSCO_Diptera_Bombyliidae_Thevenetimyia_longipalpus C:2891 [S:2617, D:274], F:123, M:271, n:3285
BUSCO_Diptera_Bombyliidae_Thraxan_patielus C:2811 [S:2660, D:151], F:121, M:353, n:3285
BUSCO_Diptera_Bombyliidae_Toxophora_sp C:2616 [S:2467, D:149], F:187, M:482, n:3285
BUSCO_Diptera_Bombyliidae_Villa_fuscicostata C:2734 [S:2542, D:192], F:155, M:396, n:3285
BUSCO_Diptera_Braulidae_Braula_coeca C:2185 [S:2070, D:115], F:409, M:691, n:3285
BUSCO_Diptera_Calliphoridae_Bixinia_sp C:2212 [S:2201, D:11], F:280, M:793, n:3285
BUSCO_Diptera_Calliphoridae_Calliphora_augur C:2908 [S:2898, D:10], F:104, M:273, n:3285
BUSCO_Diptera_Calliphoridae_Lucilia_cuprina C:3158 [S:3139, D:19], F:61, M:66, n:3285
BUSCO_Diptera_Calliphoridae_Stevenia_sp C:2783 [S:2756, D:27], F:126, M:376, n:3285
BUSCO_Diptera_Calliphoridae_Stomorhina_subapicalis C:2845 [S:2817, D:28], F:77, M:363, n:3285
BUSCO_Diptera_Canacidae_Dasyrhicnoessa_insularis C:2641 [S:2623, D:18], F:85, M:559, n:3285
BUSCO_Diptera_Carnidae_Carnus_hemapterus C:2439 [S:2416, D:23], F:219, M:627, n:3285
BUSCO_Diptera_Cecidomyiidae_Aphidoletes_aphidimyza C:2884 [S:2830, D:54], F:72, M:329, n:3285
BUSCO_Diptera_Cecidomyiidae_Catotricha_subobsoleta C:2891 [S:2825, D:66], F:145, M:249, n:3285
BUSCO_Diptera_Cecidomyiidae_Contarinia_nasturtii C:2837 [S:2771, D:66], F:96, M:352, n:3285
BUSCO_Diptera_Cecidomyiidae_Lestremia_cinerea C:2764 [S:876, D:1888], F:234, M:287, n:3285
BUSCO_Diptera_Cecidomyiidae_Mayetiola_destructor C:2754 [S:2715, D:39], F:189, M:342, n:3285
BUSCO_Diptera_Cecidomyiidae_Obolodiplosis_robiniae C:2898 [S:2828, D:70], F:55, M:332, n:3285
BUSCO_Diptera_Cecidomyiidae_Resseliella_maxima C:2877 [S:2785, D:92], F:72, M:336, n:3285
BUSCO_Diptera_Cecidomyiidae_Sitodiplosis_mosellana C:2861 [S:2797, D:64], F:88, M:336, n:3285
BUSCO_Diptera_Ceratopogonidae_Atrichopogon_sp C:2755 [S:2710, D:45], F:103, M:427, n:3285
BUSCO_Diptera_Ceratopogonidae_Culicoides_sonorensis C:2825 [S:2749, D:76], F:73, M:387, n:3285
BUSCO_Diptera_Ceratopogonidae_Dasyhelea_sp C:2689 [S:2627, D:62], F:73, M:523, n:3285
BUSCO_Diptera_Ceratopogonidae_Forcipomyia_taiwana C:2828 [S:2774, D:54], F:72, M:385, n:3285
BUSCO_Diptera_Chamaemyiidae_Pseudodinia_antennalis C:2081 [S:1739, D:342], F:323, M:881, n:3285
BUSCO_Diptera_Chaoboridae_Chaoborus_americanus C:2715 [S:2686, D:29], F:107, M:463, n:3285
BUSCO_Diptera_Chaoboridae_Chaoborus_flavidulus C:3023 [S:2950, D:73], F:63, M:199, n:3285
BUSCO_Diptera_Chaoboridae_Mochlonyx_cinctipes C:2577 [S:2558, D:19], F:410, M:298, n:3285
BUSCO_Diptera_Chironomidae_Cardiocladius_sp C:2884 [S:2816, D:68], F:99, M:302, n:3285
BUSCO_Diptera_Chironomidae_Chironomus_tentans C:3046 [S:2996, D:50], F:46, M:193, n:3285
BUSCO_Diptera_Chironomidae_Clunio_marinus C:3060 [S:3022, D:38], F:45, M:180, n:3285
BUSCO_Diptera_Chironomidae_Cricotopus_draysoni C:2683 [S:2531, D:152], F:161, M:441, n:3285
BUSCO_Diptera_Chironomidae_Diamesa_zernyi C:2837 [S:2668, D:169], F:128, M:320, n:3285
BUSCO_Diptera_Chironomidae_Kiefferophyes_invenustulus C:2891 [S:1843, D:1048], F:124, M:270, n:3285
BUSCO_Diptera_Chironomidae_Paraheptagyia_tonnoiri C:2893 [S:2832, D:61], F:94, M:298, n:3285
BUSCO_Diptera_Chironomidae_Paraphaenocladius_impensus C:2789 [S:2719, D:70], F:133, M:363, n:3285
BUSCO_Diptera_Chironomidae_Parochlus_steinenii C:2194 [S:2179, D:15], F:343, M:748, n:3285
BUSCO_Diptera_Chironomidae_Pentaneurella_katterjokki C:2905 [S:2213, D:692], F:96, M:284, n:3285
BUSCO_Diptera_Chironomidae_Podonomus_sp C:2992 [S:2869, D:123], F:68, M:225, n:3285
BUSCO_Diptera_Chironomidae_Polypedilum_pembai C:3019 [S:2932, D:87], F:75, M:191, n:3285
BUSCO_Diptera_Chironomidae_Polypedilum_vanderplanki C:3065 [S:3006, D:59], F:62, M:158, n:3285
BUSCO_Diptera_Chironomidae_Procladius_villosimanus C:2774 [S:2431, D:343], F:103, M:408, n:3285
BUSCO_Diptera_Chironomidae_Propsilocerus_akamusi C:3053 [S:3003, D:50], F:48, M:184, n:3285
BUSCO_Diptera_Chironomidae_Smittia_aterrima C:3058 [S:3013, D:45], F:46, M:181, n:3285
BUSCO_Diptera_Chironomidae_Telmatogeton_pectinata C:2602 [S:2530, D:72], F:153, M:530, n:3285
BUSCO_Diptera_Chironomidae_Telmatopelopia_nemorum C:2665 [S:2596, D:69], F:128, M:492, n:3285
BUSCO_Diptera_Chloropidae_Chlorops_oryzae C:2914 [S:2798, D:116], F:117, M:254, n:3285
BUSCO_Diptera_Chloropidae_Lipara_lucens C:2743 [S:2703, D:40], F:120, M:422, n:3285
BUSCO_Diptera_Chyromyidae_Gymnochiromyia_sp C:2001 [S:1980, D:21], F:294, M:990, n:3285
BUSCO_Diptera_Clusiidae_Clusia_lateralis C:2034 [S:2003, D:31], F:336, M:915, n:3285
BUSCO_Diptera_Coelopidae_Coelopa_frigida C:2859 [S:2846, D:13], F:77, M:349, n:3285
BUSCO_Diptera_Conopidae_Myopa_sp C:2594 [S:2224, D:370], F:153, M:538, n:3285

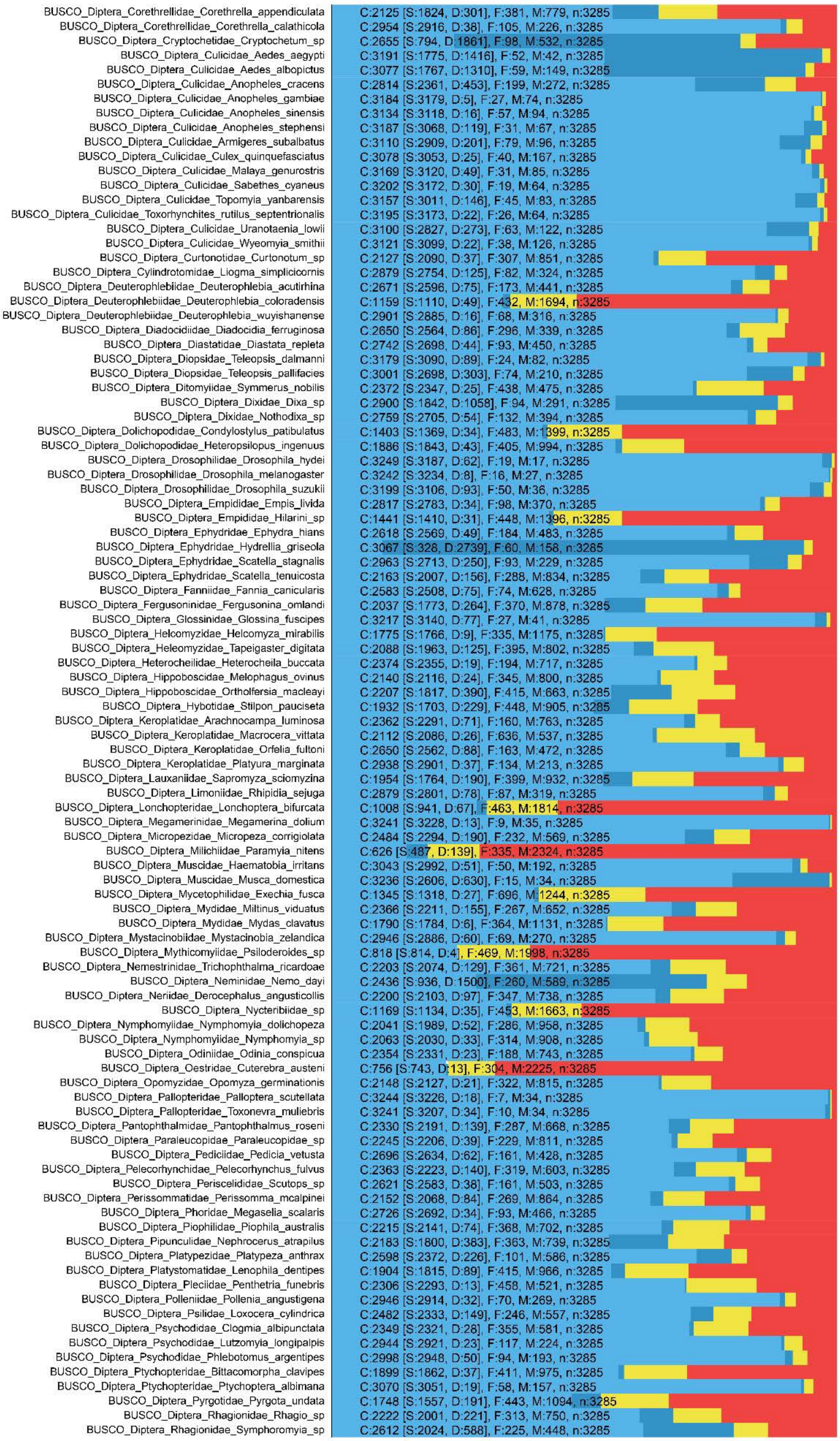
BUSCO_Diptera_Corethrellidae_Corethrella_appendiculata C:2125 [S:1824, D:301], F:381, M:779, n:3285
BUSCO_Diptera_Corethrellidae_Corethrella_calathicola C:2954 [S:2916, D:38], F:105, M:226, n:3285
BUSCO_Diptera_Cryptochetidae_Cryptochetum_sp C:2655 [S:794, D:1861], F:98, M:532, n:3285
BUSCO_Diptera_Culicidae_Aedes_aegypti C:3191 [S:1775, D:1416], F:52, M:42, n:3285
BUSCO_Diptera_Culicidae_Aedes_albopictus C:3077 [S:1767, D:1310], F:59, M:149, n:3285
BUSCO_Diptera_Culicidae_Anopheles_cracens C:2814 [S:2361, D:453], F:199, M:272, n:3285
BUSCO_Diptera_Culicidae_Anopheles_gambiae C:3184 [S:3179, D:5], F:27, M:74, n:3285
BUSCO_Diptera_Culicidae_Anopheles_sinensis C:3134 [S:3118, D:16], F:57, M:94, n:3285
BUSCO_Diptera_Culicidae_Anopheles_stephensi C:3187 [S:3068, D:119], F:31, M:67, n:3285
BUSCO_Diptera_Culicidae_Armigeres_subalbatus C:3110 [S:2909, D:201], F:79, M:96, n:3285
BUSCO_Diptera_Culicidae_Culex_quinquefasciatus C:3078 [S:3053, D:25], F:40, M:167, n:3285
BUSCO_Diptera_Culicidae_Malaya_genurostris C:3169 [S:3120, D:49], F:31, M:85, n:3285
BUSCO_Diptera_Culicidae_Sabethes_cyaneus C:3202 [S:3172, D:30], F:19, M:64, n:3285
BUSCO_Diptera_Culicidae_Topomyia_yanbarensis C:3157 [S:3011, D:146], F:45, M:83, n:3285
BUSCO_Diptera_Culicidae_Toxorhynchites_rutilus_septentrionalis C:3195 [S:3173, D:22], F:26, M:64, n:3285
BUSCO_Diptera_Culicidae_Uranotaenia_lowii C:3100 [S:2827, D:273], F:63, M:122, n:3285
BUSCO_Diptera_Culicidae_Wyeomyia_smithii C:3121 [S:3099, D:22], F:38, M:126, n:3285
BUSCO_Diptera_Curtonotidae_Curtonotum_sp C:2127 [S:2090, D:37], F:307, M:851, n:3285
BUSCO_Diptera_Cylindrotomidae_Liogma_simplicicornis C:2879 [S:2754, D:125], F:82, M:324, n:3285
BUSCO_Diptera_Deuterophlebiidae_Deuterophlebia_acutirhina C:2671 [S:2596, D:75], F:173, M:441, n:3285
BUSCO_Diptera_Deuterophlebiidae_Deuterophlebia_coloradensis C:1159 [S:1110, D:49], F:432, M:1694, n:3285
BUSCO_Diptera_Deuterophlebiidae_Deuterophlebia_wuyishanense C:2901 [S:2885, D:16], F:68, M:316, n:3285
BUSCO_Diptera_Diadocidiidae_Diadocidia_ferruginosa C:2650 [S:2564, D:86], F:296, M:339, n:3285
BUSCO_Diptera_Diastatidae_Diastata_repleta C:2742 [S:2698, D:44], F:93, M:450, n:3285
BUSCO_Diptera_Diopsidae_Teleopsis_dalmanni C:3179 [S:3090, D:89], F:24, M:82, n:3285
BUSCO_Diptera_Diopsidae_Teleopsis_pallifacies C:3001 [S:2698, D:303], F:74, M:210, n:3285
BUSCO_Diptera_Ditomyiidae_Symmerus_nobilis C:2372 [S:2347, D:25], F:438, M:475, n:3285
BUSCO_Diptera_Dixidae_Dixa_sp C:2900 [S:1842, D:1058], F:94, M:291, n:3285
BUSCO_Diptera_Dixidae_Nothodixa_sp C:2759 [S:2705, D:54], F:132, M:394, n:3285
BUSCO_Diptera_Dolichopodidae_Condylostylus_patibulatus C:1403 [S:1369, D:34], F:483, M:1399, n:3285
BUSCO_Diptera_Dolichopodidae_Heteropsilopus_ingenuus C:1886 [S:1843, D:43], F:405, M:994, n:3285
BUSCO_Diptera_Drosophilidae_Drosophila_hydei C:3249 [S:3187, D:62], F:19, M:17, n:3285
BUSCO_Diptera_Drosophilidae_Drosophila_melanogaster C:3242 [S:3234, D:8], F:16, M:27, n:3285
BUSCO_Diptera_Drosophilidae_Drosophila_suzukii C:3199 [S:3106, D:93], F:50, M:36, n:3285
BUSCO_Diptera_Empididae_Empis_livida C:2817 [S:2783, D:34], F:98, M:370, n:3285
BUSCO_Diptera_Empididae_Hilarini_sp C:1441 [S:1410, D:31], F:448, M:1396, n:3285
BUSCO_Diptera_Ephydridae_Ephydra_hians C:2618 [S:2569, D:49], F:184, M:483, n:3285
BUSCO_Diptera_Ephydridae_Hydrellia_griseola C:3067 [S:328, D:2739], F:60, M:158, n:3285
BUSCO_Diptera_Ephydridae_Scatella_stagnalis C:2963 [S:2713, D:250], F:93, M:229, n:3285
BUSCO_Diptera_Ephydridae_Scatella_tenuicosta C:2163 [S:2007, D:156], F:288, M:834, n:3285
BUSCO_Diptera_Fanniidae_Fannia_canicularis C:2583 [S:2508, D:75], F:74, M:628, n:3285
BUSCO_Diptera_Fergusoninidae_Fergusonina_omlandi C:2037 [S:1773, D:264], F:370, M:878, n:3285
BUSCO_Diptera_Glossinidae_Glossina_fuscipes C:3217 [S:3140, D:77], F:27, M:41, n:3285
BUSCO_Diptera_Helcomyzidae_Helcomyza_mirabilis C:1775 [S:1766, D:9], F:335, M:1175, n:3285
BUSCO_Diptera_Heleomyzidae_Tapeigaster_digitata C:2088 [S:1963, D:125], F:395, M:802, n:3285
BUSCO_Diptera_Heterocheilidae_Heterocheila_buccata C:2374 [S:2355, D:19], F:194, M:717, n:3285
BUSCO_Diptera_Hippoboscidae_Melophagus_ovinus C:2140 [S:2116, D:24], F:345, M:800, n:3285
BUSCO_Diptera_Hippoboscidae_Ortholfersia_macleayi C:2207 [S:1817, D:390], F:415, M:663, n:3285
BUSCO_Diptera_Hybotidae_Stilpon_pauciseta C:1932 [S:1703, D:229], F:448, M:905, n:3285
BUSCO_Diptera_Keroplatidae_Arachnocampa_luminosa C:2362 [S:2291, D:71], F:160, M:763, n:3285
BUSCO_Diptera_Keroplatidae_Macrocera_vittata C:2112 [S:2086, D:26], F:636, M:537, n:3285
BUSCO_Diptera_Keroplatidae_Orfelia_fultoni C:2650 [S:2562, D:88], F:163, M:472, n:3285
BUSCO_Diptera_Keroplatidae_Platyura_marginata C:2938 [S:2901, D:37], F:134, M:213, n:3285
BUSCO_Diptera_Lauxaniidae_Sapromyza_sciomyzina C:1954 [S:1764, D:190], F:399, M:932, n:3285
BUSCO_Diptera_Limoniidae_Rhipidia_sejuga C:2879 [S:2801, D:78], F:87, M:319, n:3285
BUSCO_Diptera_Lonchopteridae_Lonchoptera_bifurcata C:1008 [S:941, D:67], F:463, M:1814, n:3285
BUSCO_Diptera_Megamerinidae_Megamerina_dolium C:3241 [S:3228, D:13], F:9, M:35, n:3285
BUSCO_Diptera_Micropezidae_Micropeza_corrigiolata C:2484 [S:2294, D:190], F:232, M:569, n:3285
BUSCO_Diptera_Milichiidae_Paramyia_nitens C:626 [S:487, D:139], F:335, M:2324, n:3285
BUSCO_Diptera_Muscidae_Haematobia_irritans C:3043 [S:2992, D:51], F:50, M:192, n:3285
BUSCO_Diptera_Muscidae_Musca_domestica C:3236 [S:2606, D:630], F:15, M:34, n:3285
BUSCO_Diptera_Mycetophilidae_Exechia_fusca C:1345 [S:1318, D:27], F:696, M:1244, n:3285
BUSCO_Diptera_Mydidae_Miltinus_viduatus C:2366 [S:2211, D:155], F:267, M:652, n:3285
BUSCO_Diptera_Mydidae_Mydas_clavatus C:1790 [S:1784, D:6], F:364, M:1131, n:3285
BUSCO_Diptera_Mystacinobiidae_Mystacinobia_zelandica C:2946 [S:2886, D:60], F:69, M:270, n:3285
BUSCO_Diptera_Mythicomyiidae_Psiloderoides_sp C:818 [S:814, D:4], F:469, M:1998, n:3285
BUSCO_Diptera_Nemestrinidae_Trichophthalma_ricardoae C:2203 [S:2074, D:129], F:361, M:721, n:3285
BUSCO_Diptera_Neminidae_Nemo_dayi C:2436 [S:936, D:1500], F:260, M:589, n:3285
BUSCO_Diptera_Neriidae_Derocephalus_angusticollis C:2200 [S:2103, D:97], F:347, M:738, n:3285
BUSCO_Diptera_Nycteribiidae_sp C:1169 [S:1134, D:35], F:453, M:1663, n:3285
BUSCO_Diptera_Nymphomyiidae_Nymphomyia_dolichopeza C:2041 [S:1989, D:52], F:286, M:958, n:3285
BUSCO_Diptera_Nymphomyiidae_Nymphomyia_sp C:2063 [S:2030, D:33], F:314, M:908, n:3285
BUSCO_Diptera_Odiniidae_Odinia_conspicua C:2354 [S:2331, D:23], F:188, M:743, n:3285
BUSCO_Diptera_Oestridae_Cuterebra_austeni C:756 [S:743, D:13], F:304, M:2225, n:3285
BUSCO_Diptera_Opomyzidae_Opomyza_germinationis C:2148 [S:2127, D:21], F:322, M:815, n:3285
BUSCO_Diptera_Pallopteridae_Palloptera_scutellata C:3244 [S:3226, D:18], F:7, M:34, n:3285
BUSCO_Diptera_Pallopteridae_Toxonevra_muliebris C:3241 [S:3207, D:34], F:10, M:34, n:3285
BUSCO_Diptera_Pantophthalmidae_Pantophthalmus_roseni C:2330 [S:2191, D:139], F:287, M:668, n:3285
BUSCO_Diptera_Paraleucopidae_Paraleucopidae_sp C:2245 [S:2206, D:39], F:229, M:811, n:3285
BUSCO_Diptera_Pediciidae_Pedicia_vetusta C:2696 [S:2634, D:62], F:161, M:428, n:3285
BUSCO_Diptera_Pelecorhynchidae_Pelecorhynchus_fulvus C:2363 [S:2223, D:140], F:319, M:603, n:3285
BUSCO_Diptera_Periscelididae_Scutops_sp C:2621 [S:2583, D:38], F:161, M:503, n:3285
BUSCO_Diptera_Perissommatidae_Perissomma_mcalpinei C:2152 [S:2068, D:84], F:269, M:864, n:3285
BUSCO_Diptera_Phoridae_Megaselia_scalaris C:2726 [S:2692, D:34], F:93, M:466, n:3285
BUSCO_Diptera_Piophilidae_Piophila_australis C:2215 [S:2141, D:74], F:368, M:702, n:3285
BUSCO_Diptera_Pipunculidae_Nephrocerus_atrapilus C:2183 [S:1800, D:383], F:363, M:739, n:3285
BUSCO_Diptera_Platypezidae_Platypeza_anthrax C:2598 [S:2372, D:226], F:101, M:586, n:3285
BUSCO_Diptera_Platystomatidae_Lenophila_dentipes C:1904 [S:1815, D:89], F:415, M:966, n:3285
BUSCO_Diptera_Pleciidae_Penthetria_funebris C:2306 [S:2293, D:13], F:458, M:521, n:3285
BUSCO_Diptera_Polleniidae_Pollenia_angustigena C:2946 [S:2914, D:32], F:70, M:269, n:3285
BUSCO_Diptera_Psilidae_Loxocera_cylindrica C:2482 [S:2333, D:149], F:246, M:557, n:3285
BUSCO_Diptera_Psychodidae_Clogmia_albipunctata C:2349 [S:2321, D:28], F:355, M:581, n:3285
BUSCO_Diptera_Psychodidae_Lutzomyia_longipalpis C:2944 [S:2921, D:23], F:117, M:224, n:3285
BUSCO_Diptera_Psychodidae_Phlebotomus_argentipes C:2998 [S:2948, D:50], F:94, M:193, n:3285
BUSCO_Diptera_Ptychopteridae_Bittacomorpha_clavipes C:1899 [S:1862, D:37], F:411, M:975, n:3285
BUSCO_Diptera_Ptychopteridae_Ptychoptera_albimana C:3070 [S:3051, D:19], F:58, M:157, n:3285
BUSCO_Diptera_Pyrgotidae_Pyrgota_undata C:1748 [S:1557, D:191], F:443, M:1094, n:3285
BUSCO_Diptera_Rhagionidae_Rhagio_sp C:2222 [S:2001, D:221], F:313, M:750, n:3285
BUSCO_Diptera_Rhagionidae_Symphoromyia_sp C:2612 [S:2024, D:588], F:225, M:448, n:3285

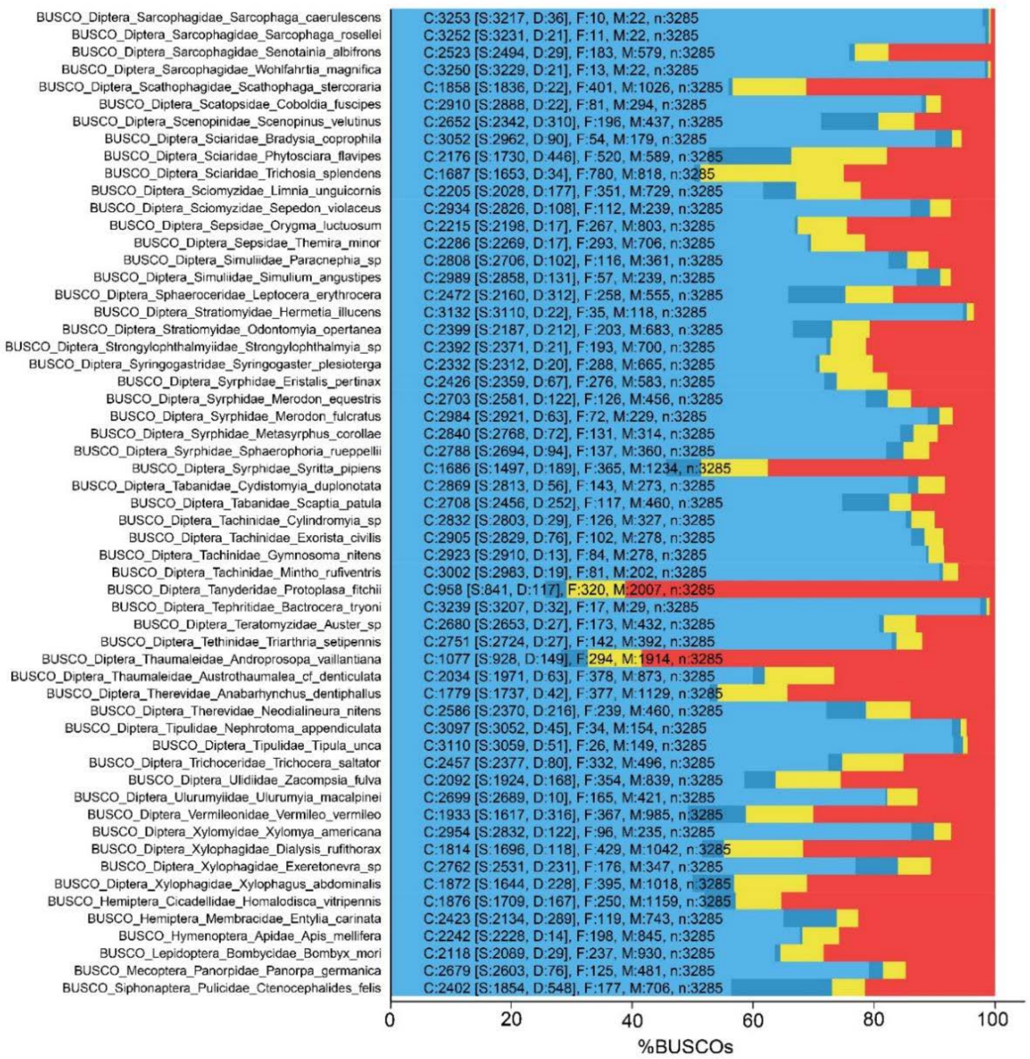


**Figure S2. BUSCO assessments of species genomic data used for reconstruction of Diptera phylogenetic trees.**

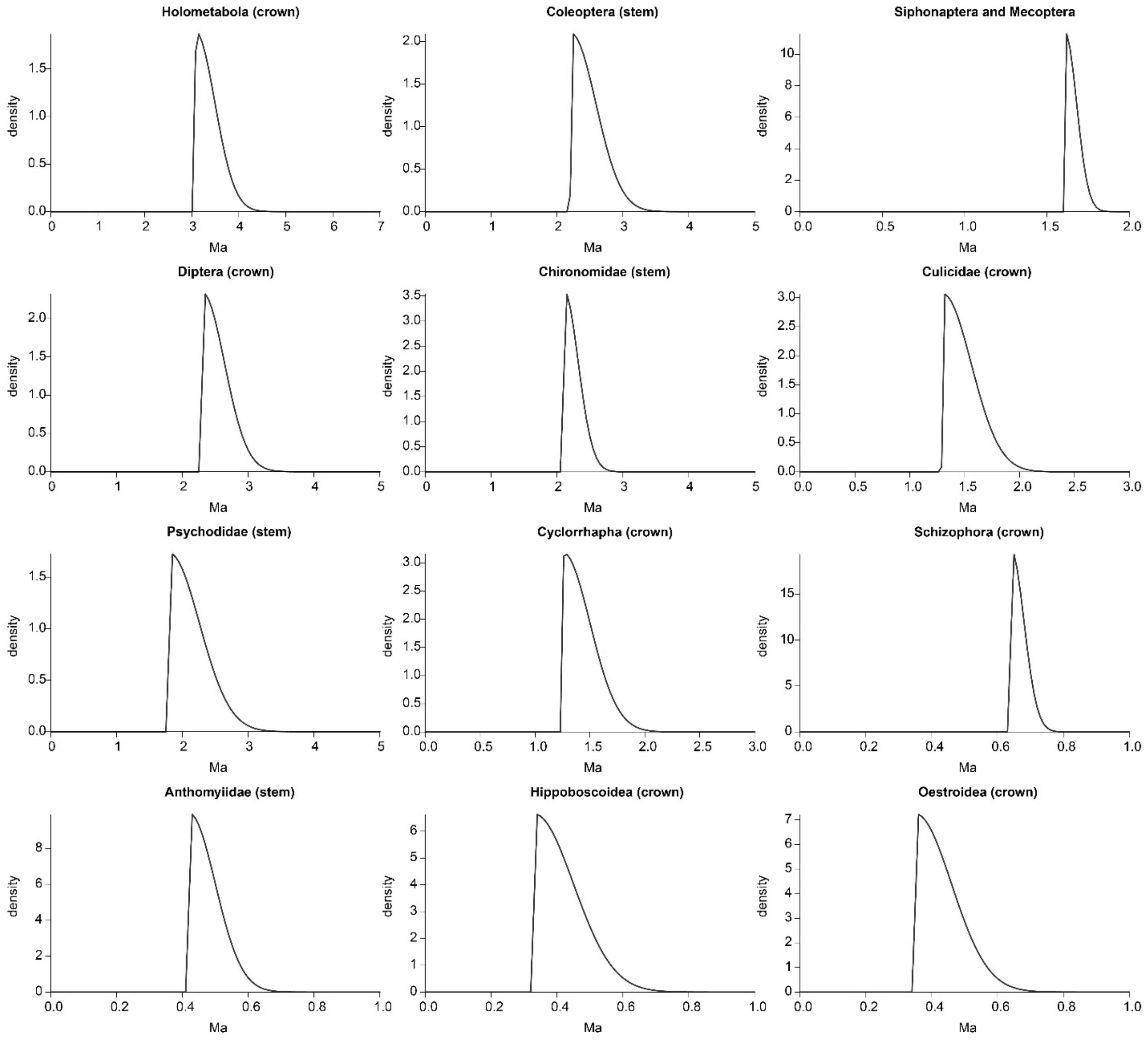


**Figure S3. Fossil Calibrations using skew-Normal distribution model.**

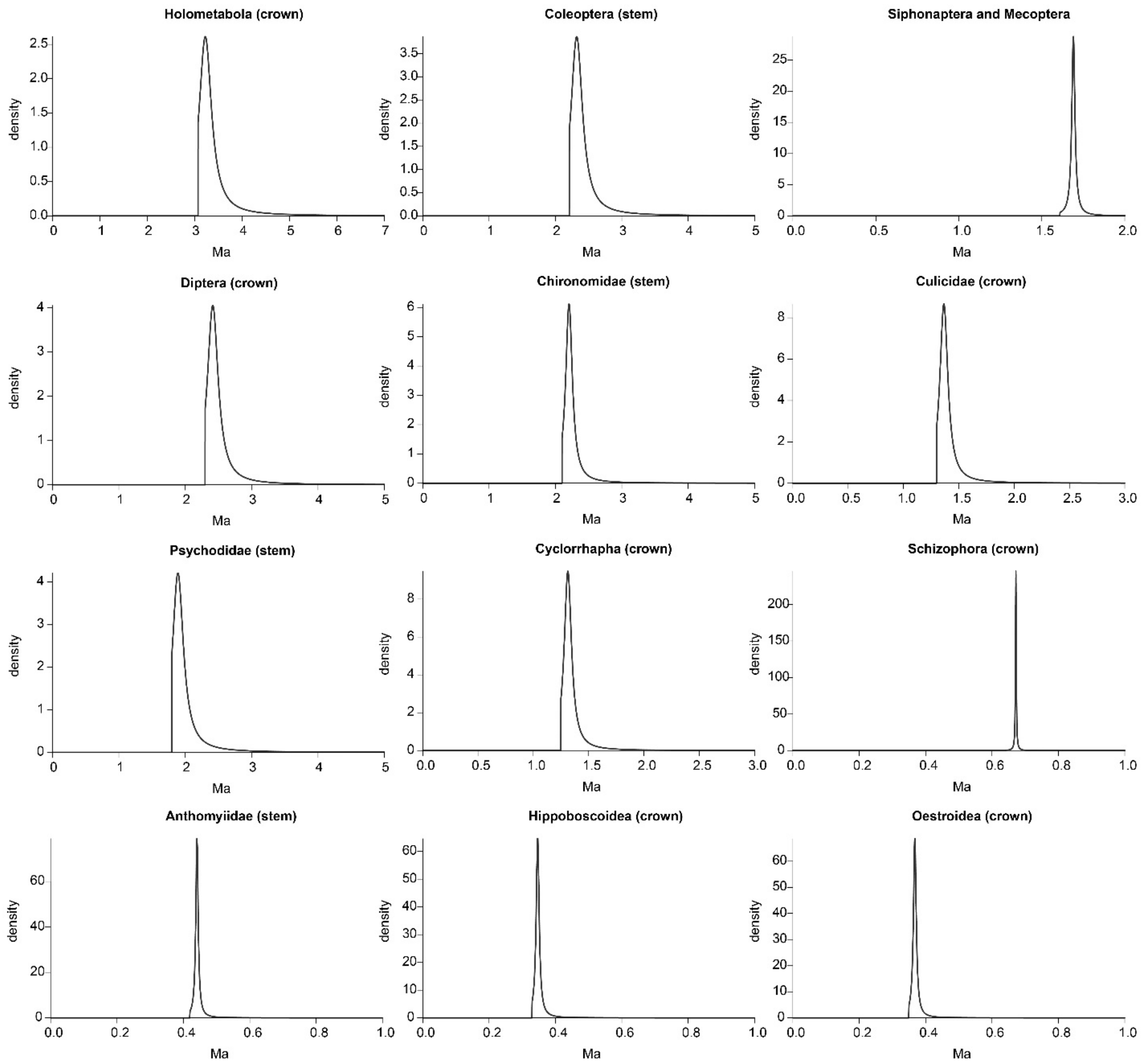


**Figure S4. Fossil Calibrations using cauchy distribution model.**

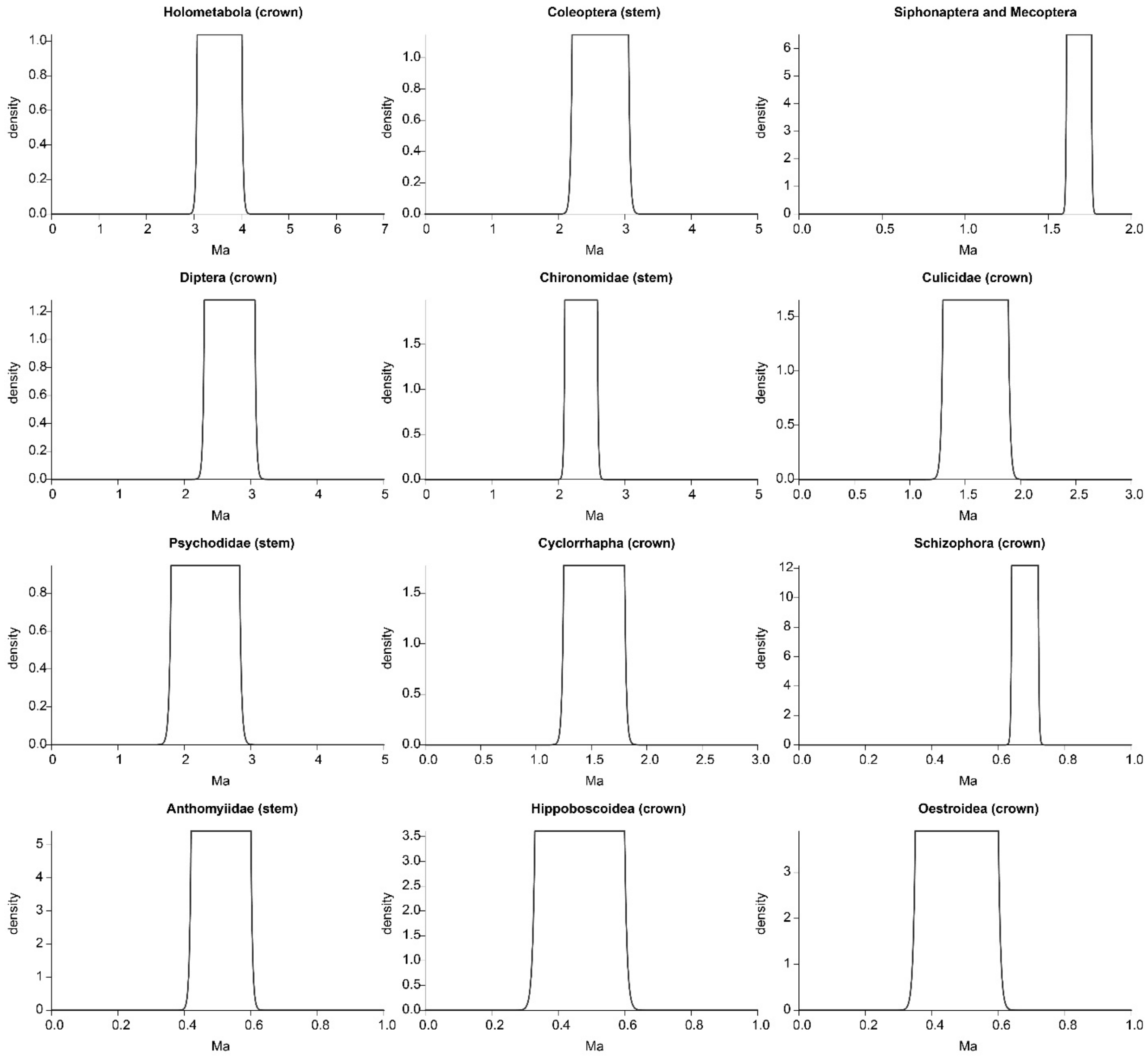


**Figure S5. Fossil Calibrations using uniform distribution model.**

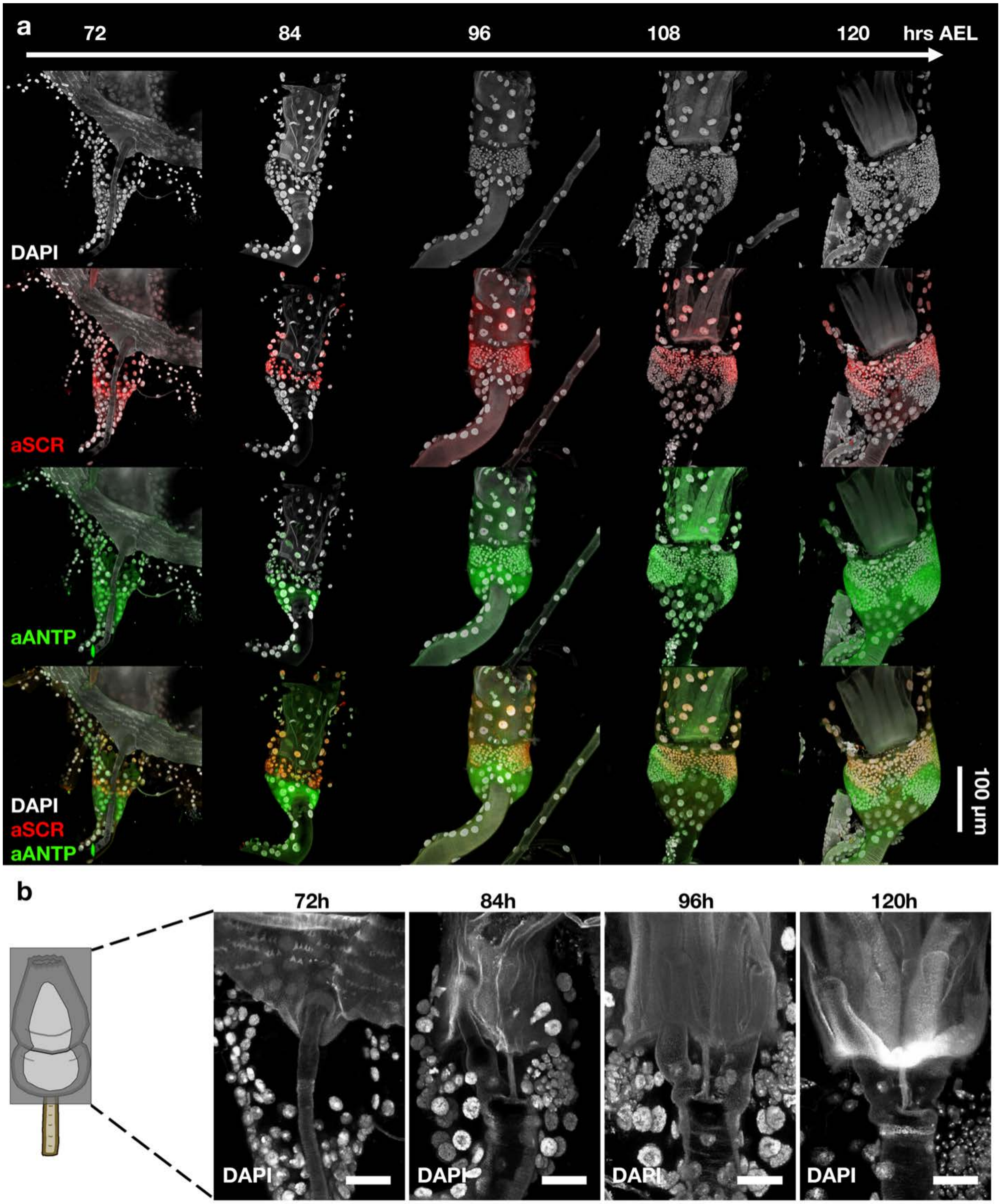


**Figure S6. The morphogenesis of the *D. melanogaster* PROD disc. (a)** The anti-SCR and anti-ANTP marking the developing *D. melanogaster* PROD disc at 72, 84, 96, 108, and 120 hrs after egg laying (AEL). **(b)** The developmental changes of the larval spiracle gradually replaced by the branched pupal spiracles. Scalebar=100 μm.

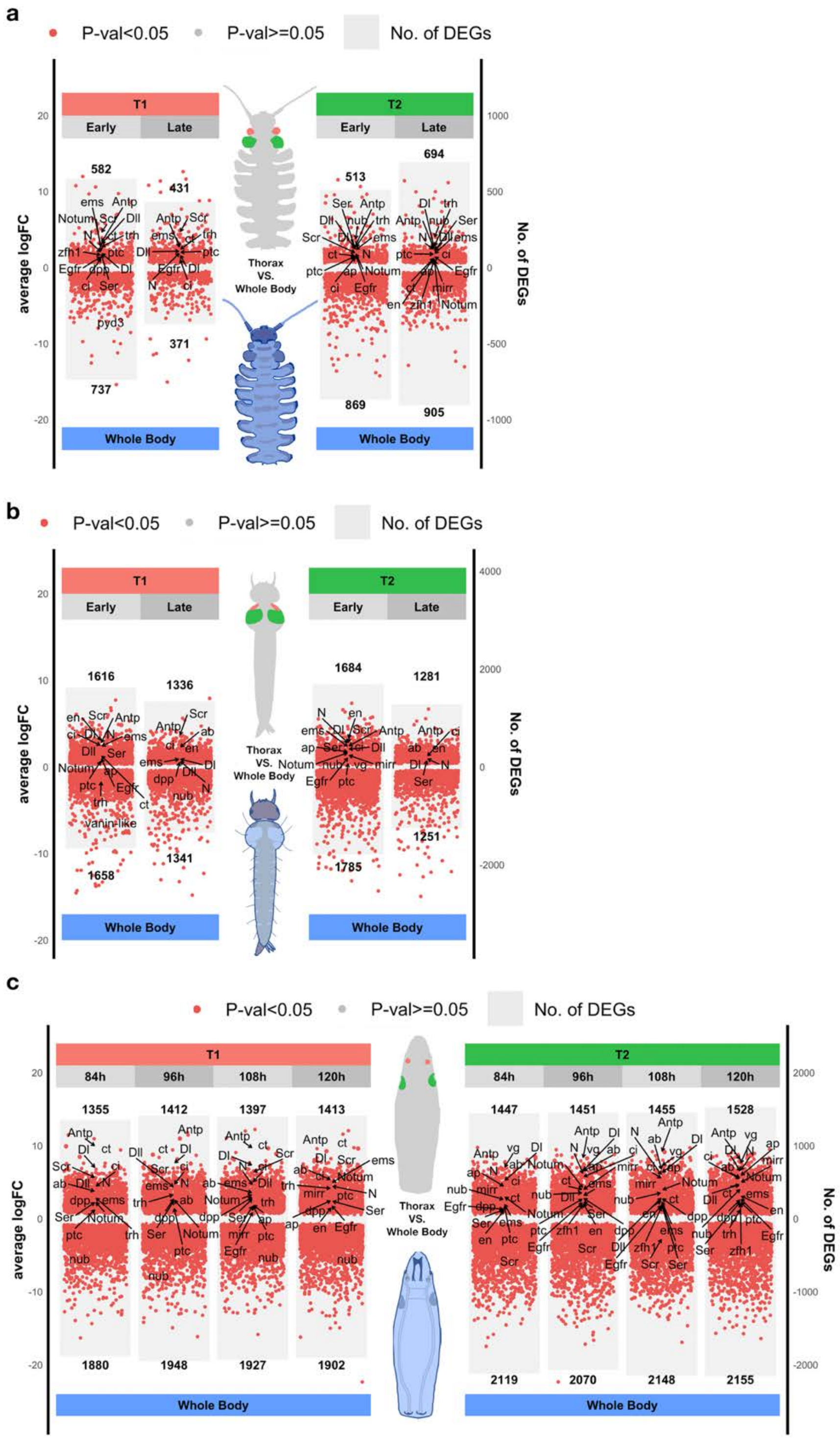


**Figure S7. Differential gene expression analysis of thoracic disc (T1, T2) vs. whole body in three dipteran species.** Two or four developmental time points were examined for *De. wuyiensis* (**A**), *A. albopictus* (**B**), *D. melanogaster* (**C**). DEGs with *p*-value < 0.05 are highlighted in red. The light gray histogram and the right y-axis indicates the number of DEGs within each comparison.

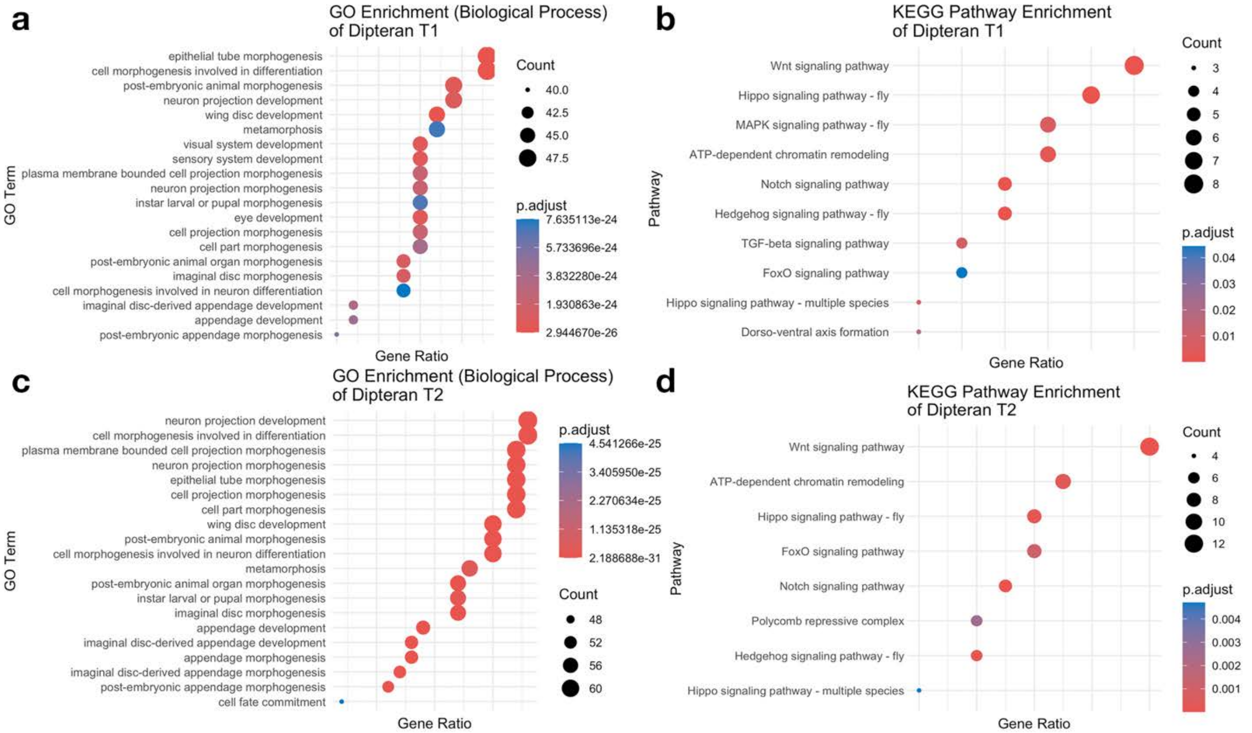


**Figure S8. PROD&Wing-shared DEGs across *De. wuyiensis*, *A. albopictus*, and *D. melanogaster.*** **(a, c)** Gene Ontology (GO) and Kyoto Encyclopedia of Genes and Genomes (KEGG) pathway enrichment analyses of 181 DEGs up-regulated in PROD vs. Body. **(b, d)** GO and KEGG pathway enrichment analyses of 270 DEGs up-regulated in wing vs. Body DEGs. GO and KEGG analyses reveal conserved genetic programs and pathways involved in thoracic development and function.

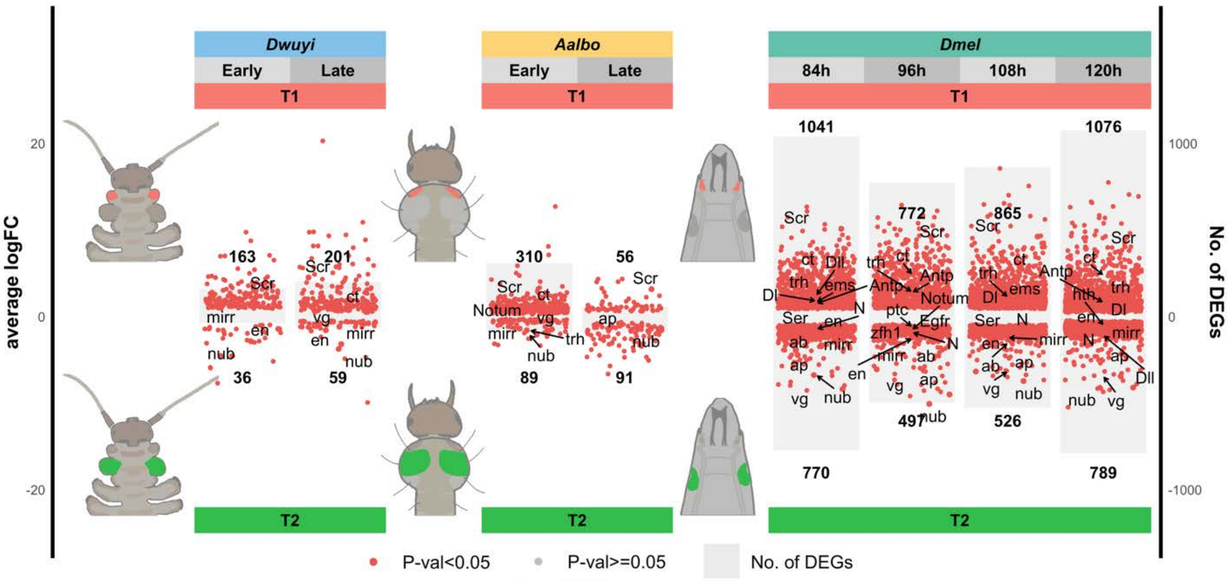


**Figure S9. Differential gene expression between T1 and T2 across dipteran species.** *De. wuyiensis*, *A. albopictus*, and *D. melanogaster.* The gray box histogram and the right y-axis denotes the number of DEGs (p<0.05).

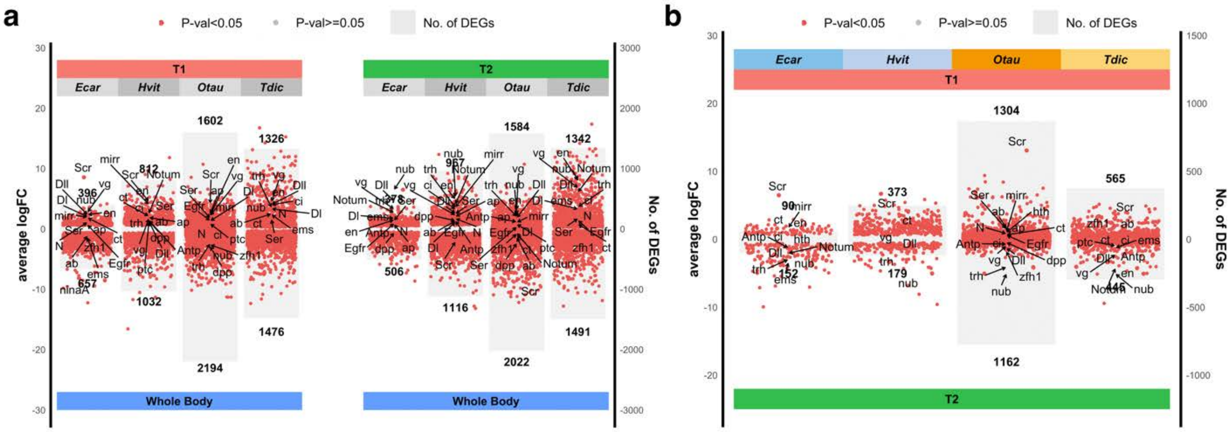


**Figure S10. Differential gene expression in outgroup species.** (**a**) DEGs between T1 or T2 versus whole-body (or control tissue fat & gut available in *T. dichotomus*) for *E. carinata* (*Ecar*), *H. vitripennis* (*Hvit*), *T. dichotomus* (*Tdic*), and *O. taurus* (*Otau*). (**b**) DEGs between T1 versus T2 for the four outgroup species. The gray box histogram on the right y-axis denotes the number of DEGs (p<0.05).

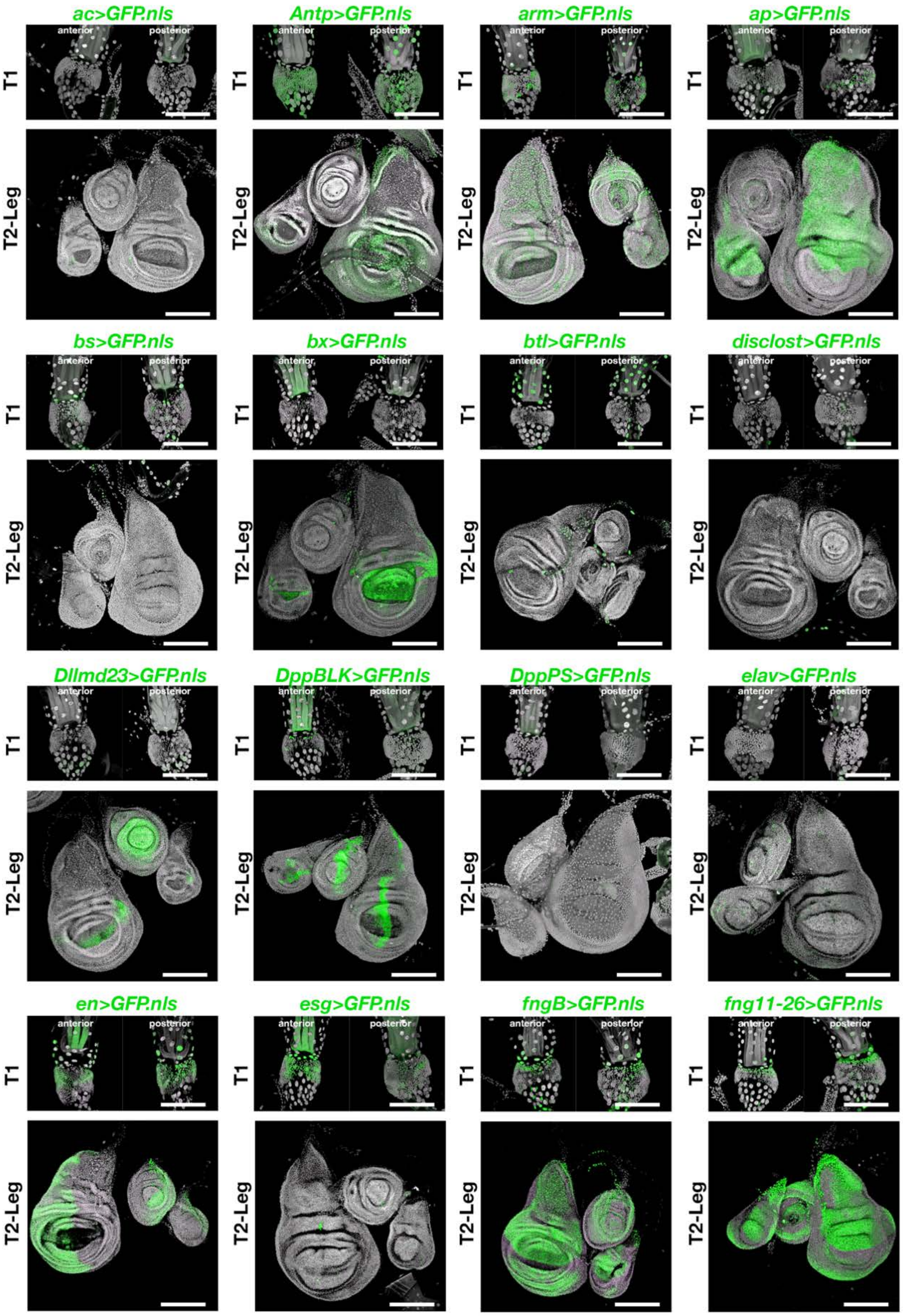


Figure S11 continued to next page

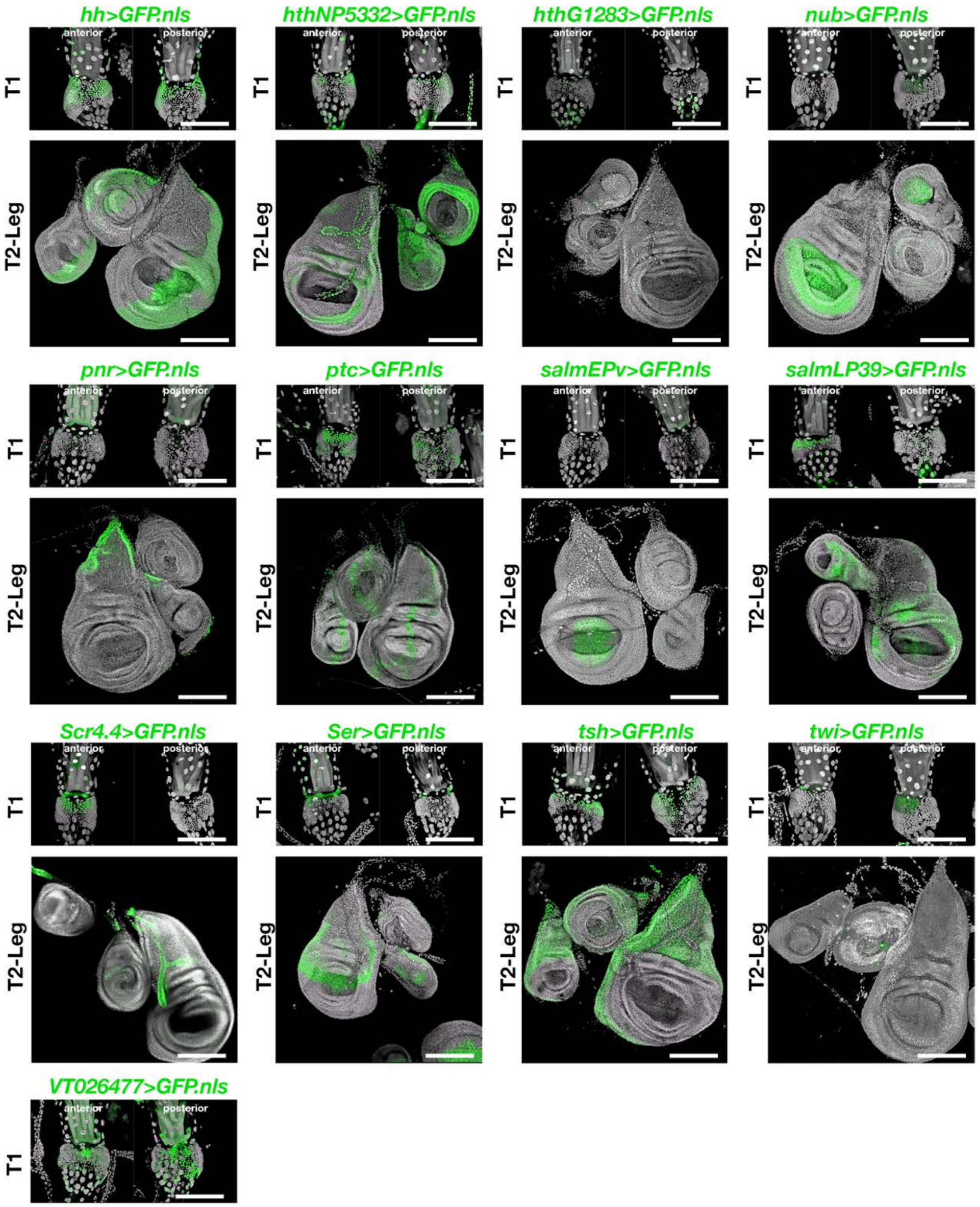


**Figure S11. GAL4 reporter assays showing *in vivo* expression patterns of genes associated with wing development in the *D. melanogaster* PROD disc, wing disc, haltere disc, and leg discs.** This is also part of screening for *GAL4* drivers (Table S5) to use for gene manipulation in PROD disc. The *VT026477-GAL4*, which drive gene expression specifically in the prothoracic dorsal discs, and *en-GAL4*, which drive gene expression serially in the PROD disc, wing disc, and haltere disc, are used in the RNAi screening. Scalebar=100 μm.

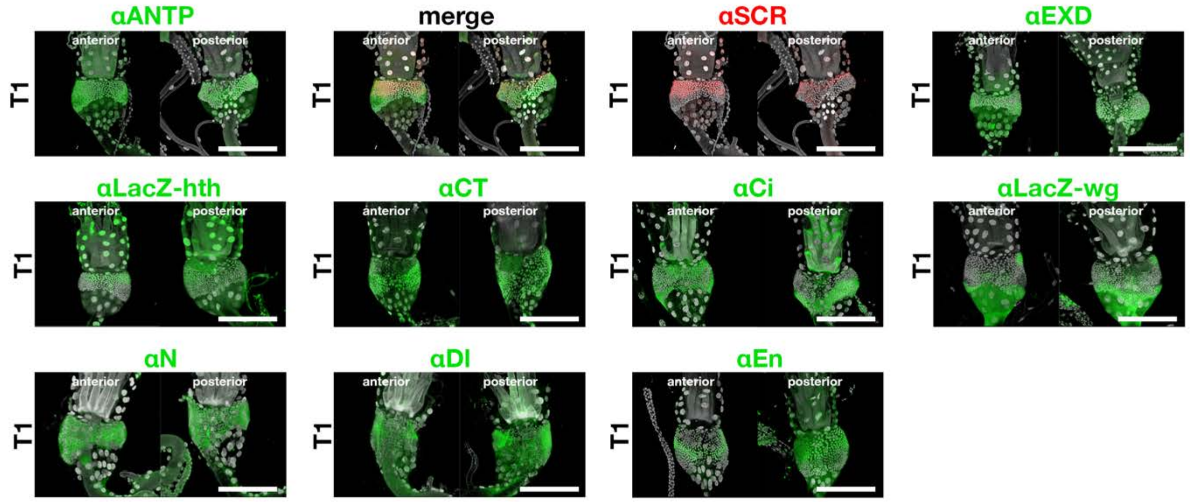


**Figure S12. The *in vivo* expression patterns of the genes associated with wing development, marked by appropriate antibodies in the *D. melanogaster* PROD disc.** *lacZ* reporter lines and antibodies detailed in Table S7 and S8. Scalebar=100 µm.

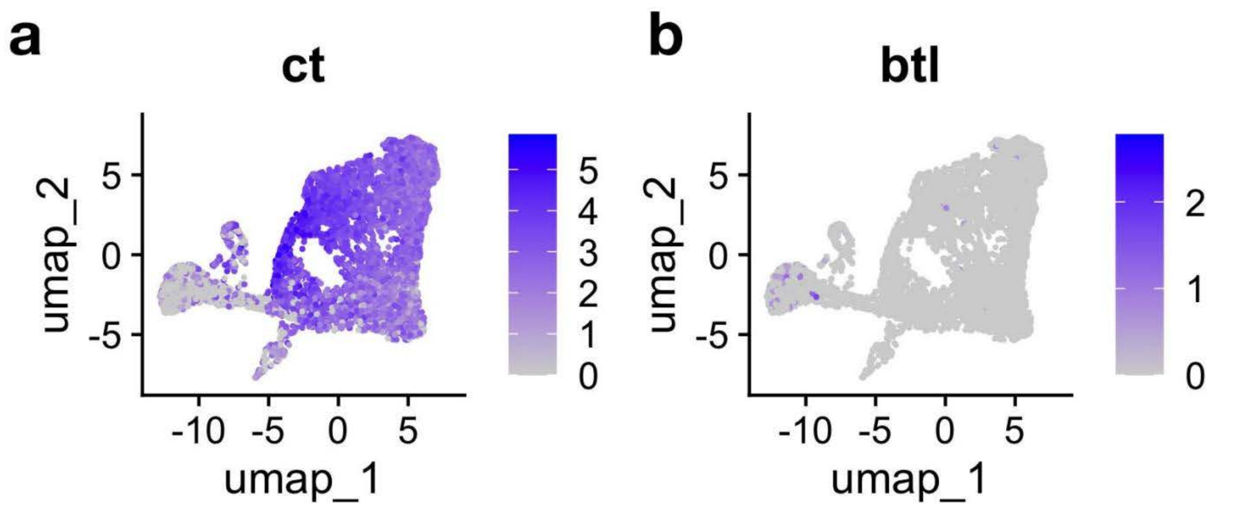


**Figure S13. Expression patterns of *btl* and *ct* genes visualized by UMAP. (a)** UMAP projection showing the expression of the homeobox gene ct across single-cell transcriptomes of the PROD disc. **(b)** UMAP projection showing the expression of the tracheal gene btl across single-cell transcriptomes of the PROD disc.

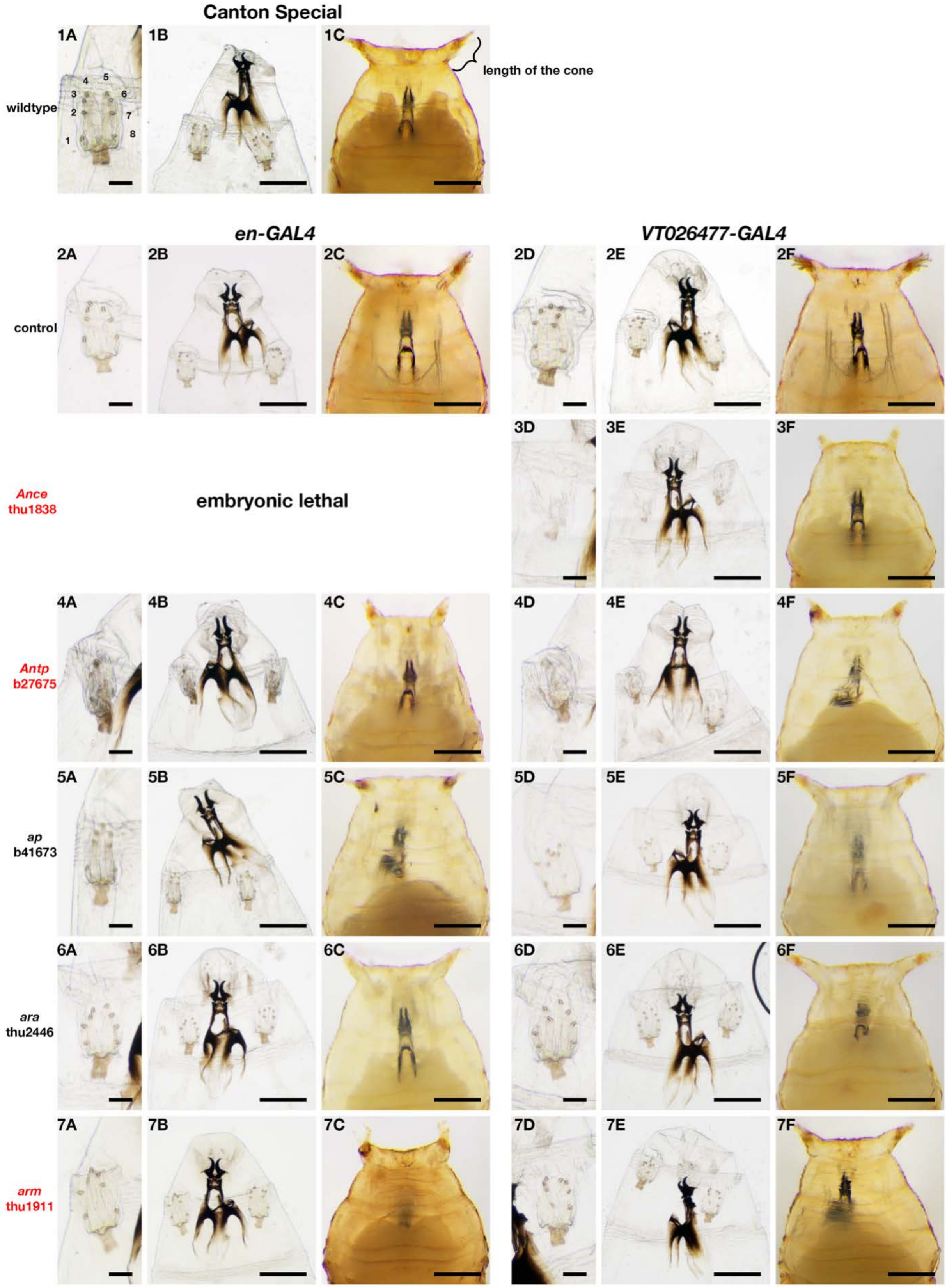


Figure S14 continued to next page

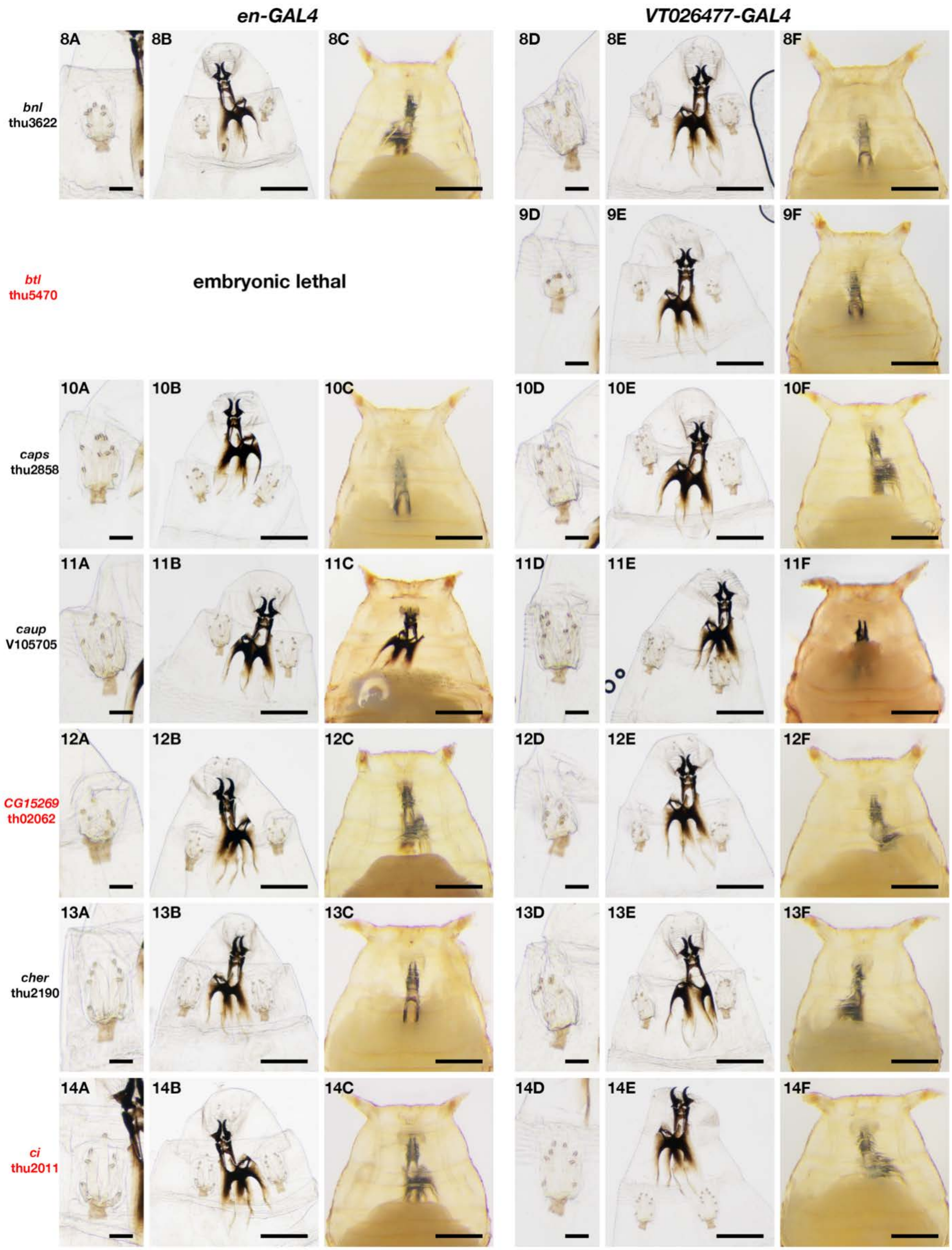


Figure S14 continued to next page

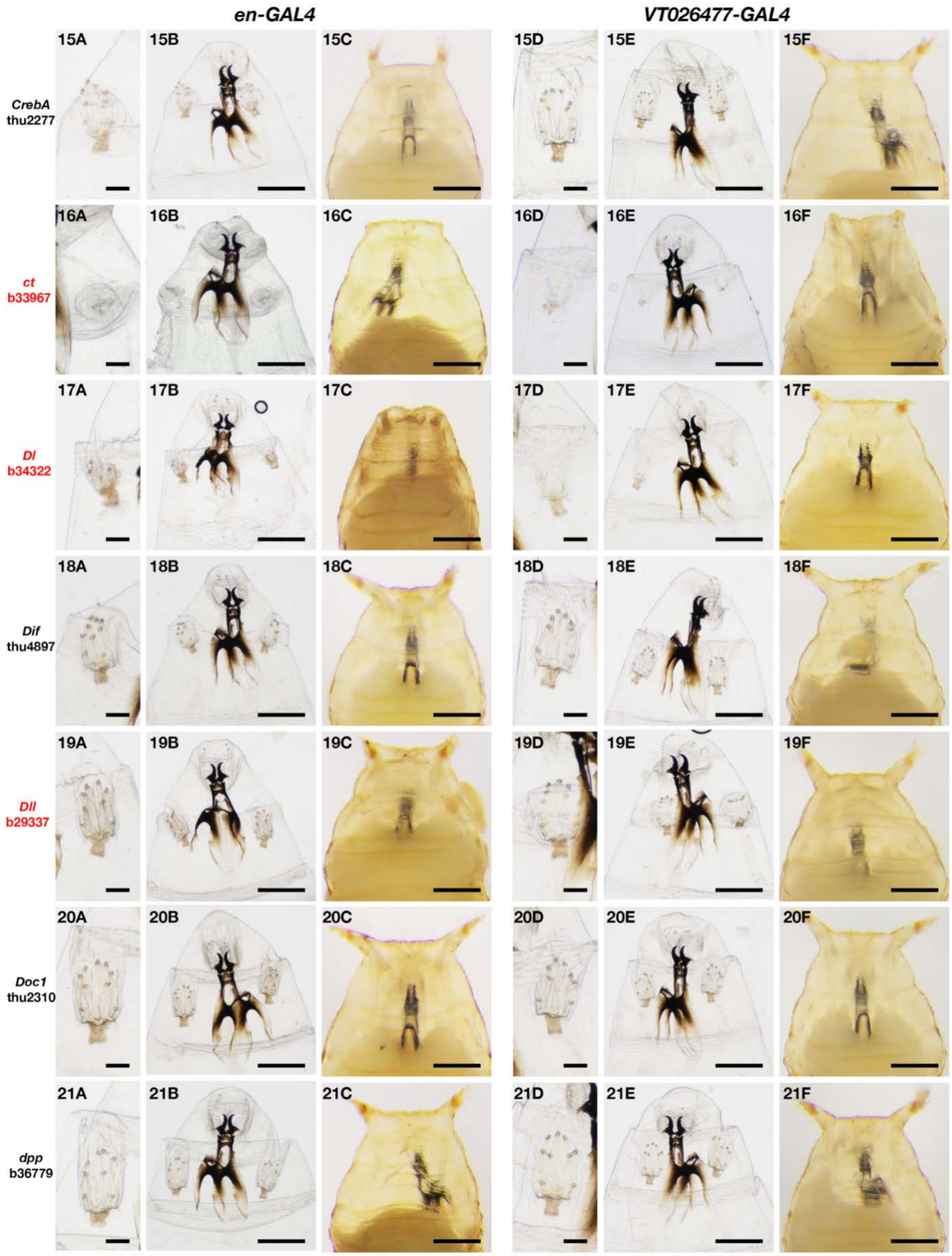


112
113 Figure S14 continued to next page

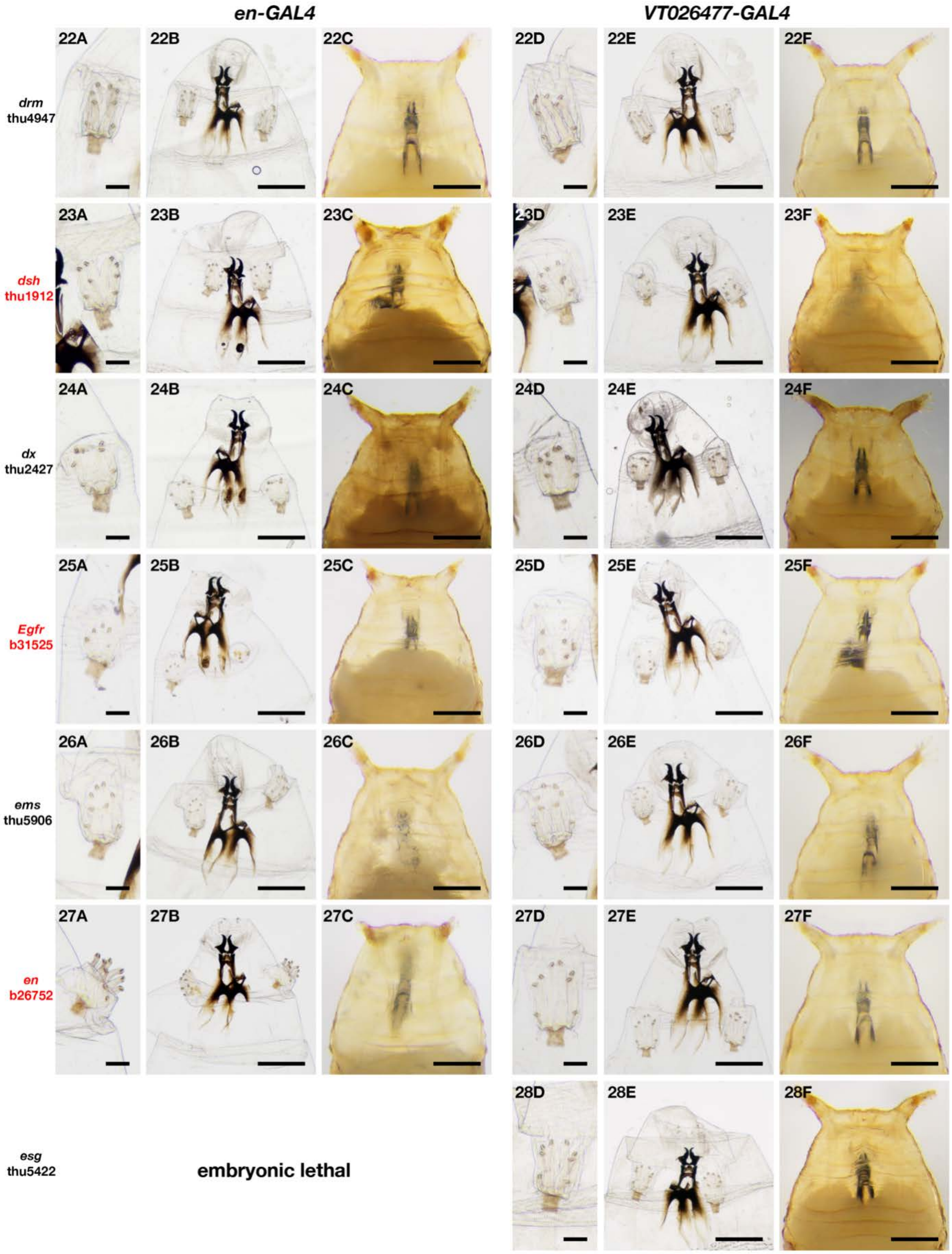


Figure S14 continued to next page

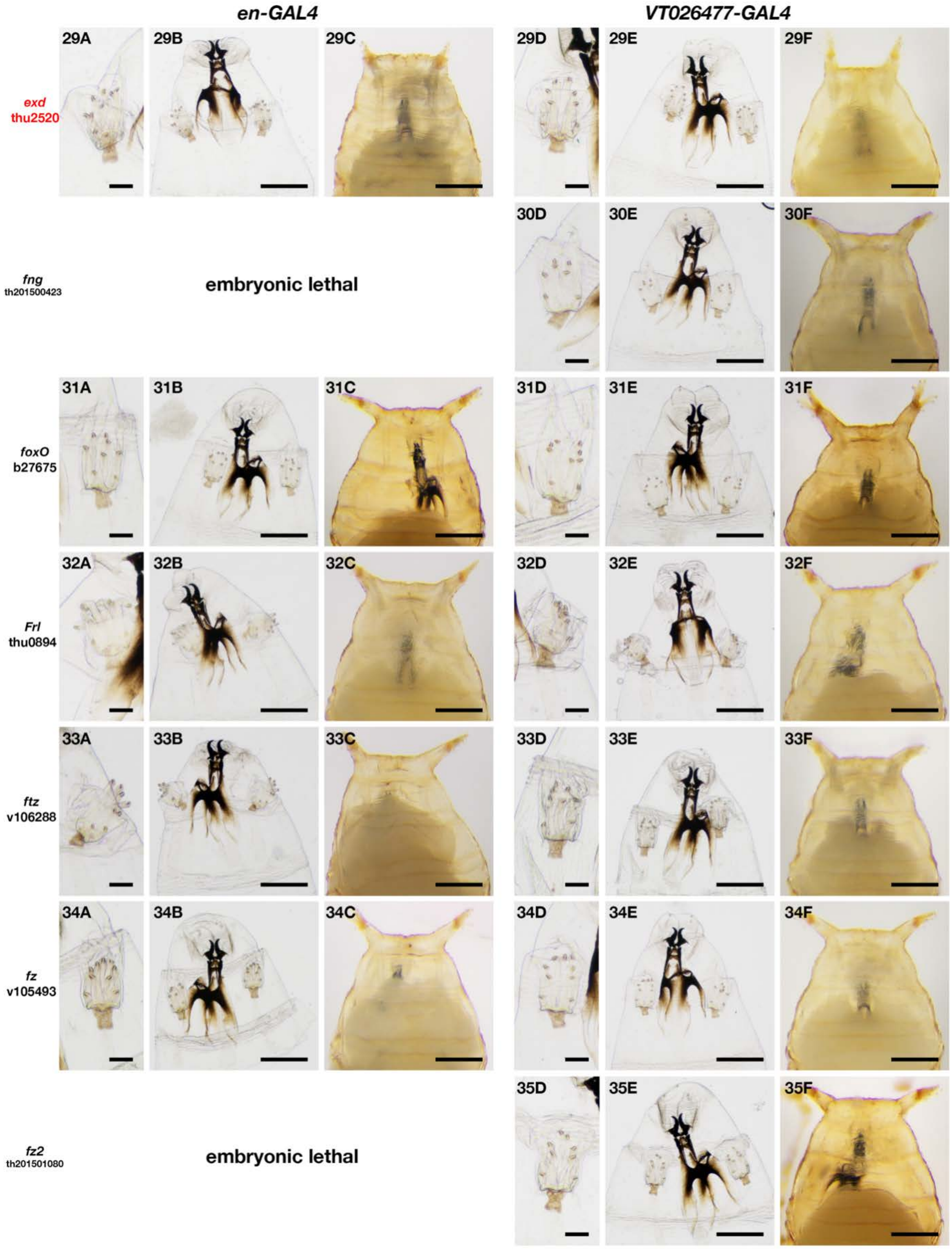


Figure S14 continued to next page

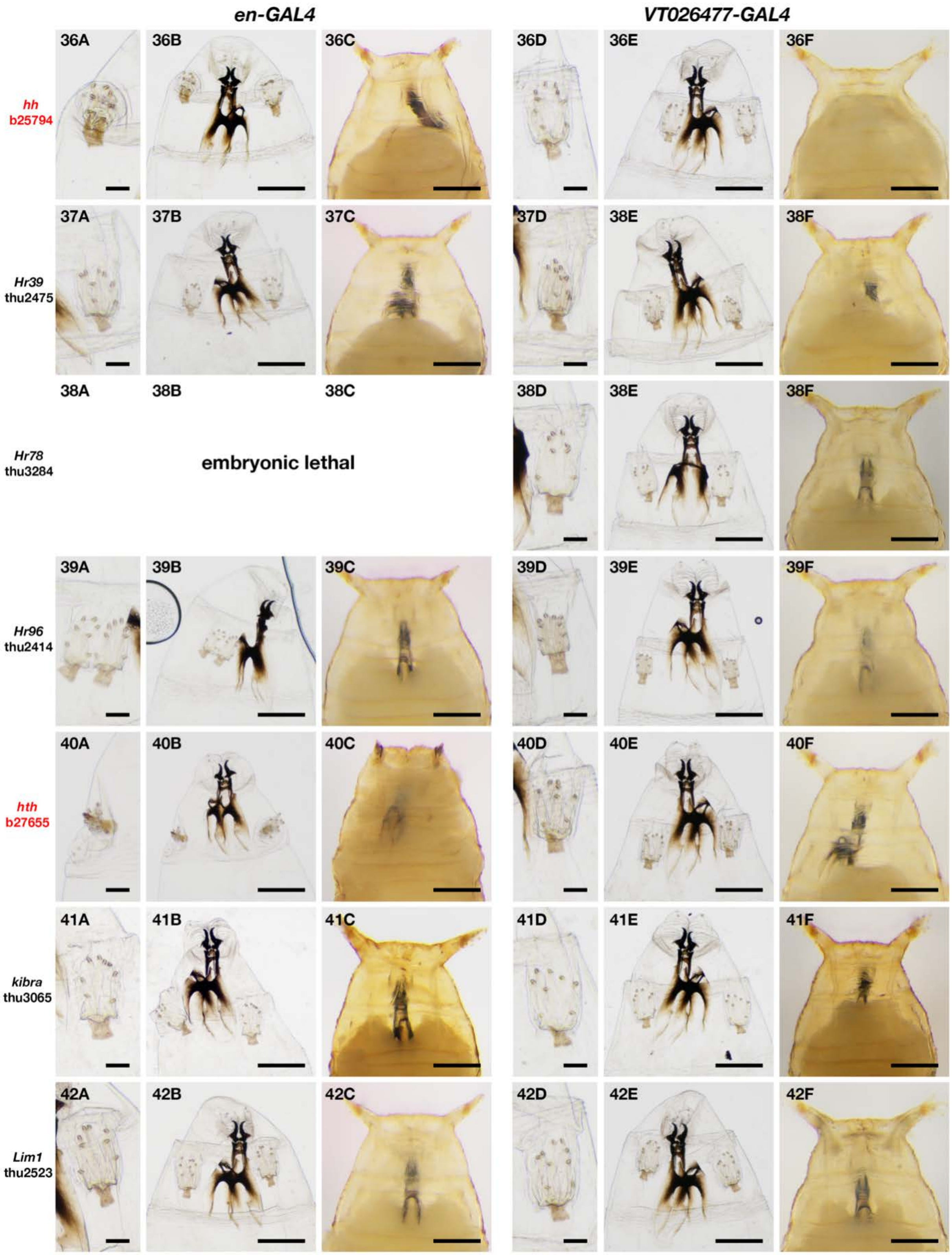


Figure S14 continued to next page

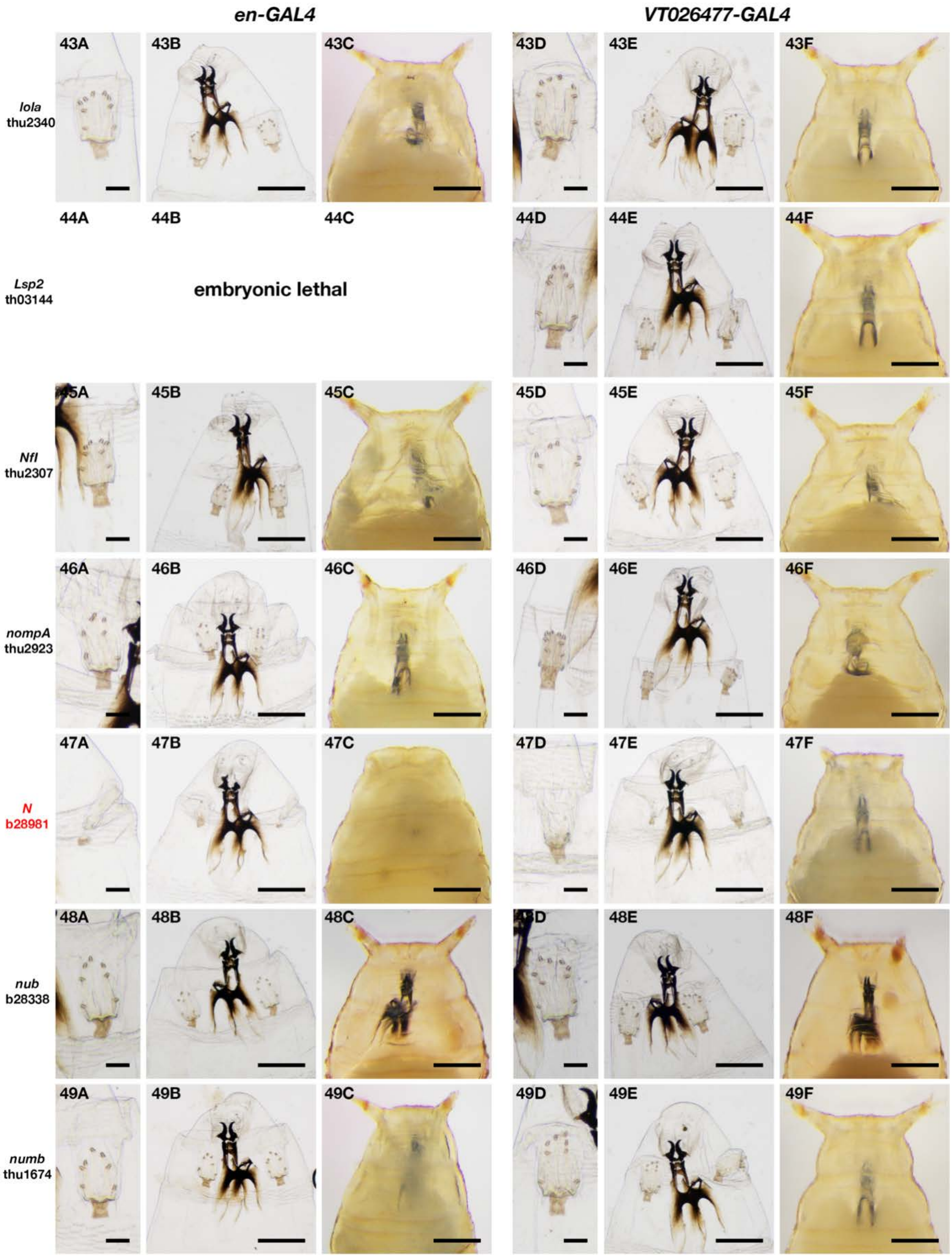


Figure S14 continued to next page

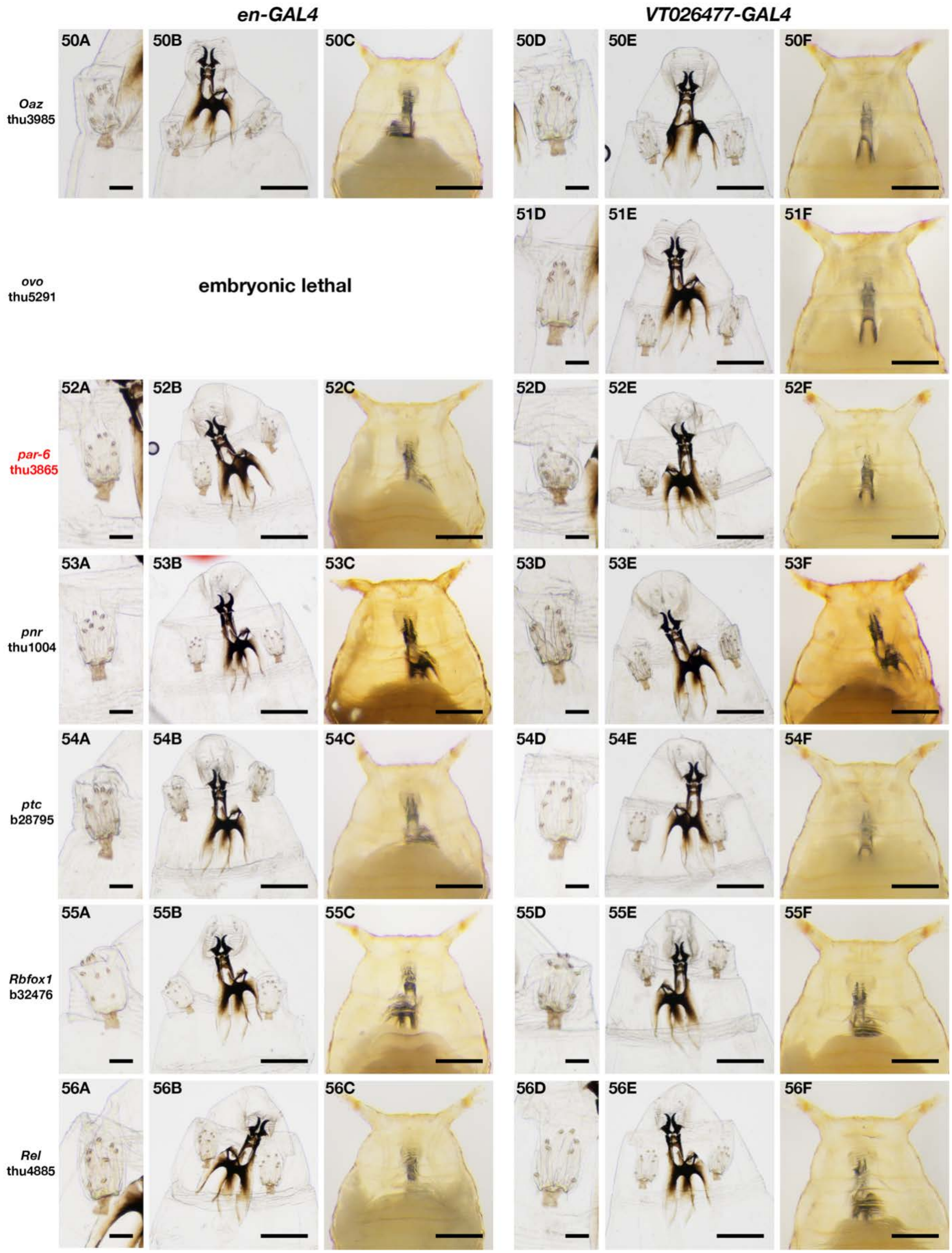


Figure S14 continued to next page

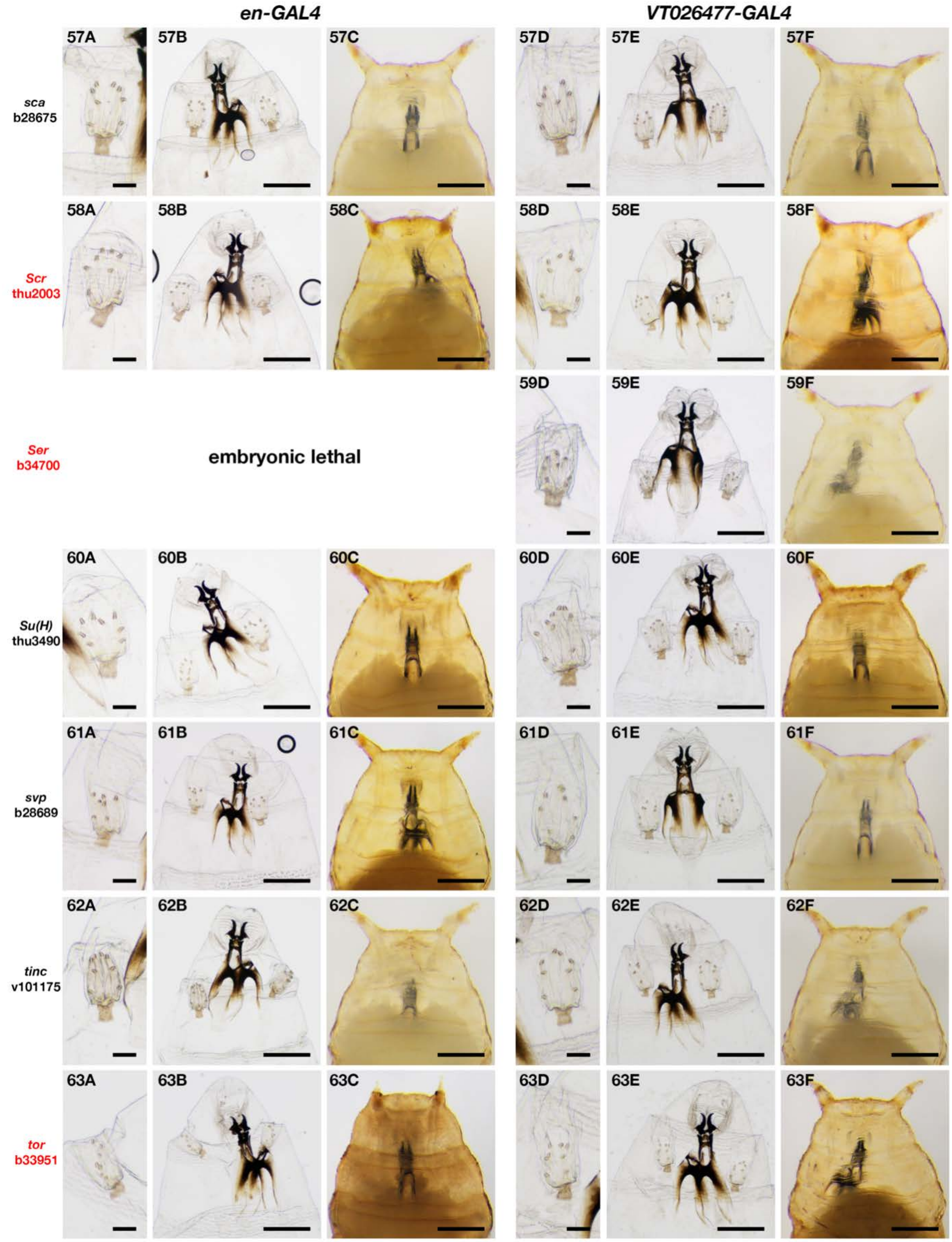


Figure S14 continued to next page

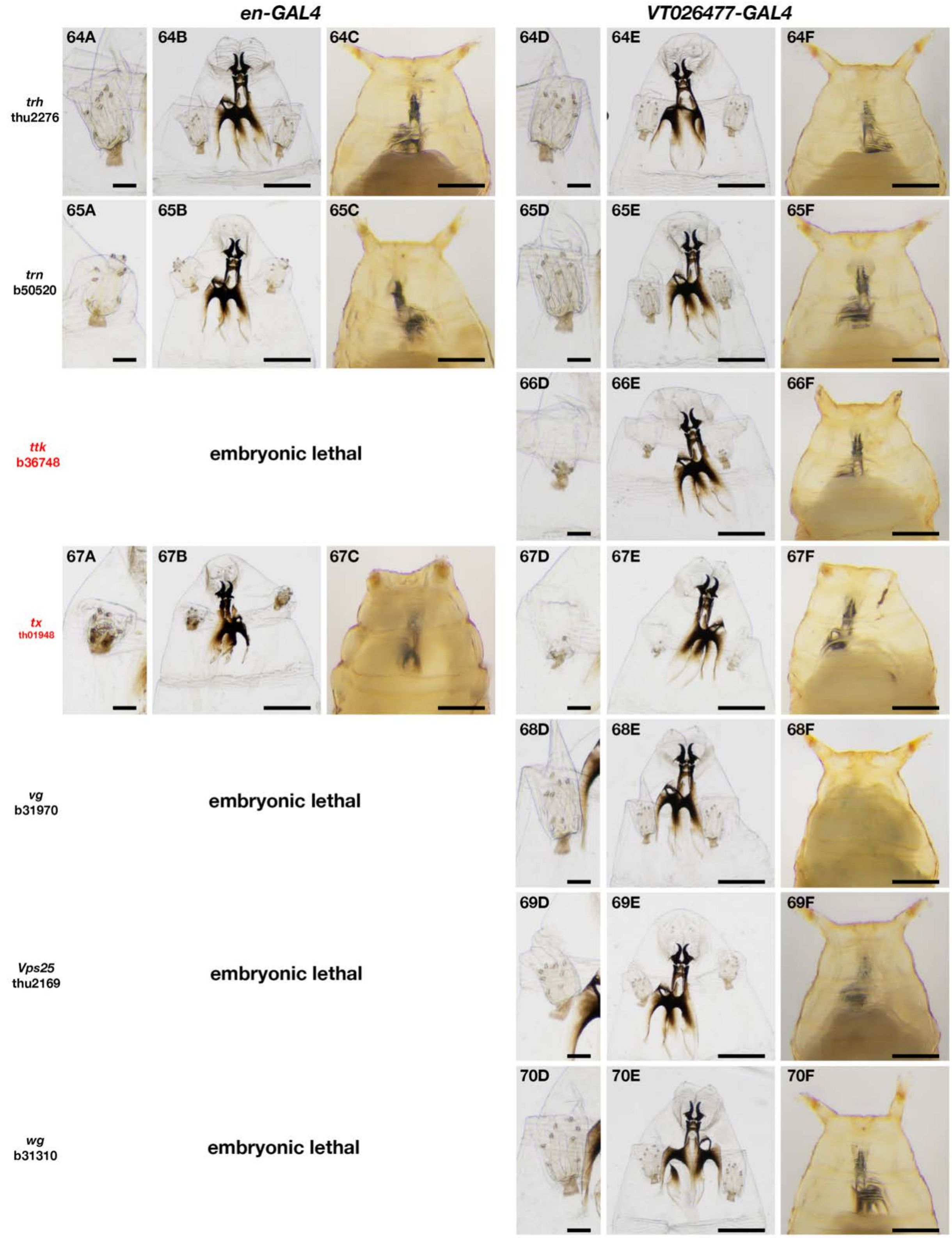


Figure S14 continued to next page

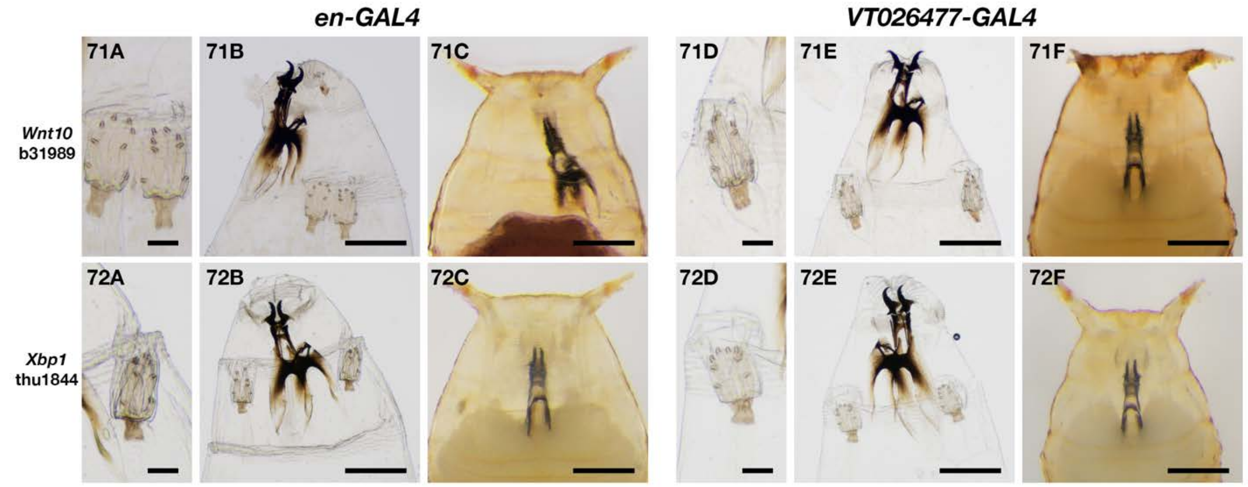


**Figure S14. Systematic assessment of gene function in the *D. melanogaster* PROD, focusing on co-opted wing genes, via RNAi driven by *en-GAL4* and *VT026477-GAL4*.** (**1A-1C**) The PROD morphologies of wildtype Canton-S *D. melanogaster* with schematic representation of expected phenotypes: the number of branches and length of the everted cone. (**2A-2F**) The PROD morphologies of the two *GAL4* controls. (**3A-3C:72A-72C**) The PROD morphologies of the *en-GAL4* RNAi knockdowns. (**3D-3F:72D-72F**) The PROD morphologies of the *VT026477-GAL4* RNAi knockdowns. Fly stock line see Table S7. Scalebar in magnified PROD panels=50 µm; Scalebar in larva and pupa images=250 µm.

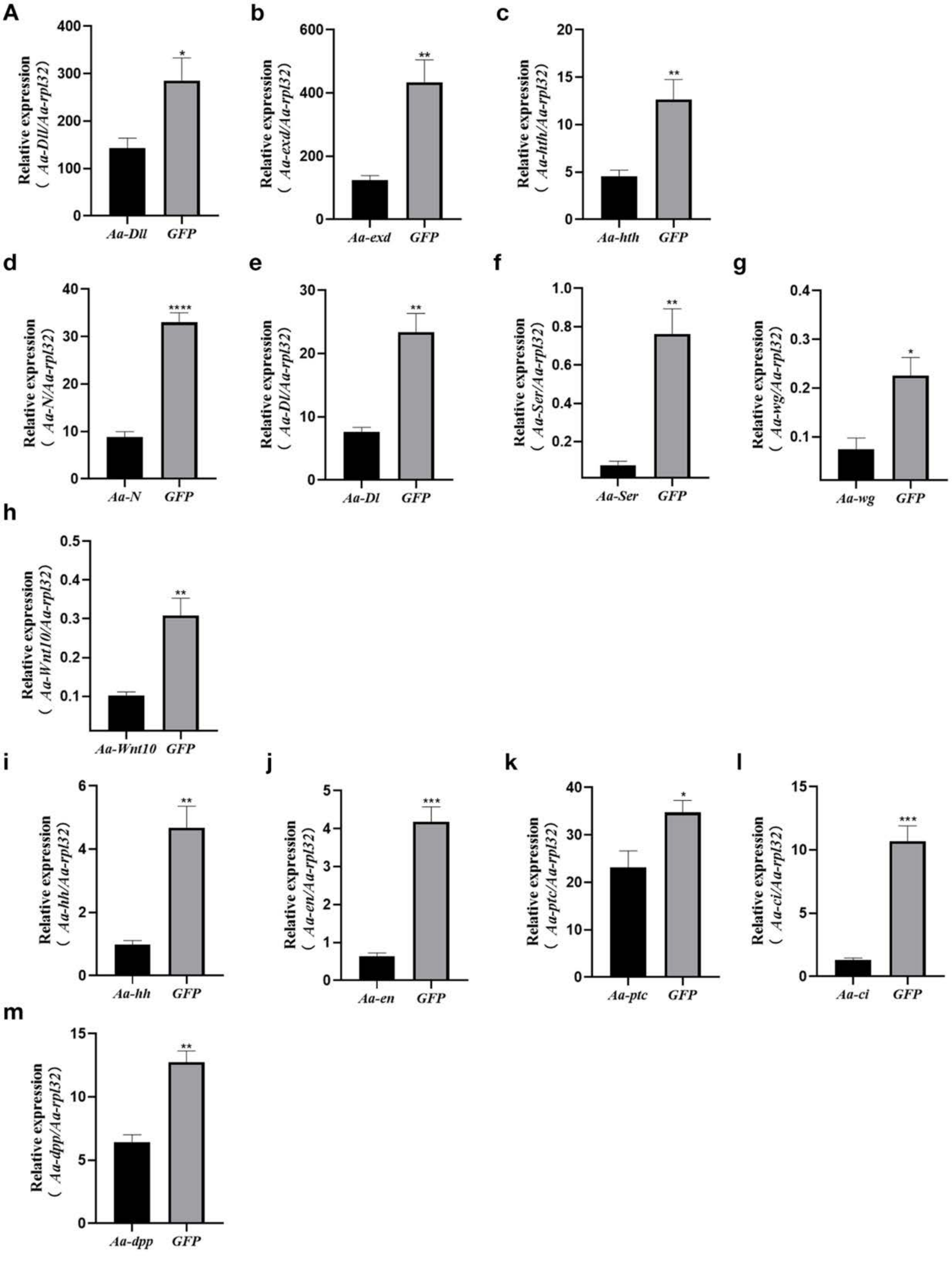


**Figure S15. qPCR assessment on the RNAi** knockdown **efficiencies of the appendage patterning genes in *Aedes*. (a-c)** The knockdown efficiency of the P/D morphogens: **(a)** *Aa-Dll*, **(b)** *Aa-exd*, and (**c**) *Aa-hth*. **(d-h)** The knockdown efficiency of the D/V morphogens: **(d)** *Aa-N*, **(e)** *Aa-Dl*, **(f)** *Aa-Ser*, **(g)** *Aa-wg*, and **(h)** *Aa-Wnt10*. **(i-m)** The knockdown efficiency of the A/P morphogens: **(i)** *Aa-hh*, **(j)** *Aa-en*, **(k)** *Aa-ptc*, **(l)** *Aa-ci*, and **(m)** *Aa-dpp*. The asterisks represent statistical significancy: (*) $p$<0.05; (**) $p$<0.01; (***) $p$<0.001; (****) $p$<0.0001.

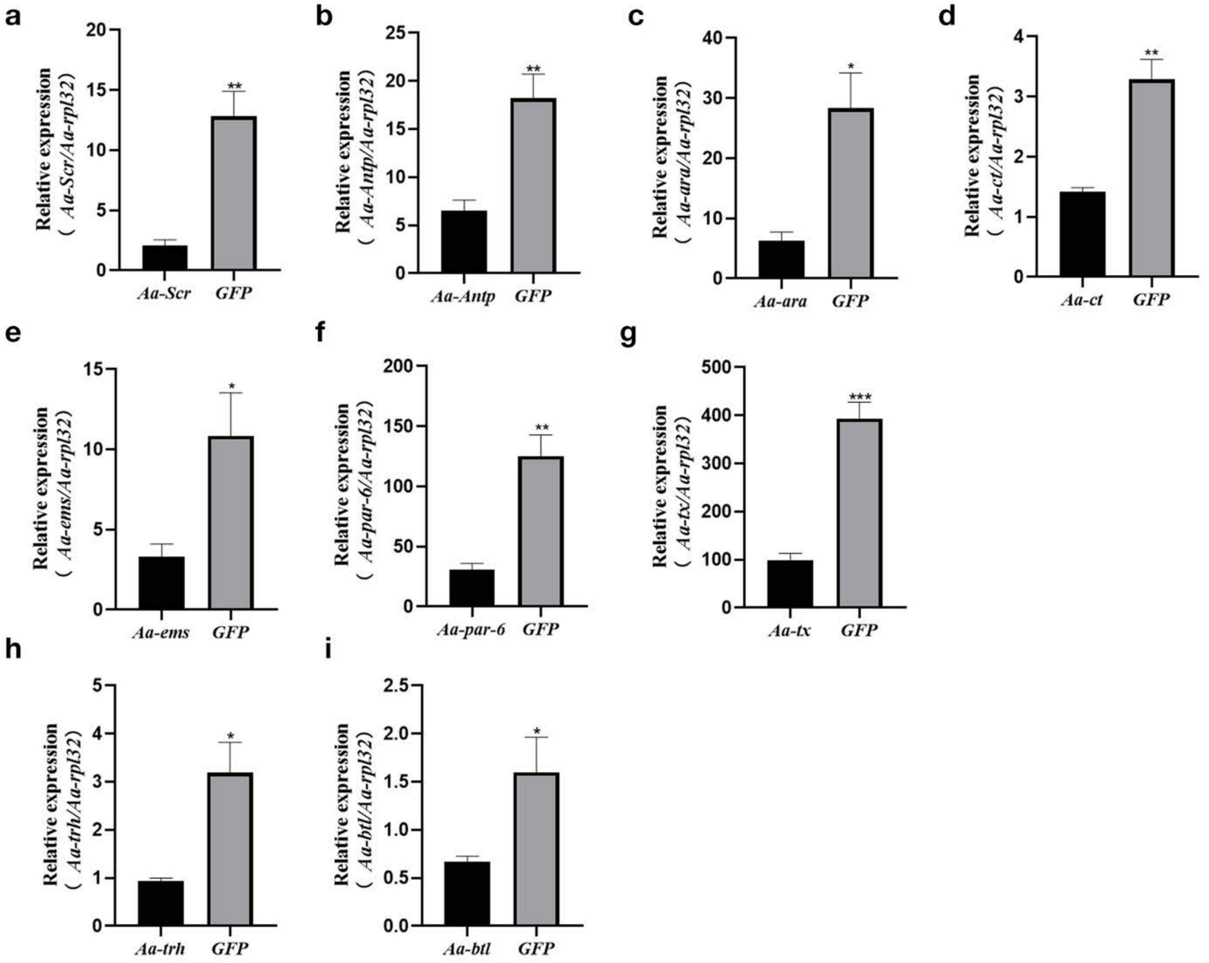


**Figure S16. qPCR assessment on the RNAi knockdown efficiency of the homeobox genes and tracheal morphogens in *Aedes*. (a-g)** The knockdown efficiency of the homeobox genes: **(a)** *Aa-Scr*, **(b)** *Aa-Antp*, **(c)** *Aa-ara*, **(d)** *Aa-ct*, **(e)** *Aa-ems*, **(f)** *Aa-par-6*, and **(g)** *Aa-tx*. **(h, i)** The knockdown efficiency of the tracheal morphogens: **(h)** *Aa-trh*, **(i)** *Aa-btl*. The asterisk symbols represent statistical significancy where: (*) $p$>0.05; (**) $p$>0.01; (***) $p$>0.001; (****) $p$>0.0001.

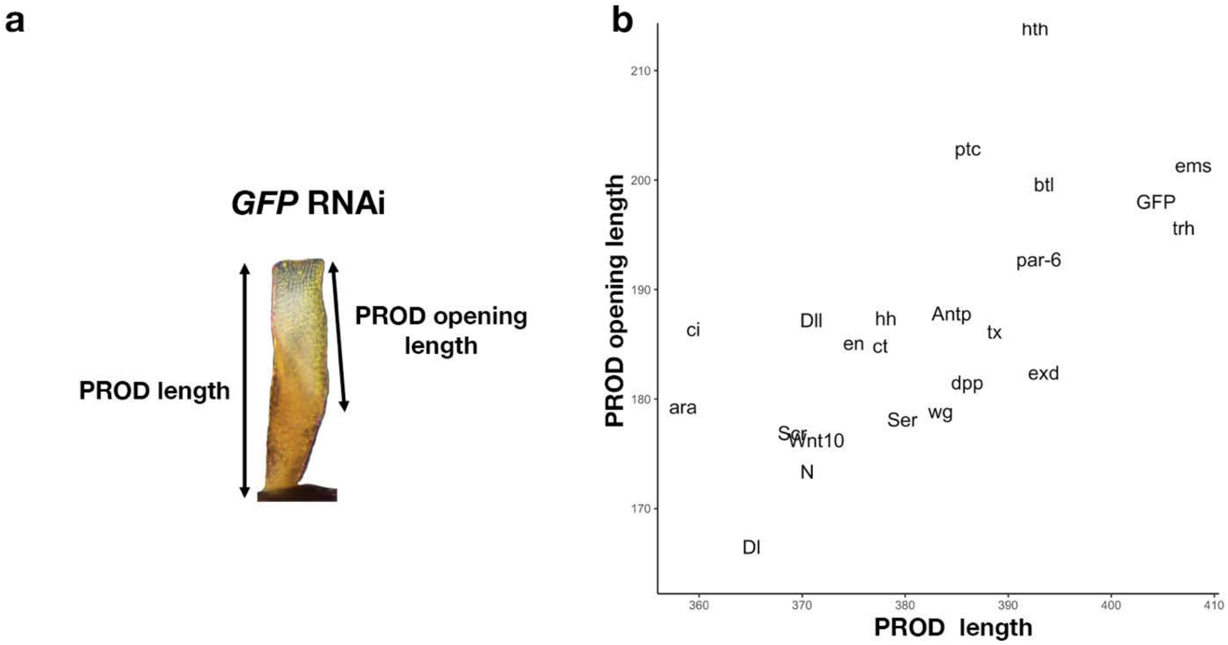


**Figure S17. Morphometric assessment of *A. albopictus* PROD in RNAi knockdown mutants. (a)** schematic representation of the two measurements: length of the PROD and the length of the PROD opening. **(b)** scatterplot of the two measurements of each gene knockdown, averaged across all replicates including those without observable phenotypes.

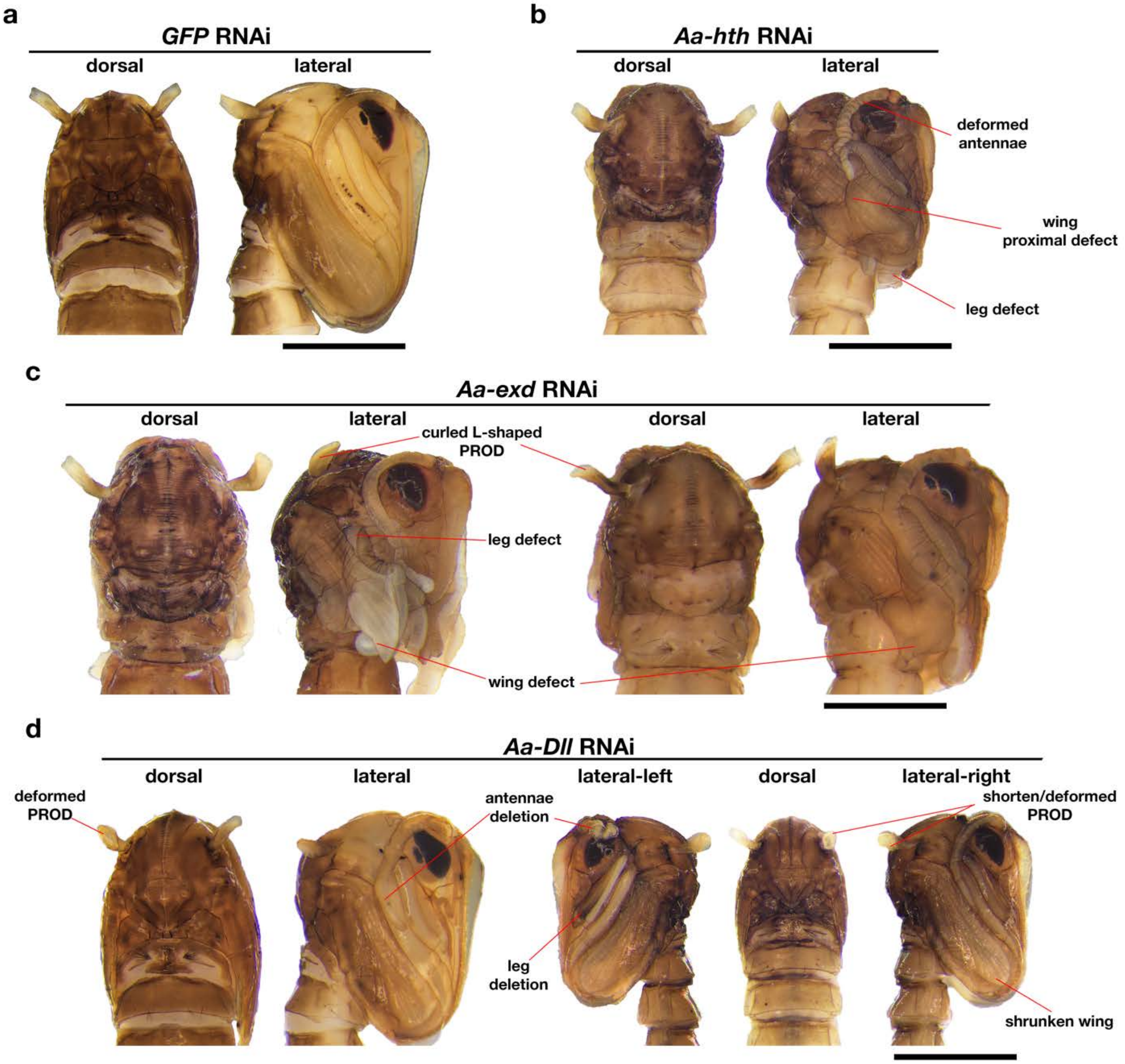


**Figure S18. RNAi reveals that the *A. albopictus* PROD development depends on the P/D patterning genes. (a)** RNAi of GFP in *A. albopictus* pupa shows no phenotype and serves as the control. **(b–d)**, Representative images of *A. albopictus* pupae of RNAi targeting the P/D axis genes: **(b)** *Aa-Dll*, **(c)** *Aa-exd*, and **(d)** *Aa-hth*. Phenotypic changes are indicated by red lines, highlighting defects in appendages (e.g., wings, legs, and antennae) including the PROD, aligning with the reported morphological defects in *D. melanogaster*. Scale bar=500 µm.

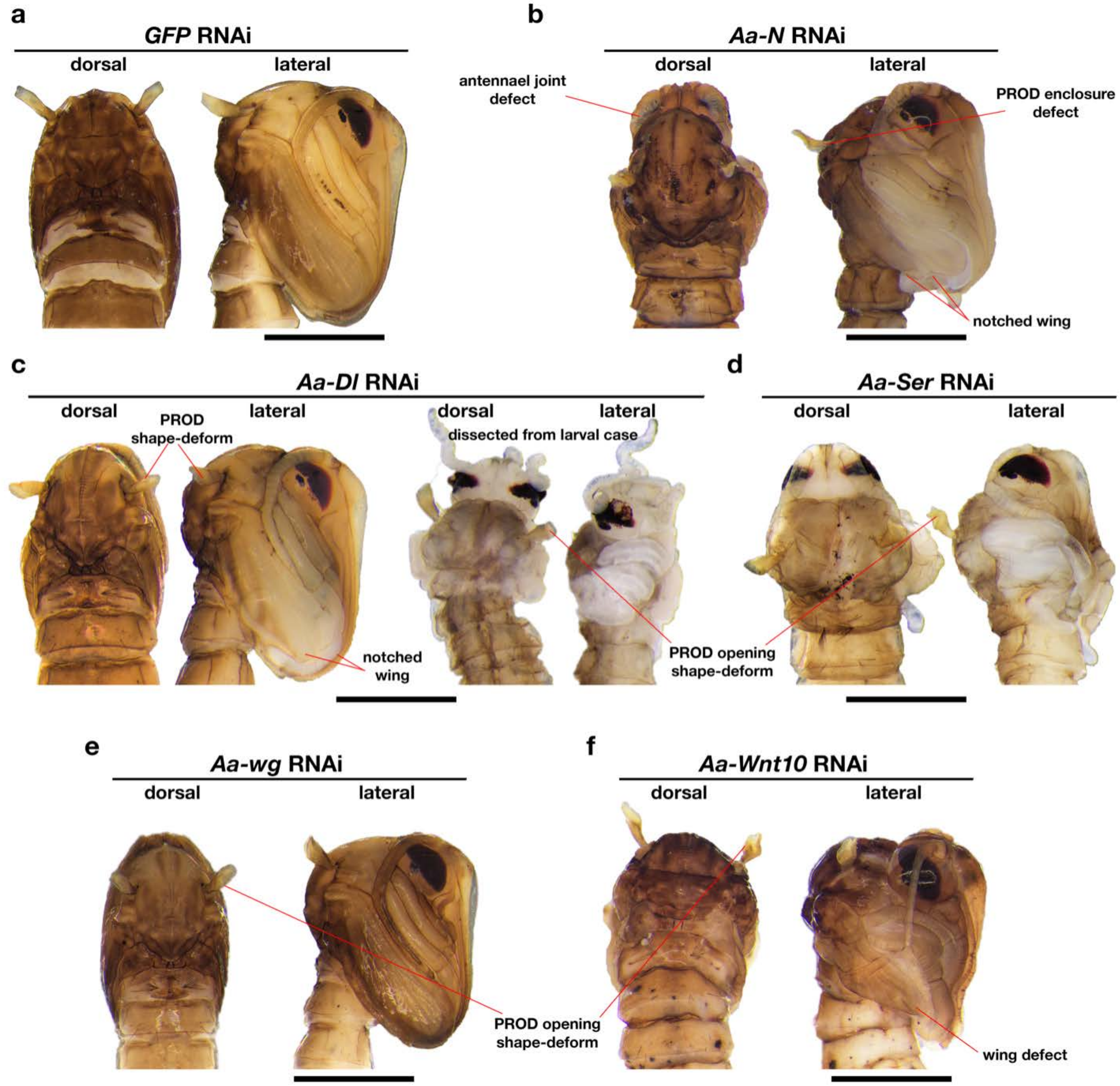


**Figure S19. RNAi reveals that the *A. albopictus* PROD development depends on the D/V patterning genes. (a)** RNAi of GFP in *A. albopictus* pupa shows no phenotype and serves as the control. **(b–d)** Representative images of *A. albopictus* pupae of RNAi targeting the *A. albopictus N/Dl* signaling pathway: **(b)** *Aa-N*, **(c)** *Aa-Dl*, and **(d)** *Aa-Ser*. **(e, f)** Representative images of *A. albopictus* pupae of RNAi targeting **(e)** *Aa-wg* and **(f)** *Aa-Wnt10*. Phenotypic changes are indicated by red lines, highlighting alterations in appendages (e.g., wings, legs, and antennae) including the PROD, aligning with the reported morphological defects in *D. melanogaster*, especially the notched wing and the joint defect of the antennal formation. Scale bar=500 μm.

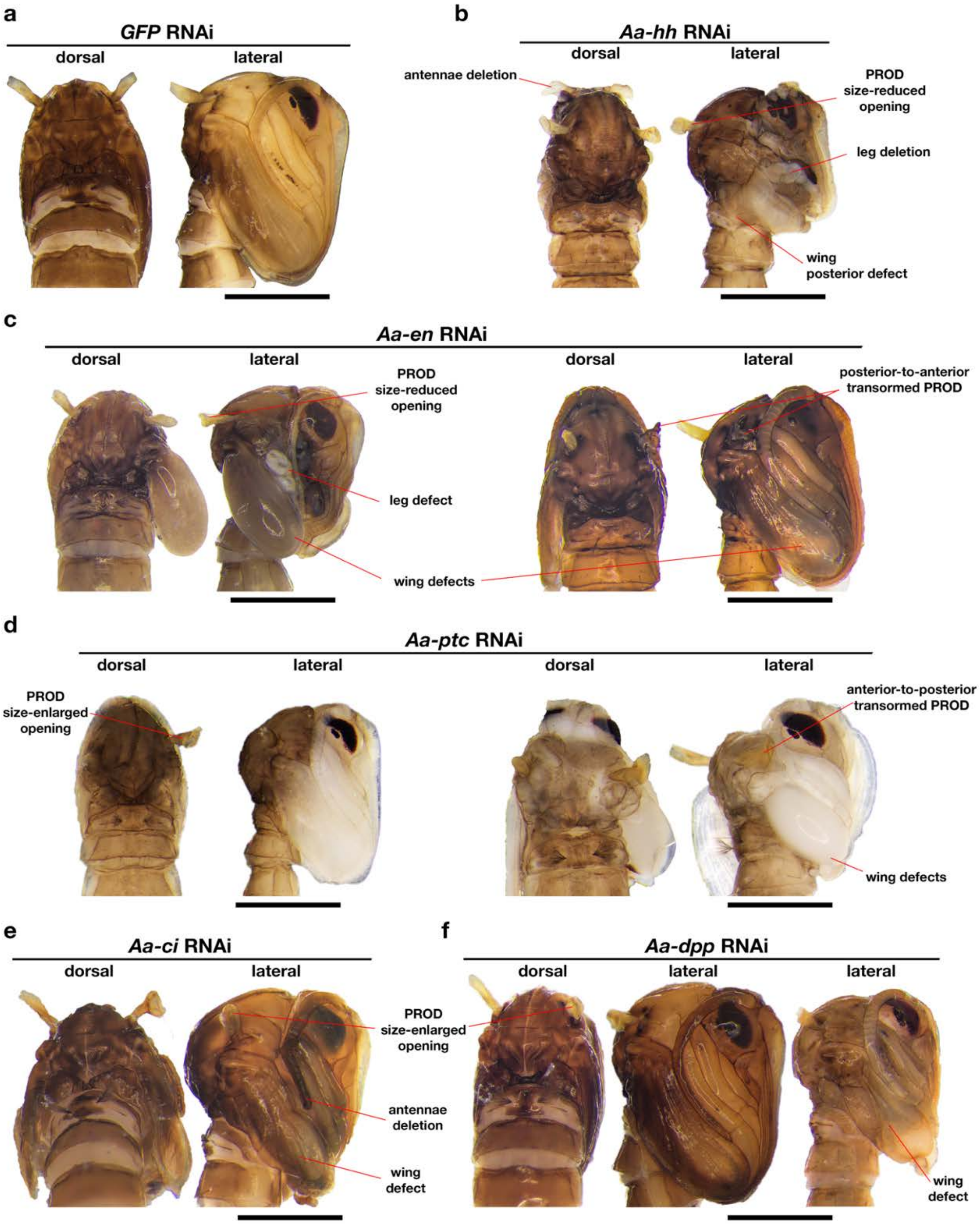


**Figure S20. RNAi reveals that the *A. albopictus* PROD development depends on the A/P patterning genes.** **(a)** RNAi of GFP in *A. albopictus* pupa shows no phenotype and serves as the control. **(b, c)** Representative images of *A. albopictus* pupae of RNAi targeting the posterior genes: **(b)** *Aa-hh* and **(c)** *Aa-en*. **(d–f)** Representative images of *A. albopictus* pupae of RNAi targeting the anterior genes: **(d)** *Aa-ptc*, **(e)** *Aa-ci*, and **(f)** *Aa-dpp*. Phenotypic changes are indicated by red lines, highlighting alterations in appendages (e.g., wings, legs, and antennae) including the PROD, aligning with reported morphological defects in *D. melanogaster*. Notably, disruption of the A/P genes in *A. albopictus* PROD produces a class of phenotype that resembles the anterior-posterior mirror image transformation. Scale bar=500 µm.

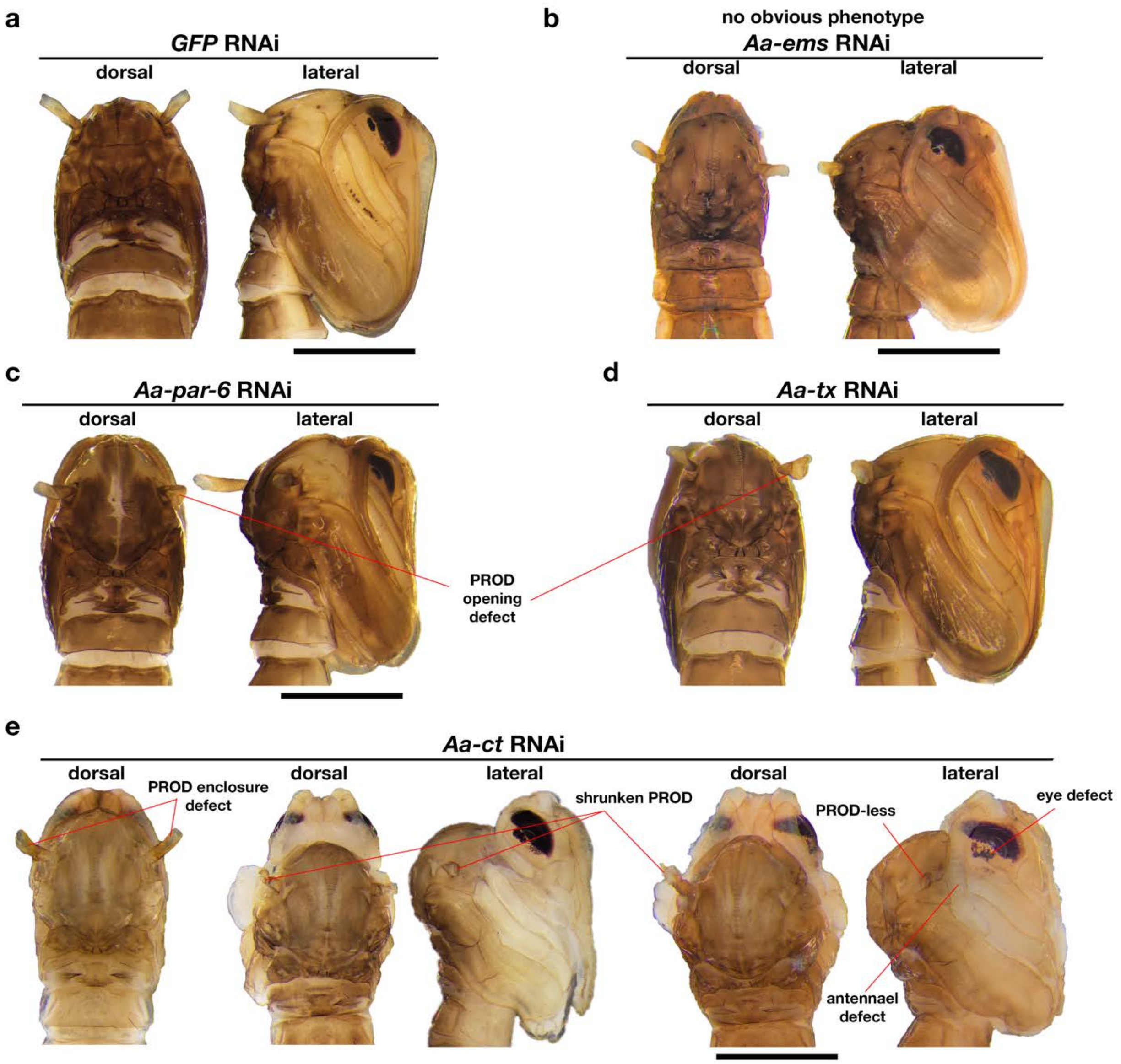


**Figure S21. RNAi of candidate transcriptional factors for finding the selector gene of *A. albopictus* PROD. (a)** The *dsGFP* injection control. **(b)** RNAi knockdowns of *Aa-ems* do not yield observable morphological phenotype in the PROD. **(c)** RNAi knockdowns of *Aa-par-6* yield deformed PROD openings. **(d)** RNAi knockdowns of *Aa-tx* yield deformed PROD openings. None of these genes cause the PROD-less phenotypes that have been observed in knockdowns of *ct* in both *A. albopictus* and *D. melanogaster*, suggesting that *ct* is the main selector gene of PROD fate. **(e)** Representative images of *A. albopictus* pupae of RNAi targeting *Aa-ct*. Phenotypic changes are indicated by red lines, highlighting alterations in appendages (e.g., wings) including the PROD, aligning with reported morphological defects in other insects. Scale bars: 500 µm.

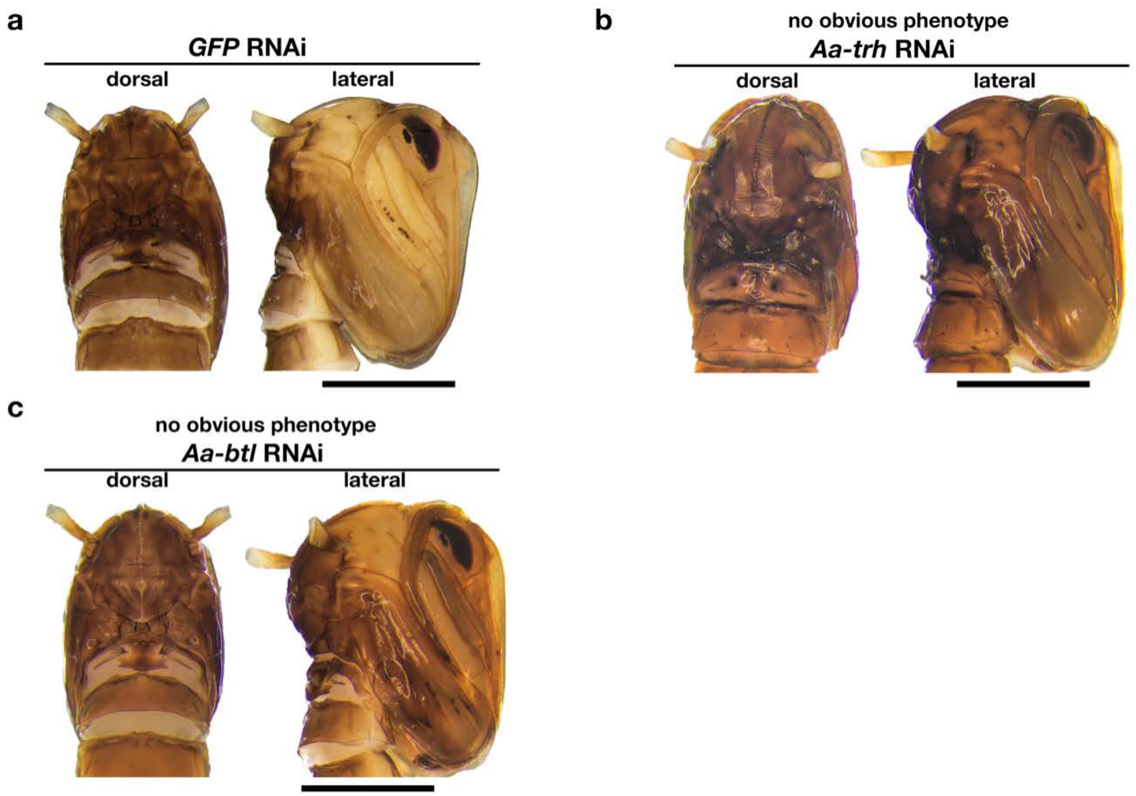


**Figure S22. Functional assessment of the tracheal GRN in *A. albopictus* PROD. (a)** RNAi of GFP in *A. albopictus* pupa shows no phenotype and serves as the control. **(b)** Representative images of *A. albopictus* pupae of RNAi targeting *Aa-trh*. **(c)** Representative images of *A. albopictus* pupae of RNAi targeting *Aa-btl*. Scalebar=500 µm.

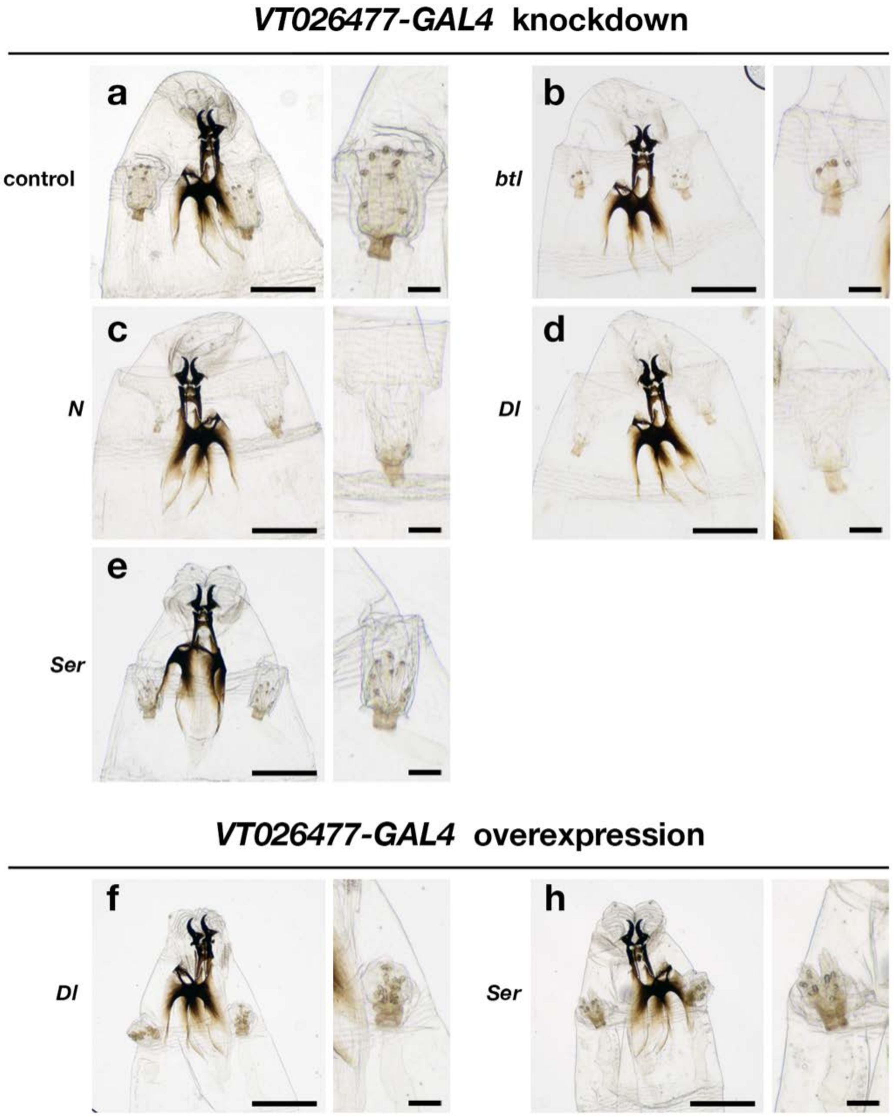


**Figure S23. Functional assessment of the tracheal GRN in *D. melanogaster* PROD. (a-e)** Functional assessment of morphogens of tracheal GRN in *D. melanogaster* using *VT026477-GAL4*. Scalebar=250 µm. **(a)** Uncrossed *VT026477-GAL4* served as control. Knockdown of *btl* **(b)**, *N* **(c)**, *Dl* **(d)**, and *Ser* **(e)** demonstrates primary GRN component that controls PROD spiracular branch development. Overexpression of *Dl* **(f)** and *Ser* **(h)** show how the number of branches and terminal openings of PROD spiracle are determined.

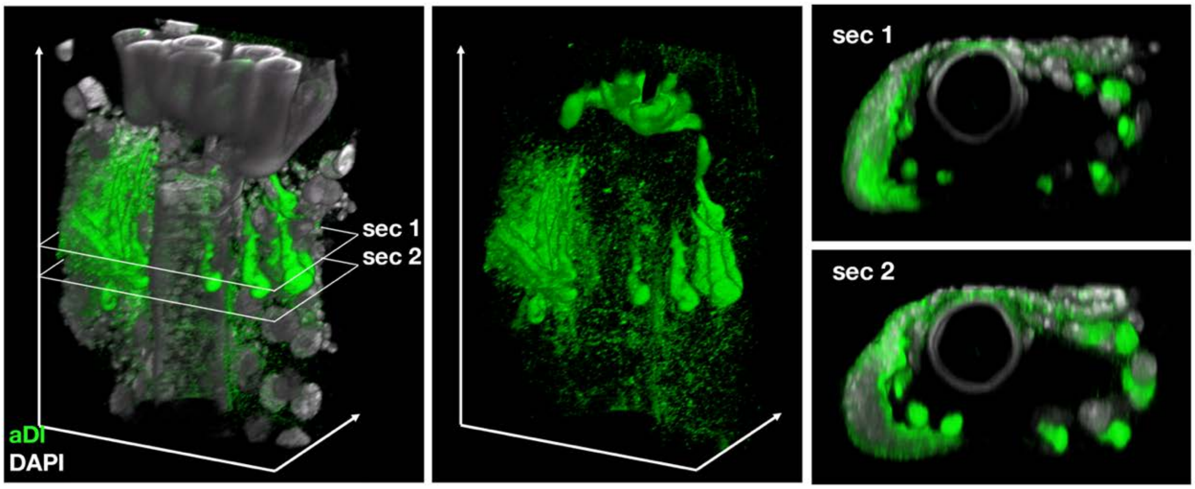


**Figure S24. Immunol staining illustrating *Dl* expression pattern foreshadows the number branch tips in the *D. melanogaster* PROD development.** Scale bars: 100 µm.

**Table S1. Species and genomic data used for phylogenetic reconstruction of Diptera.**

| Order | Family | Genus | Species | Accession |
|---|---|---|---|---|
| Coleoptera | Scarabaeidae | *Onthophagus* | *Onthophagus taurus* | GCA_000648695.2 |
| Coleoptera | Scarabaeidae | *Trypoxylus* | *Trypoxylus dichotomus* | GCA_023509865.1 |
| Coleoptera | Ostomatidae | *Tribolium* | *Tribolium castaneum* | GCA_000002335.3 |
| Diptera | Acartophthalmidae | *Acartophthalmus* | *Acartophthalmus nigrinus* | SRR10426012 |
| Diptera | Acroceridae | *Pterodontia* | *Pterodontia mellii* | SRR1695385 |
| Diptera | Agromyzidae | *Liriomyza* | *Liriomyza trifolii* | SRR1138236 |
| Diptera | Anisopodidae | *Sylvicola* | *Sylvicola fuscatus* | GCA_026546915.1 |
| Diptera | Anthomyiidae | *Delia* | *Delia floralis* | SRR17845154 |
| Diptera | Anthomyiidae | *Delia* | *Delia radicum* | SRR14802799 |
| Diptera | Anthomyzidae | *Mumetopia* | *Mumetopia occipitalis* | SRR1695368 |
| Diptera | Apioceridae | *Apiocera* | *Apiocera maritima* | SRR2046562 |
| Diptera | Apioceridae | *Apiocera* | *Apiocera moerens* | SRR1695318 |
| Diptera | Apystomyiidae | *Apystomyia* | *Apystomyia elinguis* | SRR1964413 |
| Diptera | Asilidae | *Diogmites* | *Diogmites neoternatus* | SRR4345333 |
| Diptera | Asilidae | *Eudioctria* | *Eudioctria media* | SRR10386632 |
| Diptera | Asilidae | *Philonicus* | *Philonicus albiceps* | SRR4365562 |
| Diptera | Asilidae | *Tolmerus* | *Tolmerus atricapillus* | SRR4346294 |
| Diptera | Asteiidae | *Leiomyza* | *Leiomyza laevigata* | SRR10426011 |
| Diptera | Atelestidae | *Meghyperus* | *Meghyperus* sp. | SRR10425999 |
| Diptera | Athericidae | *Atherix* | *Atherix ibis* | GCA_958298945.1 |
| Diptera | Aulacigastridae | *Aulacigaster* | *Aulacigaster mcalpinei* | SRR10425996 |
| Diptera | Australimyzidae | *Australimyza* | *Australimyza mcalpinei* | SRR10425995 |
| Diptera | Axymyiidae | *Axymyia* | *Axymyia furcata* | SRR1695322 |
| Diptera | Bibionidae | *Bibio* | *Bibio marci* | GCA_910594885.2 |
| Diptera | Bibionidae | *Dilophus* | *Dilophus febrilis* | GCA_958336335.1 |
| Diptera | Blephariceridae | *Blepharicera* | *Blepharicera* sp. | SRR1695325 |
| Diptera | Blephariceridae | *Philorus* | *Philorus* sp. | This study |
| Diptera | Blephariceridae | unknown | Blephariceridae sp. WYM | SRR32672061 |
| Diptera | Bolitophilidae | *Bolitophila* | *Bolitophila cinerea* | GCA_010015015.2 |
| Diptera | Bombyliidae | *Anthrax* | *Anthrax maculatus* | SRR10799664 |
| Diptera | Bombyliidae | *Comptosia* | *Comptosia australensis* | SRR10766074 |
| Diptera | Bombyliidae | *Conophorus* | *Conophorus* sp. | SRR10766072 |
| Diptera | Bombyliidae | *Geron* | *Geron flavocciput* | SRR10766073 |
| Diptera | Bombyliidae | *Mandella* | *Mandella* sp. | SRR10766188 |
| Diptera | Bombyliidae | *Meomyia* | *Meomyia vetusta* | SRR10766264 |
| Diptera | Bombyliidae | *Staurostichus* | *Staurostichus limbipennis* | SRR10766168 |
| Diptera | Bombyliidae | *Thevenetimyia* | *Thevenetimyia longipalpus* | SRR10766092 |
| Diptera | Bombyliidae | *Thraxan* | *Thraxan patielus* | SRR10766289 |
| Diptera | Bombyliidae | *Toxophora* | *Toxophora* sp. | SRR11700305 |
| Diptera | Bombyliidae | *Villa* | *Villa fuscicostata* | SRR10799685 |
| Diptera | Braulidae | *Braula* | *Braula coeca* | SRR2046564 |
| Diptera | Calliphoridae | *Bixinia* | *Bixinia* sp. | SRR6724163 |
| Diptera | Calliphoridae | *Calliphora* | *Calliphora augur* | SRR11309594 |
| Diptera | Calliphoridae | *Lucilia* | *Lucilia cuprina* | GCA_022045245.1 |

| Order | Family | Genus | Species | Accession |
|---|---|---|---|---|
| Diptera | Calliphoridae | *Stevenia* | *Stevenia* sp. | SRR6724161 |
| Diptera | Calliphoridae | *Stomorhina* | *Stomorhina subapicalis* | ERR10123679 |
| Diptera | Canacidae | *Dasyrhicnoessa* | *Dasyrhicnoessa insularis* | SRR10425994 |
| Diptera | Carnidae | *Carnus* | *Carnus hemapterus* | SRR10694694 |
| Diptera | Cecidomyiidae | *Aphidoletes* | *Aphidoletes aphidimyza* | GCA_030463065.1 |
| Diptera | Cecidomyiidae | *Catotricha* | *Catotricha subobsoleta* | GCA_011634745.2 |
| Diptera | Cecidomyiidae | *Contarinia* | *Contarinia nasturtii* | GCA_009176525.2 |
| Diptera | Cecidomyiidae | *Lestremia* | *Lestremia cinerea* | GCA_027564135.1 |
| Diptera | Cecidomyiidae | *Mayetiola* | *Mayetiola destructor* | GCA_000149185.1 |
| Diptera | Cecidomyiidae | *Obolodiplosis* | *Obolodiplosis robiniae* | GCA_028476595.1 |
| Diptera | Cecidomyiidae | *Resseliella* | *Resseliella maxima* | GCA_029041755.1 |
| Diptera | Cecidomyiidae | *Sitodiplosis* | *Sitodiplosis mosellana* | GCA_009176505.1 |
| Diptera | Ceratopogonidae | *Atrichopogon* | *Atrichopogon* sp. | SRR6155951 |
| Diptera | Ceratopogonidae | *Culicoides* | *Culicoides sonorensis* | SRR1174038 |
| Diptera | Ceratopogonidae | *Dasyhelea* | *Dasyhelea* sp. | SRR6155934 |
| Diptera | Ceratopogonidae | *Forcipomyia* | *Forcipomyia taiwana.merge* | ERR11267955, ERR11267956, ERR11267957 |
| Diptera | Chamaemyiidae | *Pseudodinia* | *Pseudodinia antennalis* | SRR1695384 |
| Diptera | Chaoboridae | *Chaoborus* | *Chaoborus americanus.merge* | SRR18143961, SRR18143972, SRR18143973 |
| Diptera | Chaoboridae | *Chaoborus* | *Chaoborus flavidulus.merge* | SRR1695304, SRR6155946 |
| Diptera | Chaoboridae | *Mochlonyx* | *Mochlonyx cinctipes* | GCA_001014845.1 |
| Diptera | Chironomidae | *Cardiocladius* | *Cardiocladius* sp. | SRR6155936 |
| Diptera | Chironomidae | *Chironomus* | *Chironomus tentans* | GCA_963573255.1 |
| Diptera | Chironomidae | *Clunio* | *Clunio marinus* | GCA_900005825.1 |
| Diptera | Chironomidae | *Cricotopus* | *Cricotopus draysoni* | SRR4457915 |
| Diptera | Chironomidae | *Diamesa* | *Diamesa zernyi* | SRR13641921 |
| Diptera | Chironomidae | *Kiefferophyes* | *Kiefferophyes invenustulus* | SRR6155941 |
| Diptera | Chironomidae | *Paraheptagyia* | *Paraheptagyia tonnoiri* | SRR6155948 |
| Diptera | Chironomidae | *Paraphaenocladius* | *Paraphaenocladius impensus* | SRR11637888 |
| Diptera | Chironomidae | *Parochlus* | *Parochlus steinenii.merge* | SRR3951283, SRR3951284, SRR3951285 |
| Diptera | Chironomidae | *Pentaneurella* | *Pentaneurella katterjokki* | SRR11637887 |
| Diptera | Chironomidae | *Podonomus* | *Podonomus* sp. | SRR6155949 |
| Diptera | Chironomidae | *Polypedilum* | *Polypedilum pembai* | GCA_014622435.1 |
| Diptera | Chironomidae | *Polypedilum* | *Polypedilum vanderplanki* | GCA_018290095.1 |
| Diptera | Chironomidae | *Procladius* | *Procladius villosimanus* | SRR6155953 |
| Diptera | Chironomidae | *Propsilocerus* | *Propsilocerus akamusi* | GCA_018397935.1 |
| Diptera | Chironomidae | *Smittia* | *Smittia aterrima* | GCA_033063855.1 |
| Diptera | Chironomidae | *Telmatogeton* | *Telmatogeton pectinata* | SRR6155952 |
| Diptera | Chironomidae | *Telmatopelopia* | *Telmatopelopia nemorum* | SRR6155950 |
| Diptera | Chloropidae | *Chlorops* | *Chlorops oryzae* | SRR7534658 |
| Diptera | Chloropidae | *Lipara* | *Lipara lucens* | SRR921612 |

| Order | Family | Genus | Species | Accession |
|---|---|---|---|---|
| Diptera | Chyromyidae | *Gymnochiromyia* | *Gymnochiromyia* sp. | SRR10425993 |
| Diptera | Clusiidae | *Clusia* | *Clusia lateralis* | SRR1695305 |
| Diptera | Coelopidae | *Coelopa* | *Coelopa frigida* | SRR15116463 |
| Diptera | Conopidae | *Myopa* | *Myopa* sp. | SRR1695369 |
| Diptera | Corethrellidae | *Corethrella* | *Corethrella appendiculata* | SRR951913, SRR952150 |
| Diptera | Corethrellidae | *Corethrella* | *Corethrella calathicola* | SRR6155947 |
| Diptera | Cryptochetidae | *Cryptochetum* | *Cryptochetum* sp. | SRR10425992 |
| Diptera | Culicidae | *Aedes* | *Aedes aegypti* | GCA_002204515.1 |
| Diptera | Culicidae | *Aedes* | *Aedes albopictus* | GCA_006496715.1 |
| Diptera | Culicidae | *Anopheles* | *Anopheles cracens* | GCA_002091845.1 |
| Diptera | Culicidae | *Anopheles* | *Anopheles gambiae* | GCA_943734735.2 |
| Diptera | Culicidae | *Anopheles* | *Anopheles sinensis* | GCA_000441895.2 |
| Diptera | Culicidae | *Anopheles* | *Anopheles stephensi* | GCA_013141755.1 |
| Diptera | Culicidae | *Armigeres* | *Armigeres subalbatus* | GCA_024139115.1 |
| Diptera | Culicidae | *Culex* | *Culex quinquefasciatus* | GCA_015732765.1 |
| Diptera | Culicidae | *Malaya* | *Malaya genurostris* | GCA_030247185.2 |
| Diptera | Culicidae | *Sabethes* | *Sabethes cyaneus* | GCA_943734655.2 |
| Diptera | Culicidae | *Topomyia* | *Topomyia yanbarensis* | GCA_030247195.1 |
| Diptera | Culicidae | *Toxorhynchites* | *Toxorhynchites rutilus* | GCA_029784135.1 |
| Diptera | Culicidae | *Uranotaenia* | *Uranotaenia lowii* | GCA_029784155.1 |
| Diptera | Culicidae | *Wyeomyia* | *Wyeomyia smithii* | GCA_029784165.1 |
| Diptera | Curtonotidae | *Curtonotum* | *Curtonotum* sp. | SRR10425991 |
| Diptera | Cylindrotomidae | *Liogma* | *Liogma simplicicornis* | SRR3441821 |
| Diptera | Deuterophlebiidae | *Deuterophlebia* | *Deuterophlebia acutirhina* | SRR32672060 |
| Diptera | Deuterophlebiidae | *Deuterophlebia* | *Deuterophlebia coloradensis* | SRR1695339 |
| Diptera | Deuterophlebiidae | *Deuterophlebia* | *Deuterophlebia wuyiense* | This study |
| Diptera | Diadocidiidae | *Diadocidia* | *Diadocidia ferruginosa* | GCA_027564275.1 |
| Diptera | Diastatidae | *Diastata* | *Diastata repleta* | SRR10425990 |
| Diptera | Diopsidae | *Teleopsis* | *Teleopsis dalmanni* | GCA_002237135.5 |
| Diptera | Diopsidae | *Teleopsis* | *Teleopsis pallifacies* | SRR9107624 |
| Diptera | Ditomyiidae | *Symmerus* | *Symmerus nobilis* | GCA_027564815.1 |
| Diptera | Dixidae | *Dixa* | *Dixa* sp. | SRR6155945 |
| Diptera | Dixidae | *Nothodixa* | *Nothodixa* sp. | SRR6155944 |
| Diptera | Dolichopodidae | *Condylostylus* | *Condylostylus patibulatus* | SRR5559329 |
| Diptera | Dolichopodidae | *Heteropsilopus* | *Heteropsilopus ingenuus* | SRR1695350 |
| Diptera | Drosophilidae | *Drosophila* | *Drosophila hydei* | GCA_003285905.2 |
| Diptera | Drosophilidae | *Drosophila* | *Drosophila melanogaster* | GCA_016617805.2 |
| Diptera | Drosophilidae | *Drosophila* | *Drosophila suzukii* | GCA_013340165.1 |
| Diptera | Empididae | *Empis* | *Empis livida* | ERR10378001 |
| Diptera | Empididae | *Hilarini* | *Hilarini* sp. | SRR1695351 |
| Diptera | Ephydridae | *Ephydra* | *Ephydra hians* | SRR1738666 |
| Diptera | Ephydridae | *Hydrellia* | *Hydrellia griseola* | SRR11658059 |
| Diptera | Ephydridae | *Scatella* | *Scatella stagnalis* | SRR11658065 |
| Diptera | Ephydridae | *Scatella* | *Scatella tenuicosta* | SRR1695391 |

| Order | Family | Genus | Species | Accession |
|---|---|---|---|---|
| Diptera | Fanniidae | *Fannia* | *Fannia canicularis* | SRR8236841 |
| Diptera | Fergusoninidae | *Fergusonina* | *Fergusonina omlandi* | SRR1695346 |
| Diptera | Glossinidae | *Glossina* | *Glossina fuscipes* | GCF_014805625.2 |
| Diptera | Helcomyzidae | *Helcomyza* | *Helcomyza mirabilis* | SRR10426008 |
| Diptera | Heleomyzidae | *Tapeigaster* | *Tapeigaster digitata* | SRR1695398 |
| Diptera | Heterocheilidae | *Heterocheila* | *Heterocheila buccata* | SRR10694690 |
| Diptera | Hippoboscidae | *Melophagus* | *Melophagus ovinus* | ERR3046311 |
| Diptera | Hippoboscidae | *Ortholfersia* | *Ortholfersia macleayi* | SRR1695374 |
| Diptera | Hybotidae | *Stilpon* | *Stilpon pauciseta* | SRR2046575 |
| Diptera | Keroplatidae | *Arachnocampa* | *Arachnocampa luminosa* | SRR2241413 |
| Diptera | Keroplatidae | *Macrocera* | *Macrocera vittata* | GCA_024741295.1 |
| Diptera | Keroplatidae | *Orfelia* | *Orfelia fultoni* | SRR10362330 |
| Diptera | Keroplatidae | *Platyura* | *Platyura marginata* | GCA_026745375.1 |
| Diptera | Lauxaniidae | *Sapromyza* | *Sapromyza sciomyzina* | SRR1695389 |
| Diptera | Limoniidae | *Rhipidia* | *Rhipidia sejuga* | SRR3452300 |
| Diptera | Lonchopteridae | *Lonchoptera* | *Lonchoptera bifurcata* | SRR1695357 |
| Diptera | Megamerinidae | *Megamerina* | *Megamerina dolium* | GCA_963854835.1 |
| Diptera | Micropezidae | *Micropeza* | *Micropeza corrigiolata* | SRR1695366 |
| Diptera | Milichiidae | *Paramyia* | *Paramyia nitens* | SRR1695310 |
| Diptera | Muscidae | *Haematobia* | *Haematobia irritans* | SRR9016894 |
| Diptera | Muscidae | *Musca* | *Musca domestica* | GCF_030504385.1 |
| Diptera | Mycetophilidae | *Exechia* | *Exechia fusca* | GCA_024741255.1 |
| Diptera | Mydidae | *Miltinus* | *Miltinus viduatus* | SRR1695367 |
| Diptera | Mydidae | *Mydas* | *Mydas clavatus* | SRR4345448 |
| Diptera | Mystacinobiidae | *Mystacinobia* | *Mystacinobia zelandica* | SRR6724158 |
| Diptera | Mythicomyiidae | *Psiloderoides* | *Psiloderoides* sp. | SRR10753961 |
| Diptera | Nemestrinidae | *Trichophthalma* | *Trichophthalma ricardoae* | SRR1695399 |
| Diptera | Neminidae | *Nemo* | *Nemo dayi* | SRR10426005 |
| Diptera | Neriidae | *Derocephalus* | *Derocephalus angusticollis* | SRR1695338 |
| Diptera | Nycteribiidae | unknown | Nycteribiidae sp. | PRJNA1235147 |
| Diptera | Nymphomyiidae | *Nymphomyia* | *Nymphomyia dolichopeza* | SRR1695309 |
| Diptera | Nymphomyiidae | *Nymphomyia* | *Nymphomyia* sp. | SRR32672058 |
| Diptera | Odiniidae | *Odinia* | *Odinia conspicua* | SRR10426004 |
| Diptera | Oestridae | *Cuterebra* | *Cuterebra austeni* | SRR1695306 |
| Diptera | Opomyzidae | *Opomyza* | *Opomyza germinationis* | SRR10426003 |
| Diptera | Pallopteridae | *Palloptera* | *Palloptera scutellata* | GCA_958295655.1 |
| Diptera | Pallopteridae | *Toxonevra* | *Toxonevra muliebris* | GCA_963691655.1 |
| Diptera | Pantophthalmidae | *Pantophthalmus* | *Pantophthalmus roseni* | SRR2046573 |
| Diptera | Paraleucopidae | *Paraleucopidae* | *Paraleucopidae* sp. | SRR10426002 |
| Diptera | Pediciidae | *Pedicia* | *Pedicia vetusta* | SRR3452301 |
| Diptera | Pelecorhynchidae | *Pelecorhynchus* | *Pelecorhynchus fulvus* | SRR1695377 |
| Diptera | Periscelididae | *Scutops* | *Scutops* sp. | SRR10426001 |
| Diptera | Perissommatidae | *Perissomma* | *Perissomma mcalpinei* | SRR1695312, SRR1695311 |
| Diptera | Phoridae | *Megaselia* | *Megaselia scalaris* | SRR629489 |

| Order | Family | Genus | Species | Accession |
|---|---|---|---|---|
| Diptera | Piophilidae | *Piophila* | *Piophila australis* | SRR1695380 |
| Diptera | Pipunculidae | *Nephrocerus* | *Nephrocerus atrapilus* | SRR1695371 |
| Diptera | Platypezidae | *Platypeza* | *Platypeza anthrax* | SRR1695382 |
| Diptera | Platystomatidae | *Lenophila* | *Lenophila dentipes* | SRR1695353 |
| Diptera | Pleciidae | *Penthetria* | *Penthetria funebris* | SRR6000147 |
| Diptera | Polleniidae | *Pollenia* | *Pollenia angustigena* | ERR10123660 |
| Diptera | Psilidae | *Loxocera* | *Loxocera cylindrica* | SRR1695358 |
| Diptera | Psychodidae | *Clogmia* | *Clogmia albipunctata* | GCA_001014945.1 |
| Diptera | Psychodidae | *Lutzomyia* | *Lutzomyia longipalpis* | GCF_024334085.1 |
| Diptera | Psychodidae | *Phlebotomus* | *Phlebotomus argentipes* | GCA_947086385.1 |
| Diptera | Ptychopteridae | *Bittacomorpha* | *Bittacomorpha clavipes* | SRR1695386 |
| Diptera | Ptychopteridae | *Ptychoptera* | *Ptychoptera albimana* | GCA_961205885.1 |
| Diptera | Pyrgotidae | *Pyrgota* | *Pyrgota undata* | SRR1695387 |
| Diptera | Rhagionidae | *Rhagio* | *Rhagio* sp. | SRR1695388 |
| Diptera | Rhagionidae | *Symphoromyia* | *Symphoromyia* sp. | SRR1695396 |
| Diptera | Sarcophagidae | *Sarcophaga* | *Sarcophaga caerulescens* | GCA_927399465.1 |
| Diptera | Sarcophagidae | *Sarcophaga* | *Sarcophaga rosellei* | GCA_930367235.1 |
| Diptera | Sarcophagidae | *Senotainia* | *Senotainia albifrons* | SRR10753906 |
| Diptera | Sarcophagidae | *Wohlfahrtia* | *Wohlfahrtia magnifica* | GCA_023159175.1 |
| Diptera | Scathophagidae | *Scathophaga* | *Scathophaga stercoraria* | SRR8236843 |
| Diptera | Scatopsidae | *Coboldia* | *Coboldia fuscipes* | GCA_001014335.1 |
| Diptera | Scenopinidae | *Scenopinus* | *Scenopinus velutinus* | SRR2046574 |
| Diptera | Sciaridae | *Bradysia* | *Bradysia coprophila* | GCA_014529535.1 |
| Diptera | Sciaridae | *Phytosciara* | *Phytosciara flavipes* | GCA_026413405.1 |
| Diptera | Sciaridae | *Trichosia* | *Trichosia splendens* | GCA_026413465.1 |
| Diptera | Sciomyzidae | *Limnia* | *Limnia unguicornis* | SRR1695355 |
| Diptera | Sciomyzidae | *Sepedon* | *Sepedon violaceus* | SRR11781470 |
| Diptera | Sepsidae | *Orygma* | *Orygma luctuosum* | SRR10694691 |
| Diptera | Sepsidae | *Themira* | *Themira minor* | SRR1700693 |
| Diptera | Simuliidae | *Paracnephia* | *Paracnephia* sp. | SRR6155943 |
| Diptera | Simuliidae | *Simulium* | *Simulium angustipes* | SRR6808734 |
| Diptera | Sphaeroceridae | *Leptocera* | *Leptocera erythrocera* | SRR2046566 |
| Diptera | Stratiomyidae | *Hermetia* | *Hermetia illucens* | GCA_905115235.1 |
| Diptera | Stratiomyidae | *Odontomyia* | *Odontomyia opertanea* | SRR10766057 |
| Diptera | Strongylophthalmyiidae | *Strongylophthalmyia* | *Strongylophthalmyia* sp. | SRR10426000 |
| Diptera | Syringogastridae | *Syringogaster* | *Syringogaster plesioterga* | SRR10425998 |
| Diptera | Syrphidae | *Eristalis* | *Eristalis pertinax* | SRR1695342 |
| Diptera | Syrphidae | *Merodon* | *Merodon equestris* | SRR1695363 |
| Diptera | Syrphidae | *Merodon* | *Merodon fulcratus* | SRR11658064 |
| Diptera | Syrphidae | *Metasyrphus* | *Metasyrphus corollae* | SRR3645381 |
| Diptera | Syrphidae | *Sphaerophoria* | *Sphaerophoria rueppellii* | ERR7197667 |
| Diptera | Syrphidae | *Syritta* | *Syritta pipiens* | SRR1695397 |
| Diptera | Tabanidae | *Cydistomyia* | *Cydistomyia duplonotata* | SRR11669712 |
| Diptera | Tabanidae | *Scaptia* | *Scaptia patula* | SRR10766064 |
| Diptera | Tachinidae | *Cylindromyia* | *Cylindromyia* sp. | SRR6724166 |

| Order | Family | Genus | Species | Accession |
|---|---|---|---|---|
| Diptera | Tachinidae | *Exorista* | *Exorista civilis* | SRR14252805 |
| Diptera | Tachinidae | *Gymnosoma* | *Gymnosoma nitens* | SRR6724168 |
| Diptera | Tachinidae | *Mintho* | *Mintho rufiventris* | SRR6724169 |
| Diptera | Tanyderidae | *Protoplasa* | *Protoplasa fitchii* | SRR1695383 |
| Diptera | Tephritidae | *Bactrocera* | *Bactrocera tryoni* | GCA_016617805.2 |
| Diptera | Teratomyzidae | *Auster* | *Auster* sp. | SRR10425997 |
| Diptera | Tethinidae | *Triarthria* | *Triarthria setipennis* | SRR921652 |
| Diptera | Thaumaleidae | *Androprosopa* | *Androprosopa vaillantiana* | SRR1695316 |
| Diptera | Thaumaleidae | *Austrothaumalea* | *Austrothaumalea cf* | SRR6155940 |
| Diptera | Therevidae | *Anabarhynchus* | *Anabarhynchus dentiphallus* | SRR1695314 |
| Diptera | Therevidae | *Neodialineura* | *Neodialineura nitens* | SRR1695370 |
| Diptera | Tipulidae | *Nephrotoma* | *Nephrotoma appendiculata* | GCA_947310385.1 |
| Diptera | Tipulidae | *Tipula* | *Tipula unca* | GCA_951394425.1 |
| Diptera | Trichoceridae | *Trichocera* | *Trichocera saltator* | SRR921653 |
| Diptera | Ulidiidae | *Zacompsia* | *Zacompsia fulva* | SRR1695403 |
| Diptera | Ulurumyiidae | *Ulurumyia* | *Ulurumyia macalpinei* | SRR6724160 |
| Diptera | Vermileonidae | *Vermileo* | *Vermileo vermileo* | SRR1695401 |
| Diptera | Xylomyidae | *Xylomya* | *Xylomya americana* | SRR13131721 |
| Diptera | Xylophagidae | *Dialysis* | *Dialysis rufithorax* | SRR1695340 |
| Diptera | Xylophagidae | *Exeretonevra* | *Exeretonevra* sp. | SRR10766065 |
| Diptera | Xylophagidae | *Xylophagus* | *Xylophagus abdominalis* | SRR1695402 |
| Hemiptera | Cicadellidae | *Homalodisca* | *Homalodisca vitripennis* | GCA_021130785.2 |
| Hemiptera | Membracidae | *Entylia* | *Entylia carinata* | SRP152991 |
| Hymenoptera | Apidae | *Apis* | *Apis mellifera* | GCA_003254395.2 |
| Lepidoptera | Bombycidae | *Bombyx* | *Bombyx mori* | GCA_030269925.2 |
| Mecoptera | Panorpidae | *Panorpa* | *Panorpa germanica* | GCA_963678705.1 |
| Siphonaptera | Pulicidae | *Ctenocephalides* | *Ctenocephalides felis* | GCA_003426905.1 |

**Table S2. Species and genomic data used for MCMCTREE of Diptera.**

| Order | Family | Genus | Species | Accession |
|---|---|---|---|---|
| Diptera | Acroceridae | *Acrocera* | *Acrocera orbiculus* | GCA_947359355.1 |
| Diptera | Anisopodidae | *Sylvicola* | *Sylvicola fuscatus* | GCA_026546915.1 |
| Diptera | Anthomyiidae | *Eustalomyia* | *Eustalomyia histrio* | GCA_949748255.1 |
| Diptera | Asilidae | *Philonicus* | *Philonicus albiceps* | GCA_963969385.1 |
| Diptera | Athericidae | *Atherix* | *Atherix ibis* | GCA_958298945.1 |
| Diptera | Axymyiidae | *Axymyia* | *Axymyia furcata* | SRR1695322 |
| Diptera | Bibionidae | *Dilophus* | *Dilophus febrilis* | GCA_958336335.1 |
| Diptera | Blephariceridae | *Philorus* | *Philorus* sp. | This study |
| Diptera | Bombyliidae | *Bombylius* | *Bombylius major* | GCA_932526495.1 |
| Diptera | Calliphoridae | *Lucilia* | *Lucilia cuprina* | GCA_022045245.1 |
| Diptera | Ceratopogonidae | *Culicoides* | *Culicoides sonorensis* | GCA_900258525.3 |
| Diptera | Chaoboridae | *Mochlonyx* | *Mochlonyx cinctipes* | GCA_001014845.1 |
| Diptera | Chironomidae | *Chironomus* | *Chironomus tentans* | GCA_963573255.1 |
| Diptera | Chironomidae | *Polypedilum* | *Polypedilum vanderplanki* | GCA_018290095.1 |
| Diptera | Culicidae | *Aedes* | *Aedes aegypti* | GCA_002204515.1 |
| Diptera | Culicidae | *Aedes* | *Aedes albopictus* | GCA_006496715.1 |
| Diptera | Deuterophlebiidae | *Deuterophlebia* | *Deuterophlebia wuyishanense* | This study |
| Diptera | Diadocidiidae | *Diadocidia* | *Diadocidia ferruginosa* | GCA_027564275.1 |
| Diptera | Dolichopodidae | *Poecilobothrus* | *Poecilobothrus nobilitatus* | GCA_947095535.1 |
| Diptera | Drosophilidae | *Drosophila* | *Drosophila melanogaster* | GCA_016617805.2 |
| Diptera | Empididae | *Empis* | *Empis stercorea* | GCA_949752835.1 |
| Diptera | Hippoboscidae | *Crataerina* | *Crataerina pallida* | GCA_949710015.1 |
| Diptera | Hippoboscidae | *Ornithomya* | *Ornithomya chloropus* | GCA_963971445.1 |
| Diptera | Hybotidae | *Hybos* | *Hybos culiciformis* | GCA_964007475.1 |
| Diptera | Keroplatidae | *Platyura* | *Platyura marginata* | GCA_026745375.1 |
| Diptera | Muscidae | *Musca* | *Musca domestica* | GCF_030504385.1 |
| Diptera | Mycetophilidae | *Bolitophila* | *Bolitophila cinerea* | GCA_010015015.2 |
| Diptera | Nemestrinidae | *Trichophthalma* | *Trichophthalma ricardoae* | SRR1695399 |
| Diptera | Nymphomyiidae | *Nymphomyia* | *Nymphomyia* sp. | SRR32672058 |
| Diptera | Perissommatidae | *Perissomma* | *Perissomma mcalpinei* | SRR1695312, SRR1695311 |
| Diptera | Phoridae | *Megaselia* | *Megaselia scalaris* | SRR629489 |
| Diptera | Pipunculidae | *Nephrocerus* | *Nephrocerus scutellatus* | GCA_947095585.1 |
| Diptera | Psychodidae | *Phlebotomus* | *Phlebotomus argentipes* | GCA_947086385.1 |
| Diptera | Psychodidae | *Lutzomyia* | *Lutzomyia longipalpis* | GCF_024334085.1 |
| Diptera | Ptychopteridae | *Ptychoptera* | *Ptychoptera albimana* | GCA_961205885.1 |
| Diptera | Scatopsidae | *Coboldia* | *Coboldia fuscipes* | GCA_001014335.1 |
| Diptera | Sciaridae | *Bradysia* | *Bradysia coprophila* | GCA_014529535.1 |
| Diptera | Stratiomyidae | *Hermetia* | *Hermetia illucens* | GCA_905115235.1 |
| Diptera | Syrphidae | *Episyrphus* | *Episyrphus balteatus* | GCA_945859705.1 |
| Diptera | Tabanidae | *Chrysops* | *Chrysops viduatus* | GCA_964274935.1 |
| Diptera | Tachinidae | *Nowickia* | *Nowickia ferox* | GCA_936439885.1 |
| Diptera | Tephritidae | *Bactrocera* | *Bactrocera tryoni* | GCA_016617805.2 |
| Diptera | Therevidae | *Thereva* | *Thereva nobilitata* | GCA_963855945.1 |

| Order | Family | Genus | Species | Accession |
|---|---|---|---|---|
| Diptera | Tipulidae | *Tipula* | *Tipula vernalis* | GCA_958295665.1 |
| Diptera | Trichoceridae | *Trichocera* | *Trichocera saltator* | SRR921653 |
| Diptera | Vermileonidae | *Vermileo* | *Vermileo vermileo* | SRR1695401 |
| Diptera | Xylophagidae | *Xylophagus* | *Xylophagus ater* | GCA_963422695.1 |
| Hymenoptera | Apidae | *Apis* | *Apis mellifera* | GCA_003254395.2 |
| Siphonaptera | Pulicidae | *Ctenocephalides* | *Ctenocephalides felis* | GCA_003426905.1 |
| Coleoptera | Tribolium | *castaneum* | *Tribolium castaneum* | GCA_000002335.3 |
| Lepidoptera | Bombycidae | *Bombyx* | *Bombyx mori* | GCA_030269925.2 |
| Mecoptera | Panorpidae | *Panorpa* | *Panorpa germanica* | GCA_963678705.1 |
| Hemiptera | Cicadellidae | *Homalodisca* | *Homalodisca vitripennis* | GCA_021130785.2 |

**Table S3. Statistics for genome assemblies of *De. wuyiensis* and *Philorus* sp.**

| **Assembly** | ***De. wuyiensis*** | ***Philorus* sp.** |
|---|---|---|
| # contigs (>= 10000 bp) | 37 | 76 |
| # contigs (>= 25000 bp) | 35 | 74 |
| # contigs (>= 50000 bp) | 23 | 39 |
| Total length (>= 10000 bp) | 125,473,247 | 128,985,792 |
| Total length (>= 25000 bp) | 125,429,877 | 128,941,792 |
| Total length (>= 50000 bp) | 125,019,515 | 127,809,012 |
| # contigs | 37 | 76 |
| Largest contig | 33,964,599 | 40,043,424 |
| Total length | 125,473,247 | 128,985,792 |
| GC (%) | 25 | 38 |
| Reference GC (%) | 42 | 42 |
| N50 | 30,030,989 | 29,067,738 |
| NG50 | 24,404,533 | 28,021,126 |
| N90 | 4,068,494 | 23,436,045 |
| NG90 | - | - |
| auN | 25,883,314 | 29,394,931 |
| auNG | 22,596,214 | 26,380,254 |
| L50 | 2 | 2 |
| LG50 | 3 | 3 |
| L90 | 5 | 4 |
| LG90 | - | - |
| **Third-generation sequencing** | | |
| # total reads | 1,676,880 | 2,891,147 |
| Avg. coverage depth | 233 | 492 |
| **Second-generation sequencing** | | |
| # total reads | 378,826,648 | 205,221,154 |
| Properly paired (%) | 95 | 96 |
| Avg. coverage depth | 434 | 223 |

**Table S4. Morphological and ecological characteristics of all documented Diptera families in relation to the presence of prothoracic pupal respiratory organ (PROD).**

| Label | Family | PROD | Tracheal-PROD | Habitat |
|---|---|---|---|---|
| Coleoptera_Scarabaeidae_Onthophagus_taurus_GCA_000648695.2 | Scarabaeidae | absent | absent | Terrestrial |
| Coleoptera_Scarabaeidae_Trypoxylus_dichotomus_GCA_023509865.1 | Scarabaeidae | absent | absent | Terrestrial |
| Coleoptera_Tribolium_castaneum_GCA_000002335.3 | Tribolium | absent | absent | Terrestrial |
| Hemiptera_Cicadellidae_Homalodisca_vitripennis_GCA_021130785.2 | Cicadellidae | absent | absent | Terrestrial |
| Hemiptera_Membracidae_Entylia_carinata_SRP152991 | Membracidae | absent | absent | Terrestrial |
| Hymenoptera_Apidae_Apis_mellifera_GCA_003254395.2 | Apidae | absent | absent | Terrestrial |
| Lepidoptera_Bombycidae_Bombyx_mori_GCA_030269925.2 | Bombycidae | absent | absent | Terrestrial |
| Mecoptera_Panorpidae_Panorpa_germanica_GCA_963678705.1 | Panorpidae | absent | absent | Terrestrial |
| Siphonaptera_Pulicidae_Ctenocephalides_felis_GCA_003426905.1 | Pulicidae | absent | absent | Terrestrial |
| Diptera_Deuterophlebiidae_Deuterophlebia_acutirhina_3 | Deuterophlebiidae | present | absent | Aquatic/Semi-aquatic |
| Diptera_Deuterophlebiidae_Deuterophlebia_coloradensis_SRR1695339 | Deuterophlebiidae | present | absent | Aquatic/Semi-aquatic |
| Diptera_Deuterophlebiidae_Deuterophlebia_wuyishanense_genome | Deuterophlebiidae | present | absent | Aquatic/Semi-aquatic |
| Diptera_Nymphomyiidae_Nymphomyia_dolichopeza_SRR1695309 | Nymphomyiidae | absent | absent | Aquatic/Semi-aquatic |
| Diptera_Nymphomyiidae_Nymphomyia_sp | Nymphomyiidae | absent | absent | Aquatic/Semi-aquatic |
| Diptera_Trichoceridae_Trichocera_saltator_SRR921653 | Trichoceridae | present | absent | Aquatic/Semi-aquatic |
| Diptera_Limoniidae_Rhipidia_sejuga_SRR3452300 | Limoniidae | present | absent | Aquatic/Semi-aquatic |
| Diptera_Tipulidae_Nephrotoma_appendiculata_GCA.947310385.1 | Tipulidae | present | absent | Aquatic/Semi-aquatic |
| Diptera_Tipulidae_Tipula_unca_GCA.951394425.1 | Tipulidae | present | absent | Aquatic/Semi-aquatic |
| Diptera_Pediciidae_Pedicia_vetusta_SRR3452301 | Pediciidae | present | absent | Aquatic/Semi-aquatic |

| Label | Family | PROD | Tracheal-PROD | Habitat |
| --- | --- | --- | --- | --- |
| Diptera_Cylindrotomidae_Liogma_simplicicornis_SRR3441821 | Cylindrotomidae | present | absent | Aquatic/Semi-aquatic |
| Diptera_Ptychopteridae_Bittacomorpha_clavipes_SRR1695386 | Ptychopteridae | present | absent | Aquatic/Semi-aquatic |
| Diptera_Ptychopteridae_Ptychoptera_albimana_GCA.961205885.1 | Ptychopteridae | present | absent | Aquatic/Semi-aquatic |
| Diptera_Ceratopogonidae_Atrichopogon_sp_SRR6155951 | Ceratopogonidae | present | absent | Aquatic/Semi-aquatic |
| Diptera_Ceratopogonidae_Culicoides_sonorensis_SRR1174038 | Ceratopogonidae | present | absent | Aquatic/Semi-aquatic |
| Diptera_Ceratopogonidae_Dasyhelea_sp_SRR6155934 | Ceratopogonidae | present | absent | Aquatic/Semi-aquatic |
| Diptera_Ceratopogonidae_Forcipomyia_taiwana.merge | Ceratopogonidae | present | absent | Aquatic/Semi-aquatic |
| Diptera_Chaoboridae_Chaoborus_americanus.merge | Chaoboridae | present | absent | Aquatic/Semi-aquatic |
| Diptera_Chaoboridae_Chaoborus_flavidulus.merge | Chaoboridae | present | absent | Aquatic/Semi-aquatic |
| Diptera_Chaoboridae_Mochlonyx_cinctipes_GCA.001014845.1 | Chaoboridae | present | absent | Aquatic/Semi-aquatic |
| Diptera_Chironomidae_Cardiocladius_sp_SRR6155936 | Chironomidae | present | absent | Aquatic/Semi-aquatic |
| Diptera_Chironomidae_Chironomus_tentans_GCA.963573255.1 | Chironomidae | present | absent | Aquatic/Semi-aquatic |
| Diptera_Chironomidae_Clunio_marinus_GCA.900005825.1 | Chironomidae | present | absent | Aquatic/Semi-aquatic |
| Diptera_Chironomidae_Cricotopus_draysoni_SRR4457915 | Chironomidae | present | absent | Aquatic/Semi-aquatic |
| Diptera_Chironomidae_Diamesa_zernyi_SRR13641921 | Chironomidae | present | absent | Aquatic/Semi-aquatic |
| Diptera_Chironomidae_Kiefferophyes_invenustulus_SRR6155941 | Chironomidae | present | absent | Aquatic/Semi-aquatic |
| Diptera_Chironomidae_Paraheptagyia_tonnoiri_SRR6155948 | Chironomidae | present | absent | Aquatic/Semi-aquatic |
| Diptera_Chironomidae_Paraphaenocladius_impensus_SRR11637888 | Chironomidae | present | absent | Aquatic/Semi-aquatic |

| Label | Family | PROD | Tracheal-PROD | Habitat |
|---|---|---|---|---|
| Diptera_Chironomidae_Parochlus_steinenii.merge | Chironomidae | present | absent | Aquatic/Semi-aquatic |
| Diptera_Chironomidae_Pentaneurella_katterjokki_SRR11637887 | Chironomidae | present | absent | Aquatic/Semi-aquatic |
| Diptera_Chironomidae_Podonomus_sp_SRR6155949 | Chironomidae | present | absent | Aquatic/Semi-aquatic |
| Diptera_Chironomidae_Polypedilum_pembai_GCA.014622435.1 | Chironomidae | present | absent | Aquatic/Semi-aquatic |
| Diptera_Chironomidae_Polypedilum_vanderplanki_GCA.018290095.1 | Chironomidae | present | absent | Aquatic/Semi-aquatic |
| Diptera_Chironomidae_Procladius_villosimanus_SRR6155953 | Chironomidae | present | absent | Aquatic/Semi-aquatic |
| Diptera_Chironomidae_Propsilocerus_akamusi_GCA.018397935.1 | Chironomidae | present | absent | Aquatic/Semi-aquatic |
| Diptera_Chironomidae_Smittia_aterrima_GCA.033063855.1 | Chironomidae | present | absent | Aquatic/Semi-aquatic |
| Diptera_Chironomidae_Telmatogeton_pectinata_SRR6155952 | Chironomidae | present | absent | Aquatic/Semi-aquatic |
| Diptera_Chironomidae_Telmatopelopia_nemorum_SRR6155950 | Chironomidae | present | absent | Aquatic/Semi-aquatic |
| Diptera_Culicidae_Aedes_aegypti_GCA.002204515.1 | Culicidae | present | absent | Aquatic/Semi-aquatic |
| Diptera_Culicidae_Aedes_albopictus_GCA.006496715.1 | Culicidae | present | absent | Aquatic/Semi-aquatic |
| Diptera_Culicidae_Anopheles_cracens_GCA.002091845.1 | Culicidae | present | absent | Aquatic/Semi-aquatic |
| Diptera_Culicidae_Anopheles_gambiae_GCA.943734735.2 | Culicidae | present | absent | Aquatic/Semi-aquatic |
| Diptera_Culicidae_Anopheles_sinensis_GCA.000441895.2 | Culicidae | present | absent | Aquatic/Semi-aquatic |
| Diptera_Culicidae_Anopheles_stephensi_GCA.013141755.1 | Culicidae | present | absent | Aquatic/Semi-aquatic |
| Diptera_Culicidae_Armigeres_subalbatus_GCA.024139115.1 | Culicidae | present | absent | Aquatic/Semi-aquatic |
| Diptera_Culicidae_Culex_quinquefasciatus_GCA.015732765.1 | Culicidae | present | absent | Aquatic/Semi-aquatic |

| Label | Family | PROD | Tracheal-PROD | Habitat |
|---|---|---|---|---|
| Diptera_Culicidae_Malaya_genurostris_GCA.030247185.2 | Culicidae | present | absent | Aquatic/Semi-aquatic |
| Diptera_Culicidae_Sabethes_cyaneus_GCA.943734655.2 | Culicidae | present | absent | Aquatic/Semi-aquatic |
| Diptera_Culicidae_Topomyia_yanbarensis_GCA.030247195.1 | Culicidae | present | absent | Aquatic/Semi-aquatic |
| Diptera_Culicidae_Toxorhynchites_rutilus_septentrionalis_GCA.029784135.1 | Culicidae | present | absent | Aquatic/Semi-aquatic |
| Diptera_Culicidae_Uranotaenia_lowii_GCA.029784155.1 | Culicidae | present | absent | Aquatic/Semi-aquatic |
| Diptera_Culicidae_Wyeomyia_smithii__GCA.029784165.1 | Culicidae | present | absent | Aquatic/Semi-aquatic |
| Diptera_Dixidae_Dixa_sp_SRR6155945 | Dixidae | present | absent | Aquatic/Semi-aquatic |
| Diptera_Dixidae_Nothodixa_sp_SRR6155944 | Dixidae | present | absent | Aquatic/Semi-aquatic |
| Diptera_Simuliidae_Paracnephia_sp_SRR6155943 | Simuliidae | present | absent | Aquatic/Semi-aquatic |
| Diptera_Simuliidae_Simulium_angustipes_SRR6808734 | Simuliidae | present | absent | Aquatic/Semi-aquatic |
| Diptera_Thaumaleidae_Androprosopa_vaillantiana_SRR1695316 | Thaumaleidae | present | absent | Aquatic/Semi-aquatic |
| Diptera_Thaumaleidae_Austrothaumalea_cf_denticulata_SRR6155940 | Thaumaleidae | present | absent | Aquatic/Semi-aquatic |
| Diptera_Corethrellidae_Corethrella_appendiculata.merge | Corethrellidae | present | absent | Aquatic/Semi-aquatic |
| Diptera_Corethrellidae_Corethrella_calathicola_SRR6155947 | Corethrellidae | present | absent | Aquatic/Semi-aquatic |
| Diptera_Blephariceridae_Blepharicera_sp_SRR1695325 | Blephariceridae | present | absent | Aquatic/Semi-aquatic |
| Diptera_Blephariceridae_sp_yellow | Blephariceridae | present | absent | Aquatic/Semi-aquatic |
| Diptera_Blephariceridae_sp_Red | Blephariceridae | present | absent | Aquatic/Semi-aquatic |

| Label | Family | PROD | Tracheal-PROD | Habitat |
|---|---|---|---|---|
| Diptera_Psychodidae_Clogmia_albipunctata_GCA.001014945.1 | Psychodidae | present | absent | Aquatic/Semi-aquatic |
| Diptera_Psychodidae_Lutzomyia_longipalpis_GCF.024334085.1 | Psychodidae | present | absent | Aquatic/Semi-aquatic |
| Diptera_Psychodidae_Phlebotomus_argentipes_GCA.947086385.1 | Psychodidae | present | absent | Aquatic/Semi-aquatic |
| Diptera_Tanyderidae_Protoplasa_fitchii_SRR1695383 | Tanyderidae | present | absent | Aquatic/Semi-aquatic |
| Diptera_Perissommatidae_Perissomma_mcalpinei_merged | Perissommatidae | present | absent | Terrestrial |
| Diptera_Axymyiidae_Axymyia_furcata_SRR1695322 | Axymyiidae | present | absent | Aquatic/Semi-aquatic |
| Diptera_Anisopodidae_Sylvicola_fuscatus_GCA.026546915.1 | Anisopodidae | present | present | Aquatic/Semi-aquatic |
| Diptera_Bibionidae_Bibio_marci_GCA.910594885.2 | Bibionidae | absent | absent | Aquatic/Semi-aquatic |
| Diptera_Bibionidae_Dilophus_febrilis_GCA.958336335.1 | Bibionidae | absent | absent | Aquatic/Semi-aquatic |
| Diptera_Cecidomyiidae_Aphidoletes_aphidimyza_GCA.030463065.1 | Cecidomyiidae | present | absent | Terrestrial |
| Diptera_Cecidomyiidae_Catotricha_subobsoleta_GCA.011634745.2 | Cecidomyiidae | present | absent | Terrestrial |
| Diptera_Cecidomyiidae_Contarinia_nasturtii_GCA.009176525.2 | Cecidomyiidae | present | absent | Terrestrial |
| Diptera_Cecidomyiidae_Lestremia_cinerea_GCA.027564135.1 | Cecidomyiidae | present | absent | Terrestrial |
| Diptera_Cecidomyiidae_Mayetiola_destructor_GCA.000149185.1 | Cecidomyiidae | present | absent | Terrestrial |
| Diptera_Cecidomyiidae_Obolodiplosis_robiniae_GCA.028476595.1 | Cecidomyiidae | present | absent | Terrestrial |
| Diptera_Cecidomyiidae_Resseliella_maxima_GCA.029041755.1 | Cecidomyiidae | present | absent | Terrestrial |
| Diptera_Cecidomyiidae_Sitodiplosis_mosellana_GCA.009176505.1 | Cecidomyiidae | present | absent | Terrestrial |
| Diptera_Mycetophilidae_Exechia_fusca_GCA.024741255.1 | Mycetophilidae | present | absent | Aquatic/Semi-aquatic |
| Diptera_Scatopsidae_Coboldia_fuscipes_GCA.001014335.1 | Scatopsidae | present | absent | Aquatic/Semi-aquatic |
| Diptera_Sciaridae_Bradysia_coprophila_GCA.014529535.1 | Sciaridae | present | present | Aquatic/Semi-aquatic |

| Label | Family | PROD | Tracheal-PROD | Habitat |
|---|---|---|---|---|
| Diptera_Sciaridae_Phytosciara_flavipes_GCA.026413405.1 | Sciaridae | present | present | Aquatic/Semi-aquatic |
| Diptera_Sciaridae_Trichosia_splendens_GCA.026413465.1 | Sciaridae | present | present | Aquatic/Semi-aquatic |
| Diptera_Keroplatidae_Arachnocampa_luminosa_SRR2241413 | Keroplatidae | absent | absent | Terrestrial |
| Diptera_Keroplatidae_Macrocera_vittata_GCA.024741295.1 | Keroplatidae | absent | absent | Terrestrial |
| Diptera_Keroplatidae_Orfelia_fultoni_SRR10362330 | Keroplatidae | absent | absent | Terrestrial |
| Diptera_Keroplatidae_Platyura_marginata_GCA.026745375.1 | Keroplatidae | absent | absent | Terrestrial |
| Diptera_Bolitophilidae_Bolitophila_cinerea_GCA.010015015.2 | Bolitophilidae | present | absent | Terrestrial |
| Diptera_Diadocidiidae_Diadocidia_ferruginosa_GCA.027564275.1 | Diadocidiidae | unknown | unknown | Terrestrial |
| Diptera_Ditomyiidae_Symmerus_nobilis_GCA.027564815.1 | Ditomyiidae | unknown | unknown | Unknown |
| Diptera_Pleciidae_Penthetria_funebris_SRR6000147 | Pleciidae | unknown | unknown | Unknown |
| Diptera_Nemestrinidae_Trichophthalma_ricardoae_SRR1695399 | Nemestrinidae | absent | absent | Terrestrial |
| Diptera_Stratiomyidae_Hermetia_illucens_GCA_905115235.1 | Stratiomyidae | present | absent | Aquatic/Semi-aquatic |
| Diptera_Stratiomyidae_Odontomyia_opertanea_SRR10766057 | Stratiomyidae | present | absent | Aquatic/Semi-aquatic |
| Diptera_Xylomyidae_Xylomya_americana_SRR13131721 | Xylomyidae | absent | absent | Terrestrial |
| Diptera_Pantophthalmidae_Pantophthalmus_roseni_SRR2046573 | Pantophthalmidae | absent | absent | Terrestrial |
| Diptera_Athericidae_Atherix_ibis_GCA.958298945.1 | Athericidae | present | absent | Aquatic/Semi-aquatic |
| Diptera_Pelecorhynchidae_Pelecorhynchus_fulvus_SRR1695377 | Pelecorhynchidae | absent | absent | Aquatic/Semi-aquatic |
| Diptera_Rhagionidae_Rhagio_sp_SRR1695388 | Rhagionidae | absent | absent | Aquatic/Semi-aquatic |
| Diptera_Tabanidae_Scaptia_patula_SRR10766064 | Tabanidae | present | absent | Aquatic/Semi-aquatic |
| Diptera_Vermileonidae_Vermileo_vermileo_SRR1695401 | Vermileonidae | present | present | Terrestrial |
| Diptera_Xylophagidae_Dialysis_rufithorax_SRR1695340 | Xylophagidae | present | absent | Terrestrial |

| Label | Family | PROD | Tracheal-PROD | Habitat |
|---|---|---|---|---|
| Diptera_Xylophagidae_Exeretonevra_sp._ANIC_BT03_SRR10766065 | Xylophagidae | present | absent | Terrestrial |
| Diptera_Xylophagidae_Xylophagus_abdominalis_SRR1695402 | Xylophagidae | present | absent | Terrestrial |
| Diptera_Drosophilidae_Drosophila_hydei_GCA_003285905.2 | Drosophilidae | present | present | Aquatic/Semi-aquatic |
| Diptera_Drosophilidae_Drosophila_melanogaster_GCA_016617805.2 | Drosophilidae | present | present | Aquatic/Semi-aquatic |
| Diptera_Drosophilidae_Drosophila_suzukii_GCA_013340165.1 | Drosophilidae | present | present | Aquatic/Semi-aquatic |
| Diptera_Empididae_Empis_livida_ERR10378001 | Empididae | present | absent | Aquatic/Semi-aquatic |
| Diptera_Empididae_Hilarini_sp_SRR1695351 | Empididae | present | absent | Aquatic/Semi-aquatic |
| Diptera_Muscidae_Haematobia_irritans_SRR9016894 | Muscidae | present | present | Aquatic/Semi-aquatic |
| Diptera_Muscidae_Musca_domestica_GCF_030504385.1 | Muscidae | present | absent | Aquatic/Semi-aquatic |
| Diptera_Phoridae_Megaselia_scalaris_SRR629489 | Phoridae | absent | absent | Aquatic/Semi-aquatic |
| Diptera_Sciomyzidae_Limnia_unguicornis_SRR1695355 | Sciomyzidae | present | present | Aquatic/Semi-aquatic |
| Diptera_Sciomyzidae_Sepedon_violaceus_SRR11781470 | Sciomyzidae | present | absent | Aquatic/Semi-aquatic |
| Diptera_Syrphidae_Eristalis_pertinax_SRR1695342 | Syrphidae | present | present | Terrestrial |
| Diptera_Syrphidae_Sphaerophoria_rueppellii_ERR7197667 | Syrphidae | absent | absent | Terrestrial |
| Diptera_Tephritidae_Bactrocera_tryoni_GCA_016617805.2 | Tephritidae | present | present | Terrestrial |
| Diptera_Acroceridae_Pterodontia_mellii_SRR1695385 | Acroceridae | present | absent | Terrestrial |
| Diptera_Agromyzidae_Liriomyza_trifolii_SRR1138236 | Agromyzidae | present | present | Terrestrial |
| Diptera_Anthomyzidae_Mumetopia_occipitalis_SRR1695368 | Anthomyzidae | present | present | Terrestrial |
| Diptera_Apioceridae_Apiocera_maritima_SRR2046562 | Apioceridae | present | absent | Terrestrial |
| Diptera_Apioceridae_Apiocera_moerens_SRR1695318 | Apioceridae | present | absent | Terrestrial |
| Diptera_Asilidae_Diogmites_neoternatus_SRR4345333 | Asilidae | present | absent | Terrestrial |

| Label | Family | PROD | Tracheal-PROD | Habitat |
| --- | --- | --- | --- | --- |
| Diptera_Asilidae_Eudioctria_media_SRR10386632 | Asilidae | present | absent | Terrestrial |
| Diptera_Asilidae_Philonicus_albiceps_SRR4365562 | Asilidae | present | absent | Terrestrial |

**Table S5. Summary of bulk RNAseq datasets used in this study.**

| Sample_id | Species | Data Source | Background | Stage | Tissue | RNA Library |
|---|---|---|---|---|---|---|
| G_84h_R1 | *Drosophila melanogaster* | this study | Canton Special wildtype | 84 hrs AEL | t1 drosal disc | SMARTseq |
| G_84h_R2 | *Drosophila melanogaster* | this study | Canton Special wildtype | 84 hrs AEL | t1 drosal disc | SMARTseq |
| G_84h_R3 | *Drosophila melanogaster* | this study | Canton Special wildtype | 84 hrs AEL | t1 drosal disc | SMARTseq |
| G_96h_R1 | *Drosophila melanogaster* | this study | Canton Special wildtype | 96 hrs AEL | t1 drosal disc | SMARTseq |
| G_96h_R2 | *Drosophila melanogaster* | this study | Canton Special wildtype | 96 hrs AEL | t1 drosal disc | SMARTseq |
| G_96h_R3 | *Drosophila melanogaster* | this study | Canton Special wildtype | 96 hrs AEL | t1 drosal disc | SMARTseq |
| G_108h_R1 | *Drosophila melanogaster* | this study | Canton Special wildtype | 108 hrs AEL | t1 drosal disc | SMARTseq |
| G_108h_R2 | *Drosophila melanogaster* | this study | Canton Special wildtype | 108 hrs AEL | t1 drosal disc | SMARTseq |
| G_108h_R3 | *Drosophila melanogaster* | this study | Canton Special wildtype | 108 hrs AEL | t1 drosal disc | SMARTseq |
| G_120h_R1 | *Drosophila melanogaster* | this study | Canton Special wildtype | 120 hrs AEL | t1 drosal disc | SMARTseq |
| G_120h_R2 | *Drosophila melanogaster* | this study | Canton Special wildtype | 120 hrs AEL | t1 drosal disc | SMARTseq |
| G_120h_R3 | *Drosophila melanogaster* | this study | Canton Special wildtype | 120 hrs AEL | t1 drosal disc | SMARTseq |
| W_84h_R1 | *Drosophila melanogaster* | this study | Canton Special wildtype | 84 hrs AEL | t2 drosal disc | SMARTseq |
| W_84h_R2 | *Drosophila melanogaster* | this study | Canton Special wildtype | 84 hrs AEL | t2 drosal disc | SMARTseq |
| W_84h_R3 | *Drosophila melanogaster* | this study | Canton Special wildtype | 84 hrs AEL | t2 drosal disc | SMARTseq |
| W_96h_R1 | *Drosophila melanogaster* | this study | Canton Special wildtype | 96 hrs AEL | t2 drosal disc | SMARTseq |
| W_96h_R2 | *Drosophila melanogaster* | this study | Canton Special wildtype | 96 hrs AEL | t2 drosal disc | SMARTseq |
| W_96h_R3 | *Drosophila melanogaster* | this study | Canton Special wildtype | 96 hrs AEL | t2 drosal disc | SMARTseq |

| Sample_id | Species | Data Source | Background | Stage | Tissue | RNA Library |
|---|---|---|---|---|---|---|
| W_108h_R1 | *Drosophila melanogaster* | this study | Canton Special wildtype | 108 hrs AEL | t2 drosal disc | SMARTseq |
| W_108h_R2 | *Drosophila melanogaster* | this study | Canton Special wildtype | 108 hrs AEL | t2 drosal disc | SMARTseq |
| W_108h_R3 | *Drosophila melanogaster* | this study | Canton Special wildtype | 108 hrs AEL | t2 drosal disc | SMARTseq |
| W_120h_R1 | *Drosophila melanogaster* | this study | Canton Special wildtype | 120 hrs AEL | t2 drosal disc | SMARTseq |
| W_120h_R2 | *Drosophila melanogaster* | this study | Canton Special wildtype | 120 hrs AEL | t2 drosal disc | SMARTseq |
| W_120h_R3 | *Drosophila melanogaster* | this study | Canton Special wildtype | 120 hrs AEL | t2 drosal disc | SMARTseq |
| W_120h_R4 | *Drosophila melanogaster* | this study | Canton Special wildtype | 120 hrs AEL | t2 drosal disc | SMARTseq |
| Dmel_L2_1_SRR24633458 | *Drosophila melanogaster* | SRR24633458 | unspecified wildtype | 72 hrs AEL eqv | wholebody | RNAseq |
| Dmel_L2_2_SRR24633457 | *Drosophila melanogaster* | SRR24633457 | unspecified wildtype | 72 hrs AEL eqv | wholebody | RNAseq |
| Dmel_L2_3_SRR24633456 | *Drosophila melanogaster* | SRR24633456 | unspecified wildtype | 72 hrs AEL eqv | wholebody | RNAseq |
| Dmel_WL_1_SRR24633454 | *Drosophila melanogaster* | SRR24633458 | unspecified wildtype | 120 hrs AEL eqv | wholebody | RNAseq |
| Dmel_WL_2_SRR24633453 | *Drosophila melanogaster* | SRR24633458 | unspecified wildtype | 120 hrs AEL eqv | wholebody | RNAseq |
| Dmel_WL_3_SRR24633452 | *Drosophila melanogaster* | SRR24633458 | unspecified wildtype | 120 hrs AEL eqv | wholebody | RNAseq |
| Aalbo_CK_SRR8835872 | *Aedes Albopictus* | SRR8835872 | unspecified wildtype | 3 instar larva | wholebody | RNAseq |
| Aalbo_CK_SRR8835873 | *Aedes Albopictus* | SRR8835873 | unspecified wildtype | 3 instar larva | wholebody | RNAseq |
| Aalbo_CK_SRR8835875 | *Aedes Albopictus* | SRR8835875 | unspecified wildtype | 3 instar larva | wholebody | RNAseq |
| Aalbo_HA_SRR8835870 | *Aedes Albopictus* | SRR8835870 | unspecified wildtype | 3 instar larva | wholebody | RNAseq |
| Aalbo_HA_SRR8835871 | *Aedes Albopictus* | SRR8835871 | unspecified wildtype | 3 instar larva | wholebody | RNAseq |
| Aalbo_HA_SRR8835874 | *Aedes Albopictus* | SRR8835874 | unspecified wildtype | 3 instar larva | wholebody | RNAseq |
| Aalbo_T1_Early_1 | *Aedes Albopictus* | this study | GMU iso3-3-10 | 3 instar larva | t1 drosal tissue | RNAseq |
| Aalbo_T1_Early_2 | *Aedes Albopictus* | this study | GMU iso3-3-10 | 3 instar larva | t1 drosal tissue | RNAseq |
| Aalbo_T1_Early_3 | *Aedes Albopictus* | this study | GMU iso3-3-10 | 3 instar larva | t1 drosal tissue | RNAseq |

| Sample_id | Species | Data Source | Background | Stage | Tissue | RNA Library |
|---|---|---|---|---|---|---|
| Aalbo_T1_Late_1 | *Aedes Albopictus* | this study | GMU iso3-3-10 | 4 instar larva | t1 drosal tissue | RNAseq |
| Aalbo_T1_Late_2 | *Aedes Albopictus* | this study | GMU iso3-3-10 | 4 instar larva | t1 drosal tissue | RNAseq |
| Aalbo_T1_Late_3 | *Aedes Albopictus* | this study | GMU iso3-3-10 | 4 instar larva | t1 drosal tissue | RNAseq |
| Aalbo_T2_Early_1 | *Aedes Albopictus* | this study | GMU iso3-3-10 | 3 instar larva | t2 drosal tissue | RNAseq |
| Aalbo_T2_Early_2 | *Aedes Albopictus* | this study | GMU iso3-3-10 | 3 instar larva | t2 drosal tissue | RNAseq |
| Aalbo_T2_Early_3 | *Aedes Albopictus* | this study | GMU iso3-3-10 | 3 instar larva | t2 drosal tissue | RNAseq |
| Aalbo_T2_Late_1 | *Aedes Albopictus* | this study | GMU iso3-3-10 | 4 instar larva | t2 drosal tissue | RNAseq |
| Aalbo_T2_Late_2 | *Aedes Albopictus* | this study | GMU iso3-3-10 | 4 instar larva | t2 drosal tissue | RNAseq |
| Aalbo_T2_Late_3 | *Aedes Albopictus* | this study | GMU iso3-3-10 | 4 instar larva | t2 drosal tissue | RNAseq |
| Early_g_1_3 | *Deuterophlabia wuyiensis* | this study | wuyishan wildtype | early stage of final instar larva | t1 drosal tissue | RNAseq |
| Early_g_2_2 | *Deuterophlabia wuyiensis* | this study | wuyishan wildtype | early stage of final instar larva | t1 drosal tissue | RNAseq |
| Early_g_3_5 | *Deuterophlabia wuyiensis* | this study | wuyishan wildtype | early stage of final instar larva | t1 drosal tissue | RNAseq |
| Early_w_1_3 | *Deuterophlabia wuyiensis* | this study | wuyishan wildtype | early stage of final instar larva | t2 drosal tissue | RNAseq |
| Early_w_2_1 | *Deuterophlabia wuyiensis* | this study | wuyishan wildtype | early stage of final instar larva | t2 drosal tissue | RNAseq |
| Early_w_3_5 | *Deuterophlabia wuyiensis* | this study | wuyishan wildtype | early stage of final instar larva | t2 drosal tissue | RNAseq |
| Late_g_1_2 | *Deuterophlabia wuyiensis* | this study | wuyishan wildtype | final instar larva | t1 drosal tissue | RNAseq |
| Late_g_2_3 | *Deuterophlabia wuyiensis* | this study | wuyishan wildtype | final instar larva | t1 drosal tissue | RNAseq |
| Late_g_3_6 | *Deuterophlabia wuyiensis* | this study | wuyishan wildtype | final instar larva | t1 drosal tissue | RNAseq |
| Late_w_1_2 | *Deuterophlabia wuyiensis* | this study | wuyishan wildtype | final instar larva | t2 drosal tissue | RNAseq |
| Late_w_2_3 | *Deuterophlabia wuyiensis* | this study | wuyishan wildtype | final instar larva | t2 drosal tissue | RNAseq |
| Late_w_3_6 | *Deuterophlabia wuyiensis* | this study | wuyishan wildtype | final instar larva | t2 drosal tissue | RNAseq |
| Wholebody1 | *Deuterophlabia wuyiensis* | this study | wuyishan wildtype | final instar larva | wholebody | RNAseq |

| Sample_id | Species | Data Source | Background | Stage | Tissue | RNA Library |
|---|---|---|---|---|---|---|
| Wholebody3 | *Deuterophlabia wuyiensis* | this study | wuyishan wildtype | final instar larva | wholebody | RNAseq |
| WholebodyLarva5 | *Deuterophlabia wuyiensis* | this study | wuyishan wildtype | final instar larva | wholebody | RNAseq |
| OtauM_T1NepA_SRR10121220 | *Onthophagus taurus* | SRR10121220 | wildtype | white pupa | t1 drosal imaginal tissue | RNAseq |
| OtauM_T1NepA_SRR10121230 | *Onthophagus taurus* | SRR10121230 | wildtype | white pupa | t1 drosal imaginal tissue | RNAseq |
| OtauM_T1NepA_SRR10121240 | *Onthophagus taurus* | SRR10121240 | wildtype | white pupa | t1 drosal imaginal tissue | RNAseq |
| OtauM_T1NepA_SRR10121250 | *Onthophagus taurus* | SRR10121250 | wildtype | white pupa | t1 drosal imaginal tissue | RNAseq |
| OtauM_T1NepA_SRR10121260 | *Onthophagus taurus* | SRR10121260 | wildtype | white pupa | t1 drosal imaginal tissue | RNAseq |
| OtauM_T1NepA_SRR10121270 | *Onthophagus taurus* | SRR10121270 | wildtype | white pupa | t1 drosal imaginal tissue | RNAseq |
| OtauM_T1NepD_SRR10121223 | *Onthophagus taurus* | SRR10121223 | wildtype | white pupa | t1 drosal imaginal tissue | RNAseq |
| OtauM_T1NepD_SRR10121233 | *Onthophagus taurus* | SRR10121233 | wildtype | white pupa | t1 drosal imaginal tissue | RNAseq |
| OtauM_T1NepD_SRR10121243 | *Onthophagus taurus* | SRR10121243 | wildtype | white pupa | t1 drosal imaginal tissue | RNAseq |
| OtauM_T1NepD_SRR10121253 | *Onthophagus taurus* | SRR10121253 | wildtype | white pupa | t1 drosal imaginal tissue | RNAseq |
| OtauM_T1NepD_SRR10121263 | *Onthophagus taurus* | SRR10121263 | wildtype | white pupa | t1 drosal imaginal tissue | RNAseq |
| OtauM_T1NepD_SRR10121273 | *Onthophagus taurus* | SRR10121273 | wildtype | white pupa | t1 drosal imaginal tissue | RNAseq |
| OtauM_T1NepL_SRR10121221 | *Onthophagus taurus* | SRR10121221 | wildtype | white pupa | t1 drosal imaginal tissue | RNAseq |
| OtauM_T1NepL_SRR10121231 | *Onthophagus taurus* | SRR10121231 | wildtype | white pupa | t1 drosal imaginal tissue | RNAseq |
| OtauM_T1NepL_SRR10121241 | *Onthophagus taurus* | SRR10121241 | wildtype | white pupa | t1 drosal imaginal tissue | RNAseq |
| OtauM_T1NepL_SRR10121251 | *Onthophagus taurus* | SRR10121251 | wildtype | white pupa | t1 drosal imaginal tissue | RNAseq |
| OtauM_T1NepL_SRR10121261 | *Onthophagus taurus* | SRR10121261 | wildtype | white pupa | t1 drosal imaginal tissue | RNAseq |

| Sample_id | Species | Data Source | Background | Stage | Tissue | RNA Library |
|---|---|---|---|---|---|---|
| OtauM_T1NepL_SRR10121271 | *Onthophagus taurus* | SRR10121271 | wildtype | white pupa | t1 drosal imaginal tissue | RNAseq |
| OtauM_T1NepP_SRR10121225 | *Onthophagus taurus* | SRR10121225 | wildtype | white pupa | t1 drosal imaginal tissue | RNAseq |
| OtauM_T1NepP_SRR10121235 | *Onthophagus taurus* | SRR10121235 | wildtype | white pupa | t1 drosal imaginal tissue | RNAseq |
| OtauM_T1NepP_SRR10121245 | *Onthophagus taurus* | SRR10121245 | wildtype | white pupa | t1 drosal imaginal tissue | RNAseq |
| OtauM_T1NepP_SRR10121255 | *Onthophagus taurus* | SRR10121255 | wildtype | white pupa | t1 drosal imaginal tissue | RNAseq |
| OtauM_T1NepP_SRR10121265 | *Onthophagus taurus* | SRR10121265 | wildtype | white pupa | t1 drosal imaginal tissue | RNAseq |
| OtauM_T1NepP_SRR10121275 | *Onthophagus taurus* | SRR10121275 | wildtype | white pupa | t1 drosal imaginal tissue | RNAseq |
| OtauM_T1NepR_SRR10121222 | *Onthophagus taurus* | SRR10121222 | wildtype | white pupa | t1 drosal imaginal tissue | RNAseq |
| OtauM_T1NepR_SRR10121232 | *Onthophagus taurus* | SRR10121232 | wildtype | white pupa | t1 drosal imaginal tissue | RNAseq |
| OtauM_T1NepR_SRR10121242 | *Onthophagus taurus* | SRR10121242 | wildtype | white pupa | t1 drosal imaginal tissue | RNAseq |
| OtauM_T1NepR_SRR10121252 | *Onthophagus taurus* | SRR10121252 | wildtype | white pupa | t1 drosal imaginal tissue | RNAseq |
| OtauM_T1NepR_SRR10121262 | *Onthophagus taurus* | SRR10121262 | wildtype | white pupa | t1 drosal imaginal tissue | RNAseq |
| OtauM_T1NepR_SRR10121272 | *Onthophagus taurus* | SRR10121272 | wildtype | white pupa | t1 drosal imaginal tissue | RNAseq |
| OtauM_T1_SRR10121224 | *Onthophagus taurus* | SRR10121224 | wildtype | white pupa | t1 drosal imaginal tissue | RNAseq |
| OtauM_T1_SRR10121234 | *Onthophagus taurus* | SRR10121234 | wildtype | white pupa | t1 drosal imaginal tissue | RNAseq |
| OtauM_T1_SRR10121244 | *Onthophagus taurus* | SRR10121244 | wildtype | white pupa | t1 drosal imaginal tissue | RNAseq |
| OtauM_T1_SRR10121254 | *Onthophagus taurus* | SRR10121254 | wildtype | white pupa | t1 drosal imaginal tissue | RNAseq |
| OtauM_T1_SRR10121264 | *Onthophagus taurus* | SRR10121264 | wildtype | white pupa | t1 drosal imaginal tissue | RNAseq |
| OtauM_T1_SRR10121274 | *Onthophagus taurus* | SRR10121274 | wildtype | white pupa | t1 drosal imaginal tissue | RNAseq |

| Sample_id | Species | Data Source | Background | Stage | Tissue | RNA Library |
|---|---|---|---|---|---|---|
| OtauM_T2L_SRR10121226 | *Onthophagus taurus* | SRR10121226 | wildtype | white pupa | t2 drosal imaginal tissue | RNAseq |
| OtauM_T2L_SRR10121236 | *Onthophagus taurus* | SRR10121236 | wildtype | white pupa | t2 drosal imaginal tissue | RNAseq |
| OtauM_T2L_SRR10121246 | *Onthophagus taurus* | SRR10121246 | wildtype | white pupa | t2 drosal imaginal tissue | RNAseq |
| OtauM_T2L_SRR10121256 | *Onthophagus taurus* | SRR10121256 | wildtype | white pupa | t2 drosal imaginal tissue | RNAseq |
| OtauM_T2L_SRR10121266 | *Onthophagus taurus* | SRR10121266 | wildtype | white pupa | t2 drosal imaginal tissue | RNAseq |
| OtauM_T2L_SRR10121276 | *Onthophagus taurus* | SRR10121276 | wildtype | white pupa | t2 drosal imaginal tissue | RNAseq |
| OtauM_T2R_SRR10121227 | *Onthophagus taurus* | SRR10121227 | wildtype | white pupa | t2 drosal imaginal tissue | RNAseq |
| OtauM_T2R_SRR10121237 | *Onthophagus taurus* | SRR10121237 | wildtype | white pupa | t2 drosal imaginal tissue | RNAseq |
| OtauM_T2R_SRR10121247 | *Onthophagus taurus* | SRR10121247 | wildtype | white pupa | t2 drosal imaginal tissue | RNAseq |
| OtauM_T2R_SRR10121257 | *Onthophagus taurus* | SRR10121257 | wildtype | white pupa | t2 drosal imaginal tissue | RNAseq |
| OtauM_T2R_SRR10121267 | *Onthophagus taurus* | SRR10121267 | wildtype | white pupa | t2 drosal imaginal tissue | RNAseq |
| OtauM_T2R_SRR10121277 | *Onthophagus taurus* | SRR10121277 | wildtype | white pupa | t2 drosal imaginal tissue | RNAseq |
| Otau_Body_SRR10664631 | *Onthophagus taurus* | SRR10664631 | wildtype | white pupa | wholebody | RNAseq |
| Otau_Body_SRR10664632 | *Onthophagus taurus* | SRR10664632 | wildtype | white pupa | wholebody | RNAseq |
| Otau_Body_SRR10664634 | *Onthophagus taurus* | SRR10664634 | wildtype | white pupa | wholebody | RNAseq |
| Otau_Body_SRR10664635 | *Onthophagus taurus* | SRR10664635 | wildtype | white pupa | wholebody | RNAseq |
| Otau_Body_SRR10664636 | *Onthophagus taurus* | SRR10664636 | wildtype | white pupa | wholebody | RNAseq |
| Otau_Body_SRR10664637 | *Onthophagus taurus* | SRR10664637 | wildtype | white pupa | wholebody | RNAseq |
| Otau_Body_SRR10664658 | *Onthophagus taurus* | SRR10664658 | wildtype | white pupa | wholebody | RNAseq |
| Otau_Body_SRR10664659 | *Onthophagus taurus* | SRR10664659 | wildtype | white pupa | wholebody | RNAseq |
| Otau_Body_SRR10664660 | *Onthophagus taurus* | SRR10664660 | wildtype | white pupa | wholebody | RNAseq |
| Otau_Body_SRR10664661 | *Onthophagus taurus* | SRR10664661 | wildtype | white pupa | wholebody | RNAseq |
| Otau_Body_SRR10664662 | *Onthophagus taurus* | SRR10664662 | wildtype | white pupa | wholebody | RNAseq |

| Sample_id | Species | Data Source | Background | Stage | Tissue | RNA Library |
|---|---|---|---|---|---|---|
| Otau_Body_SRR10664663 | *Onthophagus taurus* | SRR10664663 | wildtype | white pupa | wholebody | RNAseq |
| TdicM_T1_SRR7906881 | *Trypoxylus dichotomus* | SRR7906881 | wildtype | prepupating larva | t1 drosal imaginal tissue | RNAseq |
| TdicM_T1_SRR7906885 | *Trypoxylus dichotomus* | SRR7906885 | wildtype | prepupating larva | t1 drosal imaginal tissue | RNAseq |
| TdicM_T1_SRR7906896 | *Trypoxylus dichotomus* | SRR7906896 | wildtype | prepupating larva | t1 drosal imaginal tissue | RNAseq |
| TdicM_T1_SRR7906906 | *Trypoxylus dichotomus* | SRR7906906 | wildtype | prepupating larva | t1 drosal imaginal tissue | RNAseq |
| TdicM_T1_SRR7906910 | *Trypoxylus dichotomus* | SRR7906910 | wildtype | prepupating larva | t1 drosal imaginal tissue | RNAseq |
| TdicM_T1_SRR7906914 | *Trypoxylus dichotomus* | SRR7906914 | wildtype | prepupating larva | t1 drosal imaginal tissue | RNAseq |
| TdicM_T1_SRR7906924 | *Trypoxylus dichotomus* | SRR7906924 | wildtype | prepupating larva | t1 drosal imaginal tissue | RNAseq |
| TdicM_T2_SRR7906884 | *Trypoxylus dichotomus* | SRR7906884 | wildtype | prepupating larva | t2 drosal imaginal tissue | RNAseq |
| TdicM_T2_SRR7906887 | *Trypoxylus dichotomus* | SRR7906887 | wildtype | prepupating larva | t2 drosal imaginal tissue | RNAseq |
| TdicM_T2_SRR7906899 | *Trypoxylus dichotomus* | SRR7906899 | wildtype | prepupating larva | t2 drosal imaginal tissue | RNAseq |
| TdicM_T2_SRR7906903 | *Trypoxylus dichotomus* | SRR7906903 | wildtype | prepupating larva | t2 drosal imaginal tissue | RNAseq |
| TdicM_T2_SRR7906907 | *Trypoxylus dichotomus* | SRR7906907 | wildtype | prepupating larva | t2 drosal imaginal tissue | RNAseq |
| TdicM_T2_SRR7906911 | *Trypoxylus dichotomus* | SRR7906911 | wildtype | prepupating larva | t2 drosal imaginal tissue | RNAseq |
| TdicM_T2_SRR7906925 | *Trypoxylus dichotomus* | SRR7906925 | wildtype | prepupating larva | t2 drosal imaginal tissue | RNAseq |
| Tdic_Fat_DRR332810 | *Trypoxylus dichotomus* | DRR332810 | wildtype | prepupating larva | Fat | RNAseq |
| Tdic_Fat_DRR332812 | *Trypoxylus dichotomus* | DRR332812 | wildtype | prepupating larva | Fat | RNAseq |
| Tdic_Gut_DRR332806 | *Trypoxylus dichotomus* | DRR332806 | wildtype | prepupating larva | Gut | RNAseq |
| Tdic_Gut_DRR332808 | *Trypoxylus dichotomus* | DRR332808 | wildtype | prepupating larva | Gut | RNAseq |
| Ecar_Abd_SRR9942946 | *Entylia carinata* | SRR9942946 | wildtype | final nymph stage | wholebody eqv | RNAseq |
| Ecar_Abd_SRR9942958 | *Entylia carinata* | SRR9942958 | wildtype | final nymph stage | wholebody eqv | RNAseq |
| Ecar_Abd_SRR9942965 | *Entylia carinata* | SRR9942965 | wildtype | final nymph stage | wholebody eqv | RNAseq |

| Sample_id | Species | Data Source | Background | Stage | Tissue | RNA Library |
|---|---|---|---|---|---|---|
| Ecar_T1_SRR9942930 | *Entylia carinata* | SRR9942930 | wildtype | final nymph stage | t1 drosal imaginal tissue | RNAseq |
| Ecar_T1_SRR9942962 | *Entylia carinata* | SRR9942962 | wildtype | final nymph stage | t1 drosal imaginal tissue | RNAseq |
| Ecar_T1_SRR9942968 | *Entylia carinata* | SRR9942968 | wildtype | final nymph stage | t1 drosal imaginal tissue | RNAseq |
| Ecar_T2_SRR7507055 | *Entylia carinata* | SRR7507055 | wildtype | final nymph stage | t2 drosal imaginal tissue | RNAseq |
| Ecar_T2_SRR7507056 | *Entylia carinata* | SRR7507056 | wildtype | final nymph stage | t2 drosal imaginal tissue | RNAseq |
| Ecar_T2_SRR7507061 | *Entylia carinata* | SRR7507061 | wildtype | final nymph stage | t2 drosal imaginal tissue | RNAseq |
| Ecar_T2_SRR9942929 | *Entylia carinata* | SRR9942929 | wildtype | final nymph stage | t2 drosal imaginal tissue | RNAseq |
| Ecar_T2_SRR9942931 | *Entylia carinata* | SRR9942931 | wildtype | final nymph stage | t2 drosal imaginal tissue | RNAseq |
| Ecar_T2_SRR9942934 | *Entylia carinata* | SRR9942934 | wildtype | final nymph stage | t2 drosal imaginal tissue | RNAseq |
| Hvit_Wholebody_SRR10060919 | *Homalodisca vitripennis* | SRR10060919 | wildtype | final nymph stage | wholebody | RNAseq |
| Hvit_Wholebody_SRR1865088 | *Homalodisca vitripennis* | SRR1865088 | wildtype | final nymph stage | wholebody | RNAseq |
| Hvit_Wholebody_SRR1865089 | *Homalodisca vitripennis* | SRR1865089 | wildtype | final nymph stage | wholebody | RNAseq |
| Hvit_T1_SRR9942964 | *Homalodisca vitripennis* | SRR9942964 | wildtype | final nymph stage | t1 drosal imaginal tissue | RNAseq |
| Hvit_T1_SRR9942967 | *Homalodisca vitripennis* | SRR9942967 | wildtype | final nymph stage | t1 drosal imaginal tissue | RNAseq |
| Hvit_T1_SRR9942973 | *Homalodisca vitripennis* | SRR9942973 | wildtype | final nymph stage | t1 drosal imaginal tissue | RNAseq |
| Hvit_T2_SRR7507047 | *Homalodisca vitripennis* | SRR7507047 | wildtype | final nymph stage | t2 drosal imaginal tissue | RNAseq |
| Hvit_T2_SRR7507052 | *Homalodisca vitripennis* | SRR7507052 | wildtype | final nymph stage | t2 drosal imaginal tissue | RNAseq |
| Hvit_T2_SRR7507062 | *Homalodisca vitripennis* | SRR7507062 | wildtype | final nymph stage | t2 drosal imaginal tissue | RNAseq |
| Hvit_T2_SRR9942949 | *Homalodisca vitripennis* | SRR9942949 | wildtype | final nymph stage | t2 drosal imaginal tissue | RNAseq |

| Sample_id | Species | Data Source | Background | Stage | Tissue | RNA Library |
|---|---|---|---|---|---|---|
| Hvit_T2_SRR9942952 | *Homalodisca vitripennis* | SRR9942952 | wildtype | final nymph stage | t2 drosal imaginal tissue | RNAseq |
| Hvit_T2_SRR9942971 | *Homalodisca vitripennis* | SRR9942971 | wildtype | final nymph stage | t2 drosal imaginal tissue | RNAseq |

**Table S6. Top 100 marker genes that determines the cell identity of *D. melanogaster* PROD in scRNAseq.**

| Top 100 marker gene | Epithelium | Trachea | AMPs | Hemocytes |
|---|---|---|---|---|
| 1 | *Antp* | *CG13044* | *zfh1* | *CG14629* |
| 2 | *ct* | *vvl* | *SPARC* | *CG33494* |
| 3 | *CG13044* | *zfh2* | *Act57B* | *srp* |
| 4 | *CG30154* | *CG30154* | *Him* | *Had2* |
| 5 | *l(2)34Fc* | *l(2)34Fc* | *CG9650* | *zfh1* |
| 6 | *tpr* | *tpr* | *twi* | *CG4250* |
| 7 | *CG2816* | *CG2816* | *Act87E* | *lectin-28C* |
| 8 | *trol* | *CG11370* | *htl* | *CG33460* |
| 9 | *CG31997* | *ct* | *nemy* | *CG31777* |
| 10 | *zfh2* | *Gs2* | *org-1* | *CG31337* |
| 11 | *vvl* | *CG11200* | *CG9593* | *Mec2* |
| 12 | *Phk-3* | *Antp* | *CG11835* | *CG4793* |
| 13 | *CG11370* | *apt* | *Col4a1* | *Karl* |
| 14 | *apt* | *CG8630* | *vkg* | *CG4259* |
| 15 | *stumps* | *stumps* | *Mes2* | *lectin-24Db* |
| 16 | *hui* | *Spn43Aa* | *beat-IIIc* | *GlcAT-P* |
| 17 | *Gs2* | *CG13678* | *side* | *Oat* |
| 18 | *Inos* | *Inos* | *Mal-A5* | *nAChRbeta3* |
| 19 | *CG11200* | *CG9095* | *CG18557* | *NimC3* |
| 20 | *CG8630* | *trol* | *E(spl)m6-BFM* | *Arpc3B* |
| 21 | *CG13678* | *CG2016* | *unc-4* | *CG42807* |
| 22 | *CG44325* | *Tsf1* | *Mp* | *Aldh7A1* |
| 23 | *Spn43Aa* | *Pgant2* | *SerT* | *He* |
| 24 | *Tsf1* | *lncRNA:CR43302* | *sens-2* | *Tep1* |
| 25 | *CG2016* | *CG14566* | *trol* | *CG30148* |
| 26 | *mbl* | *Phk-3* | *miple1* | *firl* |
| 27 | *lncRNA:CR43302* | *CG8483* | *Slc45-1* | *eater* |
| 28 | *olf413* | *CG31997* | *kon* | *P5cr-2* |
| 29 | *Pgant2* | *CG3502* | *tow* | *CG31431* |
| 30 | *Egfr* | *hui* | *egr* | *Ilp6* |
| 31 | *CG14566* | *Ubx* | *CadN* | *Pxn* |
| 32 | *CG9095* | *CG13082* | *Nep3* | *LManI* |
| 33 | *Kank* | *spz5* | *Octalpha2R* | *CG30088* |
| 34 | *ImpE1* | *rk* | *Nna1* | *NimB4* |
| 35 | *CG8483* | *CG13639* | *Nlg1* | *CG4927* |
| 36 | *nkd* | *Ddc* | *Sox100B* | *CG30090* |
| 37 | *CG3502* | *serp* | *Oat* | *CG13315* |
| 38 | *frm* | *CG13272* | *CG45076* | *mthl2* |
| 39 | *CG17919* | *CG12009* | *grk* | *NtR* |
| 40 | *Ubx* | *mbl* | *Timp* | *CG43236* |

| Top 100 marker gene | Epithelium | Trachea | AMPs | Hemocytes |
|---|---|---|---|---|
| 41 | *Gp150* | *CG5532* | *CG12768* | *Eip93F* |
| 42 | *rk* | *bond* | *Grip* | *Sid* |
| 43 | *spz5* | *olf413* | *Poxm* | *CG43124* |
| 44 | *ara* | *CG44325* | *fau* | *CG34331* |
| 45 | *CG13082* | *Ance* | *lncRNA:CR45361* | *CG43133* |
| 46 | *ab* | *btl* | *Ilp6* | *CG7091* |
| 47 | *CG9691* | *Egfr* | *Con* | *CG10764* |
| 48 | *CG16798* | *drd* | *CG12984* | *lncRNA:CR44133* |
| 49 | *zfh1* | *ImpE1* | *CG31997* | *Hml* |
| 50 | *rgn* | *Hsp27* | *CG3168* | *CG30046* |
| 51 | *Ance* | *CG15353* | *Fas3* | *jtb* |
| 52 | *CG13639* | *ect* | *pio* | *ham* |
| 53 | *Df31* | *peb* | *Ugt317A1* | *et* |
| 54 | *Hsp27* | *RpL7A* | *Wnt4* | *CG1544* |
| 55 | *klu* | *CG13067* | *Cyp310a1* | *Nplp2* |
| 56 | *CG5532* | *Kank* | *crb* | *mthl6* |
| 57 | *CG13272* | *CG9372* | *CG42346* | *CG31778* |
| 58 | *CG12009* | *Cpr64Ad* | *ovo* | *CG15347* |
| 59 | *CG9689* | *ara* | *zyd* | *lncRNA:CR43855* |
| 60 | *Ten-a* | *CG16798* | *mspo* | *CG7470* |
| 61 | *Ddc* | *nkd* | *heph* | *CG4408* |
| 62 | *l(3)neo38* | *dpy* | *grh* | *CG42694* |
| 63 | *Oaz* | *Df31* | *CG7800* | *Lip4* |
| 64 | *heph* | *rgn* | *meso18E* | *Nepl14* |
| 65 | *caup* | *RpL5* | *Msr-110* | *CG8080* |
| 66 | *btl* | *RpL10Ab* | *GILT1* | *NimB1* |
| 67 | *pio* | *CG17919* | *Cht10* | *NimB5* |
| 68 | *svp* | *CG13003* | *Ptx1* | *CG4950* |
| 69 | *CG14439* | *frm* | *Phk-3* | *CG5958* |
| 70 | *CG4455* | *RpS4* | *sano* | *lncRNA:CR45018* |
| 71 | *bond* | *klu* | *Msp300* | *dpr3* |
| 72 | *drd* | *CG11905* | *mab-21* | *rgr* |
| 73 | *ect* | *Edg91* | *CG15628* | *CG34437* |
| 74 | *CG9372* | *l(2)k05911* | *Gp150* | *CG33458* |
| 75 | *Lac* | *ab* | *Antp* | *SPARC* |
| 76 | *mab-21* | *RpL27* | *ft* | *santa-maria* |
| 77 | *wbl* | *CG4678* | *FBti0060302* | *Col4a1* |
| 78 | *CG7970* | *RpS18* | *Lac* | *CG1092* |
| 79 | *Ppn* | *CG8925* | *ct* | *CG3831* |
| 80 | *CG15353* | *Hsp23* | *CG14516* | *CG42566* |
| 81 | *SPARC* | *magu* | *CG7675* | *NimC1* |
| 82 | *dsx-c73A* | *RpS3* | *Idgf4* | *Glt* |

| Top 100 marker gene | Epithelium | Trachea | AMPs | Hemocytes |
|---|---|---|---|---|
| 83 | *CG13067* | *Ten-a* | *CG33978* | *CG13559* |
| 84 | *sas* | *RpL36A* | *ed* | *kn* |
| 85 | *CG5397* | *miple2* | *Plp* | *LKRSDH* |
| 86 | *trn* | *CG8927* | *Inx2* | *Hexo2* |
| 87 | *Fas3* | *RpS29* | *nvy* | *CG4842* |
| 88 | *neo* | *CG14572* | *CG9691* | *Adgf-A* |
| 89 | *Oda* | *CG11147* | *FBti0019563* | *CG8620* |
| 90 | *Cpr64Ad* | *RpL9* | *verm* | *CG8501* |
| 91 | *CG42747* | *kni* | *CG17919* | *Irc* |
| 92 | *RpL7A* | *RpS2* | *GstO3* | *yellow-f* |
| 93 | *sesB* | *RpL37a* | *CG42390* | *kcc* |
| 94 | *Hsp67Ba* | *RpS23* | *dl* | *LManII* |
| 95 | *serp* | *CG7970* | *CrebA* | *Mes2* |
| 96 | *miple2* | *svp* | *hui* | *CG5321* |
| 97 | *Hsp23* | *RpS9* | *Sema2b* | *CG43125* |
| 98 | *GNBP3* | *Gp150* | *NetA* | *fat-spondin* |
| 99 | *Scr* | *Oaz* | *grn* | *CG17855* |
| 100 | *RpS4* | *CG9689* | *frm* | *alpha-Est8* |

**Table S7. The *D. melanogaster* stocks used in *GAL4* driver screening and expression pattern of morphogens in PROD.**

| Stock id/source | Genotype | Description |
|---|---|---|
| BDSC8715 | *ac-GAL4* | reporter; driver screening |
| BDSC26817 | *Antp-GAL4* | driver screening |
| BDSC3041 | *ap-GAL4* | reporter; driver screening |
| BDSC1560 | *arm-gal4* | reporter; driver screening |
| BDSC6354 | *bs-GAL4* | reporter; driver screening |
| BSDC78328 | *btl-Gal4* | reporter; driver screening |
| BDSC8860 | *bx.MS1096-GAL4* | reporter; driver screening |
| BDSC80604 | *CG34325-GAL4* | driver screening |
| this study | *ct.E-GAL4* | driver screening |
| KSC104700 | *discslost-GAL4* | reporter; driver screening |
| BDSC3038 | *Dll.md23-GAL4* | reporter; driver screening; cell lineage tracing |
| BDSC64307 | *Dll.md23-GAL4>UAS-GFP* | reporter; driver screening |
| BDSC1553 | *dpp.BLK-GAL4* | reporter; driver screening |
| BDSC7007 | *dpp.PS-GAL4* | driver screening |
| Xianjue Ma lab | *elav.c155-GAL4* | driver screening |
| BDSC6356 | *en-GAL4* | reporter; driver screening |
| BDSC67049 | *en-GAL4>UAS-GFP* | reporter; driver screening; knockdown screening & overexpression |
| Xianjue Ma lab | *esg-GAL4* | driver screening |
| BDSC9901 | *fng.11-26-GAL4* | driver screening |
| BDSC9891 | *fng.B-GAL4* | reporter; driver screening |
| Xianjue Ma lab | *hh-GAL4* | reporter; driver screening |
| BDSC65540 | *hth.1283-GAL4* | driver screening |
| Kyoto SC104957 | *hth.NP5332-GAL4* | driver screening |
| Xianjue Ma lab | *nub-GAL4* | reporter; driver screening |
| THUTB00015 | *pnr-GAL4* | reporter; driver screening |
| Xianjue Ma lab | *ptc-Gal4* | reporter; driver screening |
| BDSC80573 | *salm.EPv-GAL4* | driver screening |
| BDSC84331 | *salm.LP39-GAL4* | driver screening |
| BDSC43656 | *Scr-GAL4.4* | driver screening |
| BDSC42740 | *sens-GAL4* | reporter; driver screening |
| BDSC6791 | *Ser-GAL4* | reporter; driver screening |
| BDSC3040 | *tsh.md621-GAL4* | reporter; driver screening |
| BDSC6662 | *twi-GAL4* | reporter; driver screening |
| VDRC201689 | *VT026477-GAL4* | driver screening; knockdown screening & overexpression |
| BDSC11670 | *hth-lacZ* | reporter |
| BDSC4373 | *wg-lacZ* | reporter |
| THU1838 | *Ance RNAi* | knockdown screening |
| BDSC27675 | *Antp RNAi* | knockdown screening |
| BDSC41673 | *ap RNAi* | knockdown screening |

| Stock id/source | Genotype | Description |
|---|---|---|
| THU2446 | *ara RNAi* | knockdown screening |
| THU1911 | *arm RNAi* | knockdown screening |
| THU3622 | *bnl RNAi* | knockdown screening |
| THU5470 | *btl RNAi* | knockdown screening |
| THU2858 | *caps RNAi* | knockdown screening |
| VDRC105705 | *caup RNAi* | knockdown screening |
| TH02062.N | *CG15269 RNAi* | knockdown screening |
| THU2190 | *cher RNAi* | knockdown screening |
| THU2011 | *ci RNAi* | knockdown screening |
| THU2277 | *CrebA RNAi* | knockdown screening |
| BDSC33967 | *ct RNAi* | knockdown screening |
| BDSC34322 | *Dl RNAi* | knockdown screening |
| THU4897 | *Dif RNAi* | knockdown screening |
| BDSC29337 | *Dll RNAi* | knockdown screening |
| THU2310 | *Doc1 RNAi* | knockdown screening |
| BDSC36779 | *dpp RNAi* | knockdown screening |
| THU4947 | *drm RNAi* | knockdown screening |
| THU1912 | *dsh RNAi* | knockdown screening |
| THU2427 | *dx RNAi* | knockdown screening |
| BDSC31525 | *Egfr RNAi* | knockdown screening |
| THU5906 | *ems RNAi* | knockdown screening |
| BDSC26752 | *en RNAi* | knockdown screening |
| THU5422 | *esg RNAi* | knockdown screening |
| THU2520 | *exd RNAi* | knockdown screening |
| TH201500423.S | *fng RNAi* | knockdown screening |
| BDSC27656 | *foxO RNAi* | knockdown screening |
| THU0894 | *Frl RNAi* | knockdown screening |
| VDRC106288 | *ftz RNAi* | knockdown screening |
| VDRC105493 | *fz RNAi* | knockdown screening |
| TH201501080.S | *fz2 RNAi* | knockdown screening |
| BDSC25794 | *hh RNAi* | knockdown screening |
| THU2475 | *Hr39 RNAi* | knockdown screening |
| THU3284 | *Hr78 RNAi* | knockdown screening |
| THU2414 | *Hr96 RNAi* | knockdown screening |
| BDSC27655 | *hth RNAi* | knockdown screening |
| THU3065 | *kibra RNAi* | knockdown screening |
| THU2523 | *Lim1 RNAi* | knockdown screening |
| THU2340 | *lola RNAi* | knockdown screening |
| TH03144.N | *Lsp2 RNAi* | knockdown screening |
| THU2307 | *NfI RNAi* | knockdown screening |
| THU2923 | *nompA RNAi* | knockdown screening |
| BDSC28981 | *N RNAi* | knockdown screening |

| Stock id/source | Genotype | Description |
|---|---|---|
| BDSC28338 | *nub RNAi* | knockdown screening |
| THU1674 | *numb RNAi* | knockdown screening |
| THU3985 | *Oaz RNAi* | knockdown screening |
| THU5391 | *ovo RNAi* | knockdown screening |
| THU3865 | *par 6 RNAi* | knockdown screening |
| THU1004 | *pnr RNAi* | knockdown screening |
| BDSC28795 | *ptc RNAi* | knockdown screening |
| BDSC32476 | *Rbfox1 RNAi* | knockdown screening |
| THU4885 | *Rel RNAi* | knockdown screening |
| BDSC28675 | *sca RNAi* | knockdown screening |
| TH02003.N | *Scr RNAi* | knockdown screening |
| BDSC34700 | *Ser RNAi* | knockdown screening |
| THU3490 | *Su(H) RNAi* | knockdown screening |
| BDSC28689 | *svp RNAi* | knockdown screening |
| VDRC101175 | *tinc RNAi* | knockdown screening |
| BDSC33951 | *tor RNAi* | knockdown screening |
| THU2276 | *trh RNAi* | knockdown screening |
| BDSC50520 | *trn RNAi* | knockdown screening |
| BDSC36748 | *ttk RNAi* | knockdown screening |
| THU5846 | *tx RNAi* | knockdown screening |
| BDSC31970 | *vg RNAi* | knockdown screening |
| THU2169 | *Vps25 RNAi* | knockdown screening |
| BDSC31310 | *wg RNAi* | knockdown screening |
| BDSC31989 | *Wnt10 RNAi* | knockdown screening |
| THU1844 | *Xbp1 RNAi* | knockdown screening |
| BDSC28280 | *G-TRACE* | cell lineage tracing |
| BDSC4775 | *UAS-GFP.nls* | reporter |
| BDSC78349 | *UAS-Dll* | overexpression |
| BDSC9319 | *UAS-Dl* | overexpression |
| BDSC5815 | *UAS-Ser* | overexpression |

**Table S8. The antibodies used to investigate *D. melanogaster* PROD development.**

| Gene | Product Name | Source | Cat. no. or identifier | Host spp. | Clonality | Usage |
|---|---|---|---|---|---|---|
| *Scr* | anti-Scr | DSHB | 6H4.1 | mouse | monoclonal | 0.5-1 ug/ml |
| *Antp* | anti-Antp | DSHB | 4C3 | mouse | monoclonal | 0.5-1 ug/ml |
| *en/inv* | anti-en/inv | DSHB | 4D9 | mouse | monoclonal | 0.5-1 ug/ml |
| *exd* | anti-exd | DSHB | B11M | mouse | monoclonal | 0.5-1 ug/ml |
| *N* | anti-N | DSHB | C17.9C6 | mouse | monoclonal | 0.5-1 ug/ml |
| *Dl* | anti-Dl | DSHB | C594.9B | mouse | monoclonal | 0.5-1 ug/ml |
| *ct* | anti-ct | DSHB | 2B10 | mouse | monoclonal | 0.5-1 ug/ml |
| *ci* | anti-ci | DSHB | 2A1 | rat | monoclonal | 0.5-1 ug/ml |
| *Dll* | anti-Dll | gift from Dr. Siqian Feng | NA | guinea pig | polyclonal | 1:2000-1:4000 |
| *lacZ* | anti-lacZ | DSHB | 40-1a | mouse | monoclonal | 0.5-1 ug/ml |
| mouse *IgG* | Donkey Anti-Mouse AlexaFluor488 | Jackson | AB_2340846 | donkey | Polyclonal | 1:500 |
| mouse *IgG* | Goat anti Mouse AlexaFluor594 | Jackson | AB_2338871 | goat | Polyclonal | 1:500 |
| rat *IgG* | Goat anti Rat AlexaFluor488 | CST | 4416S | goat | Polyclonal | 1:500 |
| guinea pig *IgG* | Goat Anti-Guinea pig AlexaFluor488 | ABCAM | ab150185 | goat | Polyclonal | 1:500 |

**Table S9. Summary of phenotypic measurements in *D. melanogaster* RNAi screening to identify genes associated with PROD development.**

| Gene | Analysis | *en-gal4* | | | | | | *VT026477-gal4* | | | | | | Observable defects in branch | Observable defects in cone length |
|---|---|---|---|---|---|---|---|---|---|---|---|---|---|---|---|
| | | PROD branch | | | PROD length (μm) | | | PROD branch | | | PROD length (μm) | | | | |
| | | N | mean | SD | N | mean | SD | N | mean | SD | N | mean | SD | | |
| *Ance* | scRNAseq | 0 | NA | NA | 0 | NA | NA | 30 | 0.0 | 0.0 | 40 | 90.8 | 42.2 | √ | √ |
| *Antp* | DEGs, scRNAseq | 20 | 2.8 | 2.8 | 44 | 151.8 | 39.2 | 20 | 5.7 | 1.7 | 40 | 191.6 | 54.9 | √ | √ |
| *ap* | DEGs, scRNAseq | 18 | 7.3 | 0.6 | 24 | 160.8 | 49.2 | 21 | 8.0 | 0.7 | 40 | 248.4 | 26.9 | | |
| *ara* | DEGs, scRNAseq | 23 | 7.8 | 0.5 | 38 | 192.3 | 42.1 | 25 | 8.4 | 0.6 | 40 | 264.3 | 27.9 | | |
| *arm* | scRNAseq | 27 | 7.2 | 0.9 | 40 | 219.3 | 58.2 | 23 | 5.2 | 3.1 | 40 | 166.7 | 96.9 | | √ |
| *bnl* | association | 17 | 7.6 | 0.5 | 40 | 181.2 | 75.1 | 21 | 7.6 | 0.6 | 40 | 243.2 | 32.6 | | √ |
| *btl* | scRNAseq | 0 | NA | NA | 0 | NA | NA | 27 | 2.4 | 3.8 | 40 | 156.8 | 66.6 | √ | √ |
| *caps* | association | 23 | 8.0 | 0.8 | 40 | 236.5 | 76.4 | 27 | 8.0 | 0.5 | 40 | 244.1 | 39.7 | | |
| *caup* | scRNAseq | 10 | 7.1 | 1.1 | 40 | 215.8 | 35.9 | 26 | 8.3 | 0.5 | 40 | 232.1 | 32.5 | | |
| *CG15269* | scRNAseq | 5 | 5.6 | 0.5 | 40 | 113.3 | 59.8 | 23 | 2.9 | 3.4 | 40 | 189.3 | 54.2 | √ | √ |
| *cher* | DEGs | 27 | 7.9 | 0.5 | 40 | 252.3 | 51.0 | 23 | 8.4 | 0.8 | 40 | 234.5 | 28.3 | | |
| *ci* | DEGs | 22 | 8.0 | 0.6 | 40 | 218.1 | 22.9 | 20 | 8.5 | 0.5 | 40 | 250.9 | 26.9 | | √ |
| *CrebA* | scRNAseq | 17 | 7.6 | 0.6 | 40 | 159.1 | 29.4 | 23 | 8.2 | 0.6 | 40 | 231.2 | 27.7 | | √ |
| *ct* | DEGs, scRNAseq | 26 | 0.0 | 0.0 | 40 | 0.0 | 0.0 | 26 | 0.0 | 0.0 | 40 | 2.4 | 10.4 | √ | √ |
| *Dl* | DEGs | 24 | 0.0 | 0.0 | 40 | 0.0 | 0.0 | 22 | 0.2 | 0.6 | 40 | 103.3 | 53.1 | √ | √ |
| *Dif* | association | 26 | 7.8 | 0.7 | 40 | 230.4 | 63.5 | 23 | 7.7 | 1.7 | 40 | 269.7 | 29.2 | | |
| *Dll* | DEGs | 24 | 8.0 | 0.4 | 40 | 116.2 | 102.4 | 22 | 7.4 | 2.4 | 40 | 221.2 | 27.1 | | √ |
| *Doc1* | association | 25 | 7.8 | 0.5 | 40 | 224.1 | 75.7 | 22 | 8.3 | 0.6 | 40 | 250.9 | 29.4 | | √ |
| *dpp* | DEG | 17 | 7.5 | 0.5 | 40 | 165.8 | 88.7 | 21 | 8.3 | 0.6 | 40 | 227.4 | 32.4 | | √ |
| *drm* | association | 18 | 8.5 | 0.6 | 40 | 258.0 | 47.8 | 16 | 8.3 | 0.4 | 40 | 219.0 | 82.4 | | |
| *dsh* | association | 30 | 8.2 | 0.5 | 40 | 189.4 | 87.1 | 26 | 0.0 | 0.0 | 40 | 182.2 | 62.9 | | √ |
| *dx* | DEGs | 28 | 7.5 | 0.6 | 40 | 190.6 | 50.4 | 30 | 8.3 | 0.6 | 40 | 258.0 | 30.7 | | |

| Gene | Analysis | *en-gal4* | | | | | | *VT026477-gal4* | | | | | | Observable defects in branch | Observable defects in cone length |
|---|---|---|---|---|---|---|---|---|---|---|---|---|---|---|---|
| | | PROD branch | | | PROD length (μm) | | | PROD branch | | | PROD length (μm) | | | | |
| | | N | mean | SD | N | mean | SD | N | mean | SD | N | mean | SD | | |
| *Egfr* | DEGs, scRNAseq | 19 | 3.4 | 1.8 | 40 | 106.3 | 77.5 | 22 | 1.0 | 1.5 | 40 | 148.2 | 35.4 | √ | √ |
| *ems* | DEGs | 24 | 7.7 | 1.8 | 40 | 263.0 | 26.8 | 24 | 8.0 | 1.8 | 40 | 246.2 | 29.3 | | |
| *en* | DEGs | 23 | 7.8 | 0.7 | 40 | 135.0 | 48.6 | 22 | 7.9 | 0.6 | 40 | 244.7 | 50.8 | | √ |
| *esg* | association | 0 | NA | NA | 0 | NA | NA | 23 | 8.3 | 0.6 | 40 | 260.5 | 30.1 | | √ |
| *exd* | association | 16 | 7.2 | 2.9 | 40 | 92.6 | 75.5 | 26 | 8.0 | 0.6 | 40 | 210.8 | 27.5 | | √ |
| *fng* | association | 0 | NA | NA | 0 | 229.9 | 12.9 | 23 | 7.7 | 1.7 | 40 | 241.2 | 23.4 | | |
| *foxO* | DEGs | 24 | 8.0 | 0.5 | 40 | 242.9 | 33.0 | 23 | 8.4 | 0.6 | 40 | 272.3 | 54.5 | | |
| *Frl* | DEGs | 28 | 7.9 | 0.6 | 40 | 254.7 | 64.8 | 22 | 8.3 | 0.6 | 40 | 207.3 | 56.2 | | |
| *ftz* | DEGs | 23 | 7.6 | 0.8 | 40 | 241.1 | 25.4 | 19 | 7.7 | 2.0 | 40 | 238.5 | 26.8 | | |
| *fz* | DEGs | 23 | 7.7 | 1.1 | 40 | 245.6 | 27.3 | 27 | 8.0 | 1.7 | 40 | 233.0 | 22.8 | | |
| *fz2* | DEGs | 0 | NA | NA | 0 | NA | NA | 22 | 8.3 | 0.6 | 40 | 240.0 | 60.9 | | |
| *hh* | association | 3 | 7.0 | 1.0 | 22 | 80.1 | 78.0 | 22 | 8.0 | 0.6 | 40 | 224.0 | 50.1 | | √ |
| *Hr39* | DEGs | 22 | 8.0 | 0.5 | 40 | 236.5 | 27.4 | 20 | 8.0 | 2.0 | 40 | 237.1 | 30.0 | | |
| *Hr78* | association | 0 | NA | NA | 0 | NA | NA | 19 | 8.2 | 0.6 | 40 | 271.9 | 21.2 | | |
| *Hr96* | association | 13 | 7.8 | 0.6 | 40 | 256.3 | 37.5 | 16 | 8.3 | 0.4 | 40 | 258.1 | 51.7 | | |
| *hth* | DEGs | 20 | 2.0 | 3.5 | 40 | 0.0 | 0.0 | 21 | 8.1 | 0.4 | 40 | 229.3 | 26.8 | | √ |
| *kibra* | DEGs | 26 | 7.3 | 2.2 | 40 | 219.7 | 62.3 | 22 | 8.5 | 0.6 | 40 | 244.0 | 26.2 | | |
| *Lim1* | association | 19 | 7.2 | 2.6 | 40 | 257.5 | 26.7 | 26 | 8.2 | 0.7 | 40 | 263.1 | 27.8 | | |
| *lola* | association | 20 | 7.9 | 0.6 | 40 | 195.5 | 103.8 | 20 | 8.1 | 0.6 | 40 | 269.0 | 29.2 | | |
| *Lsp2* | association | 0 | NA | NA | 0 | NA | NA | 20 | 8.1 | 0.7 | 40 | 266.7 | 22.9 | | |
| *NfI* | DEGs | 20 | 7.8 | 0.7 | 40 | 266.3 | 26.4 | 21 | 2.8 | 4.0 | 40 | 259.8 | 23.6 | | √ |
| *nompA* | DEGs | 20 | 7.9 | 0.6 | 40 | 203.9 | 100.2 | 20 | 7.9 | 0.6 | 40 | 248.2 | 51.5 | | |
| *N* | DEGs | 26 | 0.0 | 0.0 | 40 | 0.0 | 0.0 | 24 | 0.0 | 0.0 | 24 | 79.0 | 44.5 | √ | √ |
| *nub* | DEGs | 10 | 7.7 | 0.5 | 40 | 231.5 | 54.3 | 23 | 7.6 | 1.9 | 40 | 232.8 | 56.9 | | |
| *numb* | DEGs | 11 | 7.8 | 0.6 | 40 | 205.8 | 86.0 | 20 | 6.9 | 3.0 | 40 | 260.6 | 29.0 | | |

| Gene | Analysis | *en-gal4* | | | | | | *VT026477-gal4* | | | | | | Observable defects in branch | Observable defects in cone length |
|---|---|---|---|---|---|---|---|---|---|---|---|---|---|---|---|
| | | PROD branch | | | PROD length (μm) | | | PROD branch | | | PROD length (μm) | | | | |
| | | N | mean | SD | N | mean | SD | N | mean | SD | N | mean | SD | | |
| *Oaz* | scRNAseq | 13 | 8.1 | 0.3 | 40 | 270.6 | 27.4 | 29 | 8.4 | 0.6 | 40 | 233.2 | 30.0 | | |
| *ovo* | DEGs | 0 | NA | NA | 0 | NA | NA | 18 | 8.2 | 0.7 | 40 | 243.2 | 49.6 | | |
| *par 6* | DEGs | 26 | 0.0 | 0.0 | 50 | 42.5 | 98.4 | 20 | 0.0 | 0.0 | 40 | 206.0 | 57.5 | √ | √ |
| *pnr* | association | 27 | 8.0 | 0.6 | 40 | 217.6 | 25.7 | 20 | 8.1 | 0.5 | 40 | 245.7 | 39.0 | | |
| *ptc* | DEGs | 25 | 7.8 | 0.7 | 32 | 217.1 | 67.5 | 21 | 8.0 | 0.5 | 40 | 223.7 | 41.6 | | |
| *Rbfox1* | DEGs | 23 | 8.0 | 0.5 | 40 | 257.0 | 55.1 | 20 | 8.0 | 0.0 | 40 | 251.9 | 29.9 | | |
| *Rel* | DEGs | 24 | 8.0 | 0.5 | 40 | 232.6 | 94.8 | 21 | 8.0 | 0.5 | 40 | 250.8 | 25.9 | | |
| *sca* | DEGs | 18 | 8.1 | 0.5 | 40 | 243.9 | 62.3 | 25 | 8.4 | 0.5 | 40 | 242.2 | 64.3 | | |
| *Scr* | DEGs, scRNAseq | 18 | 7.9 | 0.6 | 40 | 186.9 | 46.1 | 20 | 8.5 | 0.7 | 40 | 245.3 | 53.0 | | √ |
| *Ser* | DEGs | 0 | NA | NA | 0 | NA | NA | 23 | 6.6 | 0.9 | 40 | 223.9 | 23.3 | √ | √ |
| *Su(H)* | DEGs | 26 | 7.0 | 1.1 | 40 | 167.9 | 63.8 | 22 | 7.9 | 0.7 | 40 | 244.1 | 53.9 | | |
| *svp* | scRNAseq | 28 | 0.0 | 0.0 | 40 | 213.9 | 63.9 | 21 | 8.2 | 0.5 | 40 | 221.4 | 44.3 | | |
| *tinc* | DEGs | 17 | 7.9 | 0.4 | 40 | 256.0 | 35.0 | 22 | 8.0 | 0.6 | 40 | 214.8 | 42.2 | | |
| *tor* | association | 30 | 0.0 | 0.0 | 40 | 102.7 | 119.4 | 26 | 0.0 | 0.0 | 40 | 192.6 | 47.4 | √ | √ |
| *trh* | DEGs | 19 | 8.1 | 0.5 | 40 | 236.7 | 72.1 | 22 | 8.4 | 0.7 | 40 | 252.2 | 21.2 | | |
| *trn* | DEGs | 21 | 7.7 | 0.5 | 40 | 260.2 | 30.7 | 21 | 8.4 | 0.5 | 40 | 249.1 | 32.3 | | |
| *ttk* | DEGs | 0 | NA | NA | 0 | NA | NA | 20 | 0.0 | 0.0 | 40 | 152.2 | 30.2 | | √ |
| *tx* | DEGs | 22 | 0.0 | 0.0 | 14 | 0.0 | 0.0 | 30 | 0.0 | 0.0 | 40 | 85.1 | 59.6 | | √ |
| *vg* | DEGs | 0 | NA | NA | 0 | NA | NA | 22 | 8.0 | 0.7 | 40 | 257.4 | 31.6 | | |
| *Vps25* | DEGs | 0 | NA | NA | 0 | NA | NA | 24 | 7.8 | 2.5 | 40 | 240.8 | 33.5 | | |
| *wg* | association | 0 | NA | NA | 0 | NA | NA | 24 | 8.0 | 0.7 | 40 | 225.2 | 27.8 | | |
| *Wnt10* | DEGs | 20 | 8.0 | 0.5 | 40 | 251.9 | 37.8 | 25 | 8.0 | 0.6 | 40 | 238.4 | 49.2 | | |
| *Xbp1* | association | 23 | 7.9 | 0.7 | 40 | 220.6 | 48.9 | 22 | 8.3 | 0.7 | 40 | 203.5 | 41.2 | | |

**Table S10. List of *A. albopictus* primers used for dsRNA syntheses.**

| | Gene | Produced length | T7+forward | T7+reverse |
|---|---|---|---|---|
| 1 | *Aa-Dll* | 495 | TAATACGACTCACTATAGATGATCAGCGCCCGAAAGAT | TAATACGACTCACTATAGAAATGCTGGTACCCACTCCG |
| 2 | *Aa-exd* | 408 | TAATACGACTCACTATAGTCCCTACCCCTCGGAAGAAG | TAATACGACTCACTATAGCTACGCCGGATCAGACAGAC |
| 3 | *Aa-hth* | 351 | TAATACGACTCACTATAGGATTGACGAACGGGACACCA | TAATACGACTCACTATAGCTCGCATCACCCTGTATACTTG |
| 4 | *Aa-N* | 347 | TAATACGACTCACTATAGTCGACAAAGTCGGAGGATTCG | TAATACGACTCACTATAGCTGACACAGTCCGTTGATGC |
| 5 | *Aa-Dl* | 457 | TAATACGACTCACTATAGGTACTTCGGGTCGGGATGTG | TAATACGACTCACTATAGGGCTGCACAGTTGCTAATGC |
| 6 | *Aa-Ser* | 570 | TAATACGACTCACTATAGTCTGTGATACCAACTGGGGC | TAATACGACTCACTATAGGCGAAACTTCCCCACCATGT |
| 7 | *Aa-wg* | 303 | TAATACGACTCACTATAGCAACTTCCAGCTGAAGCCGC | TAATACGACTCACTATAGCACAGGCATGTGTGAATAATCTTCT |
| 8 | *Aa-Wnt10* | 335 | TAATACGACTCACTATAGTCCGGCAGTTGTCAGTTGAA | TAATACGACTCACTATAGCCTGGTATGTCGGACACCTG |
| 9 | *Aa-hh* | 354 | TAATACGACTCACTATAGTAGTGAGAATTCGTTGGGCG | TAATACGACTCACTATAGCCAGCCTCGACGGCTAATC |
| 10 | *Aa-en* | 393 | TAATACGACTCACTATAGCGCGAGTTTCCCCTACTTCC | TAATACGACTCACTATAGTTTGCGGTTCGTAGCAGGTT |
| 11 | *Aa-ptc* | 407 | TAATACGACTCACTATAGCGACATCGAATGGAGCCTGA | TAATACGACTCACTATAGTTAGAGTTGACGCCACGTCC |
| 12 | *Aa-ci* | 467 | TAATACGACTCACTATAGGGTCTCACCTCGGAGTACCT | TAATACGACTCACTATAGGGTTTGGTTAGCGAATGCCC |
| 13 | *Aa-dpp* | 300 | TAATACGACTCACTATAGTCACGACAAGTGGTCTCACA | TAATACGACTCACTATAGATCGCGTGATTCGTCGTGTT |
| 14 | *Aa-Scr* | 406 | TAATACGACTCACTATAGTCTTAGCACGAGGTTGCGAA | TAATACGACTCACTATAGTCAGTTGGCCATTTTAGTTGGT |
| 15 | *Aa-Antp* | 446 | TAATACGACTCACTATAGTGCAGACACAACCCAATGGTA | TAATACGACTCACTATAGGGCTCAAACATCCTCTCTGCT |
| 16 | *Aa-ara* | 270 | TAATACGACTCACTATAGGACAAACCGTATGCTCCTGC | TAATACGACTCACTATAGAGGATCGTGTTATAAAAAGCACTTG |
| 17 | *Aa-ct* | 428 | TAATACGACTCACTATAGCCGGATGAAGGGTTCTCGAC | TAATACGACTCACTATAGAAACGATGCTGCTGACACCT |
| 18 | *Aa-ems* | 338 | TAATACGACTCACTATAGTTCAACCGCGGATAGGATTT | TAATACGACTCACTATAGGGGTCCAGATGGAGAAGACG |
| 19 | *Aa-par-6* | 338 | TAATACGACTCACTATAGGCACCATAAGGCCACGGAAC | TAATACGACTCACTATAGGATACCGTTCACCTCGAGAACT |
| 20 | *Aa-tx* | 447 | TAATACGACTCACTATAGGTTGTACCAAAGGTTCGCCG | TAATACGACTCACTATAGCACGGGGTCATGGTACATCTAAA |
| 21 | *Aa-btl* | 375 | TAATACGACTCACTATAGGAAACAAAGATCGACGCTGG | TAATACGACTCACTATAGTTAAGGTTACCGTGCGGGG |
| 22 | *Aa-trh* | 359 | TAATACGACTCACTATAGATCGAGCTTTCTGCGTGAGG | TAATACGACTCACTATAGCTTCAACTTGGGCGCATCCT |
| 23 | *GFP* | 393 | TAATACGACTCACTATAGGCTACCCCGACCACATGAAG | TAATACGACTCACTATAGGGGTGCTCAGGTAGTGGTTG |

**Table S11. List of *A. albopictus* primers used for qPCR**.

| | Gene | Product length | Forward | Reverse |
|---|---|---|---|---|
| 1 | *Aa-RPL32* | 130 | TGGGGGCAAGCTTGTCATAG | GCCGCGTGTTGTACTCTGAT |
| 2 | *Aa-exd* | 136 | GTCTGTCTGATCCGGCGTAG | ACCATTCAAAGAGGAGGGCG |
| 3 | *Aa-* | 168 | TTGGTAGTCCTGGTGATGCG | AGGGGTGCGTTAAATGCTGA |
| 4 | *Aa-Dll* | 117 | TCAAATCATCACCTCCCGCC | CTCCCTGCTGAGGACTTGTG |
| 5 | *Aa-N* | 275 | ATTCCACTTCCGGCTTTCTGT | CGCGTTCTTGACCGGAATTT |
| 6 | *Aa-Dl* | 234 | TGTCAGCTGCTAGCATAAACCA | GCACGCATTTGCACATTGAAG |
| 7 | *Aa-Ser* | 215 | GTACATGTGTAAGTGTGGGCG | GTCTACGCGGTTCTCGCTTT |
| 8 | *Aa-wg* | 246 | TCTCCCGCTTTGGCTATTCC | AGGGTTGAGGAAACGGAAGTG |
| 9 | *Aa-Wnt10* | 190 | GCCATATTATGCGAGCAGCC | CATGTGGATGTGACCCCTCG |
| 10 | *Aa-hh* | 215 | ATTTGAACCGGGCGTTCTCT | ATGCTACGCTCTGAAACGCT |
| 11 | *Aa-en* | 208 | GGAGTTGGGACGCAGTTAGA | GCTACCCCAAACTCCACGAA |
| 12 | *Aa-ptc* | 215 | CCTTCTGCGTAGGCCTCAAA | GACCACTTCCAGGTGCGTTA |
| 13 | *Aa-ci* | 225 | TCGCACGTATACGGCTTCTC | TCACATCCACGCGAACAAGA |
| 14 | *Aa-dpp* | 237 | CGTCGTCAGCATCCCTCATT | CACTATCACTATCGCCGCCC |
| 15 | *Aa-Antp* | 185 | GACGGTCGGCCCAGAAAATA | TTCTGCTCTTCGCCTCGTTT |
| 16 | *Aa-Scr* | 169 | GGAAGGTCACTGACGACACA | ATTCCATGCTCGCAAACTGC |
| 17 | *Aa-ara* | 201 | TGTACCGATTCCTTGCTCTACA | TGCACTCTGAATCAGCCAAGT |
| 18 | *Aa-ct* | 176 | GCCATCCCCTCAAACTATGCT | GTCCGGGTTGTTGGTGGCT |
| 19 | *Aa-ems* | 210 | TTGGACGTTGACCAATTCTGC | CAACAGGAGGAGGGCAAGTC |
| 20 | *Aa-par-6* | 196 | AAATGTGGTGCTGCGGAAAC | AAGGCTATCTGGCGCTCATC |
| 21 | *Aa-tx* | 226 | GTAGTCGTCGGCAAAACCAT | ACGGGGTCATGGTACATCTAA |
| 22 | *Aa-trh* | 171 | GAACATCTCCAACCGAGCGA | TCAAGCATGGCATAGCACTG |
| 23 | *Aa-btl* | 210 | GCGAACGGGTACGGATGTAT | TTCCGTCGGTGCAGAATTGA |

**Table S12. Summary of *A. albopictus* RNAi results**.

| | Gene | no. of injected larvae | qPCR | Died before prepupa | no. individual reached prepupa | no phenotypic defects | Defect in other appendages | Defect in PROD | Larval lethal rate | Appendage defect rate |
|---|---|---|---|---|---|---|---|---|---|---|
| control | *GFP* | 128 | 40 | 25 | 63 | 63 | 0 | 0 | 28.41% | 0.00% |
| P/D axis | *Aa-Dll* | 109 | 40 | 29 | 40 | 14 | 11 | 15 | 42.03% | 65.00% |
| | *Aa-exd* | 138 | 40 | 44 | 54 | 6 | 19 | 29 | 44.90% | 88.89% |
| | *Aa-hth* | 121 | 40 | 33 | 48 | 17 | 7 | 24 | 40.74% | 64.58% |
| A/P axis | *Aa-en* | 208 | 40 | 112 | 56 | 23 | 6 | 27 | 66.67% | 58.93% |
| | *Aa-hh* | 181 | 40 | 100 | 41 | 10 | 9 | 22 | 70.92% | 75.61% |
| | *Aa-ci* | 175 | 40 | 87 | 48 | 9 | 13 | 26 | 64.44% | 81.25% |
| | *Aa-ptc* | 196 | 40 | 112 | 44 | 15 | 9 | 20 | 71.79% | 65.91% |
| | *Aa-dpp* | 140 | 40 | 50 | 50 | 18 | 10 | 22 | 50.00% | 64.00% |
| D/V axis | *Aa-N* | 201 | 40 | 113 | 48 | 7 | 12 | 29 | 70.19% | 85.42% |
| | *Aa-Dl* | 141 | 40 | 72 | 29 | 7 | 4 | 18 | 71.29% | 75.86% |
| | *Aa-Ser* | 175 | 40 | 74 | 61 | 29 | 9 | 23 | 54.81% | 52.46% |
| | *Aa-wg* | 117 | 40 | 49 | 28 | 11 | 4 | 13 | 63.64% | 60.71% |
| | *Aa-Wnt10* | 116 | 40 | 38 | 38 | 18 | 4 | 16 | 50.00% | 52.63% |
| Hox genes | *Aa-Antp* | 276 | 40 | 166 | 70 | 37 | 16 | 17 | 70.34% | 47.14% |
| | *Aa-Scr* | 180 | 40 | 106 | 34 | 14 | 5 | 15 | 75.71% | 58.82% |
| selector genes | *Aa-ara* | 154 | 40 | 81 | 33 | 13 | 6 | 14 | 71.05% | 60.61% |
| | *Aa-ems* | 125 | 40 | 39 | 46 | 46 | 0 | 0 | 45.88% | 0.00% |
| | *Aa-ct* | 221 | 40 | 95 | 86 | 48 | 10 | 28 | 52.49% | 44.19% |
| | *Aa-par-6* | 123 | 40 | 35 | 48 | 26 | 13 | 9 | 42.17% | 45.83% |
| | *Aa-tx* | 155 | 40 | 42 | 73 | 39 | 17 | 17 | 36.52% | 46.58% |
| tracheal genes | *Aa-btl* | 153 | 40 | 83 | 30 | 30 | 0 | 0 | 73.45% | 0.00% |
| | *Aa-trh* | 122 | 40 | 46 | 36 | 36 | 0 | 0 | 56.10% | 0.00% |

#### Table S13. Key Resources

| REAGENT or RESOURCE | SOURCE | IDENTIFIER |
|---|---|---|
| **Biological samples** | | |
| *Deuterophlebia wuyiensis* wholebody (larva) | This paper | N/A |
| *De. wuyiensis* PROD tissue (early larva) | This paper | N/A |
| *De. wuyiensis* PROD tissue (late larva) | This paper | N/A |
| *De. wuyiensis* wing disc (early larva) | This paper | N/A |
| *De. wuyiensis* wing disc (late larva) | This paper | N/A |
| *Aedes albopictus* PROD tissue (early larva) | This paper | N/A |
| *A. albopictus* PROD tissue (late larva) | This paper | N/A |
| *A. albopictus* wing disc (early larva) | This paper | N/A |
| *A. albopictus* wing disc (late larva) | This paper | N/A |
| *Drosophila melanogaster* PROD (84h AEL) | This paper | N/A |
| *D. melanogaster* PROD (96h AEL) | This paper | N/A |
| *D. melanogaster* PROD (108h AEL) | This paper | N/A |
| *D. melanogaster* PROD (120h AEL) | This paper | N/A |
| *D. melanogaster* wing disc (84h AEL) | This paper | N/A |
| *D. melanogaster* wing disc (96h AEL) | This paper | N/A |
| *D. melanogaster* wing disc (108h AEL) | This paper | N/A |
| *D. melanogaster* wing disc (120h AEL) | This paper | N/A |
| **Chemicals, peptides, and recomb** | | |
| Defibrinated sheep blood | Thermo Fisher Scientific | R54016 |
| Lactic acid | MilliporeSigma | V000448 |
| Triton X-100 | MilliporeSigma | T8787 |
| Paraformaldehyde (4%) | Beyotime Biotechnology | P0099 |
| Bovine serum albumin (BSA) | Beyotime Biotechnology | ST2249 |
| Normal goat serum | Beyotime Biotechnology | C0265 |
| ProLong Gold Antifade Mountant with DAPI | Thermo Fisher Scientific | P36931 |
| TRIzol reagent | Thermo Fisher Scientific | 15596018CN |
| RNA-later | Thermo Fisher Scientific | AM7021 |
| ClonExpress Ultra One Step Cloning Kit | Vazyme Biotech | C116-01 |
| Eastep Super Total RNA Extraction Kit | Promega | LS1040 |
| Eastep RT Master Mix | Promega | LS2050 |

| | | |
|---|---|---|
| ChamQ Universal SYBR qPCR Master Mix | Vazyme Biotech | Q711-02 |
| SMART-Seq Single Cell Kit | Takara | 634437 |
| Nextera DNA Flex Library Prep Kit | Illumina | 20018704 |
| TruSeq RNA and DNA Library Prep Kits v2 | Illumina | RS-122-2001 |
| T7 High Yield RNA Transcription Kit | Vazyme Biotech | TR101 |
| VAHTS RNA Clean Beads | Vazyme Biotech | N412 |
| DNase I | Thermo Fisher Scientific | 18047019 |
| Collagenase | MilliporeSigma | C0130 |
| Papain | MilliporeSigma | 10108014001 |
| Phosphate-buffered saline (PBS) | Beyotime Biotechnology | C0221A |
| MACs Tissue Storage Solution | Miltenyi Biotec | 130-100-008 |
| **Deposited data** | | |
| *De. wuyiensis* PacBio HiFi reads | This paper | SRA: PRJNA1226360 |
| *De. wuyiensis* Illumina reads | This paper | SRA: PRJNA1226360 |
| *De. wuyiensis* Hi-C data | This paper | SRA: PRJNA1226360 |
| *Philorus* sp. PacBio HiFi reads | This paper | SRA: PRJNA1226360 |
| *Philorus* sp. Illumina reads | This paper | SRA: PRJNA1226360 |
| *Philorus* sp. Hi-C data | This paper | SRA: PRJNA1226360 |
| *De. wuyiensis* tissue-specific RNA-seq | This paper | SRA: PRJNA1226360 |
| *A. albopictus* tissue-specific RNA-seq | This paper | SRA: PRJNA1226360 |
| *D. melanogaster* tissue-specific RNA-seq | This paper | SRA: PRJNA1226360 |
| *Philorus* sp. larval RNA-seq | This paper | SRA: PRJNA1226360 |
| *De. wuyiensis* genome files | This paper | Figshare: https://figshare.com/s/f2b77d343514b7dd17be |
| *Philorus* sp. genome files | This paper | Figshare: https://figshare.com/s/f2b77d343514b7dd17be |
| **Experimental models** | | |
| *Deuterophlebia wuyiensis* | Wild population | N/A |
| *Philorus* sp. | Wild population | N/A |
| *Aedes albopictus* | Laboratory strain | N/A |
| *Drosophila melanogaster* | Laboratory strain | N/A |
| **Software and algorithms** | | |
| Hifiasm v0.19.8 | Cheng et al., 2021 | https://github.com/chhylp123/hifiasm |
| Purge_dups v1.2.6 | Guan et al., 2020 | https://github.com/dfguan/purge_dups |
| Pilon v1.24 | Walker et al., 2014 | https://github.com/broadinstitute/pilon |

| | | |
|---|---|---|
| Juicer | Durand et al., 2016 | https://github.com/aidenlab/juicer |
| 3D-DNA v180922 | Dudchenko et al., 2017 | https://github.com/aidenlab/3d-dna |
| AllHIC v0.9.8 | Zhang et al., 2020 | https://github.com/tangerzhang/ALLHiC |
| Juicebox v2.20.00 | Durand et al., 2016 | https://github.com/aidenlab/Juicebox |
| BRAKER3 v3.0.8 | Gabriel et al., 2024 | https://github.com/Gaius-Augustus/BRAKER |
| PASA v2.5.3 | Haas et al., 2003 | https://github.com/PASApipeline/PASApipeline |
| HISAT2 v2.2.1 | Kim et al., 2019 | https://daehwankimlab.github.io/hisat2/ |
| StringTie v2.2.1 | Pertea et al., 2015 | https://ccb.jhu.edu/software/stringtie/ |
| GeMoMa v1.7.1 | Keilwagen et al., 2016 | https://www.jstacs.de/index.php/GeMoMa |
| BUSCO v5.0.0 | Manni et al., 2021 | https://busco.ezlab.org/ |
| Trinity v2.14 | Grabherr et al., 2011 | https://github.com/trinityrnaseq/trinityrnaseq |
| FastQC v0.12.0 | Andrews, 2010 | https://www.bioinformatics.babraham.ac.uk/projects/fastqc/ |
| Trimmomatic v0.39 | Bolger et al., 2014 | https://github.com/usadellab/Trimmomatic |
| DESeq2 v1.46.0 | Love et al., 2014 | https://bioconductor.org/packages/release/bioc/html/DESeq2.html |
| OrthoFinder v2.5.5 | Emms & Kelly, 2019 | https://github.com/davidemms/OrthoFinder |
| MAFFT v7.520 | Katoh & Standley, 2013 | https://mafft.cbrc.jp/alignment/software/ |
| MCMCtree v4.9 | Yang, 2007 | https://github.com/abacus-gene/paml |
| IQ-TREE v2.2.0 | Minh et al., 2020 | http://www.iqtree.org/ |
| Cell Ranger v6.0.0 | 10x Genomics | https://support.10xgenomics.com/single-cell-gene-expression/software/pipelines/latest/what-is-cell-ranger |
| Seurat v4 | Satija et al., 2015 | https://satijalab.org/seurat/ |
| Fiji (Fiji is just ImageJ) 1.54d | Schindelin et al., 2012 | https://imagej.net/software/fiji/ |
| CellSens Dimension 3.1.1 | Olympus | https://evidentscientific.com/en/products/software/cellsens |
| R v4.3.1 | R Core Team, 2023 | https://www.r-project.org/ |
| R Studio 2023.6.0.421 | Posit | https://posit.co/downloads/ |
| ggplot2 v3.5.1 | Wickham, 2016 | https://ggplot2.tidyverse.org/ |
| phytools v2.4-4 | Revell, 2012 | https://cran.r-project.org/web/packages/phytools/ |
